# Evidence for the extinction burst in grip force: A preregistered study

**DOI:** 10.64898/2026.01.23.701116

**Authors:** Marlon Palomino, María Cisneros-Plazola, Any Dubón, Valeria Pérez-Treviño, Gabriela E. López-Tolsa, Rodrigo Sosa

## Abstract

In behavioral science, “extinction” refers to the change that results when a previously rewarded pursuit no longer yields reward; accordingly, this phenomenon is central to behavioral reorganization across species. Although one might expect a monotonic transition from engagement to disengagement in goal-directed behavior, in mammals this transition is often non-monotonic whereby behavior can transiently intensify before ceasing to occur—a phenomenon known as the “extinction burst.” Detecting this effect requires a metric and time window that capture the sometimes short-lived performance increase and enable comparison to counterfactual continued-reward trajectories. Certifying an increase as genuine behavioral intensification further requires ruling out increases attributable to release from competition with reward-consummatory activities, as well as inflation of peak values by extinction-induced variability. We preregistered a study in which participants earned money by squeezing a hand dynamometer. Peak standardized force in short, time-matched blocks was pre-specified as the primary index; analyses contrasted before-to-after change at different tiers where extinction was present versus absent. Peak force increased upon extinction onset, yielding a large and robust stage-by-treatment interaction effect. Our symbolic reward regime precludes release-from-competition explanations. The continuous-force record further shows a genuine central-tendency increase, ruling out variance inflation as the source of the peak-force effect.

## 1 Introduction

Goal-directed behavior is thought to transiently intensify when mammals can no longer attain their primary goals[1, 2, 3, 4]. This phenomenon is termed the *extinction burst* [5]. In behavioral science, extinction can refer to situations in which a reward that previously followed a behavior is discontinued even though the behavior continues to occur. Many everyday situations illustrate this transient intensification. For example, a child who does not get what they requested may throw a tantrum. Or someone may press the button on a broken coffee machine harder. Similarly, when we tap the “see more” area on a touch device and nothing happens, we may find ourselves tapping faster and/or harder. Together, these examples show that burst-like phenomena are common in daily life.

Intuitively, one plausible function is that bursts serve as a probe when reward suddenly fails to occur. A brief escalation in responding may help determine whether the environment has genuinely changed or whether recent nonreward reflects incidental noise. For example, a prior attempt may have failed because effort was inadvertently insufficient[6]. This escalation can then give way to disengagement if reward remains withheld despite subsequent attempts at higher intensity[7].

A relevant question is which aspect of behavior intensifies during an extinction burst: its rate, its vigor, its duration, its variability, or some combination of these features. Historically, extinction bursts were first described in laboratory studies using rats in operant conditioning chambers, where the target response is typically lever-pressing and the reward is a small food pellet[8]. In those early studies, measurement was largely limited to how many responses occurred per unit of time; that is, response rate. Under such constraints, extinction bursts appeared as a transient increase in how closely spaced lever presses were. As a result, the concept of the extinction burst became somewhat anchored to response rate[9, 5], partly because of instrumentation limits (responses recorded as time-stamped events) and partly because of disciplinary tradition.

In recent years, the validity, or at least the generality, of extinction-burst phenomena has been called into question[10]. One account proposes that in many laboratory preparations, goal-directed behavior is not a single, isolated action but a sequence that alternates between the target response and consummatory activities (e.g., reward collection and intake). When extinction is introduced (i.e., the reward is withheld), consummatory activities collapse and arguably free up time that is then reallocated to the target response[11, 12, 13]. Accordingly, the observed increase has been argued to reflect an increased prevalence of the target response simply because it is released from competition with those reward-related activities[14]. This line of reasoning leads to the conclusion that what looks like an extinction burst does not necessarily reflect a genuine intensification of goal-directed action but could instead represent a measurement artifact. Concretely, the burst-like effect might vanish should analyses adequately control for the confounding role played by the collapse of consummatory activities.

### 1.1 Behavioral intensification during extinction beyond response rate

The extinction burst has been considered of prime relevance for interventions in behavioral management; namely, an increase in a problem behavior when trying to mitigate that behavior may hinder the management protocol itself. However, there is scant laboratory research to provide foundations for behavioral interventions[11]. A number of things that we know about the extinction burst come from studies conducted several decades ago. For example, Notterman[15] trained rats to press a lever with a specified minimum force to earn a food reward for 4 sessions; next, he applied an extinction regime and observed a clear increase in both peak force and force variability at the group level. In a similar vein, Margulies[16] trained rats to press a lever to earn a food reward and he observed an increase in both lever-press duration and duration variability. Note that increases in force such as those reported by Notterman cannot be explained by release-from-competition accounts, which predict that the target behavior increases in frequency when there are no consummatory activities competing for that time. On the other hand, increases in duration can be readily accommodated by a release-from-competition account; concretely, the collapse of consummatory activities in extinction might leave more time to execute longer lever-presses in Margulies’ study.

In a more recent study, Iversen[17] trained rats to push a hanging pole sideways to earn a reward; in this study the durations of both pole displacements and food-tray visits were recorded. In addition, a snapshot of each rat was taken every time the criterion for reward was met. Iversen found that both the average and standard deviation of pole-movement durations increased, consistent with Margulies’ study. The snapshots revealed that the more reward training rats received, the more they tended to exhibit stereotypical postures—involving only slight discrepancies between one episode and the next. Interestingly, upon extinction onset, subjects’ posture started to vary; for example, rats approached the pole from different sides or touched it with either one or two hands across episodes. At this point it is clear that nonhuman animals increase both the average and the variability of their response durations for two different target behaviors (lever pressing and pole displacement). The single study by Notterman suggests that such increases in average and variability also occur in response force. But what is shared among these three studies is an increase in variation, occurring both in dimensions (force and duration) of the target behavior itself and in other aspects of behavior like posture. Note that variation could play a role in extinction bursts different from that of duration and force; we will turn to that issue in the next subsection.

Another, yet more recent, relevant study is one by Alessandri and Lattal[18] focusing on response-force increases upon extinction onset with humans. Their behavioral task involved pressing a force sensor with the thumb to earn a reward when a criterion response was attained, and then observing variations in force once the reward was discontinued. Specifically, these authors compared the maximum force during reward against the average force in time bins of varying length during extinction. Whether an increase in force during extinction relative to reward was observed depended on the duration of the bins over which data were aggregated. Across three experiments, 100% of human participants showed an increase in pressing force when indexing average force during extinction in one-second bins. Weighted by the number of participants per experiment, about 71% of participants showed an increase in average force when using 10-second bins and 67% when using 30-second bins, both against the baseline peak, whereas only about 38% did when using 100-second bins where the benchmark was the baseline mean rather than the peak. Alessandri and Lattal[18] concluded that the increase in force they observed upon extinction onset was not systematic, consistent, or reliable, challenging the generality of the extinction burst.

Because there is no principled expectation as to how long sustained force in extinction should surpass performance during reward, we consider that any increase should constitute an extinction burst. By that criterion, the 100%-of-participants-across-three-experiments result in Alessandri and Lattal’s[18] study could be regarded as outstandingly robust support for the extinction burst; this finding would be even more impressive considering that they compared maximum force pre-extinction versus average force post-extinction, which is biased against detecting pre-post increases. There are, however, some considerations that preclude an unambiguous interpretation of the extant evidence for response-force increases upon extinction as bursts.

### 1.2 Limitations of extant evidence for extinction-induced behavioral intensification

In the previous subsections, we explained how increases in response rate during extinction can be fully accounted for by release from competition; accordingly, by themselves, they do not constitute evidence of genuine behavioral intensification, since they may instead reflect the reallocation of an already-available response into newly freed time. We went on to describe the extant evidence of behavioral increases during extinction across non-rate behavioral dimensions; now, we will examine how each of them contributes to the classic view of the extinction burst. Although increases in response duration do not imply merely more instances of the same response, they can still be accommodated by release-from-competition accounts. That points to response force increases as a clearer way to index behavioral intensification that cannot be attributed to release from competition. Hence, evidence for the extinction burst would be restricted to Notterman’s [15] and Alessandri and Lattal’s [18] studies. A feature these studies share with the others described above[16] [17]is that they lack inferential statistical treatment of their data. All the studies described above relied on summary statistics, visual inspection of the distribution or progression of measurements, and event-occurrence records in the form of timestamps. Quantitative research questions require inferential statistics to derive conclusions accompanied by estimates and their confidence.

A further limitation of the studies described above (and of most extinction-burst studies) is that they employed a quasi-experimental study design. Concretely, they feature simple baseline-treatment comparisons where the baseline is the reward stage and the treatment is extinction. Cook and Campbell[19] classified this design, which they called the one-group pretest-posttest design, among those that often do not permit reasonable causal inferences. According to their taxonomy of study designs, a true experiment, or experiment proper, would require a control group with a parallel rewarded condition while the experimental group would experience extinction; namely, such a condition would serve as a counterfactual comparison that indexes what would have happened had reward not been withheld.

This counterfactual observation would distinguish a mere increase from an extinction burst. An increase is a descriptive fact, namely, that force was higher in extinction than in reward. In contrast, a burst is a causal claim that the increase is attributable to procedural extinction. Without a continued-reward control, the studies above can establish the former but not the latter. One instance of the benefits of including such a control is that it enables indexing false-alarm rates. For example, while we considered the 100%-of-participants-across-three-experiments result in Alessandri and Lattal’s[18] study as clear evidence of an increase from reward to extinction, this conclusion would be better calibrated against what would have happened had extinction not been introduced. Assuming stable pre-extinction performance, a control condition would reveal the rate at which an increase arises by chance between two successive equivalent samples. Only against that rate can an increase be judged a genuine burst rather than a false alarm. On a related point about comparisons, what is compared across the pre- and post-extinction stages should also be equivalent—whether averages or maximum values, and whether across entire sessions or matched time windows.

One more concern is that a study’s conclusions can depend heavily on which subset of the results is emphasized. Alessandri and Lattal[18] aimed to compare several criteria for defining a burst but did not establish beforehand which one should settle their conclusion. A principled choice would be the criterion that corresponds to their own definition, a transient increase, which their short-bin analysis satisfies. Instead, they based their conclusion on the longer bins, whereas in the shorter ones every participant showed the increase. Under the stringent criteria, even the moderate positive cases at intermediate bins, around 70% of participants, were judged insufficient. This imposes a minimum-duration requirement that their adopted definition does not imply, since a transient increase need not persist to qualify, so the short-bin effect still qualifies as a burst. Choosing conservative criteria is often a defensible path; however, when several analyses are reported without designating a primary one beforehand, a reader cannot determine whether the criteria were chosen independently of the results. Committing to a burst criterion in advance would make it possible to verify that the criteria were fixed independently of the results.

Establishing extinction-burst criteria in advance is not as straightforward as one might assume. The literature on extinction bursts has increasingly expressed concerns about how this phenomenon should be measured and, more fundamentally, what should count as evidence of a burst[9, 20]. In particular, because the effect is often transient, specifying in advance the temporal window over which to track it can be difficult[20, 14, 21, 22]. This issue complicates experimental design and analysis, including decisions about how to detect and quantify burst-like changes.

The methodological concern just described stems from the putatively non-monotonic trajectory that engagement would follow in a reward-to-extinction transition that included an extinction burst. In an overly simplified view of goal-directed behavior, one might expect that when a reward is contingent on a response, organisms engage in reward pursuit, and when the reward is discontinued, they eventually disengage[13]. This transition can be idealized as a step function. Alternatively, allowing for more realism, disengagement might follow a gradual decline, as many canonical reinforcement-learning models predict[23]. By contrast, if extinction bursts are genuine, the relevant behavioral index would increase first, becoming progressively harder to detect as the response itself ceases to occur. This putative non-monotonic nature of the extinction process warrants an approach that goes beyond point-estimate summary statistics such as rates or averages to detect extinction bursts.

If summary measures are computed over a time window that includes both the rise and the subsequent decline in behavioral intensity, they will tend to wash out the very pattern that defines a burst. A straightforward solution, therefore, is to quantify peak intensity (i.e., the maximum value of the relevant behavioral index) rather than relying solely on window-averaged summaries when evaluating whether an extinction burst occurred in a given situation. However, quantifying peak intensity carries a cost arising from the variation dimension.

Our survey of the literature has pointed to increases in variation as a well-established outcome of procedural extinction. Unlike behavioral dimensions such as rate, duration, and force, increased variation could be considered at odds with the conceptualization of bursts as an intensification of the behavior that has attained a certain goal. Concretely, variation can occur without intensification. Moreover, variation might create the appearance of intensification even when behavior is trending toward attenuation. A behavioral index may vary only slightly while the target action remains effective, but once that action becomes ineffective, the index might begin to vary more, producing peaks but also equivalent downward swings, so that the net value remains unchanged (or may even be generally decreasing). It is therefore entirely plausible that high-intensity behaviors appear more often simply because behavior is varying more. This possibility would threaten the interpretation of an increase in the peak of any behavioral index as evidence of genuine intensification.

### 1.3 The present study

The purpose of the present study was to assess the extinction burst, addressing several methodological concerns raised in the subsection above. Of primary importance, we aimed to rule out release-from-competition and sheer-variation confounds. We addressed these confounds because an observed increase in behavior could reflect behavioral redistribution or variance-driven measurement artifacts, rather than a genuine increase in the behavior-generating process. Next, we describe how our study addresses these methodological and theoretical issues.

- We selected force as our dimension of interest to index behavioral intensification because rate increases are confounded by release from competition when extinction is introduced, and because duration increases remain vulnerable to the same account.
- We targeted humans as our model species, given that monetary rewards can be delivered symbolically to them. Importantly, symbolic rewards do not entail a post-reward consummatory behavior explicitly competing with the target response. Hence, any extinction burst observed following the discontinuation of such symbolic rewards cannot be explained by release from competition.
- We preregistered our study to lock in our definitional, methodological, measurement, and inferential decisions in advance, preventing their post-hoc flexibility.
- We based our conclusions on conventional statistical inference benchmarks with consideration of statistical power, rather than relying on descriptive statistics.
- We compared maximum values, rather than averages, of our measurements of interest to avoid possible washouts when observation windows include both the rising and the declining facets of the extinction process.
- Because greater variation alone can increase maximum force, a higher peak need not reflect any true intensification. We therefore included an analysis of continuous force that checks whether, at some point, there is a reliable net increase in this quantity after extinction. This analysis complements the peak-increase criterion rather than replacing it and addresses variance-driven peak inflation.
- We compared reward-to-extinction transitions against matched reward-to-reward transitions so that our analysis isolates the magnitude of increases attributable to extinction. The former captures increases observed upon extinction, whereas the latter indexes those arising in the counterfactual condition where extinction is not introduced.

Altogether, these decisions address the release-from-competition and sheer-variation confounds identified above, while the preregistration renders verifiable what we considered methodological challenges in advance.

## 2 Methods

The study protocol was reviewed and approved by the Associate Dean for Research at Universidad Panamericana (Campus Guadalajara), following evaluation of compliance with applicable scientific, technical, and ethical standards (approval issued 26 September 2025). The power-analysis script, raw data, full statistical analysis pipeline, complete participant instructions, and the task script are available in an online repository at https://github.com/RSosa-BehavioralScience/extinction_burst_force, hereafter referred to as the Electronic Supplementary Materials.

### 2.1 Sample

A total of 76 participants were recruited at the university campus. Eligible individuals were currently enrolled students. The final analyzed sample comprised 63 participants, 61 of whom were right-handed and 2 left-handed; 29 self-identified as male and 34 as female. Ages ranged from 18 to 23 (M = 19.5, SD = 1.72) years. Participants were assigned to one of four conditions; both participants and researchers were blinded to condition assignment prior to session start. Condition allocation was computer-generated and concealed within the task software. After providing informed consent, each participant completed the full procedure in a single laboratory session. Participants received compensation at a fixed per-reward rate of approximately £0.20, paid at the end of the session.

Participants were excluded under the following criteria: calibration failure (mean force across two calibration attempts *<* 10 N; see Behavioral task), non-completion (failure to complete all task-defined rewards for the assigned condition within 10 min), protocol violation (use of both hands at any time, dropping or unclasping the dynamometer, or standing before the session ended; see Assessment protocol), and reward-cue delivery failure attributable to a hardware timing limitation in the detection algorithm (details in Behavioral task). Among otherwise eligible participants, those who produced data points outside the valid hardware range were excluded after data collection.

To determine the sample size, we conducted a simulation-based power analysis tailored to our design. We aimed to detect a standardized interaction effect of 0.55 for the stage × treatment term, assuming a block effect of −0.30, participant ICC = 0.30, and one-sided *α* = 0.05 with 80% power. Because standard power-analysis tools cannot accommodate our multiple-baseline structure, we generated 10,000 simulated datasets and fitted mixed-effects models to each (for details, see Study design). With 16 participants per group (64 total), the simulations indicated a power of 0.84 to detect a median standardized interaction effect of 0.53 and a block effect of approximately −0.28, confirming that our planned sample size was sufficient.

Data collection continued until the planned target of 64 analyzable participants was reached. Participants who met prespecified exclusion criteria during ongoing data collection were excluded and replaced to maintain the planned group sizes. After recruitment closed, one additional participant was excluded for out-of-range force values identified during final data-quality checks and could not be replaced, yielding a final analyzed sample of 63 participants (group sizes reported in Study Design). In total, 13 participants met exclusion criteria: 5 due to recording artifacts, 5 for not completing all available rewarded trials within the time limit, 1 for out-of-range force values identified during final quality checks, 1 for touching a non-permitted part of the dynamometer, and 1 for turning around during the task. Of these, 12 were excluded and replaced during ongoing data collection, and 1 was excluded after recruitment closed and not replaced.

### 2.2 Materials

#### 2.2.1 Hardware

Grip responses were recorded using a Vernier Go Direct® hand dynamometer (see Figure 1a) connected via USB to a laptop. Grip-force recordings had a temporal resolution of approximately 10 Hz.

**Figure 1:**
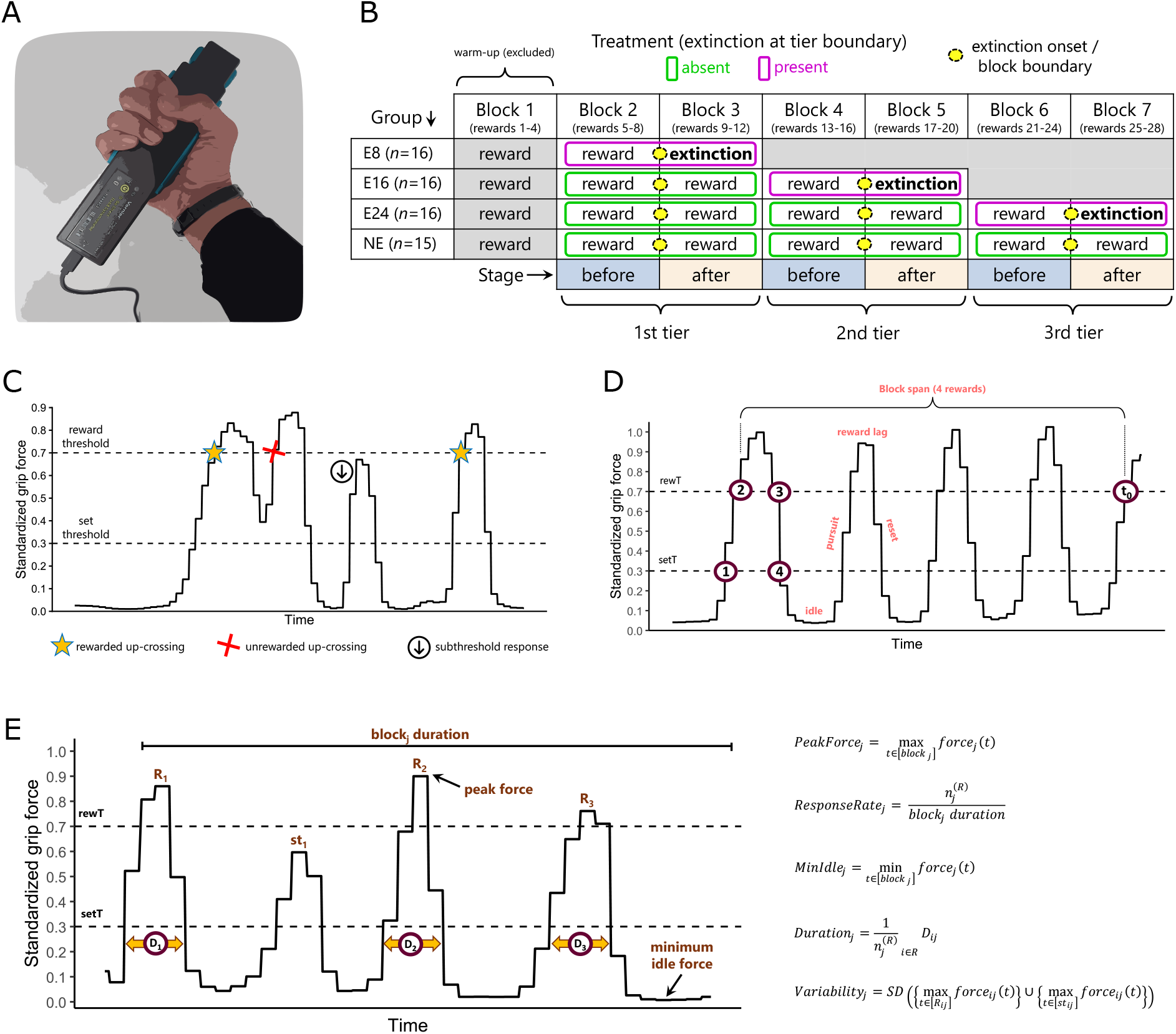
Study overview and operational definitions. (a) Hand dynamometer used to record grip force. (b) Multiple-baseline design showing reward and extinction tiers and the before/after stage coding around each tier boundary. (c) Reward criterion defined by set and reward thresholds, with examples of rewarded, unrewarded, and subthreshold responses; stars mark the exact moment of reward delivery. (d) Schematic of reward cycles and four-reward blocks. (e) Visual schematic (left) and formal definitions (right) of our blockwise behavioral metrics.

#### 2.2.2 Software

The task was programmed and presented in PsychoPy (v2024.2.4; Python 3.10)[24]. The full PsychoPy script and associated files needed to reproduce the task are provided in the Electronic Supplementary Material.

### 2.3 Study design

We used a multiple-baseline across groups design with a staggered introduction of extinction onset (after 8, 16, or 24 rewards), plus a no-extinction control (group NE). This design enhances causal inference by assessing whether the effect of interest replicates at different positions[25] in the protocol, thereby controlling for confounds such as elapsed time and reward experience. Figure 1b provides an overview of the study design.

Staggering extinction onset across groups serves two purposes. First, it separates the effect of extinction from the position at which extinction is introduced. An effect observed at a single moment cannot be confidently attributed to the manipulation, because the moment of application is confounded with everything else occurring at that moment. Introducing extinction after 8, 16, or 24 rewards yields an estimate of its effect that is pooled across positions and therefore not tied to any one of them. The resulting parameter is a net effect of extinction, aggregating information from all three onset positions. This is the rationale underlying the use of multiple-baseline designs in intervention research[25].

Second, the design is economical. Conducting three separate experiments with different onset positions would require a separate control group in each, with each control group’s data serving only its corresponding experiment. Here, a single no-extinction group provides reward→reward reference transitions for all three onset positions. Because participants were randomly assigned to conditions and run in parallel, this group contributes a counterfactual drawn from the same testing period as every extinction group, rather than from a separate experiment conducted at a different time. The later-onset groups further contribute such transitions during their pre-extinction blocks, and given that condition assignment was concealed, these blocks were indistinguishable in participants’ experience from the corresponding blocks in the no-extinction group. The design therefore optimizes the ratio of information obtained to logistical and monetary cost.

The data unit of interest was the maximum force recorded during the time span covering blocks of four rewarded responses or during which extinction was implemented. Groups E8, E16, and E24 completed 8, 16, and 24 rewarded responses, respectively (2, 4, and 6 rewarded blocks), with extinction onset immediately following the final rewarded block in each case. Group NE completed 28 rewarded responses (7 blocks) with no extinction.

For groups in which extinction was implemented (groups E8, E16, and E24), responses after extinction onset were recorded over a time window whose duration equaled the time the participant took to complete the last block before extinction. This time-matching approach enables fair comparisons by providing an equal opportunity to capture responses before and after extinction while maximizing temporal contiguity.

By design, the first block in all groups was treated as a warm-up period and thus excluded from the analysis. We paired the remaining blocks into three tiers, each consisting of two consecutive blocks that straddle a boundary at which a transition to extinction could occur in the extinction groups. Within each tier, the earlier block was coded as the *before* stage and the later block as the *after* stage. Even blocks (2, 4, 6) were therefore coded as before, and odd blocks (3, 5, 7) as after. This indexing aligned blocks around homologous boundaries for comparisons of interest, where the within-tier boundary between blocks 2↔3, 4 ↔5, 6 ↔7 includes the extinction onset for groups E8, E16, and E24, respectively (see Figure 1b).

We coded whether extinction onset was implemented within a given before–after tier as a factor with two levels: absent and present. Tiers containing extinction onset were labeled present; all others were labeled absent, representing the counterfactual scenario had the reward not been withheld. Only the last before–after tier in groups E8, E16, and E24 was labeled as present, whereas all earlier tiers in those groups—and all tiers in group NE—were labeled as absent, serving as history-matched comparisons under continued reward. A compact visual summary including block structure, tiering, boundaries, and stage/treatment coding is given in figure 1b.

### 2.4 Assessment protocol

#### 2.4.1 Participant instructions

After routine preparatory steps, participants received standardized verbal and on-screen instructions (see Electronic Supplementary Materials). The key instruction was to “discover” how to obtain rewards and to earn as many rewards as possible using only the dynamometer with their dominant hand. To discourage bimanual responding, participants held a tennis ball in the non-dominant hand throughout the session. Protocol violations (e.g., using both hands, unclasping the device, standing, or turning) triggered exclusion as prespecified. Once the experiment proper began, the computer screen displayed the following message for the remainder of the session: “Ready! Now try to earn as much money as possible! — Do not get up from your chair or release the dynamometer until the screen changes color, or you will not receive any money.” The color change mentioned in the message occurred only once, marking the end of the session, and was followed by payment of the amount of money earned by each participant.

#### 2.4.2 Physical space

All sessions were conducted individually in a quiet testing cubicle (approximately 2 × 1 m) with standard office lighting and a small table and two chairs facing each other. The cubicle door remained closed during sessions; it was transparent, allowing the researcher to monitor protocol adherence from outside. Participants were seated facing away from the door.

### 2.5 Behavioral task

#### 2.5.1 Calibration and standardization

Our primary outcome variable was the standardized peak force within each analyzed block. To enable standardization, we first calibrated each participant’s grip force. At the start of the behavioral protocol, participants were asked to perform two “very strong grips” (with a 5 s rest in between). We recorded the peak force from each attempt (*F1* and *F2*) and computed their average: *avgF* = (*F1* + *F2*)/2. All subsequent force measurements were normalized by this value: *stdF* (t) = *rawF* (t)/*avgF*, where *rawF* (*t*) is the raw force detected by the dynamometer at time *t*.

#### 2.5.2 Threshold definition and reward criterion

Participants were not provided with real-time grip-force traces on the computer screen; rewards appeared abruptly when the reward criterion was met. Rewards depended on two thresholds defined relative to each participant’s calibrated force (see Figure 1c). The set threshold was *setT* = 0.30 × *avgF* and the reward threshold was *rewT* = 0.70 × *avgF*. To earn a reward, force had to cross these two thresholds upward in succession: first an upcrossing of *setT*, then an upcrossing of *rewT*, at which point the reward was delivered immediately. After each reward, force had to fall back below *setT* before a subsequent upcrossing of *rewT* became eligible; this reset requirement prevented double counting when force oscillated around *rewT* (see Figure 1c). Rewards consisted of a coin image displayed for 0.5 s, accompanied by a brief tone. Total earnings were paid after the behavioral protocol.

During a reward event participants could go on making responses; the timer for each symbolic reward restarted whenever a new criterion was met while the reward event was already in progress, effectively extending its display (i.e., the reward event was programmed to be a retriggerable event). This was experienced as a glitch or hiccup when two rewards were attained in close succession. By design, participants were not able to tell how many rewards they had attained at any given time because there was no explicit on-screen reward counter. However, if participants kept track, they could infer the count from the number of times the reward event was triggered, since even when rewards came in close succession each retrigger interrupted the ongoing display.

#### 2.5.3 Segment classification and definition of reward cycles

Reward cycles were vital to our analyses because they allowed us to track the measurements of interest (peak force within blocks) unequivocally (see Figure 1d). We defined four event types, (1) upcrossing of *setT*; (2) upcrossing of *rewT*; (3) downcrossing of *rewT*; (4) downcrossing of *setT*. These events partition the trace into segments, which we name and interpret as follows:

(1→2) Pursuit. Interval between an upcrossing of *setT* and an upcrossing of *rewT*; represents the active build-up of force leading to reward attainment.

(2→3) Reward lag. Interval between reward delivery and force dropping below *rewT*; reflects continued exertion of force after reward, likely due to perceptual/motor lag (e.g., ballistic overshoot).

(3→4) Reset. Interval between force dropping below *rewT* and crossing *setT* downward; marks the release of force needed to re-establish eligibility for a *rewT* upcrossing to attain a reward.

(4→1) Idle. Interval between falling below *setT* and the next upcrossing of *setT*; represents transient disengagement before a new pursuit, possibly reflecting resting time.

Once all segments of a session were classified, we could define a reward cycle. A reward cycle spans from one reward delivery to the next. Formally, each cycle *j* runs from event 2 of cycle *j* to event 2 of the next cycle (*j* +1). Thus, a reward cycle is constituted by the sum of the following segments: reward lag [2 → 3]_*j*_ + reset [3 → 4]_*j*_ + idle [4 → 1]_*j*_ + pursuit [1 → 2]_*j*+1_.

#### 2.5.4 Matching windows of last rewarded block and extinction block

In tiers coded as treatment = present (those that straddle the last rewarded block and the extinction block), the boundary between its constituent blocks can be defined as extinction onset, *t*_0_. Operationally, *t*_0_ marks the timestamp when a reward that would have been delivered according to the reward criterion is first withheld. The duration of the extinction block was set equal to that of its predecessor: the total span of the last four rewarded cycles ending before *t*_0_ (see figure 1d). Thus, the blocks around *t*_0_ were time-matched. We also recorded responses during the period immediately following the extinction block—specifically, the period spanning from one window length after *t*_0_ to two window lengths after *t*_0_—for exploratory analyses of extinction progression; hereafter, we refer to this period as *late extinction*. Where no extinction onset occurred (reward→reward tiers), blocks were not time-matched; instead, we used number-of-rewards matching (blocks defined as bundles of four rewarded responses).

### 2.6 Preregistration

#### 2.6.1 Access and scope

The study was preregistered on OSF prior to data collection: https://osf.io/cn5tq/overview. Our preregistration specified the design, thresholds, and block construction around *t*_0_, exclusion criteria, the primary outcome variable of interest (standardized peak force per block), the confirmatory hypothesis, and exploratory assessments.

The confirmatory analysis plan prespecified a mixed-effects model with participant-level random effects and fixed effects for stage, treatment, and their interaction, with block entered as a covariate (see Statistical analyses). The post-onset window was limited to the time-matched extinction block; analyses beyond this window were designated exploratory. Any departures from the preregistration are described below.

#### 2.6.2 Deviations

Although exclusion criteria were prespecified, one exclusion (out-of-range force values) was identified during final quality checks after recruitment had ended; as a result, the final analyzed sample was 63 rather than the planned 64, and group sizes were not perfectly balanced. This late exclusion did not alter the confirmatory inference: effect estimates and confidence intervals remained in the same range. Moreover, our simulation-based power analysis indicated that reducing the sample size to 60 would still exceed the preregistered 80% power criterion, so the final sample size remained within the planned evidential margin.

Several exploratory analyses were modified or deferred. (1) For response rate, we implemented only the primary operationalization (responses defined as complete reward cycles) and did not implement the alternative “threshold upcrossing” definition. (2) For response duration, we operationalized duration as the full behavioral episode from set-threshold upcrossing to subsequent down-crossing, rather than isolating the reward-lag component within a cycle. (3) The planned experience-modulation analysis (stage × treatment × block interaction) was not conducted for the present submission and is reserved for future work. Finally, we added two additional exploratory outcome variables, minimum idle force and force variability, which are defined in the subsection below.

### 2.7 Behavioral metrics

Figure 1e provides a schematic and formal definitions of the blockwise behavioral metrics described below.

#### 2.7.1 Peak force

Our main confirmatory hypothesis concerned the maximum force applied by participants within blocks. Peak force was obtained by indexing force samples from the beginning to the end of each block and extracting the maximum value.

#### 2.7.2 Response rate

We included response rate because it is the main outcome variable in extinction-burst studies using free-flowing (uninterrupted) behavioral tasks, referred to in the animal conditioning literature as a free-operant procedure; it was computed by counting criterion responses within a block and dividing by block duration.

#### 2.7.3 Minimum Idle Force

As a measure of reward disengagement associated with extinction, we extracted the minimum force applied during all idle states within a given block.

#### 2.7.4 Duration

As an alternative measure of behavioral intensification, we report the duration of criterion responses. Specifically, we measured the time from a set-threshold upcrossing to the subsequent set-threshold down-crossing, indexing the full pursuit-to-reset interval.

#### 2.7.5 Variability

Finally, we quantified force variability during goal-directed responses, including both successful trials (meeting criterion) and unsuccessful ones (failing to reach the reward threshold). Force variability under reward omission is critical because it typically increases the likelihood that alternative response patterns will produce rewards through novel means.

### 2.8 Data analysis

All statistical analyses were conducted in R (v4.3.0; 2023-04-21). We used linear mixed-effects regression models for statistical inference.

#### 2.8.1 Tests for discrete metrics

The focal research question was whether extinction altered force-related performance metrics relative to matched pre-onset blocks, compared with an otherwise equivalent reward → reward condition. Accordingly, the effect of interest was the stage × treatment interaction. Rather than comparing before-after transitions within the extinction condition alone, this interaction tests whether the pre-to-post change across the onset boundary differs between tiers in which extinction is present and tiers in which it is absent, that is, whether the pre-to-post shift in reward → extinction tiers exceeds the corresponding shift in matched reward → reward tiers (see Figure 1b). For each behavioral metric, we fitted a separate model of the form:

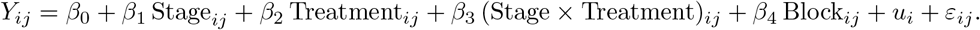

Here, *Y*_*ij*_ denotes the outcome for participant *i* in block *j*. Stage indicates whether the block occurs before versus after the onset boundary (or the homologous boundary in reward →reward tiers). Treatment indicates whether the tier includes extinction (present) or not (absent). Block was included as a numeric covariate to isolate secular change across the task. This covariate controls for gradual within-session changes unrelated to extinction (e.g., threshold-hunting adjustments, fatigue), ensuring that the stage × treatment interaction captures extinction-specific effects rather than general time-on-task trends. The participant-specific random intercept *u*_*i*_ accounts for between-participant differences in overall performance level, and *ε*_*ij*_ is the residual error term. The coefficient *β*_3_ (stage × treatment) was the parameter of interest across our metrics. For the preregistered primary outcome (peak force), we additionally report the estimated block effect and the participant-level intraclass correlation coefficient; for the remaining outcomes, we report only the standardized stage × treatment coefficient.

Because each participant can contribute multiple eligible tiers, the model pools these within-participant contrasts to yield a single interaction effect estimate. The confirmatory parameter was the stage × treatment interaction for peak force, tested directionally for a positive value. For the remaining behavioral metrics, we report the interaction parameter with two-tailed *p*-values as exploratory assessments.

We examined residuals and found that 4 of the 5 outcome variables showed departures from Gaussian assumptions, with leptokurtosis being the most prominent deviation. To address this issue while retaining the preregistered linear mixed-effects specification for effect-size calculation, we based inference on a participant-level cluster bootstrap so as to preserve the repeated-measures structure of the data[26]. Confidence intervals and hypothesis tests for the interaction of interest were obtained on a studentized bootstrap statistic[27].

#### 2.8.2 Trajectories of discrete metrics

As an additional exploratory line of inquiry, we examined how behavioral metrics evolved across extinction relative to pre-extinction reference anchors. We did so using two complementary approaches: a time-aligned analysis and an episode-aligned analysis. For the time-aligned analysis, we refit the same model but used the late extinction block, rather than the preregistered time-matched window immediately following a rewarded block. The late extinction block followed the early extinction block directly and was included to assess how the metrics progressed across extinction in equivalent time bins. However, because the number of behavioral episodes can differ between early and late extinction windows, metrics that depend on extreme values (e.g., maxima or minima) can be confounded by response density: fewer episodes reduce the likelihood of observing extreme values, which can bias apparent change across time bins.

The episode-aligned analysis was designed to complement this limitation by indexing metrics by episode number relative to extinction onset, thereby equating the number of behavioral opportunities across compared positions. Specifically, episode-wise metrics were expressed as deviations from the reference episode immediately preceding extinction onset. This episode-aligned depiction circumvents the response-density confound affecting extrema-based metrics in the time-aligned analysis, and it is presented for visual inspection only. Since later episode indices are supported by fewer participant–tier instances, the episode-aligned depiction can introduce compositional differences across episode positions; this limitation is correspondingly mitigated by the time-aligned analysis, which relies on fixed-duration bins that include information for all participants.

#### 2.8.3 Continuous-force dynamics

As a preregistered exploratory analysis, we examined mean force continuously across the reward-to-extinction transition to capture the dynamics of exerted force, which no single discrete metric reflects on its own. By indexing continuous force, we aimed to combine the dimensions captured separately by the discrete metrics into a single measure of force exerted over time, expressed relative to each participant’s own pre-extinction block (see below). Accordingly, increases in exerted force might result from higher peaks, more sustained behavioral episodes, or more closely spaced instances of either criterion or subthreshold behaviors. A net increase in continuous force upon extinction onset above that observed in the control group would indicate a rise in general force exertion attributable to reward discontinuation—regardless of which metric change underlies it within each participant.

Raw force time series were smoothed using a sliding-window average with a span equal to 25% of the pre-extinction block duration; that is, one quarter of the block capturing exactly four response cycles by design, so the window spans approximately one complete cycle on average. This window length smooths within-cycle gross oscillations while preserving between-cycle dynamics.

The time axis was normalized to place the onset of extinction at *x* = 0 and the start of the pre-extinction block at *x* = −1. This approach allows comparing pre-extinction blocks of different durations by expressing time across transitions as a proportion of the participant’s own pre-extinction block duration. Because pre-extinction and extinction blocks are time-matched (see Behavioral task subsection), the end of early extinction naturally fell at *x* = 1, and the end of late extinction at *x* = 2. The rationale is to establish the response cycle rather than clock time as the behaviorally meaningful unit to align behavioral landmarks across participants.

Force was re-standardized within each transition so that *y* = 0 corresponds to the mean of the last window falling entirely within the pre-extinction block. This equates the pre-onset force level across individuals, making deviations from zero interpretable as proportional increases or decreases relative to that reference. Transformations of the *x* and *y* axes visually funnel all participants’ performance at the same point just prior to extinction onset. The same procedure was applied to reward →reward transitions from the NE group as a counterfactual reference, and 68% bootstrap confidence intervals were derived to express uncertainty around the group mean trajectory.

## 3 Results

### 3.1 Extinction increases peak force relative to continued reward

Figure 2 shows exerted force across the session for each participant, separating blocks before versus after the onset boundary for reward →extinction tiers and reward →reward tiers. The stage × treatment interaction was positive (*β* = 0.25, 95% CI [0.15, 0.36]) and statistically significant (one-sided *p* = 9.99 × 10^−5^), supporting our preregistered directional hypothesis. This interaction suggests that peak force effectively increased following extinction onset compared to matched reward-only blocks (by about 25% of standardized force). Figure 3a confirms this pattern across all groups that included extinction-present tiers, supporting the idea that the change is linked to extinction onset rather than to a specific number of rewards obtained or time elapsed. The standardized interaction coefficient was 0.69 (95% CI [0.43, 1.00]), indicating a strong contrast between reward →extinction and reward →reward tiers.

**Figure 2:**
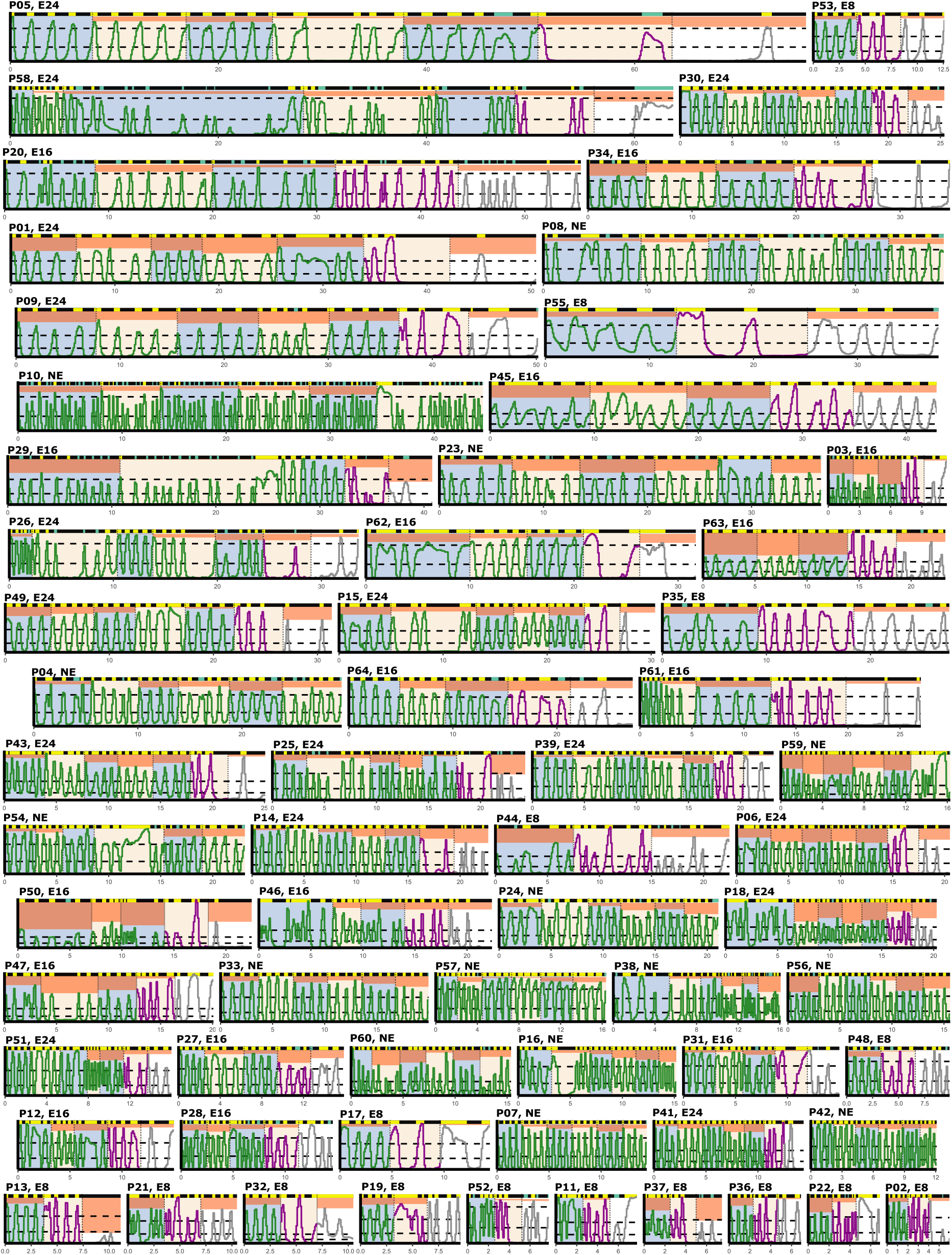
Participant-level response-force traces across blocks of interest. Each panel shows standardized grip force over time (s) for one participant. Trace color denotes block type (green = rewarded blocks; purple = early extinction; grey = late extinction). Horizontal dashed lines indicate the set and reward thresholds. Orange rectangles mark blockwise peak force; the lower edge of each rectangle indicates the maximum force recorded within that block, and the absence of an orange rectangle indicates that this block contained the highest peak force among all depicted blocks. Background shading denotes stage coding (blue = before; beige = after; unshaded = late extinction). The band above each trace provides a visual summary of the timing and duration of key behavioral events (yellow = criterion responses; aqua = subthreshold responses; black = idle state). Participant ID and group are shown at the top of each panel.

**Figure 3:**
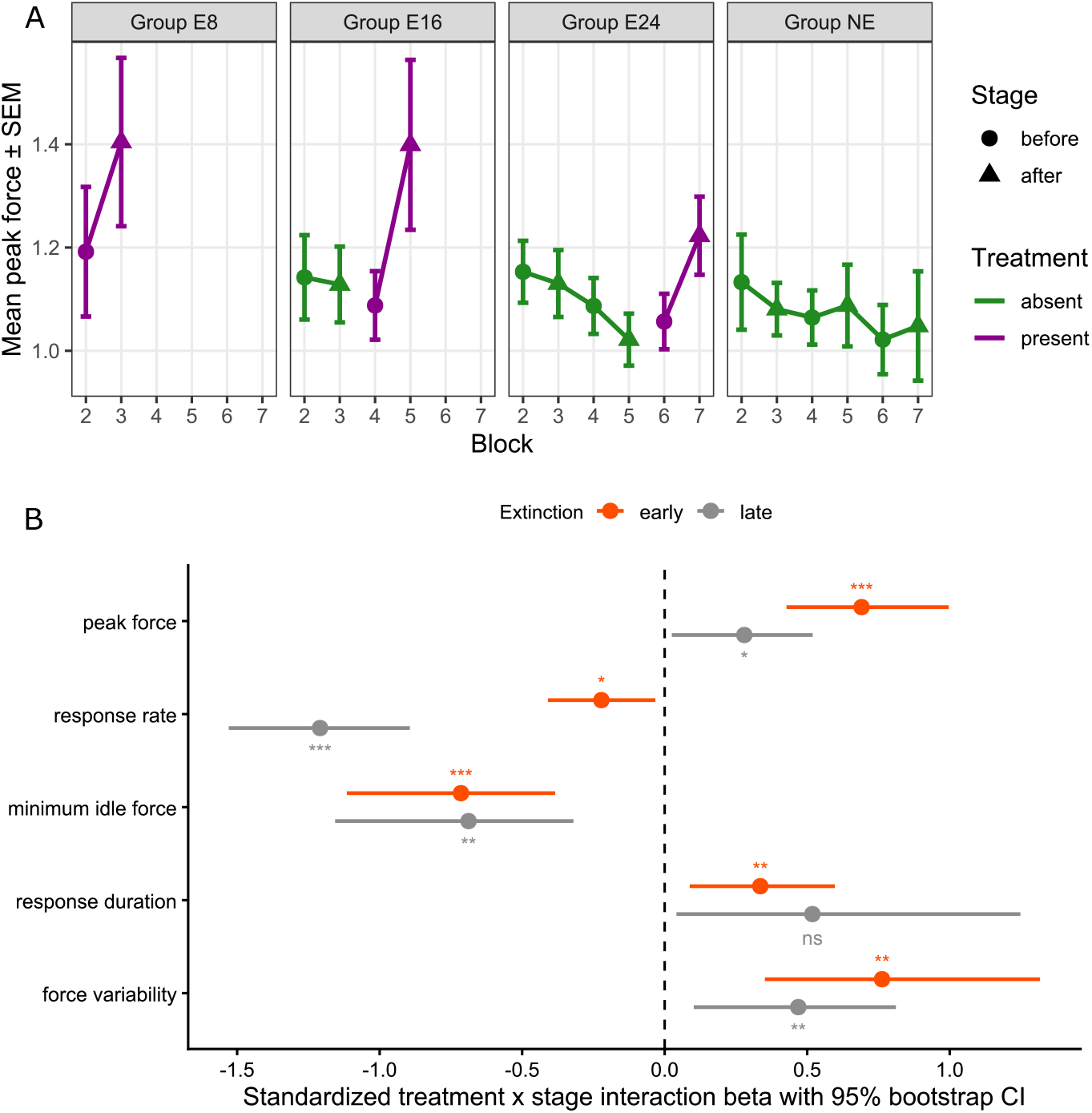
Grouped summaries and standardized interaction effects. (a) Mean blockwise peak force (*±* SEM) across groups, separated by stage (before vs after; marker shape) and treatment (absent vs present; trace color). (b) Standardized stage × treatment interaction coefficients (*β ±* 95% CI) for each behavioral metric in early (orange) and late (gray) extinction. The vertical dashed line denotes a null interaction effect (*β* = 0). Color-matched significance annotations indicate conventional *p*-value thresholds (ns (*p* 0.05), * (*p <* 0.05), ** (*p <* 0.01), *** (*p <* 0.001)).

The main effect of block was negative (*β* = −0.03, 95% CI [−0.05, −0.01]) and statistically significant (*p* = 0.007). This result indicates that participants’ peak force decreased by roughly 3% of their standardized force with each block (visible in Figure 3a as a general downward trend for extinction absent tiers). The stage × treatment interaction therefore reflects an extinction-onset increase that occurs against, and departs from, this overall downward trend. The intra-class correlation for peak force was high (ICC = 0.79, 95% CI [0.71, 0.85]) and statistically significant (*p* = 1 × 10^−4^), indicating that 79% of the total variance in peak force is robustly attributable to stable between-participant differences. As Figure 3b shows, the stage × treatment interaction was smaller in late extinction than in early extinction, but it remained significantly above zero (standardized *β* = 0.28, 95% CI [0.03, 0.52]; *p* = 0.024).

Figure 4 provides a complementary episode-aligned depiction in which each criterion-defined episode is expressed as a deviation from the last rewarded episode immediately preceding extinction onset (episode −1). In this depiction, peak force shows a clear positive increase at the first non-rewarded episode (episode +1) and continues to rise through episode +3. Beyond that point, peak force remains elevated without an evident increase or decrease over the remaining plotted episode indices. Note that later indices are supported by fewer participant–tier instances, and indices with *<* 10 cases were omitted. It is also worth noting that because the discrete metrics were computed from criterion responses only, they may underestimate any decline in force during late extinction; in that phase, subthreshold responses (see Figure 1c) accounted for about one quarter of all responses, and six participants emitted no criterion responses at all (see Figure 2). The continuous-force analysis, which is computed from the raw force over time, is not subject to these response-level and participant-level selection biases.

**Figure 4:**
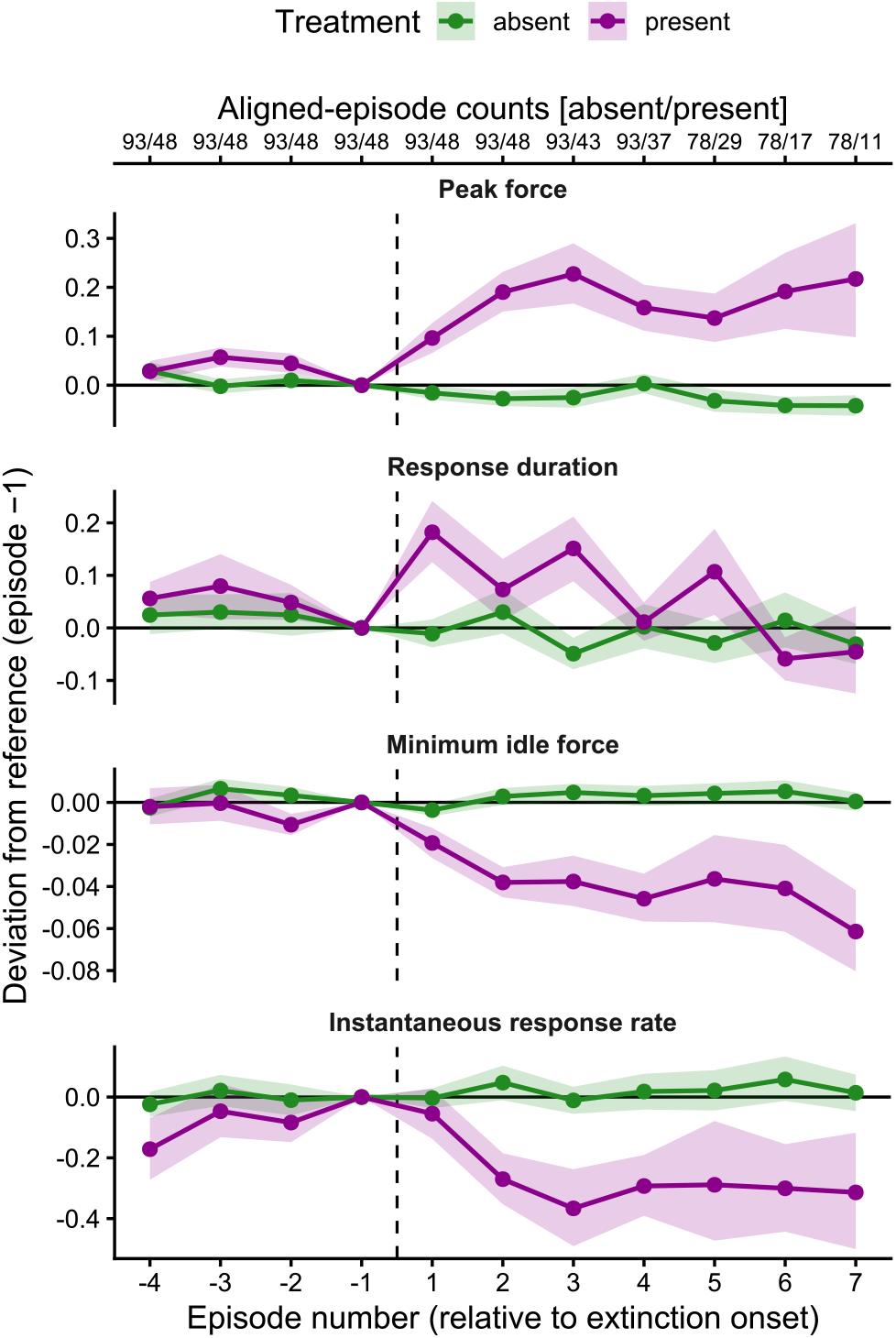
Episode-aligned analysis of behavioral metrics across extinction. Values are pooled across groups (E8, E16, E24, and NE). The x-axis shows an episode-aligned sequence of criterion-defined response episodes before (negative indices; the rewarded block immediately preceding extinction onset) and after (positive indices; spanning early and late extinction) the introduction of extinction. The dashed vertical line marks the onset boundary (between episode −1 and episode +1). For each metric, values were expressed as deviations from the reference episode immediately preceding extinction onset (episode −1 centered at zero). Green points show mean performance for tiers in which extinction was absent at the tier boundary (reward →reward), whereas purple points show mean performance for tiers in which extinction was present at the tier boundary (reward →extinction). Shaded bands depict 68% confidence intervals. Numbers above the panels indicate the number of contributing participant–tier instances at each episode index, shown separately for the absent/present conditions. Case counts can vary across episode indices because post-onset responding differs across participants, and can also differ between conditions when later episode indices are supported by fewer cases. Episode indices with fewer than 10 participant–tier instances in a given condition were omitted from plotting. Minimum idle force was defined as the minimum force observed between consecutive criterion responses. Instantaneous response rate was defined as the reciprocal of the inter-response interval[28]. The force-variability metric is omitted here because it is defined over time-binned windows rather than single episodes.

### 3.2 Exploratory metrics show distinct patterns across extinction

Figure 3b shows that the stage × treatment interaction varies across force-related metrics beyond peak force, with distinct early-versus-late extinction profiles. For response rate, the interaction was negative and significant during early extinction (standardized *β* = −0.22, 95% CI [−0.41, 0.03], *p* = 0.019), and it became strongly negative during late extinction (standardized *β* = −1.21, 95% CI [−1.53, −0.89], *p* = 1× 10^−4^). In the complementary episode-aligned depiction (Figure 4), response rate shows little change at the first non-rewarded episode (episode +1) but becomes progressively more negative over the next few episodes, reaching a clearly negative deflection by approximately episode +3 and remaining below the pre-onset reference across the remaining plotted episodes. A convergent disengagement-related pattern is visible for minimum idle force; Figure 3b shows a negative interaction in both early (standardized *β* = −0.71, 95% CI [−1.11, − 0.38]; *p* = 0.001) and late (standardized *β* =− 0.69, 95% CI [−1.16, −0.32]; *p* = 0.003) extinction, and Figure 4 shows that minimum idle force drops below the episode −1 reference immediately after onset and remains negative throughout the post-onset sequence.

Regarding indices expected to increase during extinction, response duration showed a positive stage × treatment interaction during early extinction (standardized *β* = 0.34, 95% CI [0.09, 0.60]; *p* = 0.010) that remained positive but non-significant in late extinction (standardized *β* = 0.52, 95% CI [0.04, 1.25]; *p* = 0.095). Accordingly, Figure 4 shows that response duration increases sharply at episode +1 and then exhibits fluctuations across subsequent episodes, remaining elevated relative to the episode −1 reference for several early post-onset episodes before dipping slightly below the episode −1 reference at the last plotted episodes. Force variability, for its part, displayed a stronger interaction in early extinction (standardized *β* = 0.76, 95% CI [0.35, 1.32]; *p* = 0.003) that attenuated in late extinction (standardized *β* = 0.47, 95% CI [0.10, 0.81]; *p* = 0.009); this metric is not shown in Figure 4 because it is defined over time-binned windows rather than single episodes.

### 3.3 Continuous force shows a rise-and-fall pattern

Figure 5 shows a brief pulse of continuous-force increment shortly after extinction onset. The green trace provides a reference for any increment that would have occurred in the absence of extinction. The pulse in the purple trace, attributable to extinction onset, occurs around the first quarter of the early extinction block and then quickly subsides to a level slightly below the reference trace at the beginning of late extinction. Toward the end of late extinction, force declined further below. This analysis complements the discrete-metrics trajectory analysis (Figure 4) in showing that although the maximum force of criterion responses did not detectably decline across seven behavioral episodes, general force exertion decreased over time, due to the gradual decline in response rate. In other words, we observed an intensification in our target behavior before the disengagement of that same behavior. The inset in Figure 5 shows that the location of peak continuous force in smoothed traces varied widely among participants who underwent extinction; although a sizable proportion of participants showed peak force early after extinction onset, a non-negligible number of participants reached maximum force in the second half of early extinction or in late extinction.

**Figure 5:**
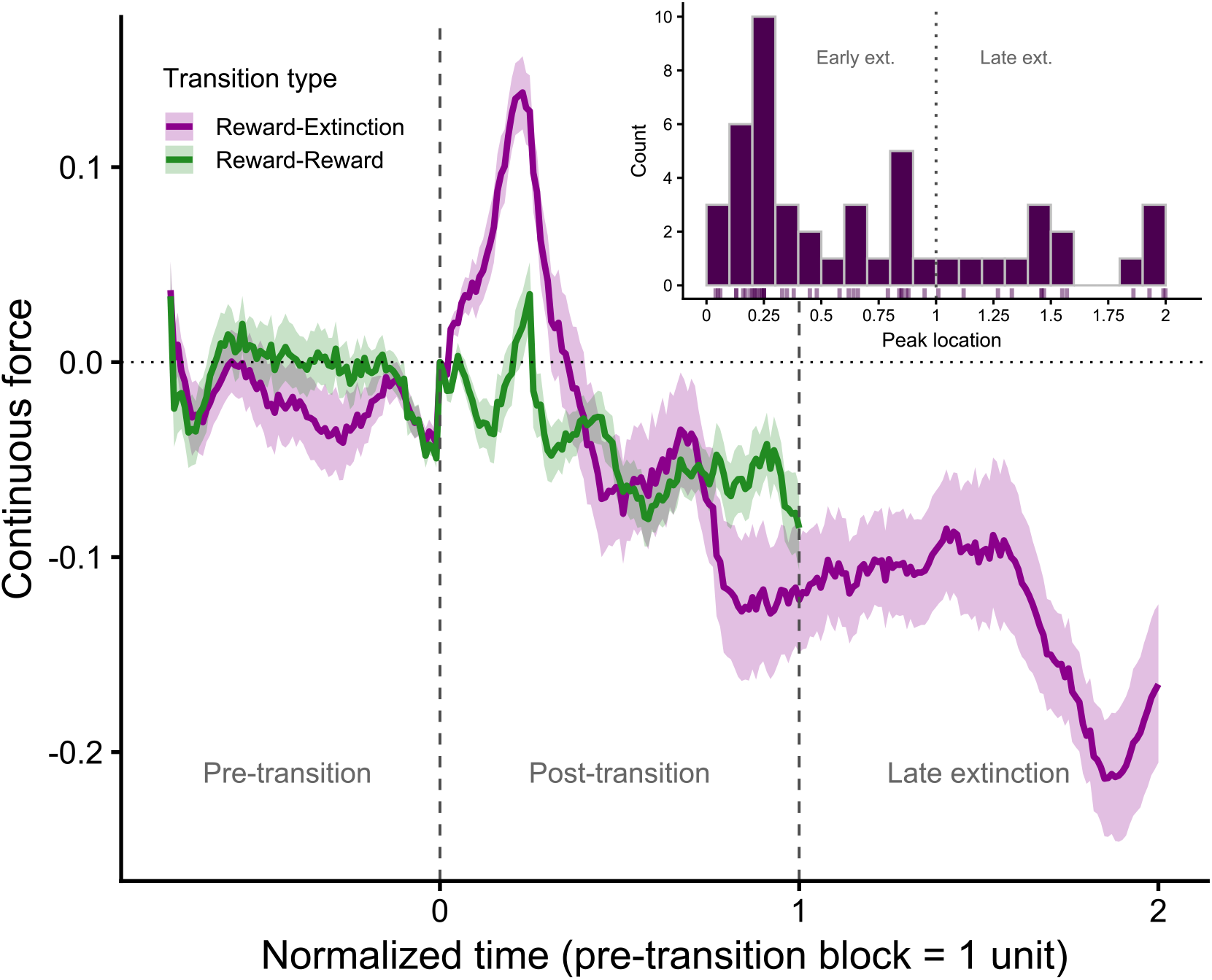
Continuous force across reward-extinction and reward-reward transitions. Shaded ribbons denote 68% bootstrap confidence intervals (1,000 resamples; approximating ± 1 *SEM*). *y* = 0 equals the mean of the last pre-transition window, so that deviations on this axis are interpretable as proportional increases or decreases from that reference. The reward →extinction trace pools the performance of participants in the three groups that involved extinction (E8, E16, and E24). The reward →reward counterfactual (NE group only; 45 transitions from 15 participants) controls for generic block-transition effects. The inset shows the distribution of individual peak locations in smoothed force traces across the normalized extinction window.

## 4 Discussion

### 4.1 Rigorous evidence for the extinction burst

The extinction process can present a non-monotonic path if it includes an extinction burst; the often short-lived nature of such a phenomenon makes its detection highly dependent on operational choices. Here, we provide evidence of an extinction burst indexed by hand-grip force in a human sample. The effect we observed was large and robust, and our design and measurement operationalization were preregistered. We selected response force, rather than response rate, as the primary index of behavioral invigoration. This choice was motivated by concerns that the target response rate can increase when the reward is discontinued due to the release of competition from alternative responses[13], an increase that does not by itself constitute evidence of genuine intensification. Furthermore, our human task enabled symbolic reward delivery that involved no handling, collection, or intake behaviors whatsoever which might otherwise be incompatible with our target behavior. Altogether, the focus on response force and the use of symbolic reward delivery jointly render release-from-competition interpretations unlikely in the present study.

Furthermore, our analysis of aggregate continuous force rules out an additional confound. An observed increase in peak force might merely reflect increased variability once responding becomes ineffective, rather than a net intensification. Force may vary slightly while the response continues to produce reward, but vary more once it no longer does, producing upward peaks accompanied by equivalent downward swings. Under that scenario, the mean force would remain unchanged even as the peaks grew, and an apparent intensification would amount to a measurement artifact. This possibility is addressed directly by averaging. Deflections that are symmetric about the mean cancel when force is averaged across participants, so a variability-driven inflation of peaks cannot produce a rise in the mean trajectory. Figure 5 shows that re-standardized, averaged, and smoothed continuous force does rise following extinction onset, and that the reward→extinction trajectory departs upward from the matched reward→reward reference. The increase we observed is therefore a net increase in force, not an artifact of greater variability.

Our study addressed several methodological limitations of previous extinction-burst investigations that used response force as the behavioral index of interest[18, 16, 15]. We examined very short time windows to capture a behavioral intensification we anticipated would be highly transient. In the same vein, as we anticipated a transient rise-and-fall of behavior, we sought to compare pre–post maximum values rather than averages because the latter are more susceptible to attenuation whenever an increase wears off. Unlike previous studies indexing extinction bursts through force, we based our conclusions on conventional statistical inference benchmarks rather than relying on descriptive statistics. Since the nature, validity, and ubiquity of extinction bursts are currently hotly debated, we preregistered our behavioral task, measurements of interest, study design, sample size, and statistical inference approach. Preregistration allowed us to disclose which decisions changed after data collection and which data/theory-driven issues we had planned for in advance. Our study is the first extinction-burst study comparing extinction against a concurrent, randomized control group matched in reward history.

Our analytic approach addresses the definitional and measurement difficulties that others have identified[13]. In our view, detecting extinction bursts requires comparing the observed reward→extinction transition against a reference trajectory derived from matched reward→reward transitions, using models that assess the stage × treatment interaction. The before–after stage factor captures change through time at the onset boundary, whereas the present/absent factor represents how behavior would have evolved had the individuals continued to be rewarded—the counterfactual trajectory. Matching is crucial because it helps account for secular trends, reducing reliance on stringent baseline stability criteria[20] prior to extinction onset and thereby lowering the risk of ceiling effects. This approach resembles the rationale of difference-in-differences estimation[29] in time-series analysis, but implemented within a fully randomized experimental design where the counterfactual is directly observed through staggered extinction onset across groups.

Beyond our preregistered confirmatory assessment, our exploratory probes shed additional light on the broader behavioral profile of extinction bursts. In line with prior work, we observed a burst-like increase in response duration[30], which may partly reflect the simple fact that generating higher force can take more time. We also observed a burst-like increase in force variability. In parallel, we found hints of behavioral disengagement—evident as early as the initial non-rewarded episode in our minimum idle force metric, and profoundly so in late extinction for response rate. Notably, we did not observe a burst in response rate. This finding is consistent with a time-allocation account in which apparent rate increases during extinction arise because reward-related activities no longer compete with the target response[11, 12]. In this context, the absence of a response-rate burst is unsurprising because in our task extinction removed little or none of the usual competing activities that can inflate gross response-rate estimates.

### 4.2 Limitations and open questions

The only one of our metrics that showed a clearly non-monotonic change around extinction onset, consistent with the transient character typically associated with burst phenomena, was response duration in our episode-aligned exploratory analysis. Transient increases followed by declines in the peak force of criterion responses might have emerged had we implemented a longer extinction period. Despite the advantages of using human participants, a limitation of our behavioral protocol is that extended reward discontinuation periods are difficult to sustain while keeping participants engaged and preserving the integrity of the task. Our protocol excluded participants who did not complete the session; with longer extinction periods, more participants might reasonably interpret the omission of reward as a task malfunction and seek clarification, thereby disrupting the extinction implementation. Accordingly, longer extinction exposures remain to be tested for force-indexed bursts, and animal preparations may offer an especially suitable setting for such work.

That said, it is worth considering that by design our episode-aligned exploratory analysis only included criterion responses. This decision reflects our priority in detecting increases attributable to extinction, rather than in detecting any subsequent decline. About a quarter of responses in late extinction did not reach the reward threshold and therefore were not included in the episode-aligned analysis. Thus, it can be said that there were indeed quite a number of weak responses past the very first stage of extinction. Those responses, and even force squeezes too weak to cross the detection threshold for subthreshold responses (see, for example, the two tiny bumps in P13’s late extinction in Figure 2), might have contributed to a decline in peak force had we included them in this analysis. We did not extend the episode-aligned analysis to subthreshold responses for two reasons. Although extinction bursts have been defined as transient increases, there is no principled criterion to set how short-lived they should be in order for them to be considered transient enough. In fact, behavioral increases upon extinction have been considered by some authors to be inconsistent because of not being sustained enough[18]. Moreover, none of the nine operational definitions compiled by Katz and Lattal [20] include a transiency criterion; historically, bursts have been certified on the basis of an increase relative to baseline alone. We believe that this is the correct way of thinking about extinction bursts. Despite transiency being a prevalent feature in conceptual definitions, extinction bursts have been detected through increases—a sound option due to the difference between bursts as an increase and the subsequent decrease as extinction learning. That is why we prioritized the detection of an increment rather than documenting a subsequent decrease. We wish to note that our dataset is available in an open repository for further analyses tracking possible subsequent declines in peak force.

The dynamics of behavioral measurements other than peak force should also be interpreted cautiously. It is plausible that the initial increase in force at extinction onset partially drove the observed changes in other indices. As noted above, the increase in response duration may, in part, reflect the increased time required to generate higher force. Similarly, our disengagement indices—minimum idle force and response rate—could, at least in part, reflect fatigue and recovery periods after extreme force exertion rather than true detachment from reward pursuit. Future work should aim to disentangle these relationships across metrics. That said, the peak-force effect we observed was pronounced and places extinction bursts, when indexed by force, as a valid and potentially adaptive behavioral phenomenon[1].

In the present study, we prioritized the assessment of maximum force in order to ascertain whether a true behavioral intensification emerged in extinction. That aim led us to treat force variation as a confound to be ruled out. Extinction-induced behavioral variation can, however, be regarded as a phenomenon of interest in its own right. Intensification could disambiguate whether the response still produces reward; disengagement could prevent the energy wasted on now-unfruitful behaviors. Between the two lies variation, which can lead to discovering new ways of attaining reward. Whereas our treatment of variation was restricted to oscillations in a single dimension (force), the literature documents a broader range of variation phenomena, including the emergence of different, non-rewarded responses[7]. We could have gathered information concerning this matter had we recorded, for example, the tilt of the dynamometer, which our hardware supports. In addition, we could have video-recorded participants as they performed the task to assess whether they adopted different postures or presented other forms of behavioral variation during extinction. Future work might explore these avenues of inquiry.

### 4.3 A need for theories beyond response competition

Among the accounts developed to explain extinction bursts, one of the most influential attributes them to frustration[2]. The key idea is that when a behavior reliably produces a reward, it comes to elicit a reward-anticipatory behavioral state; then, whenever a reward is unexpectedly omitted, frustration ensues. Frustration is then proposed to invigorate behavior, which can manifest as a burst. Importantly, the burst is only one element in a broader cluster of responses associated with frustration[2, 31]. Organisms are typically observed to vary their behavior upon extinction and, with prolonged reward discontinuation, they may eventually disengage from reward pursuit—at least in terms of the original target response.

Frustration-based accounts of extinction bursts have been criticized for inviting circular reasoning[13], in that “frustration” labels the behavioral syndrome observed during early extinction and is also offered as its explanation. This concern is not unique to frustration, however; circularity is a recurring concern for many explanatory constructs in psychology, and it does not by itself disqualify the account. Aside from circularity, frustration has been argued to produce reactions that are meaningful in their own right yet ancillary to the goal-directed activity in which burst-like changes are observed[11]. Accordingly, however important, stress-like reactions such as corticosteroid release, odor emissions, vocalizations, aggression, and escape seem not to be clearly linked to the target instrumental behavior. Moreover, frustration has been criticized as being difficult to scale, operationalize, or manipulate cleanly in many laboratory preparations[13].

These difficulties drove the search for a more parsimonious alternative that dispenses with motivational constructs altogether. On release-from-competition accounts[11, 13, 12], the burst reflects an increase in frequency without a genuine intensification of the target goal-directed behavior itself; instead, the target behavior becomes more prevalent due to a mere occupation of the gap left by its competing activities. This reconceptualization is parsimonious precisely because it requires no appeal to internal states. However, as we have argued, our results challenge this account. Increases in force upon extinction, obtained when no explicit consummatory activity competed with the target response, cannot be readily explained by release from competition.

A candidate alternative account that naturally accommodates non-linear dynamics conceives behavior as the activity of closed-loop systems, a perspective rooted in control theory[32]. On this view, reward omission can be regarded as a goal-attainment error (i.e., a mismatch between expected and obtained outcomes) that gives rise to a transient behavioral state. This state jointly determines two outcomes: (1) an increased drive or urgency to re-establish the goal[33], and (2) value updating that lowers the expected worth of the action that previously produced the reward (linked to extinction learning). These outcomes occur at different levels of a hierarchically organized control system[34], in which distinct modules pull behavior toward opposite end states[35]. Under continued reward, such a system raises the value of the rewarded action while lowering the drive to perform it. Reward omission would have an opposite effect on both, lowering value while raising drive. Because the two quantities need not change at the same pace, their relative rates of change could produce qualitatively different trajectories. Should drive rise more rapidly than value falls, we would observe a transient intensification of behavior, an escalation in goal pursuit followed by attenuation as extinction learning proceeds. Conversely, if value falls sufficiently rapidly, a more monotonic decline would ensue without any noticeable burst.

As is apparent, the control-theoretic framing is closely akin to the frustration account, though it goes beyond in distinguishing the mechanism producing the behavioral intensification (error) from the intensification itself—which may or may not occur. Such a distinction thus putatively resolves the circularity issue ascribed to frustration-based explanations. Aside from this conceptual advantage, however, our design cannot support either account over the other. Our comparisons contrast nonrewarded blocks against rewarded ones, which establishes that a force-indexed burst occurs but cannot isolate the contribution of any particular mechanism to it. Supporting a frustration-based account of extinction-induced force increase would require establishing a contrast between unexpected and expected nonreward[36]. Crucial tests should implement designs in which nonrewarded conditions are compared after different reward histories, rather than nonreward against continued reward. Manipulations of reward magnitude, reward intermittency, and the amount of training preceding extinction would afford such comparisons, and all are plausible modulators of burst intensity. Even so, none of these manipulations would disambiguate the frustration and control-theoretic accounts or favor one over the other, because surprising nonreward and goal-attainment error are functionally equivalent in this context.

A control-theoretic account of the extinction burst does make a unique prediction that diverges from frustration theory. Since behavior on this approach represents the control of perceptual variables rather than the production of specific actions[37], burst-like transients would not always take the form of an upward spike in the measured output. This possibility is at odds with the notion of reward-omission-induced invigoration, which anticipates intensification as an increase in output. Perceptions that reliably precede goal attainment become intermediate targets[38, 39]. Thus, a burst can be interpreted as a transient escalation in the pursuit of such targets when the reward unexpectedly fails to occur. In the present task, a candidate controlled perception is the haptic sensation associated with exerted effort; reward omission may therefore transiently intensify the pursuit of that perception, with increased output emerging as a consequence. Broadly, the transient change at extinction onset should occur in the direction required to bring about the perceptual state associated with goal attainment. The novel prediction deriving from this assumption is that an “upward” burst should not be expected when the task policy requires a downward shift in an output parameter. In such cases, the pursued perception would be associated with output attenuation, and the effect would be expected to manifest as a transient drop below baseline rather than as an increase. Hence, under appropriate circumstances, burst-like phenomena may materialize as a downward notch (“negative bursts”)—a prediction that remains to be tested.

## Supporting information

Electronic Supplementary Materials

## Notes

https://github.com/RSosa-BehavioralScience/extinction_burst_force

### Competing Interest Statement

The authors have declared no competing interest.

### Summary of Updates

This version incorporates revisions made in response to the final round of peer review at Royal Society Open Science. The Methods now specify that the change in screen color mentioned in the participant instructions signaled the end of the task and occurred only once, at the end of the session. The description of the continuous-force analysis was rewritten to make its relation to the episode-aligned analysis easier to follow. The Results and Discussion now explain how these two levels of analysis relate to each other. In short, the episode-aligned analysis tracks properties of individual responses that met the reward criterion, whereas the continuous-force analysis tracks deviations in force over time regardless of which response property produced them. During early extinction, peak force and response duration increased while response rate decreased. Because high-force episodes became less frequent as extinction progressed, continuous force rose and then fell even though peak force per episode did not decline. We argue that the two analyses yield complementary patterns rather than conflicting conclusions. A new paragraph in the Discussion returns to the previous studies of response force reviewed in the Introduction and states what the present study adds to them. The Discussion also addresses at greater length whether transience should be treated as a defining feature of the extinction burst. We argue that the increase in responding and the decline associated with extinction learning are separable processes, and that previous studies have not set a criterion of transience in advance. Our design prioritized the detection of increases over tracking behavior until complete disengagement. Subthreshold responses accounted for about a quarter of all responses in late extinction; they were excluded from the episode-aligned analysis by design, but including them would likely have produced a decrease in peak force in that analysis. We note that our dataset and analysis code are archived in an open repository on Zenodo (https://doi.org/10.5281/zenodo.23267915)and are available for any further analysis. The publication year of B. F. Skinner, The Behavior of Organisms, was corrected from 2038 to 1938 in the reference list.

### Funding

- Sistema Nacional de Investigadores: 64324
- Comunidad de Madrid: 2020-T2/SOC-20287
- Secretaría de Ciencia, Humanidades, Tecnología e Innovación: 1044308

