## Supplementary material for "Evidence for the extinction burst in grip force: A preregistered study": Electronic Supplementary Materials: ebf_participants_instructions.pdf

***Equipment Setup and Participant Instructions: Grip-Force Task***
***Manuscript: "Evidence for the extinction burst in grip force: A preregistered study"***

### *Equipment Setup*

*Before each participant arrived, the researcher turned on the Dell laptop, confirmed it was connected to power, verified that the Vernier Go Direct® dynamometer was correctly attached via USB, and set the system volume to level 26. Participant information (including dominant hand) was recorded in a spreadsheet before the participant entered the cubicle and took a seat. The researcher explained that initial instructions would be given verbally, followed by a series of on-screen instructions the researcher would read aloud before leaving the participant to complete the main task alone.*

### *Initial verbal instructions*

Once the participant was seated, the researcher delivered the following instructions exactly:

> "Hi, thank you for participating. First, please sit as close to the table as possible. The study involves using your dominant hand to squeeze the dynamometer when the screen asks you to do so. Later I will explain how to squeeze it. To make sure you only use that hand, we will place a tennis ball in your non-dominant hand. During the task, please follow only the instructions that appear on the screen. Please do not turn around or get up from the chair until we come back for you; otherwise, you will be excluded from the experiment and will not receive the money you earned. The experiment will last approximately five minutes, and the device does not deliver electric shocks or cause any harm."

The researcher placed a tennis ball in the participant's non-dominant hand and positioned the dynamometer in the dominant hand, ensuring the participant was comfortable.

### *On-screen instructions*

The researcher then launched the PsychoPy program, entered the participant ID, and read aloud each instruction screen:

> "Audio check. If you hear a sound, press the space bar to continue. (If you do not hear anything, tell the researcher.)"

A consent screen followed, accompanied by an information screen summarizing all study details, risks, and compensation:

> "If you give your consent, press the space bar."

Participants then received device and posture instructions:

> "Hello :) You will only need to use: –The dynamometer to squeeze, and –The space bar to continue. NOTHING ELSE. To continue, press space."

> "Hold the dynamometer throughout the ENTIRE task. You may relax your hand without squeezing as long as you continue to hold it. NEVER place it on the table or on any surface until the researcher returns. To continue, press space."

"The dynamometer measures how much force you generate when squeezing it. Always hold it with your dominant hand (the one you write with). Grab the dynamometer by the TWO long BLUE bars. To continue, press space."

"The blue circles on the dynamometer DO NOT do anything. Please do not touch them. To continue, press space."

"Rest your dominant arm on the table so it does not get tired when squeezing the dynamometer. Never release the tennis ball in your non-dominant hand. To continue, press space."

### *Calibration phase*

Participants then received the following calibration instructions:

"Next, you will have attempts to squeeze the dynamometer with a lot of force. The screen will guide you. In each attempt, you will have 5 seconds to squeeze very hard. It does not have to be your maximum force—just very hard. To continue, press space."

Two calibration trials were presented, each consisting of a 5-s squeezing interval ("Squeeze very hard") followed by a 5-s rest interval ("Relax your hand and wait"). The average force of the two trials defined the participant's 100% reference value, from which the 30% and 70% force thresholds were computed for the main task.

### *Instructions before the main task*

Participants were then told:

"Next, you will need to discover how to use the dynamometer to earn money. The money you accumulate will be given to you at the end of the experiment. The more money you earn, the more you are supporting science. To continue, press space."

"IMPORTANT: To figure out how to earn money using the dynamometer, you may ONLY USE YOUR DOMINANT HAND (NOT BOTH HANDS). To continue, press space."

"Wait until the researcher leaves the room and, once the door has closed, press the space bar to continue."

The researcher then provided an additional clarification verbally:

"On the next screen, the background will change to a yellow/orange color with two phrases—one at the top and one at the bottom. On that next screen, you must try to figure out how to earn as much money as possible. The phrase at the bottom will remind you that when the experiment ends, the background will change back to blue."

The researcher then left the cubicle and closed the door.

***Main task***
The task screen displayed:

| |
|---|
| "Ready! Now try to earn as much money as possible!" |

| |
|---|
| "Do not get up from your chair or release the dynamometer until the screen changes color, or you will not receive any money." |

The task consisted of 8, 16, 24, or 28 reinforced trials depending on group assignment. On each trial, the participant had to maintain the force below 30% of their calibration average and then exceed 70% to earn a reinforcer (a 0.5-s image of a coin accompanied by a brief tone). After the reinforced trials, groups E8, E16, and E24 transitioned to extinction for a duration equal to twice the time required to complete their final four reinforced trials. The NE group ended immediately without extinction. During extinction, squeezing produced no consequences, but responses were recorded.

Throughout the task, the researcher monitored the participant from outside the cubicle to ensure compliance with exclusion criteria (remaining seated, using only the dominant hand, maintaining continuous hold of the dynamometer, not turning, and not using body momentum).

***End of session***
When the final screen appeared, the researcher re-entered the cubicle, removed the tennis ball and dynamometer, and said:

| |
|---|
| "Thank you very much, we are finished." |

Participants were then paid the amount shown on the final screen ("Total accumulated money: X").
