## Supplementary material for "Evidence for the extinction burst in grip force: A preregistered study": Electronic Supplementary Materials: EBExperiment_force.py

```
#!/usr/bin/env python
# -*- coding: utf-8 -*-
"""
This experiment was created using PsychoPy3 Experiment Builder (v2024.2.4),
    on noviembre 28, 2025, at 09:23
If you publish work using this script the most relevant publication is:

    Peirce J, Gray JR, Simpson S, MacAskill M, Höchenberger R, Sogo H, Kastman E, Lindeløv JK. (2019) 
        PsychoPy2: Experiments in behavior made easy Behav Res 51: 195. 
        https://doi.org/10.3758/s13428-018-01193-y

"""

# --- Import packages ---
from psychopy import locale_setup
from psychopy import prefs
from psychopy import plugins
plugins.activatePlugins()
prefs.hardware['audioLib'] = 'ptb'
prefs.hardware['audioLatencyMode'] = '3'
from psychopy import sound, gui, visual, core, data, event, logging, clock, colors, layout, hardware
from psychopy.tools import environmenttools
from psychopy.constants import (NOT_STARTED, STARTED, PLAYING, PAUSED,
                                STOPPED, FINISHED, PRESSED, RELEASED, FOREVER, priority)

import numpy as np  # whole numpy lib is available, prepend 'np.'
from numpy import (sin, cos, tan, log, log10, pi, average,
                   sqrt, std, deg2rad, rad2deg, linspace, asarray)
from numpy.random import random, randint, normal, shuffle, choice as randchoice
import os  # handy system and path functions
import sys  # to get file system encoding

import psychopy.iohub as io
from psychopy.hardware import keyboard

# --- Setup global variables (available in all functions) ---
# create a device manager to handle hardware (keyboards, mice, mirophones, speakers, etc.)
deviceManager = hardware.DeviceManager()
# ensure that relative paths start from the same directory as this script
_thisDir = os.path.dirname(os.path.abspath(__file__))
# store info about the experiment session
psychopyVersion = '2024.2.4'
expName = 'exp prueba'  # from the Builder filename that created this script
# information about this experiment
expInfo = {
    'participant': '',
    'session': '1',
    'assign_seed': '123456789',
    'date|hid': data.getDateStr(),
    'expName|hid': expName,
    'psychopyVersion|hid': psychopyVersion,
}

# --- Define some variables which will change depending on pilot mode ---
'''
To run in pilot mode, either use the run/pilot toggle in Builder, Coder and Runner, 
or run the experiment with `--pilot` as an argument. To change what pilot 
#mode does, check out the 'Pilot mode' tab in preferences.
'''
# work out from system args whether we are running in pilot mode
PILOTING = core.setPilotModeFromArgs()
# start off with values from experiment settings
_fullScr = True
_winSize = [1536, 864]
# if in pilot mode, apply overrides according to preferences
if PILOTING:
    # force windowed mode
    if prefs.piloting['forceWindowed']:
        _fullScr = False
        # set window size
        _winSize = prefs.piloting['forcedWindowSize']

def showExpInfoDlg(expInfo):
    """
    Show participant info dialog.
    Parameters
    ==========
    expInfo : dict
        Information about this experiment.
    
    Returns
    ==========
    dict
        Information about this experiment.
    """
    # show participant info dialog
    dlg = gui.DlgFromDict(
        dictionary=expInfo, sortKeys=False, title=expName, alwaysOnTop=True
    )
    if dlg.OK == False:
        core.quit()  # user pressed cancel
    # return expInfo
    return expInfo


def setupData(expInfo, dataDir=None):
    """
    Make an ExperimentHandler to handle trials and saving.
    
    Parameters
    ==========
    expInfo : dict
        Information about this experiment, created by the `setupExpInfo` function.
    dataDir : Path, str or None
        Folder to save the data to, leave as None to create a folder in the current directory.    
    Returns
    ==========
    psychopy.data.ExperimentHandler
        Handler object for this experiment, contains the data to save and information about 
        where to save it to.
    """
    # remove dialog-specific syntax from expInfo
    for key, val in expInfo.copy().items():
        newKey, _ = data.utils.parsePipeSyntax(key)
        expInfo[newKey] = expInfo.pop(key)
    
    # data file name stem = absolute path + name; later add .psyexp, .csv, .log, etc
    if dataDir is None:
        dataDir = _thisDir
    filename = u'data/%s_%s_%s' % (expInfo['participant'], expName, expInfo['date'])
    # make sure filename is relative to dataDir
    if os.path.isabs(filename):
        dataDir = os.path.commonprefix([dataDir, filename])
        filename = os.path.relpath(filename, dataDir)
    
    # an ExperimentHandler isn't essential but helps with data saving
    thisExp = data.ExperimentHandler(
        name=expName, version='',
        extraInfo=expInfo, runtimeInfo=None,
        originPath='C:\\Users\\palom\\OneDrive\\Escritorio\\UP-UNED Experimento\\exp prueba_lastrun.py',
        savePickle=True, saveWideText=True,
        dataFileName=dataDir + os.sep + filename, sortColumns='time'
    )
    thisExp.setPriority('thisRow.t', priority.CRITICAL)
    thisExp.setPriority('expName', priority.LOW)
    # return experiment handler
    return thisExp


def setupLogging(filename):
    """
    Setup a log file and tell it what level to log at.
    
    Parameters
    ==========
    filename : str or pathlib.Path
        Filename to save log file and data files as, doesn't need an extension.
    
    Returns
    ==========
    psychopy.logging.LogFile
        Text stream to receive inputs from the logging system.
    """
    # set how much information should be printed to the console / app
    if PILOTING:
        logging.console.setLevel(
            prefs.piloting['pilotConsoleLoggingLevel']
        )
    else:
        logging.console.setLevel('warning')
    # save a log file for detail verbose info
    logFile = logging.LogFile(filename+'.log')
    if PILOTING:
        logFile.setLevel(
            prefs.piloting['pilotLoggingLevel']
        )
    else:
        logFile.setLevel(
            logging.getLevel('info')
        )
    
    return logFile


def setupWindow(expInfo=None, win=None):
    """
    Setup the Window
    
    Parameters
    ==========
    expInfo : dict
        Information about this experiment, created by the `setupExpInfo` function.
    win : psychopy.visual.Window
        Window to setup - leave as None to create a new window.
    
    Returns
    ==========
    psychopy.visual.Window
        Window in which to run this experiment.
    """
    if PILOTING:
        logging.debug('Fullscreen settings ignored as running in pilot mode.')
    
    if win is None:
        # if not given a window to setup, make one
        win = visual.Window(
            size=_winSize, fullscr=_fullScr, screen=0,
            winType='pyglet', allowGUI=False, allowStencil=False,
            monitor='testMonitor', color=[-0.1686, 0.0667, 0.2000], colorSpace='rgb',
            backgroundImage='', backgroundFit='none',
            blendMode='avg', useFBO=True,
            units='height',
            checkTiming=False  # we're going to do this ourselves in a moment
        )
    else:
        # if we have a window, just set the attributes which are safe to set
        win.color = [-0.1686, 0.0667, 0.2000]
        win.colorSpace = 'rgb'
        win.backgroundImage = ''
        win.backgroundFit = 'none'
        win.units = 'height'
    if expInfo is not None:
        # get/measure frame rate if not already in expInfo
        if win._monitorFrameRate is None:
            win._monitorFrameRate = win.getActualFrameRate(infoMsg='Attempting to measure frame rate of screen, please wait...')
        expInfo['frameRate'] = win._monitorFrameRate
    win.hideMessage()
    # show a visual indicator if we're in piloting mode
    if PILOTING and prefs.piloting['showPilotingIndicator']:
        win.showPilotingIndicator()
    
    return win


def setupDevices(expInfo, thisExp, win):
    """
    Setup whatever devices are available (mouse, keyboard, speaker, eyetracker, etc.) and add them to 
    the device manager (deviceManager)
    
    Parameters
    ==========
    expInfo : dict
        Information about this experiment, created by the `setupExpInfo` function.
    thisExp : psychopy.data.ExperimentHandler
        Handler object for this experiment, contains the data to save and information about 
        where to save it to.
    win : psychopy.visual.Window
        Window in which to run this experiment.
    Returns
    ==========
    bool
        True if completed successfully.
    """
    # --- Setup input devices ---
    ioConfig = {}
    
    # Setup iohub keyboard
    ioConfig['Keyboard'] = dict(use_keymap='psychopy')
    
    # Setup iohub experiment
    ioConfig['Experiment'] = dict(filename=thisExp.dataFileName)
    
    # Start ioHub server
    ioServer = io.launchHubServer(window=win, **ioConfig)
    
    # store ioServer object in the device manager
    deviceManager.ioServer = ioServer
    
    # create a default keyboard (e.g. to check for escape)
    if deviceManager.getDevice('defaultKeyboard') is None:
        deviceManager.addDevice(
            deviceClass='keyboard', deviceName='defaultKeyboard', backend='iohub'
        )
    # create speaker 'sndCheck'
    deviceManager.addDevice(
        deviceName='sndCheck',
        deviceClass='psychopy.hardware.speaker.SpeakerDevice',
        index=-1
    )
    if deviceManager.getDevice('kbAudioCheck') is None:
        # initialise kbAudioCheck
        kbAudioCheck = deviceManager.addDevice(
            deviceClass='keyboard',
            deviceName='kbAudioCheck',
        )
    if deviceManager.getDevice('key_resp_6') is None:
        # initialise key_resp_6
        key_resp_6 = deviceManager.addDevice(
            deviceClass='keyboard',
            deviceName='key_resp_6',
        )
    if deviceManager.getDevice('kbWelcome') is None:
        # initialise kbWelcome
        kbWelcome = deviceManager.addDevice(
            deviceClass='keyboard',
            deviceName='kbWelcome',
        )
    if deviceManager.getDevice('key_resp_2') is None:
        # initialise key_resp_2
        key_resp_2 = deviceManager.addDevice(
            deviceClass='keyboard',
            deviceName='key_resp_2',
        )
    if deviceManager.getDevice('kbInstrUso') is None:
        # initialise kbInstrUso
        kbInstrUso = deviceManager.addDevice(
            deviceClass='keyboard',
            deviceName='kbInstrUso',
        )
    if deviceManager.getDevice('key_resp_4') is None:
        # initialise key_resp_4
        key_resp_4 = deviceManager.addDevice(
            deviceClass='keyboard',
            deviceName='key_resp_4',
        )
    if deviceManager.getDevice('key_resp_3') is None:
        # initialise key_resp_3
        key_resp_3 = deviceManager.addDevice(
            deviceClass='keyboard',
            deviceName='key_resp_3',
        )
    if deviceManager.getDevice('kbInstrCal') is None:
        # initialise kbInstrCal
        kbInstrCal = deviceManager.addDevice(
            deviceClass='keyboard',
            deviceName='kbInstrCal',
        )
    if deviceManager.getDevice('kbInstrReinf') is None:
        # initialise kbInstrReinf
        kbInstrReinf = deviceManager.addDevice(
            deviceClass='keyboard',
            deviceName='kbInstrReinf',
        )
    if deviceManager.getDevice('key_resp') is None:
        # initialise key_resp
        key_resp = deviceManager.addDevice(
            deviceClass='keyboard',
            deviceName='key_resp',
        )
    if deviceManager.getDevice('key_resp_5') is None:
        # initialise key_resp_5
        key_resp_5 = deviceManager.addDevice(
            deviceClass='keyboard',
            deviceName='key_resp_5',
        )
    # create speaker 'sndReinf'
    deviceManager.addDevice(
        deviceName='sndReinf',
        deviceClass='psychopy.hardware.speaker.SpeakerDevice',
        index=-1
    )
    if deviceManager.getDevice('kbEnd') is None:
        # initialise kbEnd
        kbEnd = deviceManager.addDevice(
            deviceClass='keyboard',
            deviceName='kbEnd',
        )
    # return True if completed successfully
    return True

def pauseExperiment(thisExp, win=None, timers=[], playbackComponents=[]):
    """
    Pause this experiment, preventing the flow from advancing to the next routine until resumed.
    
    Parameters
    ==========
    thisExp : psychopy.data.ExperimentHandler
        Handler object for this experiment, contains the data to save and information about 
        where to save it to.
    win : psychopy.visual.Window
        Window for this experiment.
    timers : list, tuple
        List of timers to reset once pausing is finished.
    playbackComponents : list, tuple
        List of any components with a `pause` method which need to be paused.
    """
    # if we are not paused, do nothing
    if thisExp.status != PAUSED:
        return
    
    # start a timer to figure out how long we're paused for
    pauseTimer = core.Clock()
    # pause any playback components
    for comp in playbackComponents:
        comp.pause()
    # make sure we have a keyboard
    defaultKeyboard = deviceManager.getDevice('defaultKeyboard')
    if defaultKeyboard is None:
        defaultKeyboard = deviceManager.addKeyboard(
            deviceClass='keyboard',
            deviceName='defaultKeyboard',
            backend='ioHub',
        )
    # run a while loop while we wait to unpause
    while thisExp.status == PAUSED:
        # check for quit (typically the Esc key)
        if defaultKeyboard.getKeys(keyList=['escape']):
            endExperiment(thisExp, win=win)
        # sleep 1ms so other threads can execute
        clock.time.sleep(0.001)
    # if stop was requested while paused, quit
    if thisExp.status == FINISHED:
        endExperiment(thisExp, win=win)
    # resume any playback components
    for comp in playbackComponents:
        comp.play()
    # reset any timers
    for timer in timers:
        timer.addTime(-pauseTimer.getTime())


def run(expInfo, thisExp, win, globalClock=None, thisSession=None):
    """
    Run the experiment flow.
    
    Parameters
    ==========
    expInfo : dict
        Information about this experiment, created by the `setupExpInfo` function.
    thisExp : psychopy.data.ExperimentHandler
        Handler object for this experiment, contains the data to save and information about 
        where to save it to.
    psychopy.visual.Window
        Window in which to run this experiment.
    globalClock : psychopy.core.clock.Clock or None
        Clock to get global time from - supply None to make a new one.
    thisSession : psychopy.session.Session or None
        Handle of the Session object this experiment is being run from, if any.
    """
    # mark experiment as started
    thisExp.status = STARTED
    # make sure window is set to foreground to prevent losing focus
    win.winHandle.activate()
    # make sure variables created by exec are available globally
    exec = environmenttools.setExecEnvironment(globals())
    # get device handles from dict of input devices
    ioServer = deviceManager.ioServer
    # get/create a default keyboard (e.g. to check for escape)
    defaultKeyboard = deviceManager.getDevice('defaultKeyboard')
    if defaultKeyboard is None:
        deviceManager.addDevice(
            deviceClass='keyboard', deviceName='defaultKeyboard', backend='ioHub'
        )
    eyetracker = deviceManager.getDevice('eyetracker')
    # make sure we're running in the directory for this experiment
    os.chdir(_thisDir)
    # get filename from ExperimentHandler for convenience
    filename = thisExp.dataFileName
    frameTolerance = 0.001  # how close to onset before 'same' frame
    endExpNow = False  # flag for 'escape' or other condition => quit the exp
    # get frame duration from frame rate in expInfo
    if 'frameRate' in expInfo and expInfo['frameRate'] is not None:
        frameDur = 1.0 / round(expInfo['frameRate'])
    else:
        frameDur = 1.0 / 60.0  # could not measure, so guess
    
    # Start Code - component code to be run after the window creation
    
    # --- Initialize components for Routine "Init_Group" ---
    # Run 'Begin Experiment' code from code
    # ==== Asignación balanceada por cohorte (64=16 por grupo), reproducible por semilla ====
    import json, random
    from pathlib import Path
    
    groups = ["Group8","Group16","Group24","Control"]
    target_trials_map = {"Group8":8, "Group16":16, "Group24":24, "Control":28}
    
    # Carpeta de datos (del archivo de este sujeto)
    data_dir = Path(thisExp.dataFileName).parent
    data_dir.mkdir(parents=True, exist_ok=True)
    
    # Archivos de asignación
    cfg_path = data_dir / "assign_cfg.json"   # guarda la semilla usada
    idx_path = data_dir / "assign_idx.txt"    # índice del siguiente sujeto (0,1,2,...)
    
    # Semilla: de expInfo["assign_seed"] si existe; si no, la guardada; si no, default
    try:
        cfg = json.loads(cfg_path.read_text(encoding="utf-8"))
    except Exception:
        cfg = {}
    seed = int(thisExp.extraInfo.get("assign_seed", cfg.get("seed", 123456789)))
    
    # Persistir semilla (auditoría)
    try:
        cfg_path.write_text(json.dumps({"seed": seed}, ensure_ascii=False, indent=2), encoding="utf-8")
    except Exception as e:
        print(f"[Assign] WARN: no se pudo escribir {cfg_path}: {e}")
    
    # Índice actual (si no existe, inicia en 0)
    try:
        next_idx = int(idx_path.read_text(encoding="utf-8").strip())
    except Exception:
        next_idx = 0
    
    # ----- Definir cohorte balanceada -----
    COHORT_SIZE = 64  # 16 por grupo con 4 grupos
    if COHORT_SIZE % len(groups) != 0:
        raise RuntimeError("COHORT_SIZE debe ser múltiplo del número de grupos.")
    COPIES_PER_GROUP = COHORT_SIZE // len(groups)
    
    # Construir mazo balanceado y barajar con la semilla
    deck = []
    for g in groups:
        deck.extend([g] * COPIES_PER_GROUP)
    
    rng = random.Random(seed)
    rng.shuffle(deck)
    
    # Posición en el mazo (cicla cada 64): si corres 128 sujetos, tendrás 32 por grupo, etc.
    pos = next_idx % COHORT_SIZE
    group = deck[pos]
    target_trials = target_trials_map[group]
    
    # Exponer para el resto del experimento
    thisExp.extraInfo["group"]                 = group
    thisExp.extraInfo["target_trials"]         = target_trials
    thisExp.extraInfo["assign_seed"]           = seed
    thisExp.extraInfo["assign_idx"]            = next_idx
    thisExp.extraInfo["assign_pos_in_deck"]    = pos
    thisExp.extraInfo["assign_cohort_size"]    = COHORT_SIZE
    
    print(f"[Assign] seed={seed}  idx={next_idx}  pos={pos}  group={group}  target_trials={target_trials}  cohort={COHORT_SIZE}")
    
    # (Opcional) Guardar el mazo para auditoría la primera vez
    try:
        deck_path = data_dir / f"assign_deck_{COHORT_SIZE}.txt"
        if not deck_path.exists():
            deck_path.write_text("\n".join(deck), encoding="utf-8")
    except Exception as e:
        print(f"[Assign] WARN: no se pudo escribir {deck_path}: {e}")
    
    
    # --- Initialize components for Routine "txtAudioCheck" ---
    txtAudioCheckk = visual.TextStim(win=win, name='txtAudioCheckk',
        text='Comprobación de audio\n\nSi escuchas un sonido, presiona espacio para avanzar\n\n(Si no escuchas nada, avisa al evaluador)',
        font='Arial',
        pos=(0, 0), draggable=False, height=0.05, wrapWidth=1.4, ori=0.0, 
        color='white', colorSpace='rgb', opacity=None, 
        languageStyle='LTR',
        depth=0.0);
    sndCheck = sound.Sound(
        'A', 
        secs=-1, 
        stereo=True, 
        hamming=True, 
        speaker='sndCheck',    name='sndCheck'
    )
    sndCheck.setVolume(0.35)
    kbAudioCheck = keyboard.Keyboard(deviceName='kbAudioCheck')
    
    # --- Initialize components for Routine "CI" ---
    image_5 = visual.ImageStim(
        win=win,
        name='image_5', 
        image='estimulos/consentimiento.jpg', mask=None, anchor='center',
        ori=0.0, pos=(0, 0), draggable=False, size=(0.7, 0.5),
        color=[1,1,1], colorSpace='rgb', opacity=None,
        flipHoriz=False, flipVert=False,
        texRes=128.0, interpolate=True, depth=0.0)
    key_resp_6 = keyboard.Keyboard(deviceName='key_resp_6')
    text_14 = visual.TextStim(win=win, name='text_14',
        text='Si das tu consentimiento, presiona la barra espaciadora. ',
        font='Arial',
        pos=(0, -0.45), draggable=False, height=0.03, wrapWidth=None, ori=0.0, 
        color='white', colorSpace='rgb', opacity=None, 
        languageStyle='LTR',
        depth=-2.0);
    
    # --- Initialize components for Routine "Welcome" ---
    txtWelcome = visual.TextStim(win=win, name='txtWelcome',
        text='Hola :) \n\nSolamente necesitarás utilizar: \n\n-El dinamómetro para apretar, y \n-La barra espaciadora para avanzar \n\nNADA MÁS. \n\nPara avanzar, presiona espacio.\n',
        font='Arial',
        pos=(0, 0), draggable=False, height=0.05, wrapWidth=1.5, ori=0.0, 
        color='white', colorSpace='rgb', opacity=None, 
        languageStyle='LTR',
        depth=0.0);
    kbWelcome = keyboard.Keyboard(deviceName='kbWelcome')
    
    # --- Initialize components for Routine "W2" ---
    text_4 = visual.TextStim(win=win, name='text_4',
        text='Manten agarrado el dinamómetro durante TODA la tarea.\n\nPuedes relajar la mano sin apretar, siempre y cuando lo sigas teniendo en la mano. \n\nNUNCA lo dejes sobre la mesa, ni en ninguna superficie hasta que el evaluador vuelva contigo.',
        font='Arial',
        pos=(0, 0.25), draggable=False, height=0.05, wrapWidth=None, ori=0.0, 
        color='white', colorSpace='rgb', opacity=None, 
        languageStyle='LTR',
        depth=0.0);
    key_resp_2 = keyboard.Keyboard(deviceName='key_resp_2')
    text_10 = visual.TextStim(win=win, name='text_10',
        text='Para avanzar, presiona espacio.',
        font='Arial',
        pos=(0, -0.45), draggable=False, height=0.05, wrapWidth=None, ori=0.0, 
        color='white', colorSpace='rgb', opacity=None, 
        languageStyle='LTR',
        depth=-2.0);
    image_4 = visual.ImageStim(
        win=win,
        name='image_4', 
        image='estimulos/tache soltar.png', mask=None, anchor='center',
        ori=0.0, pos=(0, -0.2), draggable=False, size=(0.4, 0.4),
        color=[1,1,1], colorSpace='rgb', opacity=None,
        flipHoriz=False, flipVert=False,
        texRes=128.0, interpolate=True, depth=-3.0)
    
    # --- Initialize components for Routine "Inst_Din1" ---
    text_2 = visual.TextStim(win=win, name='text_2',
        text='El dinamómetro medirá cuánta fuerza generas al apretarlo\n\nSostenlo siempre con tu mano dominante (con la que escribes)\n\nAgarra el dinamómetro por las DOS BARRAS AZULES largas\n\nPara avanzar, presiona espacio\n\n',
        font='Arial',
        pos=(0, -0.3), draggable=False, height=0.03, wrapWidth=1.7, ori=0.0, 
        color='white', colorSpace='rgb', opacity=None, 
        languageStyle='LTR',
        depth=0.0);
    kbInstrUso = keyboard.Keyboard(deviceName='kbInstrUso')
    image_2 = visual.ImageStim(
        win=win,
        name='image_2', 
        image='estimulos/palomita agarrar.png', mask=None, anchor='center',
        ori=0.0, pos=(0, 0.2), draggable=False, size=(0.40, 0.50),
        color=[1,1,1], colorSpace='rgb', opacity=None,
        flipHoriz=False, flipVert=False,
        texRes=128.0, interpolate=True, depth=-2.0)
    
    # --- Initialize components for Routine "circulos" ---
    image_3 = visual.ImageStim(
        win=win,
        name='image_3', 
        image='estimulos/tache circulos.png', mask=None, anchor='center',
        ori=0.0, pos=(0, 0.22), draggable=False, size=(0.40, 0.45),
        color=[1,1,1], colorSpace='rgb', opacity=None,
        flipHoriz=False, flipVert=False,
        texRes=128.0, interpolate=True, depth=0.0)
    key_resp_4 = keyboard.Keyboard(deviceName='key_resp_4')
    text_11 = visual.TextStim(win=win, name='text_11',
        text='Los círculos azules del dinamómetro NO sirven para nada\n\nPor favor, no los toques\n\nPara avanzar, presiona espacio',
        font='Arial',
        pos=(0, -0.2), draggable=False, height=0.05, wrapWidth=None, ori=0.0, 
        color='white', colorSpace='rgb', opacity=None, 
        languageStyle='LTR',
        depth=-2.0);
    
    # --- Initialize components for Routine "br" ---
    text_7 = visual.TextStim(win=win, name='text_7',
        text='Reposa tu brazo dominante sobre la mesa para que no se te canse al apretar el dinamómetro.\n\nCon la otra mano no sueltes nunca la pelota de tenis que se te dio. \n\nPara avanzar, presiona espacio. ',
        font='Arial',
        pos=(0, 0), draggable=False, height=0.05, wrapWidth=None, ori=0.0, 
        color='white', colorSpace='rgb', opacity=None, 
        languageStyle='LTR',
        depth=0.0);
    key_resp_3 = keyboard.Keyboard(deviceName='key_resp_3')
    
    # --- Initialize components for Routine "txtInstrCal" ---
    text = visual.TextStim(win=win, name='text',
        text='A continuación, te daremos intentos para que aprietes el dinamómetro con mucha fuerza. \n\nLa pantalla te irá guiando.\n\nEn cada intento, tendrás 5 segundos para apretar muy fuerte.\n\nPara avanzar, presiona espacio.\n',
        font='Arial',
        pos=(0, 0), draggable=False, height=0.05, wrapWidth=1.5, ori=0.0, 
        color='white', colorSpace='rgb', opacity=None, 
        languageStyle='LTR',
        depth=0.0);
    kbInstrCal = keyboard.Keyboard(deviceName='kbInstrCal')
    
    # --- Initialize components for Routine "RegCount" ---
    text_5 = visual.TextStim(win=win, name='text_5',
        text='Vas a apretar con mucha fuerza en...',
        font='Arial',
        pos=(0, 0), draggable=False, height=0.05, wrapWidth=None, ori=0.0, 
        color='white', colorSpace='rgb', opacity=None, 
        languageStyle='LTR',
        depth=0.0);
    text_6 = visual.TextStim(win=win, name='text_6',
        text='',
        font='Arial',
        pos=(0, -0.3), draggable=False, height=0.4, wrapWidth=None, ori=0.0, 
        color='white', colorSpace='rgb', opacity=None, 
        languageStyle='LTR',
        depth=-1.0);
    
    # --- Initialize components for Routine "Calibracion" ---
    # Run 'Begin Experiment' code from codeCal
    # --- Conexión Vernier ---
    from psychopy import core
    from gdx import gdx as gdx_mod
    
    gdx = gdx_mod.gdx()
    gdx.open_usb()
    gdx.select_sensors([1])   # dinamómetro
    gdx.start(period=10)      # ~100 Hz (antes estaba en 50 ≈ 20 Hz)
    
    # ====== Sampler no-bloqueante del Vernier (actualiza última lectura, robusto a timeouts) ======
    import threading
    try:
        from types import SimpleNamespace
    except Exception:
        class SimpleNamespace:
            def __init__(self, **kw): self.__dict__.update(kw)
    
    # Crear/usar estado global una sola vez
    try:
        _gdx_state
    except NameError:
        _gdx_state = SimpleNamespace(last=0.0, stop=False)
    
    def _gdx_sampler(state):
        while not state.stop:
            try:
                s = gdx.read()
                state.last = float(s[0]) if (s and s[0] is not None) else 0.0
            except Exception:
                # Si estamos parando, salimos silenciosamente; si no, cedemos CPU y seguimos
                if getattr(state, "stop", False):
                    break
                try:
                    core.wait(0.01)
                except Exception:
                    pass
    
    # Lanzar hilo una sola vez
    try:
        _gdx_thread
    except NameError:
        _gdx_thread = threading.Thread(target=_gdx_sampler, args=(_gdx_state,), daemon=True)
        _gdx_thread.start()
    
    # ====== Variables globales de calibración ======
    avg_f = 0.0
    thr70 = 0.0   # umbral ALTO (70% de avg_f)
    thr30 = 0.0   # umbral BAJO (30% de avg_f)
    
    # (Compatibilidad por si algún trozo viejo del script aún lee estas)
    thr60 = thr70
    thr40 = thr30
    
    txtCalibHeader = visual.TextStim(win=win, name='txtCalibHeader',
        text='Calibración',
        font='Arial',
        pos=(0, 0.45), draggable=False, height=0.05, wrapWidth=None, ori=0.0, 
        color='white', colorSpace='rgb', opacity=None, 
        languageStyle='LTR',
        depth=-1.0);
    txtCalibPrompt = visual.TextStim(win=win, name='txtCalibPrompt',
        text='¡Aprieta con tu máxima fuerza!',
        font='Times New Roman',
        pos=(0, 0.20), draggable=False, height=0.07, wrapWidth=None, ori=0.0, 
        color=[1.0000, 1.0000, 1.0000], colorSpace='rgb', opacity=None, 
        languageStyle='LTR',
        depth=-2.0);
    txtCountdown = visual.TextStim(win=win, name='txtCountdown',
        text='5.0 s',
        font='Arial',
        pos=(0, -0.10), draggable=False, height=0.05, wrapWidth=None, ori=0.0, 
        color='white', colorSpace='rgb', opacity=None, 
        languageStyle='LTR',
        depth=-3.0);
    txtAttemptMsg = visual.TextStim(win=win, name='txtAttemptMsg',
        text='Any text\n\nincluding line breaks',
        font='Arial',
        pos=(0, -0.35), draggable=False, height=0.05, wrapWidth=None, ori=0.0, 
        color='white', colorSpace='rgb', opacity=None, 
        languageStyle='LTR',
        depth=-4.0);
    dotRecord = visual.ShapeStim(
        win=win, name='dotRecord',
        size=(0.08, 0.08), vertices='circle',
        ori=0.0, pos=(0, 0.32), draggable=False, anchor='center',
        lineWidth=1.0,
        colorSpace='rgb', lineColor='white', fillColor=[0.9216, 0.9216, 0.7255],
        opacity=1.0, depth=-5.0, interpolate=True)
    
    # --- Initialize components for Routine "Instructions_FaseReforzamiento" ---
    txtInstrReinf = visual.TextStim(win=win, name='txtInstrReinf',
        text='A continuación,\n\nTendrás que descubrir cómo usar el dinamómetro para ganar dinero. \n\nEl dinero que logres acumular se te dará al final del experimento. \n\nMientras más dinero acumules, más estás apoyando a la ciencia. \n\nPara avanzar, presiona espacio.\n',
        font='Arial',
        pos=(0, 0), draggable=False, height=0.05, wrapWidth=1.5, ori=0.0, 
        color='white', colorSpace='rgb', opacity=None, 
        languageStyle='LTR',
        depth=-1.0);
    kbInstrReinf = keyboard.Keyboard(deviceName='kbInstrReinf')
    
    # --- Initialize components for Routine "OJO" ---
    text_3 = visual.TextStim(win=win, name='text_3',
        text='OJO, \n\nPara descubrir cómo ganar dinero usando el dinamómetro \n\nSOLO PUEDES USAR TU MANO DOMINANTE (NO LAS 2 MANOS)\n\nPara avanzar, presiona espacio. ',
        font='Arial',
        pos=(0, 0.3), draggable=False, height=0.05, wrapWidth=None, ori=0.0, 
        color='white', colorSpace='rgb', opacity=None, 
        languageStyle='LTR',
        depth=0.0);
    key_resp = keyboard.Keyboard(deviceName='key_resp')
    image = visual.ImageStim(
        win=win,
        name='image', 
        image='estimulos/tache dos manos.png', mask=None, anchor='center',
        ori=0.0, pos=(0, -0.2), draggable=False, size=(0.40, 0.45),
        color=[1,1,1], colorSpace='rgb', opacity=None,
        flipHoriz=False, flipVert=False,
        texRes=128.0, interpolate=True, depth=-2.0)
    
    # --- Initialize components for Routine "salir" ---
    text_13 = visual.TextStim(win=win, name='text_13',
        text='Espera a que el evaluador salga del cuarto y, cuando haya cerrado la puerta: \n\nPresiona espacio para avanzar.',
        font='Arial',
        pos=(0, 0), draggable=False, height=0.05, wrapWidth=None, ori=0.0, 
        color='white', colorSpace='rgb', opacity=None, 
        languageStyle='LTR',
        depth=0.0);
    key_resp_5 = keyboard.Keyboard(deviceName='key_resp_5')
    
    # --- Initialize components for Routine "Trial_Operant" ---
    imgCoin = visual.ImageStim(
        win=win,
        name='imgCoin', 
        image='estimulos/5pesos.png', mask=None, anchor='center',
        ori=0.0, pos=(0, 0), draggable=False, size=(0.10, 0.10),
        color=[1,1,1], colorSpace='rgb', opacity=1.0,
        flipHoriz=False, flipVert=False,
        texRes=128.0, interpolate=True, depth=-1.0)
    txtOperantMsg = visual.TextStim(win=win, name='txtOperantMsg',
        text='Listo\n¡Ya intenta ganar el mayor dinero posible!',
        font='Arial',
        pos=(0, 0.35), draggable=False, height=0.05, wrapWidth=None, ori=0.0, 
        color='white', colorSpace='rgb', opacity=None, 
        languageStyle='LTR',
        depth=-2.0);
    sndReinf = sound.Sound(
        'A', 
        secs=-1, 
        stereo=True, 
        hamming=True, 
        speaker='sndReinf',    name='sndReinf'
    )
    sndReinf.setVolume(1.0)
    text_9 = visual.TextStim(win=win, name='text_9',
        text='No te levantes de tu silla ni sueltes el dinamometro hasta que la pantalla cambie de color, o no recibirás nada de dinero. ',
        font='Arial',
        pos=(0, -0.4), draggable=False, height=0.03, wrapWidth=1.7, ori=0.0, 
        color='white', colorSpace='rgb', opacity=None, 
        languageStyle='LTR',
        depth=-4.0);
    
    # --- Initialize components for Routine "Extinction" ---
    txtExtMsg = visual.TextStim(win=win, name='txtExtMsg',
        text='Intenta ganar el mayor dinero posible',
        font='Arial',
        pos=(0, 0.35), draggable=False, height=0.05, wrapWidth=None, ori=0.0, 
        color='white', colorSpace='rgb', opacity=None, 
        languageStyle='LTR',
        depth=-1.0);
    text_12 = visual.TextStim(win=win, name='text_12',
        text='No te levantes de tu silla ni sueltes el dinamometro hasta que la pantalla cambie de color, o no recibirás nada de dinero. ',
        font='Arial',
        pos=(0, -0.4), draggable=False, height=0.03, wrapWidth=1.7, ori=0.0, 
        color='white', colorSpace='rgb', opacity=None, 
        languageStyle='LTR',
        depth=-2.0);
    
    # --- Initialize components for Routine "End" ---
    txtSummary = visual.TextStim(win=win, name='txtSummary',
        text=None,
        font='Arial',
        pos=(0, 0.2), draggable=False, height=0.05, wrapWidth=None, ori=0.0, 
        color='white', colorSpace='rgb', opacity=None, 
        languageStyle='LTR',
        depth=-1.0);
    txtThanks = visual.TextStim(win=win, name='txtThanks',
        text=None,
        font='Arial',
        pos=(0, -0.2), draggable=False, height=0.05, wrapWidth=None, ori=0.0, 
        color='white', colorSpace='rgb', opacity=None, 
        languageStyle='LTR',
        depth=-2.0);
    kbEnd = keyboard.Keyboard(deviceName='kbEnd')
    
    # create some handy timers
    
    # global clock to track the time since experiment started
    if globalClock is None:
        # create a clock if not given one
        globalClock = core.Clock()
    if isinstance(globalClock, str):
        # if given a string, make a clock accoridng to it
        if globalClock == 'float':
            # get timestamps as a simple value
            globalClock = core.Clock(format='float')
        elif globalClock == 'iso':
            # get timestamps in ISO format
            globalClock = core.Clock(format='%Y-%m-%d_%H:%M:%S.%f%z')
        else:
            # get timestamps in a custom format
            globalClock = core.Clock(format=globalClock)
    if ioServer is not None:
        ioServer.syncClock(globalClock)
    logging.setDefaultClock(globalClock)
    # routine timer to track time remaining of each (possibly non-slip) routine
    routineTimer = core.Clock()
    win.flip()  # flip window to reset last flip timer
    # store the exact time the global clock started
    expInfo['expStart'] = data.getDateStr(
        format='%Y-%m-%d %Hh%M.%S.%f %z', fractionalSecondDigits=6
    )
    
    # --- Prepare to start Routine "Init_Group" ---
    # create an object to store info about Routine Init_Group
    Init_Group = data.Routine(
        name='Init_Group',
        components=[],
    )
    Init_Group.status = NOT_STARTED
    continueRoutine = True
    # update component parameters for each repeat
    # store start times for Init_Group
    Init_Group.tStartRefresh = win.getFutureFlipTime(clock=globalClock)
    Init_Group.tStart = globalClock.getTime(format='float')
    Init_Group.status = STARTED
    thisExp.addData('Init_Group.started', Init_Group.tStart)
    Init_Group.maxDuration = None
    # keep track of which components have finished
    Init_GroupComponents = Init_Group.components
    for thisComponent in Init_Group.components:
        thisComponent.tStart = None
        thisComponent.tStop = None
        thisComponent.tStartRefresh = None
        thisComponent.tStopRefresh = None
        if hasattr(thisComponent, 'status'):
            thisComponent.status = NOT_STARTED
    # reset timers
    t = 0
    _timeToFirstFrame = win.getFutureFlipTime(clock="now")
    frameN = -1
    
    # --- Run Routine "Init_Group" ---
    Init_Group.forceEnded = routineForceEnded = not continueRoutine
    while continueRoutine:
        # get current time
        t = routineTimer.getTime()
        tThisFlip = win.getFutureFlipTime(clock=routineTimer)
        tThisFlipGlobal = win.getFutureFlipTime(clock=None)
        frameN = frameN + 1  # number of completed frames (so 0 is the first frame)
        # update/draw components on each frame
        
        # check for quit (typically the Esc key)
        if defaultKeyboard.getKeys(keyList=["escape"]):
            thisExp.status = FINISHED
        if thisExp.status == FINISHED or endExpNow:
            endExperiment(thisExp, win=win)
            return
        # pause experiment here if requested
        if thisExp.status == PAUSED:
            pauseExperiment(
                thisExp=thisExp, 
                win=win, 
                timers=[routineTimer], 
                playbackComponents=[]
            )
            # skip the frame we paused on
            continue
        
        # check if all components have finished
        if not continueRoutine:  # a component has requested a forced-end of Routine
            Init_Group.forceEnded = routineForceEnded = True
            break
        continueRoutine = False  # will revert to True if at least one component still running
        for thisComponent in Init_Group.components:
            if hasattr(thisComponent, "status") and thisComponent.status != FINISHED:
                continueRoutine = True
                break  # at least one component has not yet finished
        
        # refresh the screen
        if continueRoutine:  # don't flip if this routine is over or we'll get a blank screen
            win.flip()
    
    # --- Ending Routine "Init_Group" ---
    for thisComponent in Init_Group.components:
        if hasattr(thisComponent, "setAutoDraw"):
            thisComponent.setAutoDraw(False)
    # store stop times for Init_Group
    Init_Group.tStop = globalClock.getTime(format='float')
    Init_Group.tStopRefresh = tThisFlipGlobal
    thisExp.addData('Init_Group.stopped', Init_Group.tStop)
    thisExp.nextEntry()
    # the Routine "Init_Group" was not non-slip safe, so reset the non-slip timer
    routineTimer.reset()
    
    # --- Prepare to start Routine "txtAudioCheck" ---
    # create an object to store info about Routine txtAudioCheck
    txtAudioCheck = data.Routine(
        name='txtAudioCheck',
        components=[txtAudioCheckk, sndCheck, kbAudioCheck],
    )
    txtAudioCheck.status = NOT_STARTED
    continueRoutine = True
    # update component parameters for each repeat
    sndCheck.setSound('440', secs=20, hamming=True)
    sndCheck.setVolume(0.35, log=False)
    sndCheck.seek(0)
    # create starting attributes for kbAudioCheck
    kbAudioCheck.keys = []
    kbAudioCheck.rt = []
    _kbAudioCheck_allKeys = []
    # store start times for txtAudioCheck
    txtAudioCheck.tStartRefresh = win.getFutureFlipTime(clock=globalClock)
    txtAudioCheck.tStart = globalClock.getTime(format='float')
    txtAudioCheck.status = STARTED
    thisExp.addData('txtAudioCheck.started', txtAudioCheck.tStart)
    txtAudioCheck.maxDuration = None
    # keep track of which components have finished
    txtAudioCheckComponents = txtAudioCheck.components
    for thisComponent in txtAudioCheck.components:
        thisComponent.tStart = None
        thisComponent.tStop = None
        thisComponent.tStartRefresh = None
        thisComponent.tStopRefresh = None
        if hasattr(thisComponent, 'status'):
            thisComponent.status = NOT_STARTED
    # reset timers
    t = 0
    _timeToFirstFrame = win.getFutureFlipTime(clock="now")
    frameN = -1
    
    # --- Run Routine "txtAudioCheck" ---
    txtAudioCheck.forceEnded = routineForceEnded = not continueRoutine
    while continueRoutine:
        # get current time
        t = routineTimer.getTime()
        tThisFlip = win.getFutureFlipTime(clock=routineTimer)
        tThisFlipGlobal = win.getFutureFlipTime(clock=None)
        frameN = frameN + 1  # number of completed frames (so 0 is the first frame)
        # update/draw components on each frame
        
        # *txtAudioCheckk* updates
        
        # if txtAudioCheckk is starting this frame...
        if txtAudioCheckk.status == NOT_STARTED and tThisFlip >= 0.0-frameTolerance:
            # keep track of start time/frame for later
            txtAudioCheckk.frameNStart = frameN  # exact frame index
            txtAudioCheckk.tStart = t  # local t and not account for scr refresh
            txtAudioCheckk.tStartRefresh = tThisFlipGlobal  # on global time
            win.timeOnFlip(txtAudioCheckk, 'tStartRefresh')  # time at next scr refresh
            # add timestamp to datafile
            thisExp.timestampOnFlip(win, 'txtAudioCheckk.started')
            # update status
            txtAudioCheckk.status = STARTED
            txtAudioCheckk.setAutoDraw(True)
        
        # if txtAudioCheckk is active this frame...
        if txtAudioCheckk.status == STARTED:
            # update params
            pass
        
        # *sndCheck* updates
        
        # if sndCheck is starting this frame...
        if sndCheck.status == NOT_STARTED and tThisFlip >= 0.0-frameTolerance:
            # keep track of start time/frame for later
            sndCheck.frameNStart = frameN  # exact frame index
            sndCheck.tStart = t  # local t and not account for scr refresh
            sndCheck.tStartRefresh = tThisFlipGlobal  # on global time
            # add timestamp to datafile
            thisExp.addData('sndCheck.started', tThisFlipGlobal)
            # update status
            sndCheck.status = STARTED
            sndCheck.play(when=win)  # sync with win flip
        
        # if sndCheck is stopping this frame...
        if sndCheck.status == STARTED:
            # is it time to stop? (based on global clock, using actual start)
            if tThisFlipGlobal > sndCheck.tStartRefresh + 20-frameTolerance or sndCheck.isFinished:
                # keep track of stop time/frame for later
                sndCheck.tStop = t  # not accounting for scr refresh
                sndCheck.tStopRefresh = tThisFlipGlobal  # on global time
                sndCheck.frameNStop = frameN  # exact frame index
                # add timestamp to datafile
                thisExp.timestampOnFlip(win, 'sndCheck.stopped')
                # update status
                sndCheck.status = FINISHED
                sndCheck.stop()
        
        # *kbAudioCheck* updates
        waitOnFlip = False
        
        # if kbAudioCheck is starting this frame...
        if kbAudioCheck.status == NOT_STARTED and tThisFlip >= 0.0-frameTolerance:
            # keep track of start time/frame for later
            kbAudioCheck.frameNStart = frameN  # exact frame index
            kbAudioCheck.tStart = t  # local t and not account for scr refresh
            kbAudioCheck.tStartRefresh = tThisFlipGlobal  # on global time
            win.timeOnFlip(kbAudioCheck, 'tStartRefresh')  # time at next scr refresh
            # add timestamp to datafile
            thisExp.timestampOnFlip(win, 'kbAudioCheck.started')
            # update status
            kbAudioCheck.status = STARTED
            # keyboard checking is just starting
            waitOnFlip = True
            win.callOnFlip(kbAudioCheck.clock.reset)  # t=0 on next screen flip
            win.callOnFlip(kbAudioCheck.clearEvents, eventType='keyboard')  # clear events on next screen flip
        if kbAudioCheck.status == STARTED and not waitOnFlip:
            theseKeys = kbAudioCheck.getKeys(keyList=['space'], ignoreKeys=["escape"], waitRelease=False)
            _kbAudioCheck_allKeys.extend(theseKeys)
            if len(_kbAudioCheck_allKeys):
                kbAudioCheck.keys = _kbAudioCheck_allKeys[-1].name  # just the last key pressed
                kbAudioCheck.rt = _kbAudioCheck_allKeys[-1].rt
                kbAudioCheck.duration = _kbAudioCheck_allKeys[-1].duration
                # a response ends the routine
                continueRoutine = False
        
        # check for quit (typically the Esc key)
        if defaultKeyboard.getKeys(keyList=["escape"]):
            thisExp.status = FINISHED
        if thisExp.status == FINISHED or endExpNow:
            endExperiment(thisExp, win=win)
            return
        # pause experiment here if requested
        if thisExp.status == PAUSED:
            pauseExperiment(
                thisExp=thisExp, 
                win=win, 
                timers=[routineTimer], 
                playbackComponents=[sndCheck]
            )
            # skip the frame we paused on
            continue
        
        # check if all components have finished
        if not continueRoutine:  # a component has requested a forced-end of Routine
            txtAudioCheck.forceEnded = routineForceEnded = True
            break
        continueRoutine = False  # will revert to True if at least one component still running
        for thisComponent in txtAudioCheck.components:
            if hasattr(thisComponent, "status") and thisComponent.status != FINISHED:
                continueRoutine = True
                break  # at least one component has not yet finished
        
        # refresh the screen
        if continueRoutine:  # don't flip if this routine is over or we'll get a blank screen
            win.flip()
    
    # --- Ending Routine "txtAudioCheck" ---
    for thisComponent in txtAudioCheck.components:
        if hasattr(thisComponent, "setAutoDraw"):
            thisComponent.setAutoDraw(False)
    # store stop times for txtAudioCheck
    txtAudioCheck.tStop = globalClock.getTime(format='float')
    txtAudioCheck.tStopRefresh = tThisFlipGlobal
    thisExp.addData('txtAudioCheck.stopped', txtAudioCheck.tStop)
    sndCheck.pause()  # ensure sound has stopped at end of Routine
    # check responses
    if kbAudioCheck.keys in ['', [], None]:  # No response was made
        kbAudioCheck.keys = None
    thisExp.addData('kbAudioCheck.keys',kbAudioCheck.keys)
    if kbAudioCheck.keys != None:  # we had a response
        thisExp.addData('kbAudioCheck.rt', kbAudioCheck.rt)
        thisExp.addData('kbAudioCheck.duration', kbAudioCheck.duration)
    thisExp.nextEntry()
    # the Routine "txtAudioCheck" was not non-slip safe, so reset the non-slip timer
    routineTimer.reset()
    
    # --- Prepare to start Routine "CI" ---
    # create an object to store info about Routine CI
    CI = data.Routine(
        name='CI',
        components=[image_5, key_resp_6, text_14],
    )
    CI.status = NOT_STARTED
    continueRoutine = True
    # update component parameters for each repeat
    # create starting attributes for key_resp_6
    key_resp_6.keys = []
    key_resp_6.rt = []
    _key_resp_6_allKeys = []
    # store start times for CI
    CI.tStartRefresh = win.getFutureFlipTime(clock=globalClock)
    CI.tStart = globalClock.getTime(format='float')
    CI.status = STARTED
    thisExp.addData('CI.started', CI.tStart)
    CI.maxDuration = None
    # keep track of which components have finished
    CIComponents = CI.components
    for thisComponent in CI.components:
        thisComponent.tStart = None
        thisComponent.tStop = None
        thisComponent.tStartRefresh = None
        thisComponent.tStopRefresh = None
        if hasattr(thisComponent, 'status'):
            thisComponent.status = NOT_STARTED
    # reset timers
    t = 0
    _timeToFirstFrame = win.getFutureFlipTime(clock="now")
    frameN = -1
    
    # --- Run Routine "CI" ---
    CI.forceEnded = routineForceEnded = not continueRoutine
    while continueRoutine:
        # get current time
        t = routineTimer.getTime()
        tThisFlip = win.getFutureFlipTime(clock=routineTimer)
        tThisFlipGlobal = win.getFutureFlipTime(clock=None)
        frameN = frameN + 1  # number of completed frames (so 0 is the first frame)
        # update/draw components on each frame
        
        # *image_5* updates
        
        # if image_5 is starting this frame...
        if image_5.status == NOT_STARTED and tThisFlip >= 0.0-frameTolerance:
            # keep track of start time/frame for later
            image_5.frameNStart = frameN  # exact frame index
            image_5.tStart = t  # local t and not account for scr refresh
            image_5.tStartRefresh = tThisFlipGlobal  # on global time
            win.timeOnFlip(image_5, 'tStartRefresh')  # time at next scr refresh
            # add timestamp to datafile
            thisExp.timestampOnFlip(win, 'image_5.started')
            # update status
            image_5.status = STARTED
            image_5.setAutoDraw(True)
        
        # if image_5 is active this frame...
        if image_5.status == STARTED:
            # update params
            pass
        
        # *key_resp_6* updates
        waitOnFlip = False
        
        # if key_resp_6 is starting this frame...
        if key_resp_6.status == NOT_STARTED and tThisFlip >= 0.0-frameTolerance:
            # keep track of start time/frame for later
            key_resp_6.frameNStart = frameN  # exact frame index
            key_resp_6.tStart = t  # local t and not account for scr refresh
            key_resp_6.tStartRefresh = tThisFlipGlobal  # on global time
            win.timeOnFlip(key_resp_6, 'tStartRefresh')  # time at next scr refresh
            # add timestamp to datafile
            thisExp.timestampOnFlip(win, 'key_resp_6.started')
            # update status
            key_resp_6.status = STARTED
            # keyboard checking is just starting
            waitOnFlip = True
            win.callOnFlip(key_resp_6.clock.reset)  # t=0 on next screen flip
            win.callOnFlip(key_resp_6.clearEvents, eventType='keyboard')  # clear events on next screen flip
        if key_resp_6.status == STARTED and not waitOnFlip:
            theseKeys = key_resp_6.getKeys(keyList=['space'], ignoreKeys=["escape"], waitRelease=False)
            _key_resp_6_allKeys.extend(theseKeys)
            if len(_key_resp_6_allKeys):
                key_resp_6.keys = _key_resp_6_allKeys[-1].name  # just the last key pressed
                key_resp_6.rt = _key_resp_6_allKeys[-1].rt
                key_resp_6.duration = _key_resp_6_allKeys[-1].duration
                # a response ends the routine
                continueRoutine = False
        
        # *text_14* updates
        
        # if text_14 is starting this frame...
        if text_14.status == NOT_STARTED and tThisFlip >= 0.0-frameTolerance:
            # keep track of start time/frame for later
            text_14.frameNStart = frameN  # exact frame index
            text_14.tStart = t  # local t and not account for scr refresh
            text_14.tStartRefresh = tThisFlipGlobal  # on global time
            win.timeOnFlip(text_14, 'tStartRefresh')  # time at next scr refresh
            # add timestamp to datafile
            thisExp.timestampOnFlip(win, 'text_14.started')
            # update status
            text_14.status = STARTED
            text_14.setAutoDraw(True)
        
        # if text_14 is active this frame...
        if text_14.status == STARTED:
            # update params
            pass
        
        # check for quit (typically the Esc key)
        if defaultKeyboard.getKeys(keyList=["escape"]):
            thisExp.status = FINISHED
        if thisExp.status == FINISHED or endExpNow:
            endExperiment(thisExp, win=win)
            return
        # pause experiment here if requested
        if thisExp.status == PAUSED:
            pauseExperiment(
                thisExp=thisExp, 
                win=win, 
                timers=[routineTimer], 
                playbackComponents=[]
            )
            # skip the frame we paused on
            continue
        
        # check if all components have finished
        if not continueRoutine:  # a component has requested a forced-end of Routine
            CI.forceEnded = routineForceEnded = True
            break
        continueRoutine = False  # will revert to True if at least one component still running
        for thisComponent in CI.components:
            if hasattr(thisComponent, "status") and thisComponent.status != FINISHED:
                continueRoutine = True
                break  # at least one component has not yet finished
        
        # refresh the screen
        if continueRoutine:  # don't flip if this routine is over or we'll get a blank screen
            win.flip()
    
    # --- Ending Routine "CI" ---
    for thisComponent in CI.components:
        if hasattr(thisComponent, "setAutoDraw"):
            thisComponent.setAutoDraw(False)
    # store stop times for CI
    CI.tStop = globalClock.getTime(format='float')
    CI.tStopRefresh = tThisFlipGlobal
    thisExp.addData('CI.stopped', CI.tStop)
    # check responses
    if key_resp_6.keys in ['', [], None]:  # No response was made
        key_resp_6.keys = None
    thisExp.addData('key_resp_6.keys',key_resp_6.keys)
    if key_resp_6.keys != None:  # we had a response
        thisExp.addData('key_resp_6.rt', key_resp_6.rt)
        thisExp.addData('key_resp_6.duration', key_resp_6.duration)
    thisExp.nextEntry()
    # the Routine "CI" was not non-slip safe, so reset the non-slip timer
    routineTimer.reset()
    
    # --- Prepare to start Routine "Welcome" ---
    # create an object to store info about Routine Welcome
    Welcome = data.Routine(
        name='Welcome',
        components=[txtWelcome, kbWelcome],
    )
    Welcome.status = NOT_STARTED
    continueRoutine = True
    # update component parameters for each repeat
    # create starting attributes for kbWelcome
    kbWelcome.keys = []
    kbWelcome.rt = []
    _kbWelcome_allKeys = []
    # store start times for Welcome
    Welcome.tStartRefresh = win.getFutureFlipTime(clock=globalClock)
    Welcome.tStart = globalClock.getTime(format='float')
    Welcome.status = STARTED
    thisExp.addData('Welcome.started', Welcome.tStart)
    Welcome.maxDuration = None
    # keep track of which components have finished
    WelcomeComponents = Welcome.components
    for thisComponent in Welcome.components:
        thisComponent.tStart = None
        thisComponent.tStop = None
        thisComponent.tStartRefresh = None
        thisComponent.tStopRefresh = None
        if hasattr(thisComponent, 'status'):
            thisComponent.status = NOT_STARTED
    # reset timers
    t = 0
    _timeToFirstFrame = win.getFutureFlipTime(clock="now")
    frameN = -1
    
    # --- Run Routine "Welcome" ---
    Welcome.forceEnded = routineForceEnded = not continueRoutine
    while continueRoutine:
        # get current time
        t = routineTimer.getTime()
        tThisFlip = win.getFutureFlipTime(clock=routineTimer)
        tThisFlipGlobal = win.getFutureFlipTime(clock=None)
        frameN = frameN + 1  # number of completed frames (so 0 is the first frame)
        # update/draw components on each frame
        
        # *txtWelcome* updates
        
        # if txtWelcome is starting this frame...
        if txtWelcome.status == NOT_STARTED and tThisFlip >= 0.0-frameTolerance:
            # keep track of start time/frame for later
            txtWelcome.frameNStart = frameN  # exact frame index
            txtWelcome.tStart = t  # local t and not account for scr refresh
            txtWelcome.tStartRefresh = tThisFlipGlobal  # on global time
            win.timeOnFlip(txtWelcome, 'tStartRefresh')  # time at next scr refresh
            # add timestamp to datafile
            thisExp.timestampOnFlip(win, 'txtWelcome.started')
            # update status
            txtWelcome.status = STARTED
            txtWelcome.setAutoDraw(True)
        
        # if txtWelcome is active this frame...
        if txtWelcome.status == STARTED:
            # update params
            pass
        
        # *kbWelcome* updates
        waitOnFlip = False
        
        # if kbWelcome is starting this frame...
        if kbWelcome.status == NOT_STARTED and tThisFlip >= 0.0-frameTolerance:
            # keep track of start time/frame for later
            kbWelcome.frameNStart = frameN  # exact frame index
            kbWelcome.tStart = t  # local t and not account for scr refresh
            kbWelcome.tStartRefresh = tThisFlipGlobal  # on global time
            win.timeOnFlip(kbWelcome, 'tStartRefresh')  # time at next scr refresh
            # add timestamp to datafile
            thisExp.timestampOnFlip(win, 'kbWelcome.started')
            # update status
            kbWelcome.status = STARTED
            # keyboard checking is just starting
            waitOnFlip = True
            win.callOnFlip(kbWelcome.clock.reset)  # t=0 on next screen flip
            win.callOnFlip(kbWelcome.clearEvents, eventType='keyboard')  # clear events on next screen flip
        if kbWelcome.status == STARTED and not waitOnFlip:
            theseKeys = kbWelcome.getKeys(keyList=['space'], ignoreKeys=["escape"], waitRelease=False)
            _kbWelcome_allKeys.extend(theseKeys)
            if len(_kbWelcome_allKeys):
                kbWelcome.keys = _kbWelcome_allKeys[-1].name  # just the last key pressed
                kbWelcome.rt = _kbWelcome_allKeys[-1].rt
                kbWelcome.duration = _kbWelcome_allKeys[-1].duration
                # a response ends the routine
                continueRoutine = False
        
        # check for quit (typically the Esc key)
        if defaultKeyboard.getKeys(keyList=["escape"]):
            thisExp.status = FINISHED
        if thisExp.status == FINISHED or endExpNow:
            endExperiment(thisExp, win=win)
            return
        # pause experiment here if requested
        if thisExp.status == PAUSED:
            pauseExperiment(
                thisExp=thisExp, 
                win=win, 
                timers=[routineTimer], 
                playbackComponents=[]
            )
            # skip the frame we paused on
            continue
        
        # check if all components have finished
        if not continueRoutine:  # a component has requested a forced-end of Routine
            Welcome.forceEnded = routineForceEnded = True
            break
        continueRoutine = False  # will revert to True if at least one component still running
        for thisComponent in Welcome.components:
            if hasattr(thisComponent, "status") and thisComponent.status != FINISHED:
                continueRoutine = True
                break  # at least one component has not yet finished
        
        # refresh the screen
        if continueRoutine:  # don't flip if this routine is over or we'll get a blank screen
            win.flip()
    
    # --- Ending Routine "Welcome" ---
    for thisComponent in Welcome.components:
        if hasattr(thisComponent, "setAutoDraw"):
            thisComponent.setAutoDraw(False)
    # store stop times for Welcome
    Welcome.tStop = globalClock.getTime(format='float')
    Welcome.tStopRefresh = tThisFlipGlobal
    thisExp.addData('Welcome.stopped', Welcome.tStop)
    # check responses
    if kbWelcome.keys in ['', [], None]:  # No response was made
        kbWelcome.keys = None
    thisExp.addData('kbWelcome.keys',kbWelcome.keys)
    if kbWelcome.keys != None:  # we had a response
        thisExp.addData('kbWelcome.rt', kbWelcome.rt)
        thisExp.addData('kbWelcome.duration', kbWelcome.duration)
    thisExp.nextEntry()
    # the Routine "Welcome" was not non-slip safe, so reset the non-slip timer
    routineTimer.reset()
    
    # --- Prepare to start Routine "W2" ---
    # create an object to store info about Routine W2
    W2 = data.Routine(
        name='W2',
        components=[text_4, key_resp_2, text_10, image_4],
    )
    W2.status = NOT_STARTED
    continueRoutine = True
    # update component parameters for each repeat
    # create starting attributes for key_resp_2
    key_resp_2.keys = []
    key_resp_2.rt = []
    _key_resp_2_allKeys = []
    # store start times for W2
    W2.tStartRefresh = win.getFutureFlipTime(clock=globalClock)
    W2.tStart = globalClock.getTime(format='float')
    W2.status = STARTED
    thisExp.addData('W2.started', W2.tStart)
    W2.maxDuration = None
    # keep track of which components have finished
    W2Components = W2.components
    for thisComponent in W2.components:
        thisComponent.tStart = None
        thisComponent.tStop = None
        thisComponent.tStartRefresh = None
        thisComponent.tStopRefresh = None
        if hasattr(thisComponent, 'status'):
            thisComponent.status = NOT_STARTED
    # reset timers
    t = 0
    _timeToFirstFrame = win.getFutureFlipTime(clock="now")
    frameN = -1
    
    # --- Run Routine "W2" ---
    W2.forceEnded = routineForceEnded = not continueRoutine
    while continueRoutine:
        # get current time
        t = routineTimer.getTime()
        tThisFlip = win.getFutureFlipTime(clock=routineTimer)
        tThisFlipGlobal = win.getFutureFlipTime(clock=None)
        frameN = frameN + 1  # number of completed frames (so 0 is the first frame)
        # update/draw components on each frame
        
        # *text_4* updates
        
        # if text_4 is starting this frame...
        if text_4.status == NOT_STARTED and tThisFlip >= 0.0-frameTolerance:
            # keep track of start time/frame for later
            text_4.frameNStart = frameN  # exact frame index
            text_4.tStart = t  # local t and not account for scr refresh
            text_4.tStartRefresh = tThisFlipGlobal  # on global time
            win.timeOnFlip(text_4, 'tStartRefresh')  # time at next scr refresh
            # add timestamp to datafile
            thisExp.timestampOnFlip(win, 'text_4.started')
            # update status
            text_4.status = STARTED
            text_4.setAutoDraw(True)
        
        # if text_4 is active this frame...
        if text_4.status == STARTED:
            # update params
            pass
        
        # *key_resp_2* updates
        waitOnFlip = False
        
        # if key_resp_2 is starting this frame...
        if key_resp_2.status == NOT_STARTED and tThisFlip >= 0.0-frameTolerance:
            # keep track of start time/frame for later
            key_resp_2.frameNStart = frameN  # exact frame index
            key_resp_2.tStart = t  # local t and not account for scr refresh
            key_resp_2.tStartRefresh = tThisFlipGlobal  # on global time
            win.timeOnFlip(key_resp_2, 'tStartRefresh')  # time at next scr refresh
            # add timestamp to datafile
            thisExp.timestampOnFlip(win, 'key_resp_2.started')
            # update status
            key_resp_2.status = STARTED
            # keyboard checking is just starting
            waitOnFlip = True
            win.callOnFlip(key_resp_2.clock.reset)  # t=0 on next screen flip
            win.callOnFlip(key_resp_2.clearEvents, eventType='keyboard')  # clear events on next screen flip
        if key_resp_2.status == STARTED and not waitOnFlip:
            theseKeys = key_resp_2.getKeys(keyList=['space'], ignoreKeys=["escape"], waitRelease=False)
            _key_resp_2_allKeys.extend(theseKeys)
            if len(_key_resp_2_allKeys):
                key_resp_2.keys = _key_resp_2_allKeys[-1].name  # just the last key pressed
                key_resp_2.rt = _key_resp_2_allKeys[-1].rt
                key_resp_2.duration = _key_resp_2_allKeys[-1].duration
                # a response ends the routine
                continueRoutine = False
        
        # *text_10* updates
        
        # if text_10 is starting this frame...
        if text_10.status == NOT_STARTED and tThisFlip >= 0.0-frameTolerance:
            # keep track of start time/frame for later
            text_10.frameNStart = frameN  # exact frame index
            text_10.tStart = t  # local t and not account for scr refresh
            text_10.tStartRefresh = tThisFlipGlobal  # on global time
            win.timeOnFlip(text_10, 'tStartRefresh')  # time at next scr refresh
            # add timestamp to datafile
            thisExp.timestampOnFlip(win, 'text_10.started')
            # update status
            text_10.status = STARTED
            text_10.setAutoDraw(True)
        
        # if text_10 is active this frame...
        if text_10.status == STARTED:
            # update params
            pass
        
        # *image_4* updates
        
        # if image_4 is starting this frame...
        if image_4.status == NOT_STARTED and tThisFlip >= 0.0-frameTolerance:
            # keep track of start time/frame for later
            image_4.frameNStart = frameN  # exact frame index
            image_4.tStart = t  # local t and not account for scr refresh
            image_4.tStartRefresh = tThisFlipGlobal  # on global time
            win.timeOnFlip(image_4, 'tStartRefresh')  # time at next scr refresh
            # add timestamp to datafile
            thisExp.timestampOnFlip(win, 'image_4.started')
            # update status
            image_4.status = STARTED
            image_4.setAutoDraw(True)
        
        # if image_4 is active this frame...
        if image_4.status == STARTED:
            # update params
            pass
        
        # check for quit (typically the Esc key)
        if defaultKeyboard.getKeys(keyList=["escape"]):
            thisExp.status = FINISHED
        if thisExp.status == FINISHED or endExpNow:
            endExperiment(thisExp, win=win)
            return
        # pause experiment here if requested
        if thisExp.status == PAUSED:
            pauseExperiment(
                thisExp=thisExp, 
                win=win, 
                timers=[routineTimer], 
                playbackComponents=[]
            )
            # skip the frame we paused on
            continue
        
        # check if all components have finished
        if not continueRoutine:  # a component has requested a forced-end of Routine
            W2.forceEnded = routineForceEnded = True
            break
        continueRoutine = False  # will revert to True if at least one component still running
        for thisComponent in W2.components:
            if hasattr(thisComponent, "status") and thisComponent.status != FINISHED:
                continueRoutine = True
                break  # at least one component has not yet finished
        
        # refresh the screen
        if continueRoutine:  # don't flip if this routine is over or we'll get a blank screen
            win.flip()
    
    # --- Ending Routine "W2" ---
    for thisComponent in W2.components:
        if hasattr(thisComponent, "setAutoDraw"):
            thisComponent.setAutoDraw(False)
    # store stop times for W2
    W2.tStop = globalClock.getTime(format='float')
    W2.tStopRefresh = tThisFlipGlobal
    thisExp.addData('W2.stopped', W2.tStop)
    # check responses
    if key_resp_2.keys in ['', [], None]:  # No response was made
        key_resp_2.keys = None
    thisExp.addData('key_resp_2.keys',key_resp_2.keys)
    if key_resp_2.keys != None:  # we had a response
        thisExp.addData('key_resp_2.rt', key_resp_2.rt)
        thisExp.addData('key_resp_2.duration', key_resp_2.duration)
    thisExp.nextEntry()
    # the Routine "W2" was not non-slip safe, so reset the non-slip timer
    routineTimer.reset()
    
    # --- Prepare to start Routine "Inst_Din1" ---
    # create an object to store info about Routine Inst_Din1
    Inst_Din1 = data.Routine(
        name='Inst_Din1',
        components=[text_2, kbInstrUso, image_2],
    )
    Inst_Din1.status = NOT_STARTED
    continueRoutine = True
    # update component parameters for each repeat
    # create starting attributes for kbInstrUso
    kbInstrUso.keys = []
    kbInstrUso.rt = []
    _kbInstrUso_allKeys = []
    # store start times for Inst_Din1
    Inst_Din1.tStartRefresh = win.getFutureFlipTime(clock=globalClock)
    Inst_Din1.tStart = globalClock.getTime(format='float')
    Inst_Din1.status = STARTED
    thisExp.addData('Inst_Din1.started', Inst_Din1.tStart)
    Inst_Din1.maxDuration = None
    # keep track of which components have finished
    Inst_Din1Components = Inst_Din1.components
    for thisComponent in Inst_Din1.components:
        thisComponent.tStart = None
        thisComponent.tStop = None
        thisComponent.tStartRefresh = None
        thisComponent.tStopRefresh = None
        if hasattr(thisComponent, 'status'):
            thisComponent.status = NOT_STARTED
    # reset timers
    t = 0
    _timeToFirstFrame = win.getFutureFlipTime(clock="now")
    frameN = -1
    
    # --- Run Routine "Inst_Din1" ---
    Inst_Din1.forceEnded = routineForceEnded = not continueRoutine
    while continueRoutine:
        # get current time
        t = routineTimer.getTime()
        tThisFlip = win.getFutureFlipTime(clock=routineTimer)
        tThisFlipGlobal = win.getFutureFlipTime(clock=None)
        frameN = frameN + 1  # number of completed frames (so 0 is the first frame)
        # update/draw components on each frame
        
        # *text_2* updates
        
        # if text_2 is starting this frame...
        if text_2.status == NOT_STARTED and tThisFlip >= 0.0-frameTolerance:
            # keep track of start time/frame for later
            text_2.frameNStart = frameN  # exact frame index
            text_2.tStart = t  # local t and not account for scr refresh
            text_2.tStartRefresh = tThisFlipGlobal  # on global time
            win.timeOnFlip(text_2, 'tStartRefresh')  # time at next scr refresh
            # add timestamp to datafile
            thisExp.timestampOnFlip(win, 'text_2.started')
            # update status
            text_2.status = STARTED
            text_2.setAutoDraw(True)
        
        # if text_2 is active this frame...
        if text_2.status == STARTED:
            # update params
            pass
        
        # *kbInstrUso* updates
        waitOnFlip = False
        
        # if kbInstrUso is starting this frame...
        if kbInstrUso.status == NOT_STARTED and tThisFlip >= 0.0-frameTolerance:
            # keep track of start time/frame for later
            kbInstrUso.frameNStart = frameN  # exact frame index
            kbInstrUso.tStart = t  # local t and not account for scr refresh
            kbInstrUso.tStartRefresh = tThisFlipGlobal  # on global time
            win.timeOnFlip(kbInstrUso, 'tStartRefresh')  # time at next scr refresh
            # add timestamp to datafile
            thisExp.timestampOnFlip(win, 'kbInstrUso.started')
            # update status
            kbInstrUso.status = STARTED
            # keyboard checking is just starting
            waitOnFlip = True
            win.callOnFlip(kbInstrUso.clock.reset)  # t=0 on next screen flip
            win.callOnFlip(kbInstrUso.clearEvents, eventType='keyboard')  # clear events on next screen flip
        if kbInstrUso.status == STARTED and not waitOnFlip:
            theseKeys = kbInstrUso.getKeys(keyList=['space'], ignoreKeys=["escape"], waitRelease=False)
            _kbInstrUso_allKeys.extend(theseKeys)
            if len(_kbInstrUso_allKeys):
                kbInstrUso.keys = _kbInstrUso_allKeys[-1].name  # just the last key pressed
                kbInstrUso.rt = _kbInstrUso_allKeys[-1].rt
                kbInstrUso.duration = _kbInstrUso_allKeys[-1].duration
                # a response ends the routine
                continueRoutine = False
        
        # *image_2* updates
        
        # if image_2 is starting this frame...
        if image_2.status == NOT_STARTED and tThisFlip >= 0.0-frameTolerance:
            # keep track of start time/frame for later
            image_2.frameNStart = frameN  # exact frame index
            image_2.tStart = t  # local t and not account for scr refresh
            image_2.tStartRefresh = tThisFlipGlobal  # on global time
            win.timeOnFlip(image_2, 'tStartRefresh')  # time at next scr refresh
            # add timestamp to datafile
            thisExp.timestampOnFlip(win, 'image_2.started')
            # update status
            image_2.status = STARTED
            image_2.setAutoDraw(True)
        
        # if image_2 is active this frame...
        if image_2.status == STARTED:
            # update params
            pass
        
        # check for quit (typically the Esc key)
        if defaultKeyboard.getKeys(keyList=["escape"]):
            thisExp.status = FINISHED
        if thisExp.status == FINISHED or endExpNow:
            endExperiment(thisExp, win=win)
            return
        # pause experiment here if requested
        if thisExp.status == PAUSED:
            pauseExperiment(
                thisExp=thisExp, 
                win=win, 
                timers=[routineTimer], 
                playbackComponents=[]
            )
            # skip the frame we paused on
            continue
        
        # check if all components have finished
        if not continueRoutine:  # a component has requested a forced-end of Routine
            Inst_Din1.forceEnded = routineForceEnded = True
            break
        continueRoutine = False  # will revert to True if at least one component still running
        for thisComponent in Inst_Din1.components:
            if hasattr(thisComponent, "status") and thisComponent.status != FINISHED:
                continueRoutine = True
                break  # at least one component has not yet finished
        
        # refresh the screen
        if continueRoutine:  # don't flip if this routine is over or we'll get a blank screen
            win.flip()
    
    # --- Ending Routine "Inst_Din1" ---
    for thisComponent in Inst_Din1.components:
        if hasattr(thisComponent, "setAutoDraw"):
            thisComponent.setAutoDraw(False)
    # store stop times for Inst_Din1
    Inst_Din1.tStop = globalClock.getTime(format='float')
    Inst_Din1.tStopRefresh = tThisFlipGlobal
    thisExp.addData('Inst_Din1.stopped', Inst_Din1.tStop)
    # check responses
    if kbInstrUso.keys in ['', [], None]:  # No response was made
        kbInstrUso.keys = None
    thisExp.addData('kbInstrUso.keys',kbInstrUso.keys)
    if kbInstrUso.keys != None:  # we had a response
        thisExp.addData('kbInstrUso.rt', kbInstrUso.rt)
        thisExp.addData('kbInstrUso.duration', kbInstrUso.duration)
    thisExp.nextEntry()
    # the Routine "Inst_Din1" was not non-slip safe, so reset the non-slip timer
    routineTimer.reset()
    
    # --- Prepare to start Routine "circulos" ---
    # create an object to store info about Routine circulos
    circulos = data.Routine(
        name='circulos',
        components=[image_3, key_resp_4, text_11],
    )
    circulos.status = NOT_STARTED
    continueRoutine = True
    # update component parameters for each repeat
    # create starting attributes for key_resp_4
    key_resp_4.keys = []
    key_resp_4.rt = []
    _key_resp_4_allKeys = []
    # store start times for circulos
    circulos.tStartRefresh = win.getFutureFlipTime(clock=globalClock)
    circulos.tStart = globalClock.getTime(format='float')
    circulos.status = STARTED
    thisExp.addData('circulos.started', circulos.tStart)
    circulos.maxDuration = None
    # keep track of which components have finished
    circulosComponents = circulos.components
    for thisComponent in circulos.components:
        thisComponent.tStart = None
        thisComponent.tStop = None
        thisComponent.tStartRefresh = None
        thisComponent.tStopRefresh = None
        if hasattr(thisComponent, 'status'):
            thisComponent.status = NOT_STARTED
    # reset timers
    t = 0
    _timeToFirstFrame = win.getFutureFlipTime(clock="now")
    frameN = -1
    
    # --- Run Routine "circulos" ---
    circulos.forceEnded = routineForceEnded = not continueRoutine
    while continueRoutine:
        # get current time
        t = routineTimer.getTime()
        tThisFlip = win.getFutureFlipTime(clock=routineTimer)
        tThisFlipGlobal = win.getFutureFlipTime(clock=None)
        frameN = frameN + 1  # number of completed frames (so 0 is the first frame)
        # update/draw components on each frame
        
        # *image_3* updates
        
        # if image_3 is starting this frame...
        if image_3.status == NOT_STARTED and tThisFlip >= 0.0-frameTolerance:
            # keep track of start time/frame for later
            image_3.frameNStart = frameN  # exact frame index
            image_3.tStart = t  # local t and not account for scr refresh
            image_3.tStartRefresh = tThisFlipGlobal  # on global time
            win.timeOnFlip(image_3, 'tStartRefresh')  # time at next scr refresh
            # add timestamp to datafile
            thisExp.timestampOnFlip(win, 'image_3.started')
            # update status
            image_3.status = STARTED
            image_3.setAutoDraw(True)
        
        # if image_3 is active this frame...
        if image_3.status == STARTED:
            # update params
            pass
        
        # *key_resp_4* updates
        waitOnFlip = False
        
        # if key_resp_4 is starting this frame...
        if key_resp_4.status == NOT_STARTED and tThisFlip >= 0.0-frameTolerance:
            # keep track of start time/frame for later
            key_resp_4.frameNStart = frameN  # exact frame index
            key_resp_4.tStart = t  # local t and not account for scr refresh
            key_resp_4.tStartRefresh = tThisFlipGlobal  # on global time
            win.timeOnFlip(key_resp_4, 'tStartRefresh')  # time at next scr refresh
            # add timestamp to datafile
            thisExp.timestampOnFlip(win, 'key_resp_4.started')
            # update status
            key_resp_4.status = STARTED
            # keyboard checking is just starting
            waitOnFlip = True
            win.callOnFlip(key_resp_4.clock.reset)  # t=0 on next screen flip
            win.callOnFlip(key_resp_4.clearEvents, eventType='keyboard')  # clear events on next screen flip
        if key_resp_4.status == STARTED and not waitOnFlip:
            theseKeys = key_resp_4.getKeys(keyList=['space'], ignoreKeys=["escape"], waitRelease=False)
            _key_resp_4_allKeys.extend(theseKeys)
            if len(_key_resp_4_allKeys):
                key_resp_4.keys = _key_resp_4_allKeys[-1].name  # just the last key pressed
                key_resp_4.rt = _key_resp_4_allKeys[-1].rt
                key_resp_4.duration = _key_resp_4_allKeys[-1].duration
                # a response ends the routine
                continueRoutine = False
        
        # *text_11* updates
        
        # if text_11 is starting this frame...
        if text_11.status == NOT_STARTED and tThisFlip >= 0.0-frameTolerance:
            # keep track of start time/frame for later
            text_11.frameNStart = frameN  # exact frame index
            text_11.tStart = t  # local t and not account for scr refresh
            text_11.tStartRefresh = tThisFlipGlobal  # on global time
            win.timeOnFlip(text_11, 'tStartRefresh')  # time at next scr refresh
            # add timestamp to datafile
            thisExp.timestampOnFlip(win, 'text_11.started')
            # update status
            text_11.status = STARTED
            text_11.setAutoDraw(True)
        
        # if text_11 is active this frame...
        if text_11.status == STARTED:
            # update params
            pass
        
        # check for quit (typically the Esc key)
        if defaultKeyboard.getKeys(keyList=["escape"]):
            thisExp.status = FINISHED
        if thisExp.status == FINISHED or endExpNow:
            endExperiment(thisExp, win=win)
            return
        # pause experiment here if requested
        if thisExp.status == PAUSED:
            pauseExperiment(
                thisExp=thisExp, 
                win=win, 
                timers=[routineTimer], 
                playbackComponents=[]
            )
            # skip the frame we paused on
            continue
        
        # check if all components have finished
        if not continueRoutine:  # a component has requested a forced-end of Routine
            circulos.forceEnded = routineForceEnded = True
            break
        continueRoutine = False  # will revert to True if at least one component still running
        for thisComponent in circulos.components:
            if hasattr(thisComponent, "status") and thisComponent.status != FINISHED:
                continueRoutine = True
                break  # at least one component has not yet finished
        
        # refresh the screen
        if continueRoutine:  # don't flip if this routine is over or we'll get a blank screen
            win.flip()
    
    # --- Ending Routine "circulos" ---
    for thisComponent in circulos.components:
        if hasattr(thisComponent, "setAutoDraw"):
            thisComponent.setAutoDraw(False)
    # store stop times for circulos
    circulos.tStop = globalClock.getTime(format='float')
    circulos.tStopRefresh = tThisFlipGlobal
    thisExp.addData('circulos.stopped', circulos.tStop)
    # check responses
    if key_resp_4.keys in ['', [], None]:  # No response was made
        key_resp_4.keys = None
    thisExp.addData('key_resp_4.keys',key_resp_4.keys)
    if key_resp_4.keys != None:  # we had a response
        thisExp.addData('key_resp_4.rt', key_resp_4.rt)
        thisExp.addData('key_resp_4.duration', key_resp_4.duration)
    thisExp.nextEntry()
    # the Routine "circulos" was not non-slip safe, so reset the non-slip timer
    routineTimer.reset()
    
    # --- Prepare to start Routine "br" ---
    # create an object to store info about Routine br
    br = data.Routine(
        name='br',
        components=[text_7, key_resp_3],
    )
    br.status = NOT_STARTED
    continueRoutine = True
    # update component parameters for each repeat
    # create starting attributes for key_resp_3
    key_resp_3.keys = []
    key_resp_3.rt = []
    _key_resp_3_allKeys = []
    # store start times for br
    br.tStartRefresh = win.getFutureFlipTime(clock=globalClock)
    br.tStart = globalClock.getTime(format='float')
    br.status = STARTED
    thisExp.addData('br.started', br.tStart)
    br.maxDuration = None
    # keep track of which components have finished
    brComponents = br.components
    for thisComponent in br.components:
        thisComponent.tStart = None
        thisComponent.tStop = None
        thisComponent.tStartRefresh = None
        thisComponent.tStopRefresh = None
        if hasattr(thisComponent, 'status'):
            thisComponent.status = NOT_STARTED
    # reset timers
    t = 0
    _timeToFirstFrame = win.getFutureFlipTime(clock="now")
    frameN = -1
    
    # --- Run Routine "br" ---
    br.forceEnded = routineForceEnded = not continueRoutine
    while continueRoutine:
        # get current time
        t = routineTimer.getTime()
        tThisFlip = win.getFutureFlipTime(clock=routineTimer)
        tThisFlipGlobal = win.getFutureFlipTime(clock=None)
        frameN = frameN + 1  # number of completed frames (so 0 is the first frame)
        # update/draw components on each frame
        
        # *text_7* updates
        
        # if text_7 is starting this frame...
        if text_7.status == NOT_STARTED and tThisFlip >= 0.0-frameTolerance:
            # keep track of start time/frame for later
            text_7.frameNStart = frameN  # exact frame index
            text_7.tStart = t  # local t and not account for scr refresh
            text_7.tStartRefresh = tThisFlipGlobal  # on global time
            win.timeOnFlip(text_7, 'tStartRefresh')  # time at next scr refresh
            # add timestamp to datafile
            thisExp.timestampOnFlip(win, 'text_7.started')
            # update status
            text_7.status = STARTED
            text_7.setAutoDraw(True)
        
        # if text_7 is active this frame...
        if text_7.status == STARTED:
            # update params
            pass
        
        # *key_resp_3* updates
        waitOnFlip = False
        
        # if key_resp_3 is starting this frame...
        if key_resp_3.status == NOT_STARTED and tThisFlip >= 0.0-frameTolerance:
            # keep track of start time/frame for later
            key_resp_3.frameNStart = frameN  # exact frame index
            key_resp_3.tStart = t  # local t and not account for scr refresh
            key_resp_3.tStartRefresh = tThisFlipGlobal  # on global time
            win.timeOnFlip(key_resp_3, 'tStartRefresh')  # time at next scr refresh
            # add timestamp to datafile
            thisExp.timestampOnFlip(win, 'key_resp_3.started')
            # update status
            key_resp_3.status = STARTED
            # keyboard checking is just starting
            waitOnFlip = True
            win.callOnFlip(key_resp_3.clock.reset)  # t=0 on next screen flip
            win.callOnFlip(key_resp_3.clearEvents, eventType='keyboard')  # clear events on next screen flip
        if key_resp_3.status == STARTED and not waitOnFlip:
            theseKeys = key_resp_3.getKeys(keyList=['space'], ignoreKeys=["escape"], waitRelease=False)
            _key_resp_3_allKeys.extend(theseKeys)
            if len(_key_resp_3_allKeys):
                key_resp_3.keys = _key_resp_3_allKeys[-1].name  # just the last key pressed
                key_resp_3.rt = _key_resp_3_allKeys[-1].rt
                key_resp_3.duration = _key_resp_3_allKeys[-1].duration
                # a response ends the routine
                continueRoutine = False
        
        # check for quit (typically the Esc key)
        if defaultKeyboard.getKeys(keyList=["escape"]):
            thisExp.status = FINISHED
        if thisExp.status == FINISHED or endExpNow:
            endExperiment(thisExp, win=win)
            return
        # pause experiment here if requested
        if thisExp.status == PAUSED:
            pauseExperiment(
                thisExp=thisExp, 
                win=win, 
                timers=[routineTimer], 
                playbackComponents=[]
            )
            # skip the frame we paused on
            continue
        
        # check if all components have finished
        if not continueRoutine:  # a component has requested a forced-end of Routine
            br.forceEnded = routineForceEnded = True
            break
        continueRoutine = False  # will revert to True if at least one component still running
        for thisComponent in br.components:
            if hasattr(thisComponent, "status") and thisComponent.status != FINISHED:
                continueRoutine = True
                break  # at least one component has not yet finished
        
        # refresh the screen
        if continueRoutine:  # don't flip if this routine is over or we'll get a blank screen
            win.flip()
    
    # --- Ending Routine "br" ---
    for thisComponent in br.components:
        if hasattr(thisComponent, "setAutoDraw"):
            thisComponent.setAutoDraw(False)
    # store stop times for br
    br.tStop = globalClock.getTime(format='float')
    br.tStopRefresh = tThisFlipGlobal
    thisExp.addData('br.stopped', br.tStop)
    # check responses
    if key_resp_3.keys in ['', [], None]:  # No response was made
        key_resp_3.keys = None
    thisExp.addData('key_resp_3.keys',key_resp_3.keys)
    if key_resp_3.keys != None:  # we had a response
        thisExp.addData('key_resp_3.rt', key_resp_3.rt)
        thisExp.addData('key_resp_3.duration', key_resp_3.duration)
    thisExp.nextEntry()
    # the Routine "br" was not non-slip safe, so reset the non-slip timer
    routineTimer.reset()
    
    # --- Prepare to start Routine "txtInstrCal" ---
    # create an object to store info about Routine txtInstrCal
    txtInstrCal = data.Routine(
        name='txtInstrCal',
        components=[text, kbInstrCal],
    )
    txtInstrCal.status = NOT_STARTED
    continueRoutine = True
    # update component parameters for each repeat
    # create starting attributes for kbInstrCal
    kbInstrCal.keys = []
    kbInstrCal.rt = []
    _kbInstrCal_allKeys = []
    # store start times for txtInstrCal
    txtInstrCal.tStartRefresh = win.getFutureFlipTime(clock=globalClock)
    txtInstrCal.tStart = globalClock.getTime(format='float')
    txtInstrCal.status = STARTED
    thisExp.addData('txtInstrCal.started', txtInstrCal.tStart)
    txtInstrCal.maxDuration = None
    # keep track of which components have finished
    txtInstrCalComponents = txtInstrCal.components
    for thisComponent in txtInstrCal.components:
        thisComponent.tStart = None
        thisComponent.tStop = None
        thisComponent.tStartRefresh = None
        thisComponent.tStopRefresh = None
        if hasattr(thisComponent, 'status'):
            thisComponent.status = NOT_STARTED
    # reset timers
    t = 0
    _timeToFirstFrame = win.getFutureFlipTime(clock="now")
    frameN = -1
    
    # --- Run Routine "txtInstrCal" ---
    txtInstrCal.forceEnded = routineForceEnded = not continueRoutine
    while continueRoutine:
        # get current time
        t = routineTimer.getTime()
        tThisFlip = win.getFutureFlipTime(clock=routineTimer)
        tThisFlipGlobal = win.getFutureFlipTime(clock=None)
        frameN = frameN + 1  # number of completed frames (so 0 is the first frame)
        # update/draw components on each frame
        
        # *text* updates
        
        # if text is starting this frame...
        if text.status == NOT_STARTED and tThisFlip >= 0.0-frameTolerance:
            # keep track of start time/frame for later
            text.frameNStart = frameN  # exact frame index
            text.tStart = t  # local t and not account for scr refresh
            text.tStartRefresh = tThisFlipGlobal  # on global time
            win.timeOnFlip(text, 'tStartRefresh')  # time at next scr refresh
            # add timestamp to datafile
            thisExp.timestampOnFlip(win, 'text.started')
            # update status
            text.status = STARTED
            text.setAutoDraw(True)
        
        # if text is active this frame...
        if text.status == STARTED:
            # update params
            pass
        
        # *kbInstrCal* updates
        waitOnFlip = False
        
        # if kbInstrCal is starting this frame...
        if kbInstrCal.status == NOT_STARTED and tThisFlip >= 0.0-frameTolerance:
            # keep track of start time/frame for later
            kbInstrCal.frameNStart = frameN  # exact frame index
            kbInstrCal.tStart = t  # local t and not account for scr refresh
            kbInstrCal.tStartRefresh = tThisFlipGlobal  # on global time
            win.timeOnFlip(kbInstrCal, 'tStartRefresh')  # time at next scr refresh
            # add timestamp to datafile
            thisExp.timestampOnFlip(win, 'kbInstrCal.started')
            # update status
            kbInstrCal.status = STARTED
            # keyboard checking is just starting
            waitOnFlip = True
            win.callOnFlip(kbInstrCal.clock.reset)  # t=0 on next screen flip
            win.callOnFlip(kbInstrCal.clearEvents, eventType='keyboard')  # clear events on next screen flip
        if kbInstrCal.status == STARTED and not waitOnFlip:
            theseKeys = kbInstrCal.getKeys(keyList=['space'], ignoreKeys=["escape"], waitRelease=False)
            _kbInstrCal_allKeys.extend(theseKeys)
            if len(_kbInstrCal_allKeys):
                kbInstrCal.keys = _kbInstrCal_allKeys[-1].name  # just the last key pressed
                kbInstrCal.rt = _kbInstrCal_allKeys[-1].rt
                kbInstrCal.duration = _kbInstrCal_allKeys[-1].duration
                # a response ends the routine
                continueRoutine = False
        
        # check for quit (typically the Esc key)
        if defaultKeyboard.getKeys(keyList=["escape"]):
            thisExp.status = FINISHED
        if thisExp.status == FINISHED or endExpNow:
            endExperiment(thisExp, win=win)
            return
        # pause experiment here if requested
        if thisExp.status == PAUSED:
            pauseExperiment(
                thisExp=thisExp, 
                win=win, 
                timers=[routineTimer], 
                playbackComponents=[]
            )
            # skip the frame we paused on
            continue
        
        # check if all components have finished
        if not continueRoutine:  # a component has requested a forced-end of Routine
            txtInstrCal.forceEnded = routineForceEnded = True
            break
        continueRoutine = False  # will revert to True if at least one component still running
        for thisComponent in txtInstrCal.components:
            if hasattr(thisComponent, "status") and thisComponent.status != FINISHED:
                continueRoutine = True
                break  # at least one component has not yet finished
        
        # refresh the screen
        if continueRoutine:  # don't flip if this routine is over or we'll get a blank screen
            win.flip()
    
    # --- Ending Routine "txtInstrCal" ---
    for thisComponent in txtInstrCal.components:
        if hasattr(thisComponent, "setAutoDraw"):
            thisComponent.setAutoDraw(False)
    # store stop times for txtInstrCal
    txtInstrCal.tStop = globalClock.getTime(format='float')
    txtInstrCal.tStopRefresh = tThisFlipGlobal
    thisExp.addData('txtInstrCal.stopped', txtInstrCal.tStop)
    # check responses
    if kbInstrCal.keys in ['', [], None]:  # No response was made
        kbInstrCal.keys = None
    thisExp.addData('kbInstrCal.keys',kbInstrCal.keys)
    if kbInstrCal.keys != None:  # we had a response
        thisExp.addData('kbInstrCal.rt', kbInstrCal.rt)
        thisExp.addData('kbInstrCal.duration', kbInstrCal.duration)
    thisExp.nextEntry()
    # the Routine "txtInstrCal" was not non-slip safe, so reset the non-slip timer
    routineTimer.reset()
    
    # --- Prepare to start Routine "RegCount" ---
    # create an object to store info about Routine RegCount
    RegCount = data.Routine(
        name='RegCount',
        components=[text_5, text_6],
    )
    RegCount.status = NOT_STARTED
    continueRoutine = True
    # update component parameters for each repeat
    # store start times for RegCount
    RegCount.tStartRefresh = win.getFutureFlipTime(clock=globalClock)
    RegCount.tStart = globalClock.getTime(format='float')
    RegCount.status = STARTED
    thisExp.addData('RegCount.started', RegCount.tStart)
    RegCount.maxDuration = None
    win.color = [0.6078, -0.2784, -0.2784]
    win.colorSpace = 'rgb'
    win.backgroundImage = ''
    win.backgroundFit = 'none'
    # keep track of which components have finished
    RegCountComponents = RegCount.components
    for thisComponent in RegCount.components:
        thisComponent.tStart = None
        thisComponent.tStop = None
        thisComponent.tStartRefresh = None
        thisComponent.tStopRefresh = None
        if hasattr(thisComponent, 'status'):
            thisComponent.status = NOT_STARTED
    # reset timers
    t = 0
    _timeToFirstFrame = win.getFutureFlipTime(clock="now")
    frameN = -1
    
    # --- Run Routine "RegCount" ---
    RegCount.forceEnded = routineForceEnded = not continueRoutine
    while continueRoutine and routineTimer.getTime() < 5.0:
        # get current time
        t = routineTimer.getTime()
        tThisFlip = win.getFutureFlipTime(clock=routineTimer)
        tThisFlipGlobal = win.getFutureFlipTime(clock=None)
        frameN = frameN + 1  # number of completed frames (so 0 is the first frame)
        # update/draw components on each frame
        
        # *text_5* updates
        
        # if text_5 is starting this frame...
        if text_5.status == NOT_STARTED and tThisFlip >= 0.0-frameTolerance:
            # keep track of start time/frame for later
            text_5.frameNStart = frameN  # exact frame index
            text_5.tStart = t  # local t and not account for scr refresh
            text_5.tStartRefresh = tThisFlipGlobal  # on global time
            win.timeOnFlip(text_5, 'tStartRefresh')  # time at next scr refresh
            # add timestamp to datafile
            thisExp.timestampOnFlip(win, 'text_5.started')
            # update status
            text_5.status = STARTED
            text_5.setAutoDraw(True)
        
        # if text_5 is active this frame...
        if text_5.status == STARTED:
            # update params
            pass
        
        # if text_5 is stopping this frame...
        if text_5.status == STARTED:
            # is it time to stop? (based on global clock, using actual start)
            if tThisFlipGlobal > text_5.tStartRefresh + 5-frameTolerance:
                # keep track of stop time/frame for later
                text_5.tStop = t  # not accounting for scr refresh
                text_5.tStopRefresh = tThisFlipGlobal  # on global time
                text_5.frameNStop = frameN  # exact frame index
                # add timestamp to datafile
                thisExp.timestampOnFlip(win, 'text_5.stopped')
                # update status
                text_5.status = FINISHED
                text_5.setAutoDraw(False)
        
        # *text_6* updates
        
        # if text_6 is starting this frame...
        if text_6.status == NOT_STARTED and tThisFlip >= 0.0-frameTolerance:
            # keep track of start time/frame for later
            text_6.frameNStart = frameN  # exact frame index
            text_6.tStart = t  # local t and not account for scr refresh
            text_6.tStartRefresh = tThisFlipGlobal  # on global time
            win.timeOnFlip(text_6, 'tStartRefresh')  # time at next scr refresh
            # add timestamp to datafile
            thisExp.timestampOnFlip(win, 'text_6.started')
            # update status
            text_6.status = STARTED
            text_6.setAutoDraw(True)
        
        # if text_6 is active this frame...
        if text_6.status == STARTED:
            # update params
            text_6.setText(int(6 - t), log=False)
        
        # if text_6 is stopping this frame...
        if text_6.status == STARTED:
            # is it time to stop? (based on global clock, using actual start)
            if tThisFlipGlobal > text_6.tStartRefresh + 5-frameTolerance:
                # keep track of stop time/frame for later
                text_6.tStop = t  # not accounting for scr refresh
                text_6.tStopRefresh = tThisFlipGlobal  # on global time
                text_6.frameNStop = frameN  # exact frame index
                # add timestamp to datafile
                thisExp.timestampOnFlip(win, 'text_6.stopped')
                # update status
                text_6.status = FINISHED
                text_6.setAutoDraw(False)
        
        # check for quit (typically the Esc key)
        if defaultKeyboard.getKeys(keyList=["escape"]):
            thisExp.status = FINISHED
        if thisExp.status == FINISHED or endExpNow:
            endExperiment(thisExp, win=win)
            return
        # pause experiment here if requested
        if thisExp.status == PAUSED:
            pauseExperiment(
                thisExp=thisExp, 
                win=win, 
                timers=[routineTimer], 
                playbackComponents=[]
            )
            # skip the frame we paused on
            continue
        
        # check if all components have finished
        if not continueRoutine:  # a component has requested a forced-end of Routine
            RegCount.forceEnded = routineForceEnded = True
            break
        continueRoutine = False  # will revert to True if at least one component still running
        for thisComponent in RegCount.components:
            if hasattr(thisComponent, "status") and thisComponent.status != FINISHED:
                continueRoutine = True
                break  # at least one component has not yet finished
        
        # refresh the screen
        if continueRoutine:  # don't flip if this routine is over or we'll get a blank screen
            win.flip()
    
    # --- Ending Routine "RegCount" ---
    for thisComponent in RegCount.components:
        if hasattr(thisComponent, "setAutoDraw"):
            thisComponent.setAutoDraw(False)
    # store stop times for RegCount
    RegCount.tStop = globalClock.getTime(format='float')
    RegCount.tStopRefresh = tThisFlipGlobal
    thisExp.addData('RegCount.stopped', RegCount.tStop)
    setupWindow(expInfo=expInfo, win=win)
    # using non-slip timing so subtract the expected duration of this Routine (unless ended on request)
    if RegCount.maxDurationReached:
        routineTimer.addTime(-RegCount.maxDuration)
    elif RegCount.forceEnded:
        routineTimer.reset()
    else:
        routineTimer.addTime(-5.000000)
    thisExp.nextEntry()
    
    # --- Prepare to start Routine "Calibracion" ---
    # create an object to store info about Routine Calibracion
    Calibracion = data.Routine(
        name='Calibracion',
        components=[txtCalibHeader, txtCalibPrompt, txtCountdown, txtAttemptMsg, dotRecord],
    )
    Calibracion.status = NOT_STARTED
    continueRoutine = True
    # update component parameters for each repeat
    # Run 'Begin Routine' code from codeCal
    # Parámetros
    ATTEMPTS_TOTAL = 2
    ATTEMPT_DUR = 5.0  # s por intento
    REST_DUR = 5.0     # s por descanso
    
    # Estado
    attempt_idx = 1         # 1..2
    phase = 'attempt'       # 'attempt' o 'rest'
    phaseClock = core.Clock()
    peak_force = 0.0
    peaks = []              # picos por intento
    
    # UI inicial
    txtCalibHeader.text  = f""
    txtCalibPrompt.text  = "Aprieta con mucha fuerza"
    txtCountdown.text    = f"{ATTEMPT_DUR:0.1f} s"
    txtAttemptMsg.text   = ""
    dotRecord.opacity    = 1.0  # visible durante intento
    
    IDLE_THRESHOLD_N = 8.0  # umbral para “hay presión real”
    pressing = False        # estado actual (evita parpadeo)
    
    # store start times for Calibracion
    Calibracion.tStartRefresh = win.getFutureFlipTime(clock=globalClock)
    Calibracion.tStart = globalClock.getTime(format='float')
    Calibracion.status = STARTED
    thisExp.addData('Calibracion.started', Calibracion.tStart)
    Calibracion.maxDuration = None
    # keep track of which components have finished
    CalibracionComponents = Calibracion.components
    for thisComponent in Calibracion.components:
        thisComponent.tStart = None
        thisComponent.tStop = None
        thisComponent.tStartRefresh = None
        thisComponent.tStopRefresh = None
        if hasattr(thisComponent, 'status'):
            thisComponent.status = NOT_STARTED
    # reset timers
    t = 0
    _timeToFirstFrame = win.getFutureFlipTime(clock="now")
    frameN = -1
    
    # --- Run Routine "Calibracion" ---
    Calibracion.forceEnded = routineForceEnded = not continueRoutine
    while continueRoutine:
        # get current time
        t = routineTimer.getTime()
        tThisFlip = win.getFutureFlipTime(clock=routineTimer)
        tThisFlipGlobal = win.getFutureFlipTime(clock=None)
        frameN = frameN + 1  # number of completed frames (so 0 is the first frame)
        # update/draw components on each frame
        # Run 'Each Frame' code from codeCal
        # Tiempo de fase (intento o descanso)
        t = phaseClock.getTime()
        
        # --- Lectura de fuerza actual (NO bloqueante; usar sampler) ---
        try:
            f_now = float(_gdx_state.last)
        except Exception:
            s = gdx.read()
            f_now = float(s[0]) if (s and s[0] is not None) else 0.0
        
        # --- Umbral de reposo para mostrar el círculo solo si aprietas de verdad ---
        IDLE_THRESHOLD_N = 8.0  # ajusta a 5–12 N según tu sensor/mesa
        pressing = (f_now > IDLE_THRESHOLD_N)
        
        # Círculo visible solo si hay presión real y estamos en intento
        dotRecord.opacity = 1.0 if (phase == 'attempt' and pressing) else 0.0
        
        if phase == 'attempt':
            win.color = (-0.1686, 0.0667, 0.2000) # Fondo color durante el intento
            # Actualiza pico del intento
            if f_now > peak_force:
                peak_force = f_now
        
            # UI
            remaining = max(0.0, ATTEMPT_DUR - t)
            txtCalibHeader.text = f""
            txtCalibPrompt.text = "Aprieta con mucha fuerza"
            txtCountdown.text   = f"{remaining:0.1f} s"
            txtAttemptMsg.text  = ""
        
            # Fin del intento
            if t >= ATTEMPT_DUR:
                peaks.append(peak_force)
                phase = 'rest'
                phaseClock.reset()
        
        elif phase == 'rest':
            win.color = 'darkgreen'  # Fondo color verde oscuro durante descanso
            remaining = max(0.0, REST_DUR - t)
        
            # Mensajes de descanso
            txtCalibHeader.text = f"Descanso"
            txtCalibPrompt.text = "RELAJA LA MANO Y ESPERA."
            txtCountdown.text   = f"{remaining:0.1f} s"
            txtAttemptMsg.text  = "." if t < 1.0 else ""
        
            # Fin del descanso → siguiente intento o terminar rutina
            if t >= REST_DUR:
                attempt_idx += 1
                if attempt_idx <= ATTEMPTS_TOTAL:
                    # Nuevo intento
                    peak_force = 0.0
                    phase = 'attempt'
                    phaseClock.reset()
                else:
                    # Terminar rutina
                    continueRoutine = False
        
        
        # *txtCalibHeader* updates
        
        # if txtCalibHeader is starting this frame...
        if txtCalibHeader.status == NOT_STARTED and tThisFlip >= 0.0-frameTolerance:
            # keep track of start time/frame for later
            txtCalibHeader.frameNStart = frameN  # exact frame index
            txtCalibHeader.tStart = t  # local t and not account for scr refresh
            txtCalibHeader.tStartRefresh = tThisFlipGlobal  # on global time
            win.timeOnFlip(txtCalibHeader, 'tStartRefresh')  # time at next scr refresh
            # add timestamp to datafile
            thisExp.timestampOnFlip(win, 'txtCalibHeader.started')
            # update status
            txtCalibHeader.status = STARTED
            txtCalibHeader.setAutoDraw(True)
        
        # if txtCalibHeader is active this frame...
        if txtCalibHeader.status == STARTED:
            # update params
            pass
        
        # *txtCalibPrompt* updates
        
        # if txtCalibPrompt is starting this frame...
        if txtCalibPrompt.status == NOT_STARTED and tThisFlip >= 0.0-frameTolerance:
            # keep track of start time/frame for later
            txtCalibPrompt.frameNStart = frameN  # exact frame index
            txtCalibPrompt.tStart = t  # local t and not account for scr refresh
            txtCalibPrompt.tStartRefresh = tThisFlipGlobal  # on global time
            win.timeOnFlip(txtCalibPrompt, 'tStartRefresh')  # time at next scr refresh
            # add timestamp to datafile
            thisExp.timestampOnFlip(win, 'txtCalibPrompt.started')
            # update status
            txtCalibPrompt.status = STARTED
            txtCalibPrompt.setAutoDraw(True)
        
        # if txtCalibPrompt is active this frame...
        if txtCalibPrompt.status == STARTED:
            # update params
            pass
        
        # *txtCountdown* updates
        
        # if txtCountdown is starting this frame...
        if txtCountdown.status == NOT_STARTED and tThisFlip >= 0.0-frameTolerance:
            # keep track of start time/frame for later
            txtCountdown.frameNStart = frameN  # exact frame index
            txtCountdown.tStart = t  # local t and not account for scr refresh
            txtCountdown.tStartRefresh = tThisFlipGlobal  # on global time
            win.timeOnFlip(txtCountdown, 'tStartRefresh')  # time at next scr refresh
            # add timestamp to datafile
            thisExp.timestampOnFlip(win, 'txtCountdown.started')
            # update status
            txtCountdown.status = STARTED
            txtCountdown.setAutoDraw(True)
        
        # if txtCountdown is active this frame...
        if txtCountdown.status == STARTED:
            # update params
            pass
        
        # *txtAttemptMsg* updates
        
        # if txtAttemptMsg is starting this frame...
        if txtAttemptMsg.status == NOT_STARTED and tThisFlip >= 0.0-frameTolerance:
            # keep track of start time/frame for later
            txtAttemptMsg.frameNStart = frameN  # exact frame index
            txtAttemptMsg.tStart = t  # local t and not account for scr refresh
            txtAttemptMsg.tStartRefresh = tThisFlipGlobal  # on global time
            win.timeOnFlip(txtAttemptMsg, 'tStartRefresh')  # time at next scr refresh
            # add timestamp to datafile
            thisExp.timestampOnFlip(win, 'txtAttemptMsg.started')
            # update status
            txtAttemptMsg.status = STARTED
            txtAttemptMsg.setAutoDraw(True)
        
        # if txtAttemptMsg is active this frame...
        if txtAttemptMsg.status == STARTED:
            # update params
            pass
        
        # *dotRecord* updates
        
        # if dotRecord is starting this frame...
        if dotRecord.status == NOT_STARTED and tThisFlip >= 0.0-frameTolerance:
            # keep track of start time/frame for later
            dotRecord.frameNStart = frameN  # exact frame index
            dotRecord.tStart = t  # local t and not account for scr refresh
            dotRecord.tStartRefresh = tThisFlipGlobal  # on global time
            win.timeOnFlip(dotRecord, 'tStartRefresh')  # time at next scr refresh
            # add timestamp to datafile
            thisExp.timestampOnFlip(win, 'dotRecord.started')
            # update status
            dotRecord.status = STARTED
            dotRecord.setAutoDraw(True)
        
        # if dotRecord is active this frame...
        if dotRecord.status == STARTED:
            # update params
            pass
        
        # check for quit (typically the Esc key)
        if defaultKeyboard.getKeys(keyList=["escape"]):
            thisExp.status = FINISHED
        if thisExp.status == FINISHED or endExpNow:
            endExperiment(thisExp, win=win)
            return
        # pause experiment here if requested
        if thisExp.status == PAUSED:
            pauseExperiment(
                thisExp=thisExp, 
                win=win, 
                timers=[routineTimer], 
                playbackComponents=[]
            )
            # skip the frame we paused on
            continue
        
        # check if all components have finished
        if not continueRoutine:  # a component has requested a forced-end of Routine
            Calibracion.forceEnded = routineForceEnded = True
            break
        continueRoutine = False  # will revert to True if at least one component still running
        for thisComponent in Calibracion.components:
            if hasattr(thisComponent, "status") and thisComponent.status != FINISHED:
                continueRoutine = True
                break  # at least one component has not yet finished
        
        # refresh the screen
        if continueRoutine:  # don't flip if this routine is over or we'll get a blank screen
            win.flip()
    
    # --- Ending Routine "Calibracion" ---
    for thisComponent in Calibracion.components:
        if hasattr(thisComponent, "setAutoDraw"):
            thisComponent.setAutoDraw(False)
    # store stop times for Calibracion
    Calibracion.tStop = globalClock.getTime(format='float')
    Calibracion.tStopRefresh = tThisFlipGlobal
    thisExp.addData('Calibracion.stopped', Calibracion.tStop)
    # Run 'End Routine' code from codeCal
    # Cálculo de promedio y umbrales 70/30
    calib_success = 1 if len(peaks) == 2 else 0
    avg_f = sum(peaks)/len(peaks) if peaks else 0.0
    
    thr70 = 0.70 * avg_f   # umbral ALTO para reforzar (en N)
    thr30 = 0.30 * avg_f   # umbral BAJO para rearmar (en N)
    
    # Alias por compatibilidad
    thr60 = thr70
    thr40 = thr30
    
    # Guardar en datos
    thisExp.addData('calib_n_attempts', len(peaks))
    thisExp.addData('calib_success', calib_success)
    for i, p in enumerate(peaks, start=1):
        thisExp.addData(f'calib_peak_{i}', p)
    thisExp.addData('calib_avg_f', avg_f)
    thisExp.addData('calib_thr70', thr70)
    thisExp.addData('calib_thr30', thr30)
    thisExp.addData('calib_thr_hi_percent', 0.70)
    thisExp.addData('calib_thr_lo_percent', 0.30)
    
    # Guardar en extraInfo para Trial_Operant
    thisExp.extraInfo['thr70'] = float(thr70)
    thisExp.extraInfo['thr30'] = float(thr30)
    
    # Hacerlos disponibles como globales
    globals()['thr30'] = float(thr30)
    globals()['thr70'] = float(thr70)
    
    thisExp.nextEntry()
    # the Routine "Calibracion" was not non-slip safe, so reset the non-slip timer
    routineTimer.reset()
    
    # --- Prepare to start Routine "Instructions_FaseReforzamiento" ---
    # create an object to store info about Routine Instructions_FaseReforzamiento
    Instructions_FaseReforzamiento = data.Routine(
        name='Instructions_FaseReforzamiento',
        components=[txtInstrReinf, kbInstrReinf],
    )
    Instructions_FaseReforzamiento.status = NOT_STARTED
    continueRoutine = True
    # update component parameters for each repeat
    # Run 'Begin Routine' code from code_2
    win.color = (-0.1686, 0.0667, 0.2000)
    # create starting attributes for kbInstrReinf
    kbInstrReinf.keys = []
    kbInstrReinf.rt = []
    _kbInstrReinf_allKeys = []
    # store start times for Instructions_FaseReforzamiento
    Instructions_FaseReforzamiento.tStartRefresh = win.getFutureFlipTime(clock=globalClock)
    Instructions_FaseReforzamiento.tStart = globalClock.getTime(format='float')
    Instructions_FaseReforzamiento.status = STARTED
    thisExp.addData('Instructions_FaseReforzamiento.started', Instructions_FaseReforzamiento.tStart)
    Instructions_FaseReforzamiento.maxDuration = None
    # keep track of which components have finished
    Instructions_FaseReforzamientoComponents = Instructions_FaseReforzamiento.components
    for thisComponent in Instructions_FaseReforzamiento.components:
        thisComponent.tStart = None
        thisComponent.tStop = None
        thisComponent.tStartRefresh = None
        thisComponent.tStopRefresh = None
        if hasattr(thisComponent, 'status'):
            thisComponent.status = NOT_STARTED
    # reset timers
    t = 0
    _timeToFirstFrame = win.getFutureFlipTime(clock="now")
    frameN = -1
    
    # --- Run Routine "Instructions_FaseReforzamiento" ---
    Instructions_FaseReforzamiento.forceEnded = routineForceEnded = not continueRoutine
    while continueRoutine:
        # get current time
        t = routineTimer.getTime()
        tThisFlip = win.getFutureFlipTime(clock=routineTimer)
        tThisFlipGlobal = win.getFutureFlipTime(clock=None)
        frameN = frameN + 1  # number of completed frames (so 0 is the first frame)
        # update/draw components on each frame
        
        # *txtInstrReinf* updates
        
        # if txtInstrReinf is starting this frame...
        if txtInstrReinf.status == NOT_STARTED and tThisFlip >= 0.0-frameTolerance:
            # keep track of start time/frame for later
            txtInstrReinf.frameNStart = frameN  # exact frame index
            txtInstrReinf.tStart = t  # local t and not account for scr refresh
            txtInstrReinf.tStartRefresh = tThisFlipGlobal  # on global time
            win.timeOnFlip(txtInstrReinf, 'tStartRefresh')  # time at next scr refresh
            # add timestamp to datafile
            thisExp.timestampOnFlip(win, 'txtInstrReinf.started')
            # update status
            txtInstrReinf.status = STARTED
            txtInstrReinf.setAutoDraw(True)
        
        # if txtInstrReinf is active this frame...
        if txtInstrReinf.status == STARTED:
            # update params
            pass
        
        # *kbInstrReinf* updates
        waitOnFlip = False
        
        # if kbInstrReinf is starting this frame...
        if kbInstrReinf.status == NOT_STARTED and tThisFlip >= 0.0-frameTolerance:
            # keep track of start time/frame for later
            kbInstrReinf.frameNStart = frameN  # exact frame index
            kbInstrReinf.tStart = t  # local t and not account for scr refresh
            kbInstrReinf.tStartRefresh = tThisFlipGlobal  # on global time
            win.timeOnFlip(kbInstrReinf, 'tStartRefresh')  # time at next scr refresh
            # add timestamp to datafile
            thisExp.timestampOnFlip(win, 'kbInstrReinf.started')
            # update status
            kbInstrReinf.status = STARTED
            # keyboard checking is just starting
            waitOnFlip = True
            win.callOnFlip(kbInstrReinf.clock.reset)  # t=0 on next screen flip
            win.callOnFlip(kbInstrReinf.clearEvents, eventType='keyboard')  # clear events on next screen flip
        if kbInstrReinf.status == STARTED and not waitOnFlip:
            theseKeys = kbInstrReinf.getKeys(keyList=['space'], ignoreKeys=["escape"], waitRelease=False)
            _kbInstrReinf_allKeys.extend(theseKeys)
            if len(_kbInstrReinf_allKeys):
                kbInstrReinf.keys = _kbInstrReinf_allKeys[-1].name  # just the last key pressed
                kbInstrReinf.rt = _kbInstrReinf_allKeys[-1].rt
                kbInstrReinf.duration = _kbInstrReinf_allKeys[-1].duration
                # a response ends the routine
                continueRoutine = False
        
        # check for quit (typically the Esc key)
        if defaultKeyboard.getKeys(keyList=["escape"]):
            thisExp.status = FINISHED
        if thisExp.status == FINISHED or endExpNow:
            endExperiment(thisExp, win=win)
            return
        # pause experiment here if requested
        if thisExp.status == PAUSED:
            pauseExperiment(
                thisExp=thisExp, 
                win=win, 
                timers=[routineTimer], 
                playbackComponents=[]
            )
            # skip the frame we paused on
            continue
        
        # check if all components have finished
        if not continueRoutine:  # a component has requested a forced-end of Routine
            Instructions_FaseReforzamiento.forceEnded = routineForceEnded = True
            break
        continueRoutine = False  # will revert to True if at least one component still running
        for thisComponent in Instructions_FaseReforzamiento.components:
            if hasattr(thisComponent, "status") and thisComponent.status != FINISHED:
                continueRoutine = True
                break  # at least one component has not yet finished
        
        # refresh the screen
        if continueRoutine:  # don't flip if this routine is over or we'll get a blank screen
            win.flip()
    
    # --- Ending Routine "Instructions_FaseReforzamiento" ---
    for thisComponent in Instructions_FaseReforzamiento.components:
        if hasattr(thisComponent, "setAutoDraw"):
            thisComponent.setAutoDraw(False)
    # store stop times for Instructions_FaseReforzamiento
    Instructions_FaseReforzamiento.tStop = globalClock.getTime(format='float')
    Instructions_FaseReforzamiento.tStopRefresh = tThisFlipGlobal
    thisExp.addData('Instructions_FaseReforzamiento.stopped', Instructions_FaseReforzamiento.tStop)
    # check responses
    if kbInstrReinf.keys in ['', [], None]:  # No response was made
        kbInstrReinf.keys = None
    thisExp.addData('kbInstrReinf.keys',kbInstrReinf.keys)
    if kbInstrReinf.keys != None:  # we had a response
        thisExp.addData('kbInstrReinf.rt', kbInstrReinf.rt)
        thisExp.addData('kbInstrReinf.duration', kbInstrReinf.duration)
    thisExp.nextEntry()
    # the Routine "Instructions_FaseReforzamiento" was not non-slip safe, so reset the non-slip timer
    routineTimer.reset()
    
    # --- Prepare to start Routine "OJO" ---
    # create an object to store info about Routine OJO
    OJO = data.Routine(
        name='OJO',
        components=[text_3, key_resp, image],
    )
    OJO.status = NOT_STARTED
    continueRoutine = True
    # update component parameters for each repeat
    # create starting attributes for key_resp
    key_resp.keys = []
    key_resp.rt = []
    _key_resp_allKeys = []
    # store start times for OJO
    OJO.tStartRefresh = win.getFutureFlipTime(clock=globalClock)
    OJO.tStart = globalClock.getTime(format='float')
    OJO.status = STARTED
    thisExp.addData('OJO.started', OJO.tStart)
    OJO.maxDuration = None
    # keep track of which components have finished
    OJOComponents = OJO.components
    for thisComponent in OJO.components:
        thisComponent.tStart = None
        thisComponent.tStop = None
        thisComponent.tStartRefresh = None
        thisComponent.tStopRefresh = None
        if hasattr(thisComponent, 'status'):
            thisComponent.status = NOT_STARTED
    # reset timers
    t = 0
    _timeToFirstFrame = win.getFutureFlipTime(clock="now")
    frameN = -1
    
    # --- Run Routine "OJO" ---
    OJO.forceEnded = routineForceEnded = not continueRoutine
    while continueRoutine:
        # get current time
        t = routineTimer.getTime()
        tThisFlip = win.getFutureFlipTime(clock=routineTimer)
        tThisFlipGlobal = win.getFutureFlipTime(clock=None)
        frameN = frameN + 1  # number of completed frames (so 0 is the first frame)
        # update/draw components on each frame
        
        # *text_3* updates
        
        # if text_3 is starting this frame...
        if text_3.status == NOT_STARTED and tThisFlip >= 0.0-frameTolerance:
            # keep track of start time/frame for later
            text_3.frameNStart = frameN  # exact frame index
            text_3.tStart = t  # local t and not account for scr refresh
            text_3.tStartRefresh = tThisFlipGlobal  # on global time
            win.timeOnFlip(text_3, 'tStartRefresh')  # time at next scr refresh
            # add timestamp to datafile
            thisExp.timestampOnFlip(win, 'text_3.started')
            # update status
            text_3.status = STARTED
            text_3.setAutoDraw(True)
        
        # if text_3 is active this frame...
        if text_3.status == STARTED:
            # update params
            pass
        
        # *key_resp* updates
        waitOnFlip = False
        
        # if key_resp is starting this frame...
        if key_resp.status == NOT_STARTED and tThisFlip >= 0.0-frameTolerance:
            # keep track of start time/frame for later
            key_resp.frameNStart = frameN  # exact frame index
            key_resp.tStart = t  # local t and not account for scr refresh
            key_resp.tStartRefresh = tThisFlipGlobal  # on global time
            win.timeOnFlip(key_resp, 'tStartRefresh')  # time at next scr refresh
            # add timestamp to datafile
            thisExp.timestampOnFlip(win, 'key_resp.started')
            # update status
            key_resp.status = STARTED
            # keyboard checking is just starting
            waitOnFlip = True
            win.callOnFlip(key_resp.clock.reset)  # t=0 on next screen flip
            win.callOnFlip(key_resp.clearEvents, eventType='keyboard')  # clear events on next screen flip
        if key_resp.status == STARTED and not waitOnFlip:
            theseKeys = key_resp.getKeys(keyList=['space'], ignoreKeys=["escape"], waitRelease=False)
            _key_resp_allKeys.extend(theseKeys)
            if len(_key_resp_allKeys):
                key_resp.keys = _key_resp_allKeys[-1].name  # just the last key pressed
                key_resp.rt = _key_resp_allKeys[-1].rt
                key_resp.duration = _key_resp_allKeys[-1].duration
                # a response ends the routine
                continueRoutine = False
        
        # *image* updates
        
        # if image is starting this frame...
        if image.status == NOT_STARTED and tThisFlip >= 0.0-frameTolerance:
            # keep track of start time/frame for later
            image.frameNStart = frameN  # exact frame index
            image.tStart = t  # local t and not account for scr refresh
            image.tStartRefresh = tThisFlipGlobal  # on global time
            win.timeOnFlip(image, 'tStartRefresh')  # time at next scr refresh
            # add timestamp to datafile
            thisExp.timestampOnFlip(win, 'image.started')
            # update status
            image.status = STARTED
            image.setAutoDraw(True)
        
        # if image is active this frame...
        if image.status == STARTED:
            # update params
            pass
        
        # check for quit (typically the Esc key)
        if defaultKeyboard.getKeys(keyList=["escape"]):
            thisExp.status = FINISHED
        if thisExp.status == FINISHED or endExpNow:
            endExperiment(thisExp, win=win)
            return
        # pause experiment here if requested
        if thisExp.status == PAUSED:
            pauseExperiment(
                thisExp=thisExp, 
                win=win, 
                timers=[routineTimer], 
                playbackComponents=[]
            )
            # skip the frame we paused on
            continue
        
        # check if all components have finished
        if not continueRoutine:  # a component has requested a forced-end of Routine
            OJO.forceEnded = routineForceEnded = True
            break
        continueRoutine = False  # will revert to True if at least one component still running
        for thisComponent in OJO.components:
            if hasattr(thisComponent, "status") and thisComponent.status != FINISHED:
                continueRoutine = True
                break  # at least one component has not yet finished
        
        # refresh the screen
        if continueRoutine:  # don't flip if this routine is over or we'll get a blank screen
            win.flip()
    
    # --- Ending Routine "OJO" ---
    for thisComponent in OJO.components:
        if hasattr(thisComponent, "setAutoDraw"):
            thisComponent.setAutoDraw(False)
    # store stop times for OJO
    OJO.tStop = globalClock.getTime(format='float')
    OJO.tStopRefresh = tThisFlipGlobal
    thisExp.addData('OJO.stopped', OJO.tStop)
    # check responses
    if key_resp.keys in ['', [], None]:  # No response was made
        key_resp.keys = None
    thisExp.addData('key_resp.keys',key_resp.keys)
    if key_resp.keys != None:  # we had a response
        thisExp.addData('key_resp.rt', key_resp.rt)
        thisExp.addData('key_resp.duration', key_resp.duration)
    thisExp.nextEntry()
    # the Routine "OJO" was not non-slip safe, so reset the non-slip timer
    routineTimer.reset()
    
    # --- Prepare to start Routine "salir" ---
    # create an object to store info about Routine salir
    salir = data.Routine(
        name='salir',
        components=[text_13, key_resp_5],
    )
    salir.status = NOT_STARTED
    continueRoutine = True
    # update component parameters for each repeat
    # create starting attributes for key_resp_5
    key_resp_5.keys = []
    key_resp_5.rt = []
    _key_resp_5_allKeys = []
    # store start times for salir
    salir.tStartRefresh = win.getFutureFlipTime(clock=globalClock)
    salir.tStart = globalClock.getTime(format='float')
    salir.status = STARTED
    thisExp.addData('salir.started', salir.tStart)
    salir.maxDuration = None
    # keep track of which components have finished
    salirComponents = salir.components
    for thisComponent in salir.components:
        thisComponent.tStart = None
        thisComponent.tStop = None
        thisComponent.tStartRefresh = None
        thisComponent.tStopRefresh = None
        if hasattr(thisComponent, 'status'):
            thisComponent.status = NOT_STARTED
    # reset timers
    t = 0
    _timeToFirstFrame = win.getFutureFlipTime(clock="now")
    frameN = -1
    
    # --- Run Routine "salir" ---
    salir.forceEnded = routineForceEnded = not continueRoutine
    while continueRoutine:
        # get current time
        t = routineTimer.getTime()
        tThisFlip = win.getFutureFlipTime(clock=routineTimer)
        tThisFlipGlobal = win.getFutureFlipTime(clock=None)
        frameN = frameN + 1  # number of completed frames (so 0 is the first frame)
        # update/draw components on each frame
        
        # *text_13* updates
        
        # if text_13 is starting this frame...
        if text_13.status == NOT_STARTED and tThisFlip >= 0.0-frameTolerance:
            # keep track of start time/frame for later
            text_13.frameNStart = frameN  # exact frame index
            text_13.tStart = t  # local t and not account for scr refresh
            text_13.tStartRefresh = tThisFlipGlobal  # on global time
            win.timeOnFlip(text_13, 'tStartRefresh')  # time at next scr refresh
            # add timestamp to datafile
            thisExp.timestampOnFlip(win, 'text_13.started')
            # update status
            text_13.status = STARTED
            text_13.setAutoDraw(True)
        
        # if text_13 is active this frame...
        if text_13.status == STARTED:
            # update params
            pass
        
        # *key_resp_5* updates
        waitOnFlip = False
        
        # if key_resp_5 is starting this frame...
        if key_resp_5.status == NOT_STARTED and tThisFlip >= 0.0-frameTolerance:
            # keep track of start time/frame for later
            key_resp_5.frameNStart = frameN  # exact frame index
            key_resp_5.tStart = t  # local t and not account for scr refresh
            key_resp_5.tStartRefresh = tThisFlipGlobal  # on global time
            win.timeOnFlip(key_resp_5, 'tStartRefresh')  # time at next scr refresh
            # add timestamp to datafile
            thisExp.timestampOnFlip(win, 'key_resp_5.started')
            # update status
            key_resp_5.status = STARTED
            # keyboard checking is just starting
            waitOnFlip = True
            win.callOnFlip(key_resp_5.clock.reset)  # t=0 on next screen flip
            win.callOnFlip(key_resp_5.clearEvents, eventType='keyboard')  # clear events on next screen flip
        if key_resp_5.status == STARTED and not waitOnFlip:
            theseKeys = key_resp_5.getKeys(keyList=['space'], ignoreKeys=["escape"], waitRelease=False)
            _key_resp_5_allKeys.extend(theseKeys)
            if len(_key_resp_5_allKeys):
                key_resp_5.keys = _key_resp_5_allKeys[-1].name  # just the last key pressed
                key_resp_5.rt = _key_resp_5_allKeys[-1].rt
                key_resp_5.duration = _key_resp_5_allKeys[-1].duration
                # a response ends the routine
                continueRoutine = False
        
        # check for quit (typically the Esc key)
        if defaultKeyboard.getKeys(keyList=["escape"]):
            thisExp.status = FINISHED
        if thisExp.status == FINISHED or endExpNow:
            endExperiment(thisExp, win=win)
            return
        # pause experiment here if requested
        if thisExp.status == PAUSED:
            pauseExperiment(
                thisExp=thisExp, 
                win=win, 
                timers=[routineTimer], 
                playbackComponents=[]
            )
            # skip the frame we paused on
            continue
        
        # check if all components have finished
        if not continueRoutine:  # a component has requested a forced-end of Routine
            salir.forceEnded = routineForceEnded = True
            break
        continueRoutine = False  # will revert to True if at least one component still running
        for thisComponent in salir.components:
            if hasattr(thisComponent, "status") and thisComponent.status != FINISHED:
                continueRoutine = True
                break  # at least one component has not yet finished
        
        # refresh the screen
        if continueRoutine:  # don't flip if this routine is over or we'll get a blank screen
            win.flip()
    
    # --- Ending Routine "salir" ---
    for thisComponent in salir.components:
        if hasattr(thisComponent, "setAutoDraw"):
            thisComponent.setAutoDraw(False)
    # store stop times for salir
    salir.tStop = globalClock.getTime(format='float')
    salir.tStopRefresh = tThisFlipGlobal
    thisExp.addData('salir.stopped', salir.tStop)
    # check responses
    if key_resp_5.keys in ['', [], None]:  # No response was made
        key_resp_5.keys = None
    thisExp.addData('key_resp_5.keys',key_resp_5.keys)
    if key_resp_5.keys != None:  # we had a response
        thisExp.addData('key_resp_5.rt', key_resp_5.rt)
        thisExp.addData('key_resp_5.duration', key_resp_5.duration)
    thisExp.nextEntry()
    # the Routine "salir" was not non-slip safe, so reset the non-slip timer
    routineTimer.reset()
    
    # --- Prepare to start Routine "Trial_Operant" ---
    # create an object to store info about Routine Trial_Operant
    Trial_Operant = data.Routine(
        name='Trial_Operant',
        components=[imgCoin, txtOperantMsg, sndReinf, text_9],
    )
    Trial_Operant.status = NOT_STARTED
    continueRoutine = True
    # update component parameters for each repeat
    # Run 'Begin Routine' code from codeOperant
    # ====== Parámetros de grupo / umbrales ======
    TARGET_TRIALS = int(thisExp.extraInfo.get("target_trials", 8))
    MONEY_PER_TRIAL = int(thisExp.extraInfo.get("money_per_trial", 5))
    
    # Umbrales desde calibración (globals/extraInfo)
    thr70 = float(globals().get('thr70', thisExp.extraInfo.get("thr70", -1)))
    thr30 = float(globals().get('thr30', thisExp.extraInfo.get("thr30", -1)))
    if thr70 <= 0 or thr30 < 0 or thr30 >= thr70:
        raise RuntimeError(f"Umbrales inválidos: thr70={thr70}, thr30={thr30}")
    print(f"[Operant] Umbrales cargados: thr70={thr70:.1f} N, thr30={thr30:.1f} N")
    
    # ====== Parámetros auxiliares ======
    REINF_DUR = 0.25           # reforzador 0.25 s (display y sonido)
    BAND_N    = 2.0            # banda alrededor del pico (en N)
    
    # ====== Reloj global ======
    try:
        _ = globalClock.getTime()
    except NameError:
        globalClock = core.Clock()
    
    # ====== Máquina de estados y contadores ======
    state = 'available'          # 'available' | 'wait_low'
    reinforcer_on = False
    reinforcerClock = core.Clock()
    ending_after_this_reinforcer = False
    
    trial_idx = 0                # nº de reforzadores ENTREGADOS (1..N)
    earned_money = 0
    
    # Flag: tras reforzar, mientras no haya bajado de 70 seguimos etiquetando 'lag'
    post_reinf_above70 = False
    
    # ====== Intervalos entre reforzadores ======
    interval_active = False
    interval_start_time = None
    interval_peak_force = 0.0
    interval_peak_hold = 0.0
    interval_durations = []      # solo duraciones
    interval_log = []            # (start_s, end_s, duration_s)
    
    # Para el cálculo de dt por frame
    _prev_t = globalClock.getTime()
    
    # ====== Duración ≥70% por ensayo reforzado ======
    resp70_active = False
    resp70_dur_accum = 0.0
    resp70_durations = []
    
    # ====== Moneda (robusto a rutas) ======
    from pathlib import Path
    _units = getattr(win, 'units', 'height')
    _coin_size = (220, 220) if _units == 'pix' else (0.22, 0.22)
    _candidates = [
        Path.cwd() / "estimulos" / "5pesos.png",
        Path.cwd() / "resources" / "5pesos.png",
        Path.cwd() / "stimuli" / "5pesos.png",
        r"C:\Users\palom\OneDrive\Escritorio\UP-UNED Experimento\estimulos\5pesos.png",
        r"C:\Users\palom\OneDrive\Escritorio\prueba dinamometro\estimulos\5pesos.png",
    ]
    _coin_img = None
    for p in _candidates:
        try:
            if Path(p).is_file():
                _coin_img = str(p)
                break
        except Exception:
            pass
    
    try:
        if _coin_img:
            coinStim = visual.ImageStim(
                win=win, image=_coin_img, pos=(0, 0), size=_coin_size, opacity=1.0, autoLog=False
            )
            print(f"[Operant] coin image loaded: {_coin_img}")
        else:
            coinStim = visual.Circle(
                win=win, radius=_coin_size[0]/2 if _units == 'pix' else _coin_size[0]/2,
                pos=(0, 0), lineColor=None, fillColor=[0.8, 0.8, 0.8], autoLog=False
            )
            print("[Operant] usando placeholder circular en lugar de imagen")
    except Exception as e:
        coinStim = visual.Circle(
            win=win, radius=_coin_size[0]/2 if _units == 'pix' else _coin_size[0]/2,
            pos=(0, 0), lineColor=None, fillColor=[0.8, 0.8, 0.8], autoLog=False
        )
        print(f"[Operant] WARN: no se pudo cargar imagen de moneda: {e}")
    
    # Si hubiera un componente Builder llamado imgCoin, apágalo
    try:
        imgCoin.autoDraw = False
    except Exception:
        pass
    
    # Mensaje en pantalla (opcional)
    try:
        txtOperantMsg.text = "¡Ahora, descubre cómo ganar el mayor dinero posible! Cada moneda que obtengas se va sumando al final"
    except Exception:
        pass
    
    # ====== Timeseries continuo (compartido con Extinción) ======
    try:
        ts_all_rows
    except NameError:
        ts_all_rows = []
    phase_label = 'operant'      # etiqueta de fase para el CSV continuo
    
    # Reloj relativo de la fase
    operantClock = core.Clock()
    operantClock.reset()
    
    sndReinf.setSound('A', hamming=True)
    sndReinf.setVolume(1.0, log=False)
    sndReinf.seek(0)
    # store start times for Trial_Operant
    Trial_Operant.tStartRefresh = win.getFutureFlipTime(clock=globalClock)
    Trial_Operant.tStart = globalClock.getTime(format='float')
    Trial_Operant.status = STARTED
    thisExp.addData('Trial_Operant.started', Trial_Operant.tStart)
    Trial_Operant.maxDuration = None
    win.color = [0.4431, 0.0510, -0.9137]
    win.colorSpace = 'rgb'
    win.backgroundImage = ''
    win.backgroundFit = 'none'
    # keep track of which components have finished
    Trial_OperantComponents = Trial_Operant.components
    for thisComponent in Trial_Operant.components:
        thisComponent.tStart = None
        thisComponent.tStop = None
        thisComponent.tStartRefresh = None
        thisComponent.tStopRefresh = None
        if hasattr(thisComponent, 'status'):
            thisComponent.status = NOT_STARTED
    # reset timers
    t = 0
    _timeToFirstFrame = win.getFutureFlipTime(clock="now")
    frameN = -1
    
    # --- Run Routine "Trial_Operant" ---
    Trial_Operant.forceEnded = routineForceEnded = not continueRoutine
    while continueRoutine:
        # get current time
        t = routineTimer.getTime()
        tThisFlip = win.getFutureFlipTime(clock=routineTimer)
        tThisFlipGlobal = win.getFutureFlipTime(clock=None)
        frameN = frameN + 1  # number of completed frames (so 0 is the first frame)
        # update/draw components on each frame
        # Run 'Each Frame' code from codeOperant
        # ====== Tiempo global y dt ======
        _now = globalClock.getTime()
        dt = _now - _prev_t
        if dt < 0: dt = 0
        _prev_t = _now
        
        # ====== Lectura de fuerza (NO llamar gdx.read() aquí; usar sampler) ======
        try:
            f_now = float(_gdx_state.last)
        except Exception:
            s = gdx.read()
            f_now = float(s[0]) if (s and s[0] is not None) else 0.0
        
        # ====== Tracking de duración ≥70% por ensayo ======
        if resp70_active:
            if f_now >= thr70:
                resp70_dur_accum += dt
            else:
                resp70_active = False
                resp70_durations.append(resp70_dur_accum)
        
        # ====== Métricas dentro del intervalo activo (EXCLUYENDO display de reforzador) ======
        if interval_active and (not reinforcer_on):
            if f_now > interval_peak_force:
                interval_peak_force = f_now
                interval_peak_hold = 0.0
            else:
                if f_now >= max(0.0, interval_peak_force - BAND_N):
                    interval_peak_hold += dt
        
        # ====== Detección continua de refuerzos (FR1) ======
        if state == 'available':
            if f_now >= thr70:
                # Arranca cronómetro de duración ≥70% para ESTE ensayo
                resp70_active = True
                resp70_dur_accum = 0.0
        
                # Este refuerzo cierra el intervalo del ensayo PREVIO (trial_completed)
                trial_idx += 1
                earned_money = trial_idx * MONEY_PER_TRIAL
                trial_completed = trial_idx - 1  # el que se cierra ahora
        
                # Cerrar y loguear el ensayo PREVIO (i−1), si existía
                if interval_active:
                    interval_dur = _now - interval_start_time
        
                    # ✅ Acumular para extinción
                    interval_durations.append(interval_dur)
                    interval_log.append((interval_start_time, _now, interval_dur))
        
                    # Aseguramos pico y hold
                    if f_now > interval_peak_force:
                        interval_peak_force = f_now
                        interval_peak_hold = 0.0
                    elif f_now >= max(0.0, interval_peak_force - BAND_N):
                        interval_peak_hold += dt
        
                    # Duración ≥70% del ensayo completado
                    idx = trial_completed - 1
                    dur70 = resp70_durations[idx] if 0 <= idx < len(resp70_durations) else -1.0
        
                    # Log por ensayo (no afecta timeseries)
                    thisExp.addData('group', thisExp.extraInfo.get('group','NA'))
                    thisExp.addData('trial', trial_completed)
                    thisExp.addData('reinforced', 1)
                    thisExp.addData('trial_start_global', interval_start_time)
                    thisExp.addData('trial_end_global', _now)
                    thisExp.addData('trial_dur', interval_dur)
                    thisExp.addData('force_peak', interval_peak_force)
                    thisExp.addData('force_peak_hold', interval_peak_hold)
                    thisExp.addData('resp_count', 1)
                    earned_total_before = (trial_completed - 1) * MONEY_PER_TRIAL if trial_completed >= 1 else 0
                    thisExp.addData('earned_total_before', earned_total_before)
                    thisExp.addData('earned_in_trial',    MONEY_PER_TRIAL)
                    thisExp.addData('earned_total_after', trial_completed * MONEY_PER_TRIAL)
                    thisExp.addData('earned_money_total', trial_completed * MONEY_PER_TRIAL)
                    thisExp.addData('resp70_dur', dur70)
                    thisExp.addData('force_at_reinf', f_now)
                    thisExp.addData('thr70_used_N', thr70)
                    thisExp.addData('thr30_used_N', thr30)
                    thisExp.nextEntry()
        
                # Inicia NUEVO intervalo (ensayo actual i)
                interval_active = True
                interval_start_time = _now
                interval_peak_force = f_now
                interval_peak_hold = 0.0
        
                # Reforzador no-bloqueante (display + sonido)
                reinforcer_on = True
                reinforcerClock.reset()
                try:
                    sndReinf.stop()
                except Exception:
                    pass
                try:
                    sndReinf.play()
                except Exception:
                    pass
        
                # >>> Flag: desde ahora y hasta bajar de 70 etiquetamos 'lag'
                post_reinf_above70 = True
        
                if trial_idx >= TARGET_TRIALS:
                    ending_after_this_reinforcer = True
        
        elif state == 'wait_low':
            # Re-arme: esperar caer ≤30% para volver a permitir refuerzo
            if f_now <= thr30:
                state = 'available'
        
        # Si justo después del refuerzo sigue arriba de 30%, quedamos en 'wait_low'
        if reinforcer_on and state == 'available' and f_now > thr30:
            state = 'wait_low'
        
        # Terminar display del reforzador
        if reinforcer_on:
            coinStim.draw()
            if reinforcerClock.getTime() >= REINF_DUR:
                reinforcer_on = False
                try:
                    sndReinf.stop()
                except Exception:
                    pass
        
        # >>> Apagar el flag cuando baje por primera vez de 70 después del refuerzo
        if post_reinf_above70 and (f_now < thr70):
            post_reinf_above70 = False
        
        # ====== Etiqueta de estado (lag/reset/dwell/pursuit) ======
        if (f_now >= thr70) and (post_reinf_above70 or state == 'available'):
            ph_state = 'lag'
        else:
            if state == 'wait_low':
                ph_state = 'reset' if f_now > thr30 else 'dwell'
            else:  # available (rearmado)
                ph_state = 'pursuit' if (thr30 < f_now < thr70) else ('dwell' if f_now <= thr30 else 'lag')
        
        # ====== Etiqueta de bloque/ventana para timeseries ======
        try:
            N = int(TARGET_TRIALS)
        except Exception:
            N = int(thisExp.extraInfo.get("target_trials", 8))
        block_label = ""
        lo = max(1, N - 3)  # últimos 4 ensayos: N-3..N
        if lo <= int(trial_idx) <= N:
            block_label = "block 1"
        
        # ====== Timeseries continuo (NO generes el CSV aquí) ======
        t_op = operantClock.getTime()
        ts_all_rows.append((
            globalClock.getTime(), t_op, 'operant',
            float(f_now), ph_state, int(trial_idx), block_label
        ))
        
        # Cierre tras cumplir objetivo
        if ending_after_this_reinforcer and (not reinforcer_on) and (state != 'wait_low' or f_now <= thr30):
            continueRoutine = False
        
        
        # *imgCoin* updates
        
        # if imgCoin is starting this frame...
        if imgCoin.status == NOT_STARTED and tThisFlip >= 0-frameTolerance:
            # keep track of start time/frame for later
            imgCoin.frameNStart = frameN  # exact frame index
            imgCoin.tStart = t  # local t and not account for scr refresh
            imgCoin.tStartRefresh = tThisFlipGlobal  # on global time
            win.timeOnFlip(imgCoin, 'tStartRefresh')  # time at next scr refresh
            # add timestamp to datafile
            thisExp.timestampOnFlip(win, 'imgCoin.started')
            # update status
            imgCoin.status = STARTED
            imgCoin.setAutoDraw(True)
        
        # if imgCoin is active this frame...
        if imgCoin.status == STARTED:
            # update params
            imgCoin.setOpacity(0.0, log=False)
        
        # *txtOperantMsg* updates
        
        # if txtOperantMsg is starting this frame...
        if txtOperantMsg.status == NOT_STARTED and tThisFlip >= 0.0-frameTolerance:
            # keep track of start time/frame for later
            txtOperantMsg.frameNStart = frameN  # exact frame index
            txtOperantMsg.tStart = t  # local t and not account for scr refresh
            txtOperantMsg.tStartRefresh = tThisFlipGlobal  # on global time
            win.timeOnFlip(txtOperantMsg, 'tStartRefresh')  # time at next scr refresh
            # add timestamp to datafile
            thisExp.timestampOnFlip(win, 'txtOperantMsg.started')
            # update status
            txtOperantMsg.status = STARTED
            txtOperantMsg.setAutoDraw(True)
        
        # if txtOperantMsg is active this frame...
        if txtOperantMsg.status == STARTED:
            # update params
            pass
        
        # *sndReinf* updates
        
        # if sndReinf is stopping this frame...
        if sndReinf.status == STARTED:
            if bool(False) or sndReinf.isFinished:
                # keep track of stop time/frame for later
                sndReinf.tStop = t  # not accounting for scr refresh
                sndReinf.tStopRefresh = tThisFlipGlobal  # on global time
                sndReinf.frameNStop = frameN  # exact frame index
                # add timestamp to datafile
                thisExp.timestampOnFlip(win, 'sndReinf.stopped')
                # update status
                sndReinf.status = FINISHED
                sndReinf.stop()
        
        # *text_9* updates
        
        # if text_9 is starting this frame...
        if text_9.status == NOT_STARTED and tThisFlip >= 0.0-frameTolerance:
            # keep track of start time/frame for later
            text_9.frameNStart = frameN  # exact frame index
            text_9.tStart = t  # local t and not account for scr refresh
            text_9.tStartRefresh = tThisFlipGlobal  # on global time
            win.timeOnFlip(text_9, 'tStartRefresh')  # time at next scr refresh
            # add timestamp to datafile
            thisExp.timestampOnFlip(win, 'text_9.started')
            # update status
            text_9.status = STARTED
            text_9.setAutoDraw(True)
        
        # if text_9 is active this frame...
        if text_9.status == STARTED:
            # update params
            pass
        
        # check for quit (typically the Esc key)
        if defaultKeyboard.getKeys(keyList=["escape"]):
            thisExp.status = FINISHED
        if thisExp.status == FINISHED or endExpNow:
            endExperiment(thisExp, win=win)
            return
        # pause experiment here if requested
        if thisExp.status == PAUSED:
            pauseExperiment(
                thisExp=thisExp, 
                win=win, 
                timers=[routineTimer], 
                playbackComponents=[sndReinf]
            )
            # skip the frame we paused on
            continue
        
        # check if all components have finished
        if not continueRoutine:  # a component has requested a forced-end of Routine
            Trial_Operant.forceEnded = routineForceEnded = True
            break
        continueRoutine = False  # will revert to True if at least one component still running
        for thisComponent in Trial_Operant.components:
            if hasattr(thisComponent, "status") and thisComponent.status != FINISHED:
                continueRoutine = True
                break  # at least one component has not yet finished
        
        # refresh the screen
        if continueRoutine:  # don't flip if this routine is over or we'll get a blank screen
            win.flip()
    
    # --- Ending Routine "Trial_Operant" ---
    for thisComponent in Trial_Operant.components:
        if hasattr(thisComponent, "setAutoDraw"):
            thisComponent.setAutoDraw(False)
    # store stop times for Trial_Operant
    Trial_Operant.tStop = globalClock.getTime(format='float')
    Trial_Operant.tStopRefresh = tThisFlipGlobal
    thisExp.addData('Trial_Operant.stopped', Trial_Operant.tStop)
    setupWindow(expInfo=expInfo, win=win)
    # Run 'End Routine' code from codeOperant
    # ====== Ventana de extinción basada en R_N → t0 ======
    ext_last7_durations = interval_durations[-7:]  # puede ser <7 si hubo <8 refuerzos
    ext_last_open_start = interval_start_time if interval_active else None
    
    print(f"[Reinf→Ext] last7_durations={ext_last7_durations}")
    print(f"[Reinf→Ext] last_open_interval_start={ext_last_open_start}")
    print(f"[Reinf] N={trial_idx}  len(interval_durations)={len(interval_durations)}  "
          f"last7_sum={sum(ext_last7_durations) if ext_last7_durations else 0:.3f}  "
          f"open_start={ext_last_open_start}")
    
    # Ya no escribimos un CSV propio aquí; el CSV continuo se escribe al final de Extinción.
    thisExp.addData('reinforcement_trials_delivered', trial_idx)
    thisExp.addData('earned_money_total_endReinf', trial_idx * MONEY_PER_TRIAL)
    
    sndReinf.pause()  # ensure sound has stopped at end of Routine
    thisExp.nextEntry()
    # the Routine "Trial_Operant" was not non-slip safe, so reset the non-slip timer
    routineTimer.reset()
    
    # --- Prepare to start Routine "Extinction" ---
    # create an object to store info about Routine Extinction
    Extinction = data.Routine(
        name='Extinction',
        components=[txtExtMsg, text_12],
    )
    Extinction.status = NOT_STARTED
    continueRoutine = True
    # update component parameters for each repeat
    # Run 'Begin Routine' code from codeExt
    # ====== Parámetros para detección de t0 y ventana ======
    DETECT_HOLD = 0.0    # t0 = primer cruce a ≥70% (SIN hold)
    
    # ====== Grupo: el Control (28) NO hace ventana de extinción ======
    try:
        group = str(thisExp.extraInfo.get('group', 'NA'))
    except Exception:
        group = 'NA'
    CONTROL_NO_EXT = (group == 'Control')
    
    # ====== Insumos desde Trial_Operant ======
    try:
        last7 = list(ext_last7_durations)
    except NameError:
        last7 = []
    try:
        last_open_start = float(ext_last_open_start)
    except Exception:
        last_open_start = None
    
    # ====== Umbrales ======
    thr70 = float(globals().get('thr70', thisExp.extraInfo.get('thr70', -1)))
    thr30 = float(globals().get('thr30', thisExp.extraInfo.get('thr30', -1)))
    assert thr70 > 0, f"Falta thr70 de calibración (thr70={thr70})."
    
    # ====== Parámetros auxiliares ======
    TARGET_TRIALS   = int(thisExp.extraInfo.get("target_trials", 8))
    MONEY_PER_TRIAL = int(thisExp.extraInfo.get("money_per_trial", 5))
    BAND_N          = 2.0
    
    # ====== Relojes y estado ======
    try:
        _ = globalClock.getTime()
    except NameError:
        globalClock = core.Clock()
    
    detectClock = core.Clock()     # espera t0
    _prev_detect_t = 0.0
    _hold_accum = 0.0
    
    # Métricas tramo R_N → t0
    last_peak_force = 0.0
    last_peak_hold  = 0.0
    
    # Ventana desde t0
    extWindowClock = core.Clock()
    awaiting_t0 = True
    t0_rel = None
    t0_global = None
    EXT_DUR = None
    EXT_4_DUR = None
    
    # Flags/marcas
    ext_mark4_fired = False
    ext_mark8_fired = False
    ext_mark4_rel = -1.0
    ext_mark8_rel = -1.0
    ext_mark4_global = -1.0
    ext_mark8_global = -1.0
    
    # Acumuladores ventana
    _prev_t_win = 0.0
    auc_ext = 0.0
    peak_ext = 0.0
    time_above70 = 0.0
    
    # ====== FSM 70→30→30→70 para detectar respuestas en extinción ======
    ext_resp_active = False
    ext_resp_dur = 0.0
    ext_resp_peak = 0.0
    ext_responses = []
    arm_state = 'armed'  # se reajusta al confirmar t0
    
    # Continuidad del tracking ≥70% del ensayo N
    try:
        _ = resp70_active
    except NameError:
        resp70_active = False
        resp70_dur_accum = 0.0
    try:
        _ = resp70_durations
    except NameError:
        resp70_durations = []
    
    # ====== UI (opcional) ======
    try:
        txtExtMsg.text = "¡Ahora, descubre cómo ganar el mayor dinero posible! Cada moneda que obtengas se va sumando al final"
    except Exception:
        pass
    
    # ====== Timeseries continuo (compartido con Operante) ======
    try:
        ts_all_rows
    except NameError:
        ts_all_rows = []
    phase_label = 'extinction'
    
    print(f"[Ext] last7_len={len(last7)}  last7_sum={(sum(last7) if last7 else 0.0):.3f} s  last_open_start={last_open_start}  thr70={thr70:.2f}")
    
    # store start times for Extinction
    Extinction.tStartRefresh = win.getFutureFlipTime(clock=globalClock)
    Extinction.tStart = globalClock.getTime(format='float')
    Extinction.status = STARTED
    thisExp.addData('Extinction.started', Extinction.tStart)
    Extinction.maxDuration = None
    win.color = [0.4431, 0.0510, -0.9137]
    win.colorSpace = 'rgb'
    win.backgroundImage = ''
    win.backgroundFit = 'none'
    # keep track of which components have finished
    ExtinctionComponents = Extinction.components
    for thisComponent in Extinction.components:
        thisComponent.tStart = None
        thisComponent.tStop = None
        thisComponent.tStartRefresh = None
        thisComponent.tStopRefresh = None
        if hasattr(thisComponent, 'status'):
            thisComponent.status = NOT_STARTED
    # reset timers
    t = 0
    _timeToFirstFrame = win.getFutureFlipTime(clock="now")
    frameN = -1
    
    # --- Run Routine "Extinction" ---
    Extinction.forceEnded = routineForceEnded = not continueRoutine
    while continueRoutine:
        # get current time
        t = routineTimer.getTime()
        tThisFlip = win.getFutureFlipTime(clock=routineTimer)
        tThisFlipGlobal = win.getFutureFlipTime(clock=None)
        frameN = frameN + 1  # number of completed frames (so 0 is the first frame)
        # update/draw components on each frame
        # Run 'Each Frame' code from codeExt
        # ====== A) Esperar t0 (primer cruce a ≥70%, SIN hold) ======
        if awaiting_t0:
            t_rel = detectClock.getTime()
            dt_det = t_rel - _prev_detect_t
            if dt_det < 0:
                dt_det = 0.0
            _prev_detect_t = t_rel
        
            # Lectura fuerza (usar sampler)
            try:
                f_now = float(_gdx_state.last)
            except Exception:
                s = gdx.read()
                f_now = float(s[0]) if (s and s[0] is not None) else 0.0
        
            # Continuar/terminar resp70 del ensayo N (mismo comportamiento que antes)
            if resp70_active:
                if f_now >= thr70:
                    resp70_dur_accum += dt_det
                else:
                    resp70_active = False
                    resp70_durations.append(resp70_dur_accum)
        
            # Métricas R_N→t0 (pico y 'hold' alrededor del pico)
            if f_now > last_peak_force:
                last_peak_force = f_now
                last_peak_hold  = 0.0
            else:
                if f_now >= max(0.0, last_peak_force - BAND_N):
                    last_peak_hold += dt_det
        
            # ---------- t0 instantáneo (SIN hold) ----------
            if f_now >= thr70:
                t0_rel = t_rel
                t0_global = globalClock.getTime()
        
                # Cerrar el último intervalo abierto (ensayo N reforzado) justo en t0
                if (last_open_start is not None) and (t0_global >= last_open_start):
                    interval_dur = t0_global - last_open_start
                    N = TARGET_TRIALS
                    earned_total_before = (N - 1) * MONEY_PER_TRIAL
                    earned_total_after  = earned_total_before + MONEY_PER_TRIAL
                    idxN = N - 1
                    dur70_N = resp70_durations[idxN] if 0 <= idxN < len(resp70_durations) else -1.0
        
                    thisExp.addData('group', thisExp.extraInfo.get('group','NA'))
                    thisExp.addData('trial', N)
                    thisExp.addData('reinforced', 1)
                    thisExp.addData('trial_start_global', last_open_start)
                    thisExp.addData('trial_end_global', t0_global)
                    thisExp.addData('trial_dur', interval_dur)
                    thisExp.addData('force_peak', last_peak_force)
                    thisExp.addData('force_peak_hold', last_peak_hold)
                    thisExp.addData('resp_count', 1)
                    thisExp.addData('earned_total_before', earned_total_before)
                    thisExp.addData('earned_in_trial',    MONEY_PER_TRIAL)
                    thisExp.addData('earned_total_after', earned_total_after)
                    thisExp.addData('earned_money_total', earned_total_after)
                    thisExp.addData('resp70_dur', dur70_N)
                    thisExp.addData('closed_in_extinction', 1)
                    thisExp.nextEntry()
        
                # Grupo Control: no ventana; acabar en t0
                if CONTROL_NO_EXT:
                    thisExp.addData('ext_skipped_control', 1)
                    thisExp.addData('ext_t0_rel', t0_rel if t0_rel is not None else -1)
                    thisExp.addData('ext_t0_global', t0_global if t0_global is not None else -1)
                    try:
                        last_cycle_RN_to_t0 = (t0_global - last_open_start) if (t0_global is not None and last_open_start is not None) else -1.0
                    except NameError:
                        last_cycle_RN_to_t0 = -1.0
                    thisExp.addData('ext_last_cycle_RN_to_t0', last_cycle_RN_to_t0)
                    thisExp.addData('ext_dur', 0.0)
                    thisExp.addData('ext_resp_count', 0)
                    thisExp.nextEntry()
        
                    continueRoutine = False  # Fin inmediato en grupo Control
                else:
                    # Duración ventana: “últimos 4 × 2”
                    if (last_open_start is not None) and (t0_global >= last_open_start):
                        rn_to_t0   = (t0_global - last_open_start)
                        closed3_sum = sum(last7[-3:]) if last7 else 0.0
                        EXT_4_DUR  = closed3_sum + rn_to_t0
                        EXT_DUR    = 2.0 * EXT_4_DUR
                    else:
                        rn_to_t0   = 0.0
                        closed3_sum = sum(last7[-3:]) if last7 else 0.0
                        EXT_4_DUR  = closed3_sum if closed3_sum > 0 else 70.0
                        EXT_DUR    = 2.0 * EXT_4_DUR
        
                    print(f"[Ext] t0_rel={t0_rel:.3f}s  t0_global={t0_global:.3f}s  EXT_4_DUR={EXT_4_DUR:.3f}s  EXT_DUR(=2×4)={EXT_DUR:.3f}s")
        
                    # Arrancar ventana
                    extWindowClock.reset()
                    awaiting_t0 = False
                    _prev_t_win = 0.0
                    arm_state = 'need_below70'  # criterio 70→30→30→70
        
                # Nota: no añadimos fila pre-t0 en este frame; pasa directo a ventana.
            else:
                # ------- Pre-t0: distinguir dwell/pursuit (sin 'lag' antes de t0) -------
                if f_now <= thr30:
                    ph_state = 'dwell'
                elif f_now < thr70:
                    ph_state = 'pursuit'
                else:
                    # En teoría no entramos aquí porque f_now≥thr70 dispara t0 arriba
                    ph_state = 'reset'
        
                block_label = "block 1"  # seguimos dentro del ensayo N hasta t0
                ts_all_rows.append((
                    globalClock.getTime(), t_rel, 'extinction',
                    float(f_now), ph_state, int(TARGET_TRIALS), block_label
                ))
        
        # ====== B) Ventana activa (t0 → t0 + EXT_DUR) ======
        else:
            if CONTROL_NO_EXT:
                continueRoutine = False
            else:
                t_win = extWindowClock.getTime()
                if t_win >= float(EXT_DUR):
                    if not ext_mark8_fired:
                        ext_mark8_fired = True
                        ext_mark8_rel = float(EXT_DUR)
                        ext_mark8_global = (t0_global + float(EXT_DUR)) if (t0_global is not None) else -1.0
        
                    if ext_resp_active:
                        ext_responses.append((ext_resp_dur, ext_resp_peak))
                        ext_resp_active = False
                    continueRoutine = False
                else:
                    # Lectura fuerza
                    try:
                        f_now = float(_gdx_state.last)
                    except Exception:
                        s = gdx.read()
                        f_now = float(s[0]) if (s and s[0] is not None) else 0.0
        
                    # dt (relativo a la ventana)
                    dt = t_win - _prev_t_win
                    if dt < 0:
                        dt = 0.0
                    _prev_t_win = t_win
        
                    # Métricas ventana
                    auc_ext += f_now * dt
                    if f_now > peak_ext:
                        peak_ext = f_now
                    if f_now >= thr70:
                        time_above70 += dt
        
                    # Marca “últimos 4”
                    if (not ext_mark4_fired) and (EXT_4_DUR is not None) and (t_win >= float(EXT_4_DUR)):
                        ext_mark4_fired = True
                        ext_mark4_rel = t_win
                        ext_mark4_global = (t0_global + t_win) if (t0_global is not None) else -1.0
        
                    # ====== FSM 70→30→30→70 para respuestas en ventana ======
                    if arm_state == 'need_below70':
                        if f_now < thr70:
                            arm_state = 'need_below30'
                    elif arm_state == 'need_below30':
                        if f_now <= thr30:
                            arm_state = 'need_above30'
                    elif arm_state == 'need_above30':
                        if f_now >= thr30:
                            arm_state = 'armed'
                    elif arm_state == 'armed':
                        if (not ext_resp_active) and (f_now >= thr70):
                            # Nueva respuesta
                            ext_resp_active = True
                            ext_resp_dur = 0.0
                            ext_resp_peak = f_now
                            arm_state = 'need_below70'
        
                    # Actualización de respuesta activa
                    if ext_resp_active:
                        if f_now >= thr70:
                            ext_resp_dur += dt
                            if f_now > ext_resp_peak:
                                ext_resp_peak = f_now
                        else:
                            ext_responses.append((ext_resp_dur, ext_resp_peak))
                            ext_resp_active = False
        
                    # ====== Etiqueta de estado (ventana) ======
                    if arm_state == 'need_below70':
                        ph_state = 'lag' if f_now >= thr70 else 'reset'
                    elif arm_state == 'need_below30':
                        ph_state = 'reset'
                    elif arm_state == 'need_above30':
                        ph_state = 'dwell' if f_now <= thr30 else 'pursuit'
                    elif arm_state == 'armed':
                        if f_now >= thr70:
                            ph_state = 'lag'
                        elif f_now <= thr30:
                            ph_state = 'dwell'
                        else:
                            ph_state = 'pursuit'
                    else:
                        ph_state = 'pursuit'
        
                    # ----- Etiqueta de bloque/ventana para timeseries
                    if (EXT_4_DUR is not None) and (t_win < float(EXT_4_DUR)):
                        block_label = "extinction_conf"
                    else:
                        block_label = "extinction_expl"
        
                    # Serie temporal continua
                    ts_all_rows.append((
                        globalClock.getTime(), t_win, 'extinction',
                        float(f_now), ph_state, int(TARGET_TRIALS), block_label
                    ))
        
        
        # *txtExtMsg* updates
        
        # if txtExtMsg is starting this frame...
        if txtExtMsg.status == NOT_STARTED and tThisFlip >= 0.0-frameTolerance:
            # keep track of start time/frame for later
            txtExtMsg.frameNStart = frameN  # exact frame index
            txtExtMsg.tStart = t  # local t and not account for scr refresh
            txtExtMsg.tStartRefresh = tThisFlipGlobal  # on global time
            win.timeOnFlip(txtExtMsg, 'tStartRefresh')  # time at next scr refresh
            # add timestamp to datafile
            thisExp.timestampOnFlip(win, 'txtExtMsg.started')
            # update status
            txtExtMsg.status = STARTED
            txtExtMsg.setAutoDraw(True)
        
        # if txtExtMsg is active this frame...
        if txtExtMsg.status == STARTED:
            # update params
            pass
        
        # *text_12* updates
        
        # if text_12 is starting this frame...
        if text_12.status == NOT_STARTED and tThisFlip >= 0.0-frameTolerance:
            # keep track of start time/frame for later
            text_12.frameNStart = frameN  # exact frame index
            text_12.tStart = t  # local t and not account for scr refresh
            text_12.tStartRefresh = tThisFlipGlobal  # on global time
            win.timeOnFlip(text_12, 'tStartRefresh')  # time at next scr refresh
            # add timestamp to datafile
            thisExp.timestampOnFlip(win, 'text_12.started')
            # update status
            text_12.status = STARTED
            text_12.setAutoDraw(True)
        
        # if text_12 is active this frame...
        if text_12.status == STARTED:
            # update params
            pass
        
        # check for quit (typically the Esc key)
        if defaultKeyboard.getKeys(keyList=["escape"]):
            thisExp.status = FINISHED
        if thisExp.status == FINISHED or endExpNow:
            endExperiment(thisExp, win=win)
            return
        # pause experiment here if requested
        if thisExp.status == PAUSED:
            pauseExperiment(
                thisExp=thisExp, 
                win=win, 
                timers=[routineTimer], 
                playbackComponents=[]
            )
            # skip the frame we paused on
            continue
        
        # check if all components have finished
        if not continueRoutine:  # a component has requested a forced-end of Routine
            Extinction.forceEnded = routineForceEnded = True
            break
        continueRoutine = False  # will revert to True if at least one component still running
        for thisComponent in Extinction.components:
            if hasattr(thisComponent, "status") and thisComponent.status != FINISHED:
                continueRoutine = True
                break  # at least one component has not yet finished
        
        # refresh the screen
        if continueRoutine:  # don't flip if this routine is over or we'll get a blank screen
            win.flip()
    
    # --- Ending Routine "Extinction" ---
    for thisComponent in Extinction.components:
        if hasattr(thisComponent, "setAutoDraw"):
            thisComponent.setAutoDraw(False)
    # store stop times for Extinction
    Extinction.tStop = globalClock.getTime(format='float')
    Extinction.tStopRefresh = tThisFlipGlobal
    thisExp.addData('Extinction.stopped', Extinction.tStop)
    setupWindow(expInfo=expInfo, win=win)
    # Run 'End Routine' code from codeExt
    # ====== Parar el sampler antes de guardar ======
    try:
        _gdx_state.stop = True
    except Exception:
        pass
    try:
        if '_gdx_thread' in globals() and _gdx_thread is not None:
            _gdx_thread.join(timeout=1.0)
    except Exception:
        pass
    
    # ====== Resumen de extinción ======
    if CONTROL_NO_EXT:
        print(f"[Ext] Control: cerrado ensayo N en t0 y saltado End.")
    else:
        if (EXT_DUR is None) or (EXT_DUR <= 0):
            EXT_DUR = 0.0
        mean_ext = (auc_ext / EXT_DUR) if EXT_DUR > 0 else 0.0
    
        thisExp.addData('ext_t0_rel', t0_rel if t0_rel is not None else -1)
        thisExp.addData('ext_t0_global', t0_global if t0_global is not None else -1)
        thisExp.addData('ext_dur', EXT_DUR)
        thisExp.addData('ext_peakN', peak_ext)
        thisExp.addData('ext_meanN', mean_ext)
        thisExp.addData('ext_timeAboveThr70', time_above70)
    
        try:
            last_cycle_RN_to_t0 = (t0_global - last_open_start) if (t0_global is not None and last_open_start is not None) else -1.0
        except NameError:
            last_cycle_RN_to_t0 = -1.0
        thisExp.addData('ext_last_cycle_RN_to_t0', last_cycle_RN_to_t0)
    
        # Guardar respuestas en extinción (criterio 70→30→30→70)
        try:
            if ext_resp_active:
                ext_responses.append((ext_resp_dur, ext_resp_peak))
                ext_resp_active = False
        except NameError:
            pass
        thisExp.addData('ext_resp_count', len(ext_responses))
        for i, (dur, peak) in enumerate(ext_responses, start=1):
            thisExp.addData(f'ext_resp{i}_dur', dur)
            thisExp.addData(f'ext_resp{i}_peak', peak)
    
        # Marcas 4/8
        thisExp.addData('ext_mark4_rel', ext_mark4_rel)
        thisExp.addData('ext_mark4_global', ext_mark4_global)
        if ext_mark8_rel < 0 and EXT_DUR is not None:
            ext_mark8_rel = float(EXT_DUR)
            ext_mark8_global = (t0_global + float(EXT_DUR)) if (t0_global is not None) else -1.0
        thisExp.addData('ext_mark8_rel', ext_mark8_rel)
        thisExp.addData('ext_mark8_global', ext_mark8_global)
    
        print(f"[Ext] End window at t = {extWindowClock.getTime():.3f}s  (planned {EXT_DUR:.3f}s)")
        thisExp.nextEntry()
    
    # ====== Guardar CSV CONTINUO (operant + extinction) ======
    ts_path = thisExp.dataFileName + "_timeseries_all.csv"
    try:
        with open(ts_path, 'w', encoding='utf-8') as f:
            # Sin t_rel_phase_s ni phase
            f.write("t_abs_s,force_N,state,block,trial_idx\n")
            for row in ts_all_rows:
                if not isinstance(row, (list, tuple)):
                    continue
    
                # Formatos posibles que generamos:
                # 6 campos:  [t_abs, t_rel, phase, fN, state, trial]
                # 7 campos:  [t_abs, t_rel, phase, fN, state, trial, block]
                #            [t_abs, t_rel, phase, fN, state, block, trial]
                if len(row) >= 6:
                    t_abs = float(row[0])
                    fN    = float(row[3])
                    st    = str(row[4])
    
                    blk = ""
                    tr  = -1
    
                    if len(row) == 6:
                        tr = int(row[5])
                    else:
                        a, b = row[5], row[6]
                        # Detecta orden de (block, trial_idx)
                        if isinstance(a, (int, float)) and isinstance(b, str):
                            tr, blk = int(a), str(b)
                        elif isinstance(a, str) and isinstance(b, (int, float)):
                            blk, tr = str(a), int(b)
                        else:
                            # Fallback robusto
                            try:
                                tr = int(a); blk = str(b)
                            except Exception:
                                try:
                                    tr = int(b); blk = str(a)
                                except Exception:
                                    blk = str(a); tr = -1
    
                    blk_str = "" if blk is None else str(blk)
                    f.write(f"{t_abs:.6f},{fN:.4f},{st},{blk_str},{tr}\n")
                # Si llega una fila con otro largo, la ignoramos de forma segura.
    
        thisExp.addData('timeseries_all_file', ts_path)
    except Exception as e:
        thisExp.addData('timeseries_all_file_error', str(e))
    
    thisExp.nextEntry()
    # the Routine "Extinction" was not non-slip safe, so reset the non-slip timer
    routineTimer.reset()
    
    # --- Prepare to start Routine "End" ---
    # create an object to store info about Routine End
    End = data.Routine(
        name='End',
        components=[txtSummary, txtThanks, kbEnd],
    )
    End.status = NOT_STARTED
    continueRoutine = True
    # update component parameters for each repeat
    # Run 'Begin Routine' code from codeEnd
    # ------- Mostrar dinero total y mensaje final -------
    
    # 1) Intenta tomar el total desde extraInfo (si lo guardamos en la fase de reforzamiento)
    earned_total = thisExp.extraInfo.get('earned_money_total', None)
    
    # 2) Si no existe, caer al número de ensayos objetivo (aprox. correcta si completó todos)
    if earned_total is None:
        try:
            earned_total = int(thisExp.extraInfo.get('target_trials', 0)) * 5
        except Exception:
            earned_total = 0
    
    # 3) Construir textos
    txtSummary.text = f"Dinero total acumulado: ${earned_total} pesos"
    txtThanks.text  = "Pantera, ¡muchas gracias por ayudar a la ciencia! :)\n\nPresiona ESPACIO para finalizar."
    
    # create starting attributes for kbEnd
    kbEnd.keys = []
    kbEnd.rt = []
    _kbEnd_allKeys = []
    # store start times for End
    End.tStartRefresh = win.getFutureFlipTime(clock=globalClock)
    End.tStart = globalClock.getTime(format='float')
    End.status = STARTED
    thisExp.addData('End.started', End.tStart)
    End.maxDuration = None
    # keep track of which components have finished
    EndComponents = End.components
    for thisComponent in End.components:
        thisComponent.tStart = None
        thisComponent.tStop = None
        thisComponent.tStartRefresh = None
        thisComponent.tStopRefresh = None
        if hasattr(thisComponent, 'status'):
            thisComponent.status = NOT_STARTED
    # reset timers
    t = 0
    _timeToFirstFrame = win.getFutureFlipTime(clock="now")
    frameN = -1
    
    # --- Run Routine "End" ---
    End.forceEnded = routineForceEnded = not continueRoutine
    while continueRoutine:
        # get current time
        t = routineTimer.getTime()
        tThisFlip = win.getFutureFlipTime(clock=routineTimer)
        tThisFlipGlobal = win.getFutureFlipTime(clock=None)
        frameN = frameN + 1  # number of completed frames (so 0 is the first frame)
        # update/draw components on each frame
        
        # *txtSummary* updates
        
        # if txtSummary is starting this frame...
        if txtSummary.status == NOT_STARTED and tThisFlip >= 0.0-frameTolerance:
            # keep track of start time/frame for later
            txtSummary.frameNStart = frameN  # exact frame index
            txtSummary.tStart = t  # local t and not account for scr refresh
            txtSummary.tStartRefresh = tThisFlipGlobal  # on global time
            win.timeOnFlip(txtSummary, 'tStartRefresh')  # time at next scr refresh
            # add timestamp to datafile
            thisExp.timestampOnFlip(win, 'txtSummary.started')
            # update status
            txtSummary.status = STARTED
            txtSummary.setAutoDraw(True)
        
        # if txtSummary is active this frame...
        if txtSummary.status == STARTED:
            # update params
            pass
        
        # *txtThanks* updates
        
        # if txtThanks is starting this frame...
        if txtThanks.status == NOT_STARTED and tThisFlip >= 0.0-frameTolerance:
            # keep track of start time/frame for later
            txtThanks.frameNStart = frameN  # exact frame index
            txtThanks.tStart = t  # local t and not account for scr refresh
            txtThanks.tStartRefresh = tThisFlipGlobal  # on global time
            win.timeOnFlip(txtThanks, 'tStartRefresh')  # time at next scr refresh
            # add timestamp to datafile
            thisExp.timestampOnFlip(win, 'txtThanks.started')
            # update status
            txtThanks.status = STARTED
            txtThanks.setAutoDraw(True)
        
        # if txtThanks is active this frame...
        if txtThanks.status == STARTED:
            # update params
            pass
        
        # *kbEnd* updates
        waitOnFlip = False
        
        # if kbEnd is starting this frame...
        if kbEnd.status == NOT_STARTED and tThisFlip >= 0.0-frameTolerance:
            # keep track of start time/frame for later
            kbEnd.frameNStart = frameN  # exact frame index
            kbEnd.tStart = t  # local t and not account for scr refresh
            kbEnd.tStartRefresh = tThisFlipGlobal  # on global time
            win.timeOnFlip(kbEnd, 'tStartRefresh')  # time at next scr refresh
            # add timestamp to datafile
            thisExp.timestampOnFlip(win, 'kbEnd.started')
            # update status
            kbEnd.status = STARTED
            # keyboard checking is just starting
            waitOnFlip = True
            win.callOnFlip(kbEnd.clock.reset)  # t=0 on next screen flip
            win.callOnFlip(kbEnd.clearEvents, eventType='keyboard')  # clear events on next screen flip
        if kbEnd.status == STARTED and not waitOnFlip:
            theseKeys = kbEnd.getKeys(keyList=['space'], ignoreKeys=["escape"], waitRelease=False)
            _kbEnd_allKeys.extend(theseKeys)
            if len(_kbEnd_allKeys):
                kbEnd.keys = _kbEnd_allKeys[-1].name  # just the last key pressed
                kbEnd.rt = _kbEnd_allKeys[-1].rt
                kbEnd.duration = _kbEnd_allKeys[-1].duration
                # a response ends the routine
                continueRoutine = False
        
        # check for quit (typically the Esc key)
        if defaultKeyboard.getKeys(keyList=["escape"]):
            thisExp.status = FINISHED
        if thisExp.status == FINISHED or endExpNow:
            endExperiment(thisExp, win=win)
            return
        # pause experiment here if requested
        if thisExp.status == PAUSED:
            pauseExperiment(
                thisExp=thisExp, 
                win=win, 
                timers=[routineTimer], 
                playbackComponents=[]
            )
            # skip the frame we paused on
            continue
        
        # check if all components have finished
        if not continueRoutine:  # a component has requested a forced-end of Routine
            End.forceEnded = routineForceEnded = True
            break
        continueRoutine = False  # will revert to True if at least one component still running
        for thisComponent in End.components:
            if hasattr(thisComponent, "status") and thisComponent.status != FINISHED:
                continueRoutine = True
                break  # at least one component has not yet finished
        
        # refresh the screen
        if continueRoutine:  # don't flip if this routine is over or we'll get a blank screen
            win.flip()
    
    # --- Ending Routine "End" ---
    for thisComponent in End.components:
        if hasattr(thisComponent, "setAutoDraw"):
            thisComponent.setAutoDraw(False)
    # store stop times for End
    End.tStop = globalClock.getTime(format='float')
    End.tStopRefresh = tThisFlipGlobal
    thisExp.addData('End.stopped', End.tStop)
    # check responses
    if kbEnd.keys in ['', [], None]:  # No response was made
        kbEnd.keys = None
    thisExp.addData('kbEnd.keys',kbEnd.keys)
    if kbEnd.keys != None:  # we had a response
        thisExp.addData('kbEnd.rt', kbEnd.rt)
        thisExp.addData('kbEnd.duration', kbEnd.duration)
    thisExp.nextEntry()
    # the Routine "End" was not non-slip safe, so reset the non-slip timer
    routineTimer.reset()
    # Run 'End Experiment' code from codeEnd
    # ------- Commit de la asignación (avanzar índice SOLO al terminar) -------
    from pathlib import Path
    try:
        used_idx = int(thisExp.extraInfo.get("assign_idx"))
    except Exception:
        used_idx = None
    
    try:
        data_dir = Path(thisExp.dataFileName).parent
        idx_path = data_dir / "assign_idx.txt"
        if used_idx is not None:
            try:
                old_val = int(idx_path.read_text(encoding="utf-8").strip()) if idx_path.exists() else used_idx
            except Exception:
                old_val = used_idx
            # Avanza si el archivo no está por delante (robusto si solo hay una estación)
            if old_val <= used_idx:
                idx_path.write_text(str(used_idx + 1), encoding="utf-8")
                print(f"[Assign] committed index → {used_idx + 1}")
            else:
                print(f"[Assign] idx ya avanzó a {old_val}; no se modifica.")
    except Exception as e:
        print(f"[Assign] WARN: no se pudo avanzar indice: {e}")
    
    # ------- Cierre ordenado del dinamómetro / sampler -------
    try:
        if '_gdx_state' in globals() and hasattr(_gdx_state, 'stop') and (_gdx_state.stop is not True):
            _gdx_state.stop = True
            if '_gdx_thread' in globals() and _gdx_thread is not None:
                _gdx_thread.join(timeout=0.5)
    except Exception:
        pass
    
    try:
        gdx.stop()
        gdx.close()
    except Exception:
        pass
    
    
    # mark experiment as finished
    endExperiment(thisExp, win=win)


def saveData(thisExp):
    """
    Save data from this experiment
    
    Parameters
    ==========
    thisExp : psychopy.data.ExperimentHandler
        Handler object for this experiment, contains the data to save and information about 
        where to save it to.
    """
    filename = thisExp.dataFileName
    # these shouldn't be strictly necessary (should auto-save)
    thisExp.saveAsWideText(filename + '.csv', delim='auto')
    thisExp.saveAsPickle(filename)


def endExperiment(thisExp, win=None):
    """
    End this experiment, performing final shut down operations.
    
    This function does NOT close the window or end the Python process - use `quit` for this.
    
    Parameters
    ==========
    thisExp : psychopy.data.ExperimentHandler
        Handler object for this experiment, contains the data to save and information about 
        where to save it to.
    win : psychopy.visual.Window
        Window for this experiment.
    """
    if win is not None:
        # remove autodraw from all current components
        win.clearAutoDraw()
        # Flip one final time so any remaining win.callOnFlip() 
        # and win.timeOnFlip() tasks get executed
        win.flip()
    # return console logger level to WARNING
    logging.console.setLevel(logging.WARNING)
    # mark experiment handler as finished
    thisExp.status = FINISHED
    logging.flush()


def quit(thisExp, win=None, thisSession=None):
    """
    Fully quit, closing the window and ending the Python process.
    
    Parameters
    ==========
    win : psychopy.visual.Window
        Window to close.
    thisSession : psychopy.session.Session or None
        Handle of the Session object this experiment is being run from, if any.
    """
    thisExp.abort()  # or data files will save again on exit
    # make sure everything is closed down
    if win is not None:
        # Flip one final time so any remaining win.callOnFlip() 
        # and win.timeOnFlip() tasks get executed before quitting
        win.flip()
        win.close()
    logging.flush()
    if thisSession is not None:
        thisSession.stop()
    # terminate Python process
    core.quit()


# if running this experiment as a script...
if __name__ == '__main__':
    # call all functions in order
    expInfo = showExpInfoDlg(expInfo=expInfo)
    thisExp = setupData(expInfo=expInfo)
    logFile = setupLogging(filename=thisExp.dataFileName)
    win = setupWindow(expInfo=expInfo)
    setupDevices(expInfo=expInfo, thisExp=thisExp, win=win)
    run(
        expInfo=expInfo, 
        thisExp=thisExp, 
        win=win,
        globalClock='float'
    )
    saveData(thisExp=thisExp)
    quit(thisExp=thisExp, win=win)
```
