## Supplementary material for "Evidence for the extinction burst in grip force: A preregistered study": Electronic Supplementary Materials: gdx.py

```
# To use Go Direct sensors with Python 3 you must install the godirect module
# with the command: pip3 install godirect

from godirect import GoDirect

import logging
import time    

# gdx_vpython.py contains functions for a canvas with data collection buttons
from gdx import gdx_vpython
vp = gdx_vpython.ver_vpython()


logging.basicConfig()
#logging.getLogger('godirect').setLevel(logging.DEBUG)
#logging.getLogger('bleak').setLevel(logging.DEBUG)
#logging.getLogger('pygatt').setLevel(logging.DEBUG)


class gdx:

    # 1.0.0 is the first time a version was created. This was when the VPython functions were added
    VERSION = "1.0.0"    
    
    # Variables passed between the gdx functions.

    # devices - a 1D list of the connected Go Direct device objects.
    devices = []
    # device_sensors - a 2D list of the sensor numbers selected by the user [[1,2],[1]]
    device_sensors = []
    # enabled_sensors - a 2D list of the sensor objects that have been enabled for data collection.
    enabled_sensors = []
    # buffer - a 2D list to store the excess data from a sensor when multi-points are collected from a read due to fast collection.
    buffer = []
    # ble_open - this is a flag to keep track of when godirect is asked to open ble, to make sure it's not asked twice.
    ble_open = False
    # is this a vpython program
    vpython = False
    # is there a vpython buttons?
    vpython_buttons = False
    # is there a vpython chart?
    vpython_chart = False
    # is there a vpython meters?
    vpython_meters = False
    # is there a vpython slider for sample period?
    vpython_slider = False
    # is this the first time calling start() in a vpython program?
    vp_first_start = True
    # store the period. may be used in vp_start_button()
    period = 100
    # vp start button flag. Use this to determine if gdx.start() or gdx.stop() need to be called
    vp_start_button_flag = False

    def __init__(self):

        self.godirect = GoDirect(use_ble=False, use_usb=False) 

    def get_version(self):
        """ get the version of the gdx module
        """
        version = self.VERSION
        
        return version
    
    # this open() function combines the original open_ble() and open_usb() 
    def open(self, connection='usb', device_to_open=None):
        """ Open a Go Direct device via Bluetooth or USB for data collection. 
        
        Args: 
            connection (str): set as 'usb' or 'ble'

            device_to_open: Leave this argument blank to provide a list in the terminal of all 
            discovered Go Direct devices. The user then chooses the device or devices from 
            the prompt. To run code without a prompt, the argument can be set to a specific 
            Go Direct device name or names. For example,  "GDX-FOR 071000U9" or 
            "GDX-FOR 071000U9, GDX-HD 151000C1". In addition, if connection='ble', the argument 
            can be set to "proximity_pairing" to open the device with the highest 
            rssi (closest proximity).
		""" 
              
        if connection == 'ble' or connection == 'BLE':
            self.open_ble(device_to_open)
        elif connection == 'usb' or connection == 'USB':
            self.open_usb(device_to_open)
        else:
            print("Unknown value for connection in gdx.open(). Use 'usb' or 'ble'.")

    def open_usb(self, device_to_open=None):
        """ Discovers all Go Direct devices with a USB connection and opens those devices
        for data collection. 
		""" 

        # Call the godirect module to open a USB connection
        self.godirect.__init__(use_ble=False, use_usb=True)

        found_devices, number_found_devices = self.find_devices()
        
        if number_found_devices >= 1: 
            # need to have the usb device open in order to get its name
            open_usb_devices = self.open_all_usb_devices_to_get_name(found_devices)  
            if open_usb_devices >= 1:                            
                if device_to_open != None: 
                    self.select_dev_using_sn(found_devices, device_to_open)
                else:
                    # if just one device is connected, then automatically connect that device (no prompt)
                    if open_usb_devices == 1:
                        gdx.devices = found_devices
                    else:
                        self.user_chooses_device(found_devices) 
            else:
                print("USB device found but error trying to open")
                print(f"The number of USB devices found is {number_found_devices}")
                print("If this is more than connected Go Direct devices, there might")
                print("be another USB device (like a hub) that is being detected")
                print("Try moving the hub to a different port.")
                print("Otherwise, open Graphical Analysis to verify a connection") 
        else:
            str1 = "\nNo Go Direct device found \n"
            str2 = "Troubleshooting tips... \n\n"
            str3 = " - Reconnect the USB cable \n"
            str4 = " - Try a different USB port \n"
            str5 = " - Try a different USB cable \n"
            str6 = " - Open GA (Graphical Analysis) to verify a good connection \n" 
            print(str1 + str2 + str3 +str4 +str5 +str6)     

    def open_ble(self, device_to_open=None):
        """ Open a Go Direct device via bluetooth for data collection. 
        
        Args: 
            device_to_open: Leave this argument blank to provide a list in the termial of all discovered 
            Go Direct devices. The user then chooses the device or devices from the prompt. To run code 
            without a prompt, the argument can be set to a specific Go Direct device name or names. 
            For example,  "GDX-FOR 071000U9" or "GDX-FOR 071000U9, GDX-HD 151000C1". In addition, the argument
            can be set to "proximity_pairing" to open the device with the highest rssi (closest proximity).
		""" 

        if gdx.ble_open == True:
            #print("open_ble() - ble already open")
            return

        print("wait for bluetooth initialization...")
        
        # Tell godirect you want to use ble. If you need to use the Bluegiga dongle, set the 
        # use_ble_bg equal to True (uncomment the command)
        self.godirect.__init__(use_ble=True, use_ble_bg=False, use_usb=False)
        #self.godirect.__init__(use_ble=True, use_ble_bg=True, use_usb=False)
        found_devices, number_found_devices = self.find_devices()
             
        # Was there 1 or more Go Direct ble devices found? 
        # print("found " +str(number_found_devices) + " devices:")     
        if number_found_devices >= 1: 
                           
            if device_to_open == "proximity_pairing": 
                self.proximity_pairing(found_devices, number_found_devices)                 
            elif device_to_open != None: 
                self.select_dev_using_sn(found_devices, device_to_open)
            else:
                # if it is just 1 device, then connect without the popup
                if number_found_devices == 1:
                    gdx.devices = found_devices
                else:
                    self.user_chooses_device(found_devices) 

            open_success = self.open_selected_device()  
            if open_success == False:
                print("Error while trying to open device. ")
                print("Troubleshoot by opening Graphical Analysis to test")               
                      
        else:
            str1 = "No Go Direct device found \n\n"
            str2 = "Troubleshooting tips... \n"
            str3 = "Make sure device is powered on \n"
            str4 = "Confirm computer Bluetooth is on \n"
            str5 = "Open GA (Graphical Analysis) to verify a good connection \n" 
            print(str1 + str2 + str3 +str4 +str5)
    
    def find_devices(self):
        """ determine how many Go Direct devices are found (usb or ble). Returns a list 
        of GoDirectDevice objects and the number of devices.
        """
        try:
            found_devices = self.godirect.list_devices()
            number_found_devices = len(found_devices)
            #print("number of devices found = " +str(number_found_devices))
        except:
            #print("No Go Direct devices found")
            found_devices = 0
            number_found_devices = 0
            gdx.devices = []
        if number_found_devices == 0:
            gdx.devices = []
        return found_devices, number_found_devices

    def open_all_usb_devices_to_get_name(self, found_devices):
        """ Unfortunately, cannot get the name (like, 'GDX-FOR 071000U9') from
        a USB device until it is open. So, open all available USB devices.
        """

        #print("attempting to open", len(found_devices), "device(s)...")
        i = 0
        open_usb_devices = 0
        while i < len(found_devices): 
            try:
                open_device_success = found_devices[i].open()
                if open_device_success:
                    open_usb_devices += 1
                #print("open device ",i, " = ", open_device_success, sep="")
                i += 1 
            except:
                open_usb_devices = 0
                break

        return open_usb_devices
    
    def select_dev_using_sn(self, found_devices, device_to_open):
        """ The case below occurs when the device_to_open argument is given a specific device
            name or names, such as "GDX-FOR 071000U9" or "GDX-FOR 071000U9, GDX-HD 151000C1"
            In the for loop each device to open is compared to the devices found in the list of 
            found_devices. If the names match, then we store the device as a device to open.
        """

        device_name_list = []
        for device in found_devices: 
            #print("name of available device: ", str(device.name))
            device_name_list.append(str(device.name))

        device_to_open_list = device_to_open.split(", ")
        for x in device_to_open_list:
            #print("name of device wanting to open: ", x)
            for device in found_devices: 
                if x == str(device.name):
                    #print("device names match = True")
                    gdx.devices.append(device) 
                else:
                    #print("device names match = False") 
                    pass

        if len(device_to_open_list) == len(gdx.devices):
            pass
        else:
            print("serial number matching error. Check for typos in device_to_open")
            print("device_to_open = ", device_to_open_list)
            print("found devices = ", device_name_list)

    def user_chooses_device(self, found_devices):
        """ The case below occurs when there is no device_to_open argument. In this case, provide 
            a list of all discovered ble sensors and the user chooses which device or devices to open.
        """

        i=1
        print('\n')
        print("List of found devices")
        for d in found_devices:
            print(str(i)+": "+str(d))
            i += 1
        
        if len(found_devices) == 1:
            # If there is just 1 usb or ble device the user only has to hit Enter
            print('\n')
            print("One device found. Press 'enter' to connect", end=' ')
            input()
            gdx.devices.append(found_devices[0])
        else:
            print('\n')
            print("- If connecting a single device, type the number (e.g., 1) that")
            print("corresponds with the device, and then press 'enter'.")
            print("- If connnecting multiple devices, type in each number")
            print("separated with commas with no spaces(e.g., 1,2), and then ")
            print("press 'enter':", end=' ')
        
            user_selected_device = []
            for s in input().split(','):
                user_selected_device.append(int(s))
            for selected in user_selected_device: 
                gdx.devices.append(found_devices[selected-1])
            print('\n')

    def proximity_pairing(self, found_devices, number_found_devices):
        """ The case below occurs when the device_to_open parameter = "proximity_pairing"
            In the for loop each device, in the list of found_devices, is pulled out one at a time.
            That device's rssi is compared to the previous highest rssi.
            The device with the highest rssi is stored as the device to open  
        """

        print ("begin proximity pairing")
        i=1
        rmax=-99
        dmax=0
        for device in found_devices:
            print(str(i)+": "+str(device))
            v=device.rssi
            if v>rmax:
                dmax=i
                rmax=v
                #print("rmax: ", rmax," dmax: ", dmax)
            i+= 1
        x=dmax
        selected = int(x)
        if selected <= number_found_devices:
            gdx.devices.append(found_devices[selected-1])
            print("proximity device to open = ", found_devices[selected-1] )
        else:
            print("Error in proximity selection")

    def open_selected_device(self):
        """ Open the device or devices that were selected in one of the cases above.
        """
        
        open_success = False
        i = 0
        print("attempting to open", len(gdx.devices), "device(s)...")
        while i < len(gdx.devices): 
            try:
                open_device_success = gdx.devices[i].open()
                print("open device ",i, " = ", open_device_success, sep="")
                if open_device_success:
                    open_success = True
                    gdx.ble_open = True
                else:
                    open_success = False
                    return open_success
                time.sleep(1)
                i +=1     
            except:
                open_success = False
                break

        return open_success 

    def select_sensors(self, sensors=None):
        """ Select the sensors you wish to enable for data collection. 
        
        Args: 
            sensors []: if the sensors argument is left blank, a list of all available sensors is provided
            by a prompt in the terminal for the user to select from. To run code without a prompt, set this argument 
            as a 1D list or a 2D list of lists of the sensors you wish to enable, such as [1,2,3] to enable 
            sensors 1,2 and 3 for one device, or [[1,2,3],[1,2]] to enable sensors 1,2 and 3 for one device and 
            sensors 1 and 2 for a second device. 
		""" 

        # First check to make sure there are devices connected.     
        if not gdx.devices:
            print("select_sensors() - no device connected")
            return
        
        # If the sensors argument is left blank provide an input prompt for the user to select sensors
        if sensors == None: 
            i = 0
            while i < len(gdx.devices):
                selected_sensors = []
                print('\n')
                print("List of sensors for", gdx.devices[i])
                sensors = gdx.devices[i].list_sensors()
                for s in sensors:
                    c = sensors[s]
                    print(str(c))  
                
                print('\n')
                print("- If connecting a single sensor, type the number (e.g., 1) that")
                print("corresponds with the sensor, and then press 'enter'.")
                print("- If connecting multiple sensors, type in each number")
                print("separated with commas with no spaces(e.g., 1,2), and then ")
                print("click 'enter':", end=' ')
                
                for x in input().split(','):
                    selected_sensors.append(int(x))                    
                gdx.device_sensors.append(selected_sensors)
                i += 1   

        # If there is a sensor argument, it could be an integer, a list (1D), or a list of lists (2D). 
        else:
            # Checks if the variable is a list
            if type(sensors) == list:
                # It is a list. Checks if it is a 2D list
                if isinstance(sensors[0], list):
                    # Does this 2D sensor list have a list of sensors for each device? 
                    if len(sensors)!= len(gdx.devices):
                        print("the sensor parameter in select_sensors() does not match number of devices")
                        gdx.devices = []
                        return
                    else:
                        # Save the 2D list of sensors in device_sensors, such as [[1],[1,2,3]]
                        #print("2d list of sensors = ", sensors)
                        gdx.device_sensors = sensors
                # it is a 1D list
                else:
                     # A 1D list is appropriate if one device is connected. Make sure just one device is connected
                    if len(gdx.devices)!= 1:
                            print("the sensor parameter in select_sensors() does not match number of devices")
                            gdx.devices = [] 
                            return                 
                    else:
                        # Save the 1D list as a 2D list in device_sensors - [[1,2]]
                        gdx.device_sensors.append(sensors)
            # It's not a list. Check if it is an integer
            else:
                if isinstance(sensors, int):
                    # sensors are stored as a list, so change the int to a list
                    sensors = [sensors]
                    # Save the 1D list as a 2D list in device_sensors - [[1,2]]
                    gdx.device_sensors.append(sensors)

        #print("sensors for data collection = ", gdx.device_sensors)

        # check to make sure the user setup the device with a valid sensor number
        valid_sensor_num = self.check_sensor_number()
        if valid_sensor_num:
            # Enable the sensors that were selected for data collection.
            i = 0
            while i < len(gdx.devices):
                #print("device ",i, " enabled sensors = ", gdx.device_sensors[i], sep="")
                gdx.devices[i].enable_sensors(sensors = gdx.device_sensors[i])
                i +=1

            # The enabled sensor objects are stored in a variable, to be used in the read() function. 
            i = 0
            while i < len(gdx.devices):
                # The variable "enabled_sensors" is a 2D list that stores each device's enabled sensor objects [[obj],[obj,obj]].
                gdx.enabled_sensors.append(gdx.devices[i].get_enabled_sensors())
                i +=1
        # if it's not a valid number then empty the gdx.devices array so that no other functions are called
        else:
            gdx.devices = []

    def check_sensor_number(self):
        """ check to see if the user set an appropriate, available sensor number for this
        device. 
        """
            
        i = 0
        # Get the sensors from each device, one device at a time 
        while i < len(gdx.devices):
            all_sensor_numbers = []
            sensors = gdx.devices[i].list_sensors()
            
            # the all_sensor_numbers list will be used in the code below to determine incompatible sensors
            for x in sensors:
                c = sensors[x]
                number = c.sensor_number
                all_sensor_numbers.append(number)

            sensors_selected_by_user = gdx.device_sensors[i]
            for sensor_selected in sensors_selected_by_user:
                #print(f"sensor selected = {sensor_selected}, available sensors = {all_sensor_numbers}")
                if sensor_selected in all_sensor_numbers:
                    valid_sensor_num = True
                else:
                    valid_sensor_num = False
                    print('select_sensors() setup error')
                    print("The value ", sensor_selected, " in select_sensors() is not valid")
                    print(f"Valid sensor values for device{i}:", '\n')
                    for x in sensors:
                        c = sensors[x]
                        number = c.sensor_number
                        description = c.sensor_description
                        units = c.sensor_units
                        all_sensor_numbers.append(number)
                        print(f'{number} - {description} ({units})')
                    print('\n')
            i += 1

        return valid_sensor_num

    def start(self, period=None):
        """ Start collecting data from the sensors that were selected in the select_sensors() function. 
        
        Args: 
            period (int): If period is left blank, a prompt in the terminal allows the user to enter
            the period (time between samples). To run the code without this prompt, set this argument to 
            a period in milliseconds, e.g. period=1000
		"""        
        
        # First check to make sure there are devices connected.  
        if not gdx.devices:
            print("start() - no device connected")
            return 

        # if they are using vpython and the slider then the period is controlled there
        if gdx.vpython_slider:
            # if this is the very first call, then configure the slider with a starting value
            if gdx.vp_first_start == True:
                # if the period arg is left blank, just set a default starting value for the slider
                if period == None:
                    vp.slider_set(sample_rate=10)
                    period = 100
                # otherwise, use the period to set the slider with an initial value
                else:
                    sample_rate = (1/period) * 1000
                    vp.slider_set(sample_rate)
            # if it is not the first start(), then just get the period value from the slider
            else:
                period = vp.slider_get()

        # This is not a vpython program.
        else:
            # If the period argument is left blank provide an input prompt for the user to enter the period.
            if period == None: 
                print('\n')
                print("Enter the sampling period (milliseconds):", end=' ')
                period = int(input())
                print('\n')
                sample_rate = 1/(period/1000)
                #print("sample rate = ", sample_rate, "samples/second")
                
            # Provide a warning message if the user is attempting fast data collection
            if period < 10:
                input("Be aware that sampling at a period less than 10ms may be problemeatic. Press Enter to continue ")
        
        # if this is a vpython program, and this is the very first time that start() has
        # been called, do not start data collection yet. Why? Because the user's vpython 
        # code will have a start() call in their code, but they don't want to actually 'start' 
        # data collection at that point. Instead they will want to 'start' data collection 
        # when they click the 'collect' button. 
        if gdx.vpython == True and gdx.vpython_buttons == True and gdx.vp_first_start == True:
            # set the period variable in gdx_vpython.py, but do not start the devices
            gdx_vpython.ver_vpython.period = period
            gdx.vp_first_start = False
            
        # Start the devices collecting data.
        else:
            if gdx.vpython:
                # if this is a vpython program, then clear the chart, if there is a chart
                if gdx.vpython_chart:
                    column_headers= self.enabled_sensor_info()
                    vp.chart_clear(column_headers)
                gdx_vpython.ver_vpython.time = 0
                # store the period in case it is needed in vp_start_button()
                gdx.period = period
                # if this is the first call to start() change this flag 
                if gdx.vp_first_start == True:
                    gdx.vp_first_start = False

            # Start all devices
            i = 0
            while i < len(gdx.devices):
                #print("start device ", i, sep="")
                gdx.devices[i].start(period=period)
                i +=1 
        
    def read(self):             
        """ Take single point readings from the enabled sensors.

        Returns:
		    retvalues[]: a 1D list of sensor readings. A single data point 
            for each enabled sensor.
		"""
        
        retvalues = []  
        values = []       
        
        # First check to make sure there are devices connected.  
        if not gdx.devices:
            print("read() - no device connected")
            return 
        
        # Are there data in the buffer? If so, pull data from the buffer to get the retvalues
        if gdx.buffer:
            i = 0
            for i in range(len(gdx.buffer)):
                pop_values = gdx.buffer[i].pop(0)
                retvalues.append(pop_values)
            # if this was the last value in the buffer, clear the list so that it is not a list of empty lists
            if not gdx.buffer[0]:
                gdx.buffer = []     

        # The buffer is empty, so take readings from the sensor to get retvalues
        else:
            gdx.buffer = [] 
            i = 0
            # Read from each device, one at a time
            while i < len(gdx.devices):
                if gdx.devices[i].read():  
                    sensors = gdx.enabled_sensors[i]
                    # Take readings from each sensor in the device, one at time
                    if sensors:
                        for sensor in sensors: 
                            # The sensor.values call may read one sensor value, or multiple sensor values (if fast sampling)
                            values[:] = sensor.values
                            # Pull the first value off the values list
                            pop_values = values.pop(0)
                            # Build a list of each sensors' first value (this builds the return list)
                            retvalues.append(pop_values)
                            # Build a list of lists for each sensors data that is not returned and put it in the buffer
                            if values:
                                gdx.buffer.append(values)
                            sensor.clear()
                            values = []
                i +=1  
  
        if not retvalues:
            return None
        else:
            # if this is vpython and there are meters, update them with the retvalues. Note that we do
            # not need to know if the start button has been clicked (like the chart, below) because we 
            # update the meters, even when data collection is not occuring.
            if gdx.vpython: 
                if gdx.vpython_meters:
                    column_headers= self.enabled_sensor_info()
                    vp.meter_data(column_headers, retvalues)
            # if there is a chart AND the start button has been clicked
                if gdx.vpython_chart == True and gdx.vp_start_button_flag == True:
                    vp.chart_plot(retvalues)

            return retvalues
            
    def readValues(self):             
        """ Take multiple point readings from the enabled sensors and return the readings as a 2D list.

        Returns:
		    retvalues[]: a 2D list of sensor readings. Multiple points for each enabled sensor.
		"""
        
        retvalues = []
        i = 0
        # Read from each device, one at a time
        while i < len(gdx.devices):
            if gdx.devices[i].read():  
                sensors = gdx.enabled_sensors[i]
                # Take readings from each sensor in the device, one at time
                if sensors:
                    for sensor in sensors: 
                        # The sensor.values call may read one sensor value, or multiple sensor values (if fast sampling)
                        retvalues[:] = sensor.values
                        sensor.clear()
            i +=1
        return retvalues

    def listOfListsReadValues(self, dev2=False):             #Ex 11

        """ Same functionality as read() above, however value sensor.values is copied into
        values[] by value instead of by reference, allowing sensor.clear() to be called.
        Only the most recent measurements are returned from readValues() and then cleared
        from both sensor.values and values[]
        Returns:
            value[]: a list that includes a data point from each of the enabled sensors
        """
        '''if dev2 == False: #this first code sets up dev1. If dev2 = True, then do not call this code.
            if gdx.selected_device == None:
                return 
            device = gdx.selected_device

        elif dev2 == True: #is this function being called to configure dev2? then use this code
            if gdx.selected_device2 == None:
                return 
            device = gdx.selected_device2

        retValues = []  

        #if self.selected_device == None:
            #return None
        #if self.selected_device.read():
        if device.read():
            sensors = device.get_enabled_sensors()
            if sensors != None:
                for sensor in sensors: 
                    values = []
                    values[:] = sensor.values                   #New Examples
                    sensor.clear()
                    retValues.append(values)
                return retValues 
        else:
            return None'''
 
    def stop(self):
        """ Stop data collection on the enabled sensors.
		"""       

        # First check to make sure there are devices connected.  
        if not gdx.devices:
            print("stop() - no device connected")
            return
        
        i = 0
        while i < len(gdx.devices):
           # print("stop device ",i, sep="")
            gdx.devices[i].stop()
            i+=1

    def close(self):
        """ Disconnect the USB or BLE device and quit godirect.
        """

        # First check to make sure there are devices connected.  
        if not gdx.devices:
            print("close() - no device connected")
            return

        i = 0
        while i < len(gdx.devices):
            #print("close device ", i, sep="")
            gdx.devices[i].close()
            i+=1
        gdx.devices = []

        gdx.ble_open = False
        self.godirect.quit()  
        #print("quit godirect")

    def device_info(self):
        """ Returns information about the device. The device must be opened first, 
        using the open() function, before this function can be called.

		Returns:
		    device_info[]: a 1D list for one device or a 2D list for multiple. The list
            includes name, description, battery %, charger state, rssi
		"""         

        if not gdx.devices:
            print("device_info - no device connected")
            return

        # The elements in the device_info list are: 0 = name, 1 = description, 2 = battery %, 3 = charger state, 4 = rssi
        device_info = []  
        
        # If there is just one device connected, package the info in a 1D list [device info]
        if len(gdx.devices) ==1:
            device_info.append(gdx.devices[0]._name)
            device_info.append(gdx.devices[0]._description)
            device_info.append(gdx.devices[0]._battery_level_percent) 
            charger_state = ["Idle", "Charging", "Complete", "Error"]  
            device_info.append(charger_state[gdx.devices[0]._charger_state])
            device_info.append(gdx.devices[0]._rssi)
            return device_info  

        # If there is more than one device connected, package the info in a 2D list [[device0 info], [device1 info]]
        else:
            i = 0
            while i < len(gdx.devices):
                one_device_info = []
                one_device_info.append(gdx.devices[i]._name)
                one_device_info.append(gdx.devices[i]._description)
                one_device_info.append(gdx.devices[i]._battery_level_percent) 
                charger_state = ["Idle", "Charging", "Complete", "Error"]  
                one_device_info.append(charger_state[gdx.devices[i]._charger_state])
                one_device_info.append(gdx.devices[i]._rssi)
                i+=1
                device_info.append(one_device_info)
            return device_info            
                
    def enabled_sensor_info(self):
        """ Returns each enabled sensors' description and units (good for column headers).

		Returns:
		    sensor_info[]: a 1D list that includes each enabled sensors' description 
            with units, e.g. ['Force (N)', 'X-axis acceleration (m/s²)']                 
		"""                

        if not gdx.devices:
            print("enabled_sensor_info() - no device connected")
            return

        sensor_info = []

        i = 0
        # Get the enabled sensors from each device, one device at a time
        while i < len(gdx.devices):
            sensors = gdx.enabled_sensors[i] 
            for sensor in sensors:
                info = sensor.sensor_description + " (" + sensor.sensor_units + ")"
                sensor_info.append(info)
            i += 1

        return sensor_info
        # if it is just a single value, don't send it as a list
        # if len(sensor_info) == 1:
        #     return sensor_info[0]
        # else:
        #     return sensor_info

    def sensor_info(self):
        """ Information about all of the available sensors on a connected Go Direct device.

		Returns:
		    available_sensors[]: a 2D list containing information about each 
            sensor found on the device. This includes sensor number, description, units, and 
            a list of incompatible sensors (if any). An incompatible sensor is a sensor that can
            not run at the same time as this sensor. For example, Go Direct EKG cannot run the EKG
            sensor at the same time as the EMG sensor. 
		"""         

        if not gdx.devices:
            print("sensor_info() - no device connected")
            return

        available_sensors = []  
        all_sensor_numbers = []

        i = 0
        # Get the sensors from each device, one device at a time
        while i < len(gdx.devices):
            sensors = gdx.devices[i].list_sensors()
            
            # the all_sensor_numbers list will be used in the code below to determine incompatible sensors
            for x in sensors:
                c = sensors[x]
                number = c.sensor_number
                all_sensor_numbers.append(number)
        
            for x in sensors:
                incompatible_sensors = []
                s = sensors[x]
                number = s.sensor_number
                description = s.sensor_description
                units = s.sensor_units

                # The exclusion_mask is a number that represents sensor numbers that are incompatible with this sensor.
                exclusion_mask = s._mutual_exclusion_mask
                # Convert the exclusion_mask number to a list of Trues and Falses representing the mask.
                bin_string = format(exclusion_mask, '32b')  
                # Reverse the bin_string (with [::-1]) to format it with the most significant bit first.
                # The answer is a True False list [TRUE, TRUE, FALSE]
                answer =  [x == '1' for x in bin_string[::-1]] 
                
                e = 0
                # Change the True/False list to a list of sensor numbers. e.g, [TRUE, TRUE, FALSE] = [1,2]
                # Pull out the True/False values of the list one at a time and if it is TRUE, then add it to 
                # the list of incompatible sensors. 
                for channel in answer: 
                    # If this value of the list is TRUE and it is a confirmed sensor number. 
                    if channel == True and e in all_sensor_numbers: 
                        incompatible_sensors.append(e)
                    e+=1  

                available_sensors.append([number, description, units, incompatible_sensors])
                
            i+=1
        # Return the available_sensor list [0 = sensor number, 1 = description, 2 = units, 3 = incompatible sensors[]]
        return available_sensors

    def discover_ble_devices(self, init=True):
        """ Enables bluetooth, and returns the name and rssi of all discovered GoDirect devices. 
        This function should be called prior to opening a device. The name returned 
        by this function can be used as an argurment in the ble_open() function to open a specific device. 

		Returns:
		    discovered_ble_devices[]: a 2D list. A list containing a list of name and rssi for each device
            [[name1,rssi1],[name2,rssi2],[name3,rssi3]]
		""" 
        
        # If you are going to call this several times, there might be a reason to only call 
        # the init code once.
        # The first time you call this function set init = True, the following times set init = False.
        if init == True:
            self.godirect.__init__(use_ble=True, use_usb=False)
            gdx.ble_open = False
            print("Begin search for ble devices...")
 
        # Find all available bluetooth devices 
        found_devices = self.godirect.list_devices() 
        number_found_devices = len(found_devices)
        #print("Number of ble devices found = " +str(number_found_devices))
        discovered_ble_devices = []
                
        if number_found_devices >= 1:
            for device in found_devices:
                device_name = device.name
                # Note that you can get the rssi from this call before opening the device
                device_rssi = device.rssi
                discovered_ble_devices.append([device_name, device_rssi])
            
        return discovered_ble_devices

#### VPYTHON FUNCTIONS ####

    def vp_vernier_canvas(self, buttons=True, slider=True, meters=True, chart=False, cvs=True):
        """ Create vptyhon objects that are used for controlling data collection. 
        
        Args: 
            buttons (bool): Create a Collect/Stop and Close button
            slider (bool): Create a slider to control sampling rate
            meters (bool): Create meters to display live sensor data
            chart (bool): Create a chart to plot live sensor data
            cvs (bool): Create a default canvas, ready for vpython objects
        
        """
        # keep track of what objects have been selected
        gdx.vpython = True
        gdx.vpython_buttons = buttons
        gdx.vpython_chart = chart
        gdx.vpython_meters = meters
        gdx.vpython_slider = slider

        # setup the canvas based on what was selected
        if buttons or slider:
            vp.setup_canvas(buttons, slider)
        if chart:
            column_headers= self.enabled_sensor_info()
            vp.chart_init(column_headers)
        if meters:
            vp.meter_init()
        if cvs:
            vp.create_default_canvas()

    def vp_close_is_pressed(self):
        """ Monitor the state of the vpython canvas Close button. When true, 
        a gdx.stop() and gdx.close() are called to stop data collection and 
        disconnect the device. When false, and if there are meters, they are 
        updated with live readings.

        Returns:
            close_button_state (bool): True if Close button has been pressed
        """
        # Note that there is no vpython rate() call here, but there is in the
        # gdx_vpython.py module in the collect_button() function

        # First check to make sure there are devices connected.      
        if not gdx.devices:
            print("vp_close_button() - no device connected")
            close_button_state = True
        else:
            # get the state of the vpython Close button
            close_button_state = vp.closed_button()
        
        # If the user has clicked the Close button
        if close_button_state == True:
            self.stop()
            self.close()
            if gdx.vpython_chart:
                vp.chart_delete()
            if gdx.vpython_meters:
                vp.meter_delete()
            if gdx.vpython_slider:
                vp.slider_delete()
            if gdx.vpython_buttons:
                vp.button_delete()
            vp.canvas_delete()
        # the Close button has not been pressed
        else:
            # if there are meters, update their values using a short data collection loop
            if gdx.vpython_meters:
                for device in gdx.devices:
                    device.start(period=250)
                for x in range(4):
                    # The read() function has code to send the value to the meter
                    self.read()
                self.stop()
                
        return close_button_state

    def vp_collect_is_pressed(self):
        """ Monitor the state of the vpython canvas Collect/Stop button. When Collect
        is clicked, a gdx.start() is called. When Stop is clicked, a gdx.stop() is
        called.

        Returns:
            collect_button_state (bool): True if button is in the 'COLLECT' state. False
            if the button is in the 'STOP' state.
        """

        # First check to make sure there are devices connected.      
        if not gdx.devices:
            print("vp_collect_button() - no device connected")
            return
        
        # get the state of the collect button
        collect_button_state = vp.collect_button()

        # It is not enough to know the state of the button, it is also 
        # important to know if the button was just clicked (just been pressed)
        if gdx.vp_start_button_flag != collect_button_state:
            # if it was just clicked into the True state, start data collection
            if collect_button_state == True:
                self.start(gdx.period)
                gdx.vp_start_button_flag = True
            # if it was just clicked into the False state, stop data collection
            else:
                self.stop()
                gdx.vp_start_button_flag = False

        return collect_button_state

    def vp_get_slider_period(self):
        """ Get the value of the slider as the period (time between samples).
            Returns the value in milliseconds.
        """
        period = vp.slider_get()
        return period
```
