## Supplementary material for "Evidence for the extinction burst in grip force: A preregistered study": Electronic Supplementary Materials: gdx_vpython.py

```
import logging
  
# from gdx_modules.gdx import gdx_class
# gdx = gdx_class()
#import gdx_modules.gdx


#Are these variable required outside the class? 
# woutput = wtext(text='')
# collectbutton = None
# closebutton = None

class ver_vpython:


    closed = False
    collect_button_state = False
    period = 100   # period in ms
    time = 0
    plot_1 = None    # vpython gcurve for the graph canvas
    plot_2 = None
    plot_3 = None
    plot_4 = None
    plot_5 = None
    graph_canvas = None
    meter_canvas = None
    button_canvas = None
    meter_text = None
    slider_text = None
    cb = None    # collect button
    clsb = None    # close button
    sl = None    # slider

    def __init__(self):  
        """
        """

    def setup_canvas(self, vp_button=False, slider_control=False):

        # Can we install the vpython library, with godirect?
        #from vpython import canvas, button, box, wtext, checkbox, rate
        from vpython import button, scene, slider, wtext, canvas, color
        #global collectbutton, closebutton
        # if they are using vpython, then create the canvas and collect/stop/close buttons
        
    
        ver_button_scene = canvas()
        ver_vpython.button_canvas = ver_button_scene
        ver_button_scene.width = 0
        ver_button_scene.height = 10

        ver_button_scene.append_to_title('\n')
        if vp_button:
            collectbutton = button(text='<b style="color:green; font-size:26px"> COLLECT </b>', 
                                pos=ver_button_scene.title_anchor, bind=vp_collect_stop)
            ver_vpython.cb = collectbutton
            ver_button_scene.append_to_title('  ')
            closebutton = button(text='<b style="color:red; font-size:26px">   CLOSE   </b>', 
                                pos=ver_button_scene.title_anchor, bind=vp_closed)
            ver_vpython.clsb = closebutton
        
        if slider_control:
            ver_button_scene.append_to_title('  ')
            slider_control = slider(pos=ver_button_scene.title_anchor, min=1, max=100, value=10, step=1, length=200, bind=vp_slider)
            ver_vpython.sl = slider_control
            #scene.append_to_caption('\n\n') 
            #scene.append_to_title('\n')
            slider_text = wtext(pos=ver_button_scene.title_anchor, text='10 samples/second')
            ver_vpython.period = 100
            ver_vpython.slider_text = slider_text


        ver_button_scene.append_to_title('\n')

    def create_default_canvas(self):
        """ Add a small canvas below the button canvas and the meter canvas. This should allow
        the user to create vpython objects without having to create a scene. If they do create
        a scene it should simply overwrite this canvas.        
        """

        from vpython import canvas, scene, color

        scene = canvas(width=800, height=150)
        scene.background = color.black
    
    def button_delete(self):
        ver_vpython.cb.delete()
        ver_vpython.clsb.delete()

    def slider_delete(self):
        from vpython import canvas
        
        canvas.delete(ver_vpython.slider_text)
        ver_vpython.sl.delete()

    def canvas_delete(self):
        from vpython import canvas, scene

        # this also seems to work for deleting the button object
        #canvas.delete(ver_vpython.cb)
        # ver_vpython.cb.delete()
        # ver_vpython.clsb.delete()
        ver_vpython.button_canvas.delete()
        
        scene.delete()
        current = canvas.get_selected()
        if current:
            current.delete()

    def slider_set(self, sample_rate):
        ver_vpython.sl.value = sample_rate
        ver_vpython.period = (1/sample_rate) * 1000
        ver_vpython.slider_text.text = f'{sample_rate} samples/second'

    def slider_get(self):
        period = ver_vpython.period
        return period
    
    def chart_init(self, column_headers):
        from vpython import graph, gcurve, color, vector
        
        if column_headers == None:
            column_headers = 'Data'
        gd = graph(xtitle='Time', ytitle=column_headers, scroll=True,
        width=500, height=300, xmin=0, xmax=5, fast=False)
        ver_vpython.graph_canvas = gd
        plot_1 = gcurve(color=vector(0.37, 0.57, 0.74))
        ver_vpython.plot_1 = plot_1
        ver_vpython.plot_1.plot(0,0)
        plot_2 = gcurve(color=vector(0.75, 0.75, 0.3))
        ver_vpython.plot_2 = plot_2
        ver_vpython.plot_2.plot(0,0)
        plot_3 = gcurve(color=vector(0, 0, 0.57))
        ver_vpython.plot_3 = plot_3
        ver_vpython.plot_3.plot(0,0)
        plot_4 = gcurve(color=vector(0.8, 0.38, 0.44))
        ver_vpython.plot_4 = plot_4
        ver_vpython.plot_4.plot(0,0)
        plot_5 = gcurve(color=vector(0.35, 0.2, 0.49))
        ver_vpython.plot_5 = plot_5
        ver_vpython.plot_5.plot(0,0)

    def chart_plot(self, data):
        if data == None:
            return
        else:
            if not isinstance(data, list):
                # measurements needs to be a list, if it is not, change it to a list
                data = [data]
            len_data = len(data)
            if len_data == 1:
                ver_vpython.plot_1.plot(ver_vpython.time, data[0])
            elif len_data == 2:
                ver_vpython.plot_1.plot(ver_vpython.time, data[0])
                ver_vpython.plot_2.plot(ver_vpython.time, data[1])
            elif len_data == 3:
                ver_vpython.plot_1.plot(ver_vpython.time, data[0])
                ver_vpython.plot_2.plot(ver_vpython.time, data[1])
                ver_vpython.plot_3.plot(ver_vpython.time, data[2])
            elif len_data == 4:
                ver_vpython.plot_1.plot(ver_vpython.time, data[0])
                ver_vpython.plot_2.plot(ver_vpython.time, data[1])
                ver_vpython.plot_3.plot(ver_vpython.time, data[2])
                ver_vpython.plot_4.plot(ver_vpython.time, data[3])
            else:
                ver_vpython.plot_1.plot(ver_vpython.time, data[0])
                ver_vpython.plot_2.plot(ver_vpython.time, data[1])
                ver_vpython.plot_3.plot(ver_vpython.time, data[2])
                ver_vpython.plot_4.plot(ver_vpython.time, data[3])
                ver_vpython.plot_5.plot(ver_vpython.time, data[4])

        ver_vpython.time = ver_vpython.time + (ver_vpython.period/1000)
        

    def chart_clear(self, column_headers):
        if column_headers == None:
            column_headers = 'Data'
        ver_vpython.plot_1.delete()
        ver_vpython.plot_2.delete()
        ver_vpython.plot_3.delete()
        ver_vpython.plot_4.delete()
        ver_vpython.plot_5.delete()
        ver_vpython.graph_canvas.ytitle = column_headers
        ver_vpython.graph_canvas.xmin = 0
        ver_vpython.graph_canvas.xmax = 5
        
    def chart_delete(self):
        ver_vpython.graph_canvas.delete()

    def meter_init(self):
        from vpython import canvas, wtext, scene, color
        
        ver_meter_canvas = canvas(width=0, height=20)
        ver_vpython.meter_canvas = ver_meter_canvas
        ver_meter_canvas.append_to_title('\n')
        woutput = wtext(text='', pos=ver_meter_canvas.title_anchor)
        ver_vpython.meter_text = woutput
        #<b>mass <i>M</i></b>
        #woutput.text = "<b>{ch_string}</b>\n".format(ch_string)
        
        woutput.text = f""  

    def meter_data(self, column_headers, data):
        if data == None:
            meter_string = 'No data'
        else:
            if not isinstance(data, list):
                # data needs to be a list, if it is not, change it to a list
                data = [data]
            if not isinstance(column_headers, list):
                # column_headers needs to be a list, if it is not, change it to a list
                column_headers = [column_headers]

            meter_string = ' '
            for (ch, d) in zip(column_headers, data):
                round_data = str(round(d, 2))
                meter_string = meter_string + ch + ": " + round_data + '    '
        
        ver_vpython.meter_text.text = f"{meter_string}"

    def meter_delete(self):
        from vpython import canvas
        
        canvas.delete(ver_vpython.meter_text)
        ver_meter_canvas = ver_vpython.meter_canvas
        # this worked
        ver_meter_canvas.delete()
        # this worked too
        #canvas.delete(ver_meter_canvas)
    
    # def slider_init(self):
    #     from vpython import canvas, slider, wtext, scene
        
    #     sc = canvas(width=0, height=20)
    #     ver_vpython.slider_canvas = sc
    #     # when the slider is changed, the function vp_slider is called
    #     sl = slider(align='right', min=1, max=100, value=10, step=1, length=200, bind=vp_slider, left=20)
    #     #scene.append_to_caption('\n\n') 
    #     #scene.append_to_title('\n')
    #     slider_text = wtext(text='')
    #     ver_vpython.slider_text = slider_text

    # def print_to_canvas(self):
    #     """ Feedback to the user on the vpython screen 
	# 	"""  
    #     from vpython import canvas

    #     canvas.get_selected().append_to_caption('Must specify device sensors.')
    #     canvas.get_selected().caption = 'test caption'
    #     raise AttributeError('Must specify device sensors.')     

    def collect_button(self):
        """ Return value = True if the button is in the Collect state. Return
        value = False if it is in the Stop state.
        """
        from vpython import rate, color

        if ver_vpython.collect_button_state:
            return True
            
        else:
            # assumption is that rate() is only needed when data collection is not occurring
            # When it is, the computer will automatically be slowed because of the time
            # required to talk to the hardware.
            rate(50)
            return False

    def closed_button(self):
        if ver_vpython.closed:
            return True
        else:
            return False

    

    
def vp_collect_stop(f):
    """ This function gets called only when the button has been pressed. Return
    value = True if the button is in the Collect state.
    """
    
    if f.text == '<b style="color:green; font-size:26px"> COLLECT </b>':
        f.text = '<b style="color:black; font-size:26px">    STOP     </b>'
        #
        # "Change the class variable’s value using the class name only."
        ver_vpython.collect_button_state = True

    else:
        f.text = '<b style="color:green; font-size:26px"> COLLECT </b>'
        ver_vpython.collect_button_state = False

def vp_closed():

    ver_vpython.closed = True
    ver_vpython.collect_button_state = False

def vp_slider(s):
    ver_vpython.period = (1/s.value) * 1000
    ver_vpython.slider_text.text = f'{s.value} samples/second'

  
    


   
```
