## Supplementary material for "Evidence for the extinction burst in grip force: A preregistered study": Electronic Supplementary Materials: ReadMe — PsychoPy task code.txt

README — PsychoPy task code

Files included
EBExperiment_force.py: Python script generated by PsychoPy implementing the experimental task used in the study.
gdx/: Package required to interface the hand dynamometer with PsychoPy.
assign_cfg.json: Configuration file containing the randomization seed used for participant assignment to experimental groups.
Stimulus files. Individual image files used by the task, including: informed consent stimulus, monetary reinforcer image (5 Mexican pesos), instructional images for correct hand dynamometer use.

Description
This PsychoPy task was designed to study extinction-burst-like changes in force output in human participants, measured via a hand dynamometer.
The task includes an operant reinforcement phase followed by extinction, as well as automatic assignment to four experimental groups, with group-specific parameters defined within the script.

Software requirements
PsychoPy (Python environment provided by PsychoPy).
The gdx package included in this folder (required when running the task with the dynamometer).

Hardware
Hand dynamometer connected via USB (as specified in the script).
The task includes a calibration routine used to determine force thresholds during the experiment.

Output
When executed, the script saves task data to a local data/ directory created by PsychoPy.
No participant data are included in this shared material.
