## Supplementary material for "Evidence for the extinction burst in grip force: A preregistered study": Electronic Supplementary Materials: 0_ebf_power_analysis.R

```
#Sample size calculation via statistical power analysis

#Load required packages
if (!require('dplyr'))      install.packages('dplyr');      library(dplyr)
if (!require('ggplot2'))    install.packages('ggplot2');    library(ggplot2)
if (!require('lme4'))       install.packages('lme4');       library(lme4)
if (!require('lmerTest'))   install.packages('lmerTest');   library(lmerTest)
if (!require('effectsize')) install.packages('effectsize'); library(effectsize)

set.seed(1401)

#Set target parameters for the simulation
alpha            <- 0.05 #significance threshold
nsim             <- 10000        #number of simulated data sets
n_per_group      <- 16        #number of participants per group
#NOTE: If you decrease n_per_group from 16 to 15 (i.e., 60 total across 4 groups),
#...the simulated power remains > 0.80 under the preregistered assumptions, indicating that a modest
#...shortfall in analyzable N does not materially compromise the planned confirmatory test.


beta_int_std_tgt <- 0.55 #target standardized β for stage×treatment interaction
beta_block_std   <- -0.30      #target standardized β for block
ICC_target       <- 0.30 #target intra-class correlation  for ID random intercept

beta_stage_std   <-  0.00 #before/after main effect set to 0 by 
#assumption: after conditioning on numeric block (secular trend), we assume no 
#...additional global shift from before→after unless treatment is present
beta_trt_std     <-  0.00 #cft/treat main effect set to 0 by 
#assumption: no global treat–cft difference across stages; the effect of 
#treatment is expressed only when it co-occurs with ‘after’ (the interaction).

####HELPER FUNCTIONS

#Bootstraped confidence intervals
boot_ci_mean <- function(x, R = 2000L) {
  if (length(x) <= 1L) return(rep(mean(x), 3L))
  b <- replicate(R, mean(sample(x, replace = TRUE)))
  as.numeric(quantile(b, c(.025, .5, .975), names = FALSE))
}

#Viz 1: interaction plot on pooled data
plot_interaction_last <- function(d_last, offset = 0.06) {
  pal <- c(cft = "#4E79A7", treat = "#E15759") 
  
  d_last <- d_last %>%
    mutate(
      stage     = factor(stage, levels = c("before","after")),
      treatment = factor(treatment, levels = c("cft","treat")),
      x_base    = as.numeric(stage),
      x_off     = ifelse(treatment == "treat", +offset, -offset),
      x         = x_base + x_off
    )
  
  id_itxn <- d_last %>%
    group_by(ID, treatment, stage, x) %>%
    summarise(mean_y = mean(force_obs), .groups = "drop")
  
  sum_itxn <- d_last %>%
    group_by(stage, treatment) %>%
    summarise(
      n   = n(),
      ci  = list(boot_ci_mean(force_obs, R = 2000L)),
      mean = ci[[1]][2], lo = ci[[1]][1], hi = ci[[1]][3],
      .groups = "drop"
    ) %>%
    mutate(
      x_base = as.numeric(stage),
      x_off  = ifelse(treatment == "treat", +offset, -offset),
      x      = x_base + x_off
    )
  
  ggplot() +
    geom_line(
      data = id_itxn,
      aes(x = x, y = mean_y, group = interaction(ID, treatment), color = treatment),
      alpha = 0.22, linewidth = 1.5
    ) +
    geom_point(
      data = id_itxn,
      aes(x = x, y = mean_y, color = treatment),
      alpha = 0.35, size = 2
    ) +
    geom_line(
      data = sum_itxn,
      aes(x = x, y = mean, group = treatment, color = treatment),
      linewidth = 2
    ) +
    geom_point(
      data = sum_itxn,
      aes(x = x, y = mean, color = treatment),
      size = 3
    ) +
    geom_errorbar(
      data = sum_itxn,
      aes(x = x, ymin = lo, ymax = hi, color = treatment),
      width = 0.06, linewidth = 0.6
    ) +
    scale_x_continuous(breaks = c(1, 2), labels = c("before", "after")) +
    scale_color_manual(values = pal) +
    labs(
      x = "Stage", y = "Observed outcome", color = "Treatment",
      title = "stage × treatment with individual trajectories and mean ± 95% CI"
    ) +
    theme_minimal(base_size = 12)
}

#Viz 2: mean data faceted by group
plot_faceted_last <- function(d_last) {
  pal <- c(cft = "#4E79A7", treat = "#E15759")
  lw  <- 1.0  
  
  agg_pts <- d_last %>%
    mutate(
      Group     = factor(Group, levels = c("E8","E16","E24","NE")),
      Block     = as.integer(Block),
      stage     = factor(stage, levels = c("before","after")),
      treatment = factor(treatment, levels = c("cft","treat"))
    ) %>%
    arrange(Group, Block, treatment, stage) %>%
    group_by(Group, Block, treatment, stage) %>%
    summarise(
      mean_y = mean(force_obs),
      se_y   = sd(force_obs)/sqrt(n()),
      .groups = "drop"
    )
  
  agg_line <- agg_pts %>%
    group_by(Group, Block, treatment) %>%
    summarise(mean_y = mean(mean_y), .groups = "drop")
  
  seg <- agg_line %>%
    group_by(Group, treatment) %>%
    arrange(Block, .by_group = TRUE) %>%
    mutate(Block_next = lead(Block),
           mean_next  = lead(mean_y)) %>%
    ungroup() %>%
    filter(!is.na(Block_next) & (Block_next - Block == 1))
  
  ggplot() +
    geom_segment(
      data = seg,
      aes(x = Block, y = mean_y, xend = Block_next, yend = mean_next, color = treatment),
      linewidth = lw
    ) +
    geom_point(
      data = agg_pts,
      aes(x = Block, y = mean_y, color = treatment, shape = stage),
      size = 2
    ) +
    geom_errorbar(
      data = agg_pts,
      aes(x = Block, ymin = mean_y - se_y, ymax = mean_y + se_y, color = treatment),
      width = 0.1, linewidth = lw
    ) +
    facet_wrap(~ Group, nrow = 1, drop = FALSE) +
    scale_color_manual(values = pal) +
    scale_x_continuous(breaks = sort(unique(agg_pts$Block))) +
    labs(
      x = "Block (numeric)", y = "Mean observed outcome",
      title = "Means ± SE by Group × Block (faceted); adjacent blocks connected per treatment",
      color = "Treatment", shape = "Stage"
    ) +
    theme_bw(base_size = 12) +
    theme(panel.grid.minor = element_blank())
}

#Study design (group, number of blocks, types of block sequence)
#E8: 2 blocks (B1=reward, B2=extinction)
#E16: 4 blocks (B1-3=reward, B4=extinction)
#E24: 6 blocks (B1-5=reward, B6=extinction)
#NE: 6 blocks (B1-6=reward)

#Set the number of before/after pairs per group
k_by_group <- c(E8 = 1L, E16 = 2L, E24 = 3L, NE = 3L)

#Create simulated participants and assign to groups
build_ids <- function(n_pg) {
  mk_ids <- function(prefix, n) paste0(prefix, "_", seq_len(n))
  tibble(
    ID    = c(mk_ids("E8", n_pg), mk_ids("E16", n_pg), mk_ids("E24", n_pg), mk_ids("NE", n_pg)),
    Group = c(rep("E8", n_pg), rep("E16", n_pg), rep("E24", n_pg), rep("NE", n_pg))
  )
}

#Create one simulated data set
simulate_one <- function() {
  ids <- build_ids(n_per_group)
  
  expand_subject <- function(id, grp) {
    K <- k_by_group[[grp]] #number of BEFORE/AFTER pairs: E8=1, E16=2, E24=3, NE=3
    out <- vector("list", K)
    for (b in seq_len(K)) {
      trt <- if (grp == "NE") "cft" else if (b == K) "treat" else "cft"
      out[[b]] <- data.frame(
        ID = id, Group = grp,
        Block = c(2*b - 1, 2*b),          #before = odd, after = even
        stage = c("before","after"),
        treatment = trt,
        stringsAsFactors = FALSE
      )
    }
    bind_rows(out)
  }
  
  d <- bind_rows(lapply(seq_len(nrow(ids)), function(i) {
    expand_subject(ids$ID[i], ids$Group[i])
  }))
  
  
  d <- d %>%
    mutate(
      stage     = factor(stage, levels = c("before","after")),
      treatment = factor(treatment, levels = c("cft","treat")),
      Block     = as.integer(Block)
    )
  
  d <- d %>%
    mutate(
      block_std = as.numeric(scale(Block)),
      is_after  = as.integer(stage == "after"),
      is_treat  = as.integer(treatment == "treat"),
      after_xtreat = is_after * is_treat
    )
  
  #random-effect & residual SD used to hit ICC target approximately
  #ICC ≈ var_id / (var_id + var_eps).
  sd_id  <- sqrt(ICC_target)
  sd_eps <- sqrt(1 - ICC_target)
  
  #Add random intercepts
  re_id <- rnorm(n = n_distinct(d$ID), mean = 0, sd = sd_id)
  names(re_id) <- unique(d$ID)
  
  #Add fixed effects
  fixed_z <- beta_block_std * d$block_std +
    beta_stage_std * d$is_after    +
    beta_trt_std   * d$is_treat    +
    beta_int_std_tgt * d$after_xtreat
  
  #Generate outcome
  d$force_z  <- fixed_z + re_id[d$ID] + rnorm(nrow(d), 0, sd_eps)
  d$force_obs <- d$force_z 
  
  #Fit  model for hypothesis testing
  m <- lmer(force_obs ~ stage * treatment + Block + (1 | ID), data = d)
  
  #get interaction stats 
  coefs <- summary(m)$coefficients
  p_int <- unname(coefs["stageafter:treatmenttreat", "Pr(>|t|)"])
  b_raw <- unname(fixef(m)["stageafter:treatmenttreat"])
  
  #get standardized regression coefficients
  std_tab <- suppressWarnings(try(standardize_parameters(m, method = "refit"), silent = TRUE))
  b_int_std <- NA_real_
  b_blk_std <- NA_real_
  if (!inherits(std_tab, "try-error")) {
    std_col <- grep("^Std", names(std_tab), value = TRUE)[1]
    if (!is.na(std_col)) {
      #interaction effect size
      row_int <- std_tab$Parameter == "stageafter:treatmenttreat"
      if (any(row_int)) b_int_std <- unname(std_tab[[std_col]][row_int])
      #block effect size
      row_blk <- std_tab$Parameter == "Block"
      if (any(row_blk)) b_blk_std <- unname(std_tab[[std_col]][row_blk])
    }
  }
  
  #intra class correlation of participant ID
  vc      <- as.data.frame(VarCorr(m))
  var_id  <- vc$vcov[vc$grp == "ID"]
  var_res <- sigma(m)^2
  icc_id  <- var_id / (var_id + var_res)
  
  list(
    d = d, m = m,
    p = p_int, b_raw = b_raw, b_int_std = b_int_std, b_blk_std = b_blk_std,
    icc_id = icc_id
  )
}

#Power analysis loop (collect distributions); keep last run for viz
power_sim <- function(nsim) {
  pb <- txtProgressBar(min = 0, max = nsim, style = 3)
  pvals <- betas_raw <- b_int_std <- b_blk_std <- icc_ids <- rep(NA_real_, nsim)
  last_d <- NULL; last_m <- NULL
  
  for (i in seq_len(nsim)) {
    res <- simulate_one()
    pvals[i]     <- res$p
    betas_raw[i] <- res$b_raw
    b_int_std[i] <- res$b_int_std
    b_blk_std[i] <- res$b_blk_std
    icc_ids[i]   <- res$icc_id
    if (i == nsim) { last_d <- res$d; last_m <- res$m }
    if (i %% max(1, floor(nsim/50)) == 0) setTxtProgressBar(pb, i)
  }
  close(pb)
  
  list(
    #one sided testing
    power = mean((pvals/2 < alpha) & (betas_raw > 0), na.rm = TRUE),
    pvals   = pvals,
    b_raw   = betas_raw,
    b_int_std = b_int_std,
    b_blk_std = b_blk_std,
    icc_id  = icc_ids,
    last_d  = last_d,
    last_m  = last_m
  )
}

#Run simulation with specified parameters
cat("Running power simulation...\n")
note <- paste0(
  "Even with a fixed random seed and the computational environment recorded (e.g., via sessionInfo()), ",
  "small run-to-run differences are expected and do not change the qualitative conclusions. ",
  "Accordingly, exact numeric outputs (such as power estimates or median standardized coefficients) ",
  "may differ slightly from the preregistered values, typically at the third decimal place. ",
  "In addition, some environments (different R/package versions or systems) may occasionally produce ",
  "minor mixed-model convergence warnings where the optimizer stops just above the tolerance; ",
  "these are typically benign and should not be interpreted as evidence that the results have changed."
)
cat(note, "\n")

# WARNING: This step may take a while to run.
sim_out <- power_sim(nsim)

q <- function(x, p) quantile(x, p, na.rm = TRUE)

cat("\n================ Simulation results ================\n")
cat(sprintf("Targets:  β_int(std)=%.3f  β_block(std)=%.3f   ICC=%.3f\n",
            beta_int_std_tgt, beta_block_std, ICC_target))
cat(sprintf("n_per_group: %d   nsim: %d   alpha: %.3f\n", n_per_group, nsim, alpha))
cat("----------------------------------------------------\n")
cat(sprintf("POWER (stage×treatment, lmerTest): %.3f\n", sim_out$power))
cat(sprintf("Median standardized β (interaction): %.3f  [IQR %.3f, %.3f]\n",
            median(sim_out$b_int_std, na.rm = TRUE), q(sim_out$b_int_std,.25), q(sim_out$b_int_std,.75)))
cat(sprintf("Median standardized β (Block):       %.3f  [IQR %.3f, %.3f]\n",
            median(sim_out$b_blk_std, na.rm = TRUE), q(sim_out$b_blk_std,.25), q(sim_out$b_blk_std,.75)))
cat(sprintf("Median ICC(ID):                      %.3f  [IQR %.3f, %.3f]\n",
            median(sim_out$icc_id,    na.rm = TRUE), q(sim_out$icc_id,.25),  q(sim_out$icc_id,.75)))
cat("====================================================\n")

#Last run checks
last_d <- sim_out$last_d
last_m <- sim_out$last_m

cat("\nLast run — head(last_d):\n"); print(head(last_d, 12))
cat("\nLast run — counts by Group × Block × treatment:\n")
print(last_d %>% count(Group, Block, treatment))
cat("\nLast run — stage × treatment cross-tab:\n")
print(with(last_d, table(stage, treatment)))

#Last run visualizations
#interaction plot
plot_interaction_last(last_d, offset = 0.06)

#faceted Group × Block plot
plot_faceted_last(last_d)

##last-run model summaries
print(summary(last_m))
standardize_parameters(last_m)
```
