## Supplementary material for "Evidence for the extinction burst in grip force: A preregistered study": Electronic Supplementary Materials: 1_ebf_compiling_inspection_segmentation.R

```
#Compile raw files into the source data frame and create helpers to 
#...inspect full sessions or selected within-session segments

#Load required packages
if (!require('readxl')) install.packages('readxl'); library(readxl)
if (!require('dplyr')) install.packages('dplyr'); library(dplyr)
if (!"tools" %in% (.packages())) library(tools)
if (!require('ggplot2')) install.packages('ggplot2'); library(ggplot2)

#IMPORTANT: set ebf_raw_data as working directory

#helper to normalize file names
keyify <- function(x) {
  x <- basename(x)
  x <- tools::file_path_sans_ext(x)
  tolower(trimws(x))
}

#Load thresholds.xlsx (col1 = filename, col2 = reward threshold)
th <- read_excel("thresholds.xlsx", col_names = FALSE)
colnames(th) <- c("file_name", "reward_th")
th$file_key <- keyify(th$file_name)

#Named vector: reward threshold by normalized key
reward_by_key <- setNames(th$reward_th, th$file_key)

#List participant CSVs
csv_files <- list.files(pattern = "\\.csv$", full.names = TRUE)

#Helper to return source data frame
compile_one <- function(fpath) {
  df <- tryCatch(read.csv(fpath, stringsAsFactors = FALSE), error = function(e) NULL)
  if (is.null(df)) return(NULL)
  
  df$block_original <- if ("block" %in% names(df)) df$block else NA
  
  fkey <- keyify(fpath)
  if (!fkey %in% names(reward_by_key)) {
    warning(sprintf("No reward threshold found for %s; force_std will be NA.", basename(fpath)))
    calib <- NA_real_
  } else {
    reward_th <- as.numeric(reward_by_key[[fkey]])
    calib <- reward_th / 0.7
  }
  df$force_std <- if (is.finite(calib)) df$force_N / calib else NA_real_
  df$calibration_force <- calib
  
  if (!"trial_idx" %in% names(df)) stop("Column 'trial_idx' is missing.")
  block_from_trials <- floor((as.integer(df$trial_idx) - 1L) / 4L) + 1L
  
  blk_raw <- df$block_original
  replace_mask <- is.na(blk_raw) | blk_raw == "" | tolower(trimws(blk_raw)) == "block 1"
  
  existing_numeric <- suppressWarnings(as.integer(gsub("[^0-9]", "", blk_raw)))
  df$block_final <- existing_numeric
  df$block_final[replace_mask] <- block_from_trials[replace_mask]
  if (all(is.na(df$block_final))) df$block_final <- block_from_trials
  
  df$stage <- NA_character_
  df$stage[df$block_final %in% c(2L, 4L, 6L)] <- "before"
  df$stage[df$block_final %in% c(3L, 5L, 7L)] <- "after"
  
  max_trials <- suppressWarnings(max(as.integer(df$trial_idx), na.rm = TRUE))
  grp <- if (is.finite(max_trials)) {
    if (max_trials == 28L) "NE"
    else if (max_trials == 24L) "E24"
    else if (max_trials == 16L) "E16"
    else if (max_trials == 8L)  "E8"
    else NA_character_
  } else NA_character_
  df$group <- grp
  
  df$condition <- NA_character_
  
  ext_flag <- rep(FALSE, nrow(df))
  if ("block_original" %in% names(df)) {
    ext_flag <- !is.na(df$block_original) & tolower(trimws(df$block_original)) == "extinction_conf"
  }
  
  blk <- suppressWarnings(as.integer(df$block_final))
  
  if (!is.na(grp) && grp == "NE") {
    df$condition[!is.na(blk) & blk %in% 2:7] <- "ctfl"
    
  } else if (!is.na(grp) && grp == "E8") {
    df$condition[!is.na(blk) & blk == 2L] <- "treat"
    df$condition[ext_flag] <- "treat"
    
  } else if (!is.na(grp) && grp == "E16") {
    df$condition[!is.na(blk) & blk %in% 2:3] <- "ctfl"
    df$condition[!is.na(blk) & blk == 4L] <- "treat"
    df$condition[ext_flag] <- "treat"
    
  } else if (!is.na(grp) && grp == "E24") {
    df$condition[!is.na(blk) & blk %in% 2:5] <- "ctfl"
    df$condition[!is.na(blk) & blk == 6L] <- "treat"
    df$condition[ext_flag] <- "treat"
  }
  
  st <- as.character(df$state)
  if (!all(is.na(st))) {
    is_lag <- !is.na(st) & tolower(st) == "lag"
    lag_starts_idx <- which(c(TRUE, head(is_lag, -1) != tail(is_lag, 1)) & is_lag)
    tvec <- df$t_abs_s
    
    pick <- c(1, 5, 9, 13, 17, 21, 25)
    if (length(lag_starts_idx) > 0) {
      run_times <- tvec[lag_starts_idx]
      have_ord  <- pick[pick <= length(run_times)]
      if (length(have_ord) >= 1) {
        starts_times <- run_times[have_ord]
        breaks <- c(starts_times, Inf)
        labels_map <- c("1"="1","5"="2","9"="3","13"="4","17"="5","21"="6","25"="7")
        labs <- labels_map[as.character(have_ord)]
        
        alt_blk_chr <- cut(
          x = tvec,
          breaks = breaks,
          labels = labs,
          right  = FALSE,
          include.lowest = TRUE
        )
        df$block_alt <- suppressWarnings(as.integer(as.character(alt_blk_chr)))
  
        if (!(!is.na(grp) && grp == "NE")) {
          df$block_alt[df$block_alt == 7L] <- NA_integer_
        }
      } else {
        df$block_alt <- NA_integer_
      }
    } else {
      df$block_alt <- NA_integer_
    }
  } else {
    df$block_alt <- NA_integer_
  }
  
  df$file_name <- basename(fpath)
  df
}

all_list <- lapply(csv_files, compile_one)
all_data <- do.call(rbind, all_list[!vapply(all_list, is.null, logical(1))])

desired <- c("file_name", "group", "condition", "stage",
             "block_original", "block_final", "block_alt",
             "trial_idx", "state",
             "t_abs_s", "force_N", "force_std", "calibration_force")
keep <- intersect(desired, names(all_data))
all_data <- all_data[, c(keep, setdiff(names(all_data), keep))]

ext_mask <- !is.na(all_data$block_original) &
  tolower(trimws(all_data$block_original)) == "extinction_conf"

all_data$block_final[ext_mask & all_data$group == "E24"] <- 7L
all_data$block_final[ext_mask & all_data$group == "E16"] <- 5L
all_data$block_final[ext_mask & all_data$group == "E8"]  <- 3L

all_data$stage <- NA_character_
all_data$stage[all_data$block_final %in% c(2L, 4L, 6L)] <- "before"
all_data$stage[all_data$block_final %in% c(3L, 5L, 7L)] <- "after"

################INSPECT DATA THROUGH INDIVIDUAL PLOTS

contiguous_spans <- function(time, group) {
  ok <- !(is.na(time) | is.na(group))
  time <- time[ok]; group <- as.character(group[ok])
  if (!length(time)) return(data.frame(xmin=numeric(0), xmax=numeric(0), grp=character(0)))
  o <- order(time); t <- time[o]; g <- group[o]
  cuts <- c(1, which(g[-length(g)] != g[-1]) + 1)
  data.frame(
    xmin = t[cuts],
    xmax = c(t[pmax(cuts[-1] - 1, 1)], max(t)),
    grp  = g[cuts],
    stringsAsFactors = FALSE
  )
}

plot_participant_force_std <- function(all_data, file_pattern, use_block = c("block_final", "block_alt")) {
  use_block <- match.arg(use_block)
  hit <- grepl(file_pattern, all_data$file_name, ignore.case = TRUE)
  if (!any(hit)) stop("No rows matched file_pattern in all_data$file_name")
  
  df <- all_data[hit, , drop = FALSE]
  df <- df[order(df$t_abs_s), ]
  df$block <- df[[use_block]]
  df_force <- within(df, panel <- "Force")
  
  state_spans <- contiguous_spans(df$t_abs_s, df$state)
  if (nrow(state_spans)) {
    names(state_spans)[names(state_spans) == "grp"] <- "state"
    state_spans$state_clean <- tolower(trimws(state_spans$state))
    state_spans$state_key   <- paste0("state:", state_spans$state_clean)
    state_spans$panel <- "Force"
  }
  
  block_spans <- contiguous_spans(df$t_abs_s, df$block)
  if (nrow(block_spans)) {
    names(block_spans)[names(block_spans) == "grp"] <- "block"
    block_spans$block_fac <- factor(block_spans$block, levels = sort(unique(block_spans$block)))
    block_spans$block_key <- paste0("block:", block_spans$block_fac)
    block_spans$panel <- "Block"
    block_spans$ymin <- 0
    block_spans$ymax <- 1
  }
  
  #Thresholds
  hline_df <- data.frame(y = c(0.3, 0.7), panel = "Force")
  
  #Color-code behavioral states
  state_pal <- c(
    "state:lag"    = "yellow", # light grey
    "state:pursue" = "blue", # soft blue
    "state:idle"  = "black", # soft orange
    "state:reset"  = "brown"  # soft violet
  )
  #Color-code blocks
  block_lvls <- if (nrow(block_spans)) paste0("block:", levels(block_spans$block_fac)) else character(0)
  block_pal_full <- c("antiquewhite","#D9D9D9","#BDBDBD","#969696","#737373","#525252","#252525")
  block_pal <- setNames(block_pal_full[seq_along(block_lvls)], block_lvls)
  
  fill_values <- c(state_pal, block_pal)
  
  state_lvls_present <- if (nrow(state_spans)) unique(state_spans$state_key) else character(0)
  block_lvls_present <- block_lvls
  fill_breaks <- c(sort(state_lvls_present), block_lvls_present)
  fill_labels <- c(gsub("^state:", "", sort(state_lvls_present)),
                   paste("Block", gsub("^block:", "", block_lvls_present)))
  
  ggplot() +
    { if (nrow(state_spans))
      geom_rect(
        data = state_spans,
        aes(xmin = xmin, xmax = xmax, ymin = -Inf, ymax = Inf, fill = state_key),
        alpha = 0.5, inherit.aes = FALSE
      ) else NULL } +
    #Thresholds
    geom_hline(
      data = hline_df,
      aes(yintercept = y),
      linetype = "dashed", linewidth = 0.5
    ) +
    #Force trace (standardized)
    geom_line(
      data = df_force,
      aes(x = t_abs_s, y = force_std, color = factor(trial_idx), group = trial_idx),
      linewidth = 1.5, alpha = 0.99
    ) +
    
    { if (nrow(block_spans))
      geom_rect(
        data = block_spans,
        aes(xmin = xmin, xmax = xmax, ymin = ymin, ymax = ymax, fill = block_key),
        inherit.aes = FALSE
      ) else NULL } +
    facet_grid(rows = vars(panel), scales = "free_y", switch = "y") +
    labs(
      x = "Absolute time (s)",
      y = NULL,
      color = "Trial",
      fill  = "State / Block",
      title = unique(df$file_name)
    ) +
    theme_classic(base_size = 14) +
    theme(
      strip.placement = "outside",
      strip.text.y.left = element_text(angle = 0),
      legend.position = "bottom",
      legend.box = "vertical",
      panel.spacing.y = grid::unit(6, "pt"),
      axis.title.y = element_blank()
    ) +
    scale_y_continuous(expand = expansion(mult = c(0.02, 0.02))) +
    scale_fill_manual(values = fill_values, breaks = fill_breaks, labels = fill_labels, drop = FALSE) +
    guides(fill = guide_legend(order = 1), color = guide_legend(order = 2))
}

#Remove invalidated participant
all_data <- all_data %>%
  dplyr::filter(file_name != "40_timeseries_all.csv")

#Example: change the two-digit participant ID (01–64; skip 40) to plot a different file
plot_participant_force_std(all_data, file_pattern = "57_timeseries_all\\.csv$", use_block = "block_final")


###################INSPECT SELECTED TIMEFRAMES WITHIN A SESSION-PARTICIPANT UNIT

#Extract a time segment for one participant by ID and absolute-time window
segment_from_all_data <- function(all_data, id, t_start, t_end, pad = 0, ignore_case = TRUE, strict = TRUE) {
  stopifnot(is.data.frame(all_data), "file_name" %in% names(all_data),
            "t_abs_s" %in% names(all_data), "force_std" %in% names(all_data))
  
  if (t_end < t_start) { tmp <- t_start; t_start <- t_end; t_end <- tmp }
  
  #Match file_name
  pattern <- paste0("^", id, "_timeseries_all\\.csv$")
  hit <- grepl(pattern, all_data$file_name, ignore.case = ignore_case)
  
  if (!any(hit)) {
    stop(sprintf("No rows matched ID '%s' (expected file_name '%s').", id, paste0(id, "_timeseries_all.csv")))
  }
  
  df <- all_data[hit, c("t_abs_s", "force_std"), drop = FALSE]
  df <- df[order(df$t_abs_s), , drop = FALSE]
  
  lo <- t_start - pad
  hi <- t_end   + pad
  seg <- df[df$t_abs_s >= lo & df$t_abs_s <= hi, , drop = FALSE]
  
  names(seg) <- c("time_s", "force_std")
  rownames(seg) <- NULL
  
  if (strict && nrow(seg) == 0) {
    stop(sprintf("No samples in [%.3f, %.3f] seconds for ID '%s'.", t_start, t_end, id))
  }
  
  seg
}

#Examples

#To illustrate reward criterion
seg <- segment_from_all_data(all_data, id = "08", t_start = 217.5, t_end = 224.5)

ggplot(seg, aes(x = time_s, y = force_std)) +
  geom_line(linewidth = 0.8) +
  geom_hline(yintercept = c(0.3, 0.7), linetype = "dashed", linewidth = 0.6) +
  labs(x = "Time", y = "Standardized grip force") +
  scale_y_continuous(breaks = seq(0, 2, 0.1), expand = c(0, 0)) +
  coord_cartesian(ylim = c(0, 1.05)) +
  theme_classic(base_size = 13) +
  theme(
    axis.text.x  = element_blank(),
    axis.ticks.x = element_blank()
  )
#700 x 300

#To illustrate behavioral segments and block span calculation
seg <- segment_from_all_data(all_data, id = "01", t_start = 250, t_end = 256)

ggplot(seg, aes(x = time_s, y = force_std)) +
  geom_line(linewidth = 0.8) +
  geom_hline(yintercept = c(0.3, 0.7), linetype = "dashed", linewidth = 0.6) +
  labs(x = "Time", y = "Standardized grip force") +
  scale_y_continuous(breaks = seq(0, 2, 0.1), expand = c(0, 0)) +
  coord_cartesian(ylim = c(0, 1.05)) +
  theme_classic(base_size = 13) +
  theme(
    axis.text.x  = element_blank(),
    axis.ticks.x = element_blank()
  )
#700 x 300

#To illustrate metric calculation
seg <- segment_from_all_data(all_data, id = "48", t_start = 220.25, t_end = 224.25)

ggplot(seg, aes(x = time_s, y = force_std)) +
  geom_line(linewidth = 0.8) +
  geom_hline(yintercept = c(0.3, 0.7), linetype = "dashed", linewidth = 0.6) +
  labs(x = "Time", y = "Standardized grip force") +
  scale_y_continuous(breaks = seq(0, 2, 0.1), expand = c(0, 0)) +
  coord_cartesian(ylim = c(-0.025, 1.05)) +
  theme_classic(base_size = 13) +
  theme(
    axis.text.x  = element_blank(),
    axis.ticks.x = element_blank()
  )
```
