## Supplementary material for "Evidence for the extinction burst in grip force: A preregistered study": Electronic Supplementary Materials: 2_ebf_confirmatory_testing_n_group_viz.R

```
#Confirmatory statistical testing and group-level visualization

#Load packages
if (!require('dplyr'))      install.packages('dplyr');      library(dplyr)
if (!require('ggplot2'))    install.packages('ggplot2');    library(ggplot2)
if (!require('lme4'))       install.packages('lme4');       library(lme4)
if (!require('lmerTest'))   install.packages('lmerTest');   library(lmerTest)
if (!require('effectsize')) install.packages('effectsize'); library(effectsize)
if (!require("rptR")) install.packages("rptR"); library(rptR)

#Set alpha
alpha <- 0.05

#Build empirical summary_data from all_data
#drop blocks 0 and 1 (warm-up)
#obtain max(force_std) per file × group × condition × stage × block

summary_data <- all_data %>%
  dplyr::filter(!block_final %in% c(0, 1)) %>%
  dplyr::filter(stage %in% c("before", "after")) %>%
  dplyr::group_by(file_name, group, condition, stage, block_final) %>%
  dplyr::summarise(
    force_obs = {
      x <- force_std
      x <- x[is.finite(x)]
      if (length(x) == 0L) NA_real_ else max(x)
    },
    .groups = "drop"
  ) %>%
  dplyr::filter(!is.na(force_obs)) %>%
  dplyr::mutate(
    ID    = file_name,
    Group = factor(group, levels = c("E8","E16","E24","NE")),
    Block = as.integer(block_final),
    
    stage = ifelse(stage == "before", "before", "after"),
    stage = factor(stage, levels = c("before","after")),
    
    treatment = ifelse(tolower(condition) == "treat", "treat", "cft"),
    treatment = factor(treatment, levels = c("cft","treat"))
  ) %>%
  dplyr::select(ID, Group, Block, stage, treatment, force_obs)

#Plot faceted Group × Block means ± SE
plot_faceted_last <- function(d_last) {
  
  pal <- c(absent = "#228B22", present = "#8B008B")
  lw  <- 1.0
  
  agg_pts <- d_last %>%
    dplyr::mutate(
      Group     = factor(Group, levels = c("E8","E16","E24","NE")),
      Block     = as.integer(Block),
      stage     = factor(stage, levels = c("before","after")),
      treatment = factor(treatment, levels = c("cft","treat"),
                         labels = c("absent","present"))
    ) %>%
    dplyr::arrange(Group, Block, treatment, stage) %>%
    dplyr::group_by(Group, Block, treatment, stage) %>%
    dplyr::summarise(
      mean_y = mean(force_obs),
      se_y   = sd(force_obs)/sqrt(dplyr::n()),
      .groups = "drop"
    )
  
  agg_line <- agg_pts %>%
    dplyr::group_by(Group, Block, treatment) %>%
    dplyr::summarise(mean_y = mean(mean_y), .groups = "drop")
  
  seg <- agg_line %>%
    dplyr::group_by(Group, treatment) %>%
    dplyr::arrange(Block, .by_group = TRUE) %>%
    dplyr::mutate(Block_next = dplyr::lead(Block),
                  mean_next  = dplyr::lead(mean_y)) %>%
    dplyr::ungroup() %>%
    dplyr::filter(!is.na(Block_next) & (Block_next - Block == 1))
  
  ggplot() +
    geom_segment(
      data = seg,
      aes(x = Block, y = mean_y, xend = Block_next, yend = mean_next, color = treatment),
      linewidth = lw
    ) +
    geom_point(
      data = agg_pts,
      aes(x = Block, y = mean_y, color = treatment, shape = stage),
      size = 3.2,
      show.legend = c(color = FALSE, shape = TRUE)
    ) +
    geom_errorbar(
      data = agg_pts,
      aes(x = Block, ymin = mean_y - se_y, ymax = mean_y + se_y, color = treatment),
      width = 0.3, linewidth = lw
    ) +
    facet_wrap(~ Group, nrow = 1, drop = FALSE,
               labeller = labeller(Group = function(x) paste0("Group ", x))) +
    scale_color_manual(values = pal, breaks = c("absent","present")) +
    scale_x_continuous(breaks = sort(unique(agg_pts$Block))) +
    labs(
      x = "Block", y = "Mean peak force ± SEM",
      color = "Treatment", shape = "Stage"
    ) +
    theme_bw(base_size = 12) +
    theme(panel.grid.minor = element_blank())
}

#Mixed model for confirmatory preregistered hypothesis
last_m <- lmer(
  force_obs ~ stage * treatment + Block + (1 | ID),
  data = summary_data
)

print(summary(last_m))
confint(last_m)

#Obtain standardized betas
std_tab <- standardize_parameters(last_m, method = "refit")
print(std_tab)

coefs <- summary(last_m)$coefficients

b_int_raw <- unname(fixef(last_m)["stageafter:treatmenttreat"])
p_int_two <- unname(coefs["stageafter:treatmenttreat", "Pr(>|t|)"])
p_int_one <- ifelse(b_int_raw > 0, p_int_two/2, 1 - p_int_two/2)

std_col   <- grep("^Std", names(std_tab), value = TRUE)[1]
b_int_std <- std_tab[[std_col]][std_tab$Parameter == "stageafter:treatmenttreat"]
b_blk_std <- std_tab[[std_col]][std_tab$Parameter == "Block"]

cat("\n================ Results ================\n")
cat(sprintf("Interaction β (raw):          %.2f\n", b_int_raw))
cat(sprintf("Interaction β (standardized): %.2f\n", b_int_std))
cat(sprintf("Block β (standardized):       %.2f\n", b_blk_std))
cat(sprintf("Interaction p (two-sided):    %.10f\n", p_int_two))
cat(sprintf("Interaction p (one-sided +):  %.10f\n", p_int_one))

#Obtain repeatability (intra-class correlation) with confidence intervals
rpt_id <- rptGaussian(
  force_obs ~ stage * treatment + Block + (1 | ID),
  grname  = "ID",
  data    = summary_data,
  nboot   = 2000,
  npermut = 0
)

print(rpt_id)
plot(rpt_id)

#Build and print plot
p_facets <- plot_faceted_last(summary_data)
print(p_facets)
#700x300
```
