## Supplementary material for "Evidence for the extinction burst in grip force: A preregistered study": Electronic Supplementary Materials: 3_ebf_data_refinement.R

```
#Event-wise and block-wise data frame construction for downstream analyses

#Load packages
if (!require('dplyr')) install.packages('dplyr'); library(dplyr)

#Helper that builds an events data frame
build_events <- function(all_data) {
  stopifnot(
    all(c("file_name","group","condition","stage","block_final","block_original",
          "t_abs_s","force_std") %in% names(all_data))
  )
  
  all_data <- all_data %>%
    mutate(
      id = sub("_timeseries_all\\.csv$", "", file_name, ignore.case = TRUE),
      condition_lab = dplyr::case_when(
        tolower(condition) %in% c("treat")      ~ "present",
        tolower(condition) %in% c("ctfl","cft") ~ "absent",
        TRUE ~ NA_character_
      )
    )
  
  sp <- split(all_data, all_data$id)
  neq_na <- function(a, b) {
    (is.na(a) & !is.na(b)) | (!is.na(a) & is.na(b)) | (!is.na(a) & !is.na(b) & a != b)
  }
  safe_min <- function(x) { x <- x[is.finite(x)]; if (length(x)) min(x) else NA_real_ }
  safe_max <- function(x) { x <- x[is.finite(x)]; if (length(x)) max(x) else NA_real_ }
  
  ev_list <- lapply(sp, function(df) {
    df <- df[order(df$t_abs_s), , drop = FALSE]
    conf_blk <- df$block_final[tolower(trimws(df$block_original)) == "extinction_conf"]
    conf_blk <- suppressWarnings(as.integer(conf_blk[which(!is.na(conf_blk))[1]]))
    block_adj <- suppressWarnings(as.integer(df$block_final))
    is_expl   <- tolower(trimws(df$block_original)) == "extinction_expl"
    if (!is.na(conf_blk) && any(is_expl, na.rm = TRUE)) block_adj[is_expl] <- conf_blk + 1L
    df$block_adj <- block_adj
    
    b  <- df$block_adj
    ch <- c(TRUE, neq_na(b[-1], b[-length(b)]))
    start_idx <- which(ch & !is.na(b))
    
    blk_info <- data.frame(
      block_adj   = b[start_idx],
      block_start = df$t_abs_s[start_idx]
    ) |>
      dplyr::group_by(block_adj) |>
      dplyr::summarise(
        block_start = safe_min(block_start),
        block_end   = safe_max(df$t_abs_s[df$block_adj == first(block_adj)]),
        block_duration = ifelse(is.finite(block_start) & is.finite(block_end),
                                block_end - block_start, NA_real_),
        .groups = "drop"
      )
    
    t <- df$t_abs_s; f <- df$force_std; n <- nrow(df)
    ok <- is.finite(t) & is.finite(f); lagf <- c(NA_real_, f[-n])
    
    up03   <- ok & !is.na(lagf) & (lagf < 0.3) & (f >= 0.3)
    up07   <- ok & !is.na(lagf) & (lagf < 0.7) & (f >= 0.7)
    down03 <- ok & !is.na(lagf) & (lagf >= 0.3) & (f < 0.3)
    
    up03_i   <- which(up03)
    up07_i   <- which(up07)
    down03_i <- which(down03)
    
    if (is.finite(f[1]) && f[1] < 0.3) down03_i <- sort(unique(c(1L, down03_i)))
    
    next_after <- function(idx_vec, i) { j <- idx_vec[idx_vec > i]; if (length(j)) j[1] else NA_integer_ }
    first_between <- function(idx_vec, lo, hi) {
      j <- idx_vec[idx_vec >= lo & idx_vec < hi]   # inclusive lower bound
      if (length(j)) j[1] else NA_integer_
    }
    
    rows <- list()
    
    for (u in up03_i) {
      d <- next_after(down03_i, u); if (is.na(d)) next
      m <- first_between(up07_i, u, d)
      
      rng <- u:d
      dur <- t[d] - t[u]
      mx  <- suppressWarnings(max(f[rng], na.rm = TRUE)); if (!is.finite(mx)) next
      
      if (is.na(m)) { at <- u; ev <- "subthreshold" } else { at <- m; ev <- "criterion" }
      
      this_blk   <- df$block_adj[at]
      this_stage <- df$stage[at]
      this_group <- df$group[at]
      this_cond  <- df$condition_lab[at]
      
      bi <- blk_info[match(this_blk, blk_info$block_adj), ]
      bstart <- bi$block_start; bdur <- bi$block_duration
      
      rows[[length(rows) + 1L]] <- data.frame(
        id             = df$id[at],
        group          = this_group,
        condition      = this_cond,            
        stage          = this_stage,
        block          = this_blk,
        block_starts   = bstart,
        block_duration = bdur,
        event          = ev,
        timestamp      = t[at],
        max            = mx,                    
        duration       = dur,                  
        stringsAsFactors = FALSE
      )
    }
    
    for (d in down03_i) {
      u2 <- next_after(up03_i, d)
      if (is.na(u2) && is.finite(f[n]) && f[n] < 0.3) u2 <- n
      if (is.na(u2)) next
      
      rng <- d:u2
      mn  <- suppressWarnings(min(f[rng], na.rm = TRUE))
      dur <- t[u2] - t[d]
      
      this_blk   <- df$block_adj[d]
      this_stage <- df$stage[d]
      this_group <- df$group[d]
      this_cond  <- df$condition_lab[d]
      
      bi <- blk_info[match(this_blk, blk_info$block_adj), ]
      bstart <- bi$block_start; bdur <- bi$block_duration
      
      rows[[length(rows) + 1L]] <- data.frame(
        id             = df$id[d],
        group          = this_group,
        condition      = this_cond,        
        stage          = this_stage,
        block          = this_blk,
        block_starts   = bstart,
        block_duration = bdur,
        event          = "idle",
        timestamp      = t[d],              
        max            = mn,                 
        duration       = dur,                 
        stringsAsFactors = FALSE
      )
    }
    
    out <- if (length(rows)) do.call(rbind, rows) else
      data.frame(id=character(0), group=character(0), condition=character(0),
                 stage=character(0), block=integer(0),
                 block_starts=numeric(0), block_duration=numeric(0),
                 event=character(0), timestamp=numeric(0),
                 max=numeric(0), duration=numeric(0))
    
    all_blks <- sort(unique(blk_info$block_adj))
    present_blks <- sort(unique(out$block))
    missing_blks <- setdiff(all_blks, present_blks)
    
    if (length(missing_blks)) {
      none_rows <- lapply(missing_blks, function(bb) {
        ii <- which(df$block_adj == bb)
        at <- if (length(ii)) ii[1] else NA_integer_
        
        bi <- blk_info[match(bb, blk_info$block_adj), ]
        bstart <- bi$block_start
        bdur   <- bi$block_duration
        
        data.frame(
          id             = if (!is.na(at)) df$id[at] else df$id[1],
          group          = if (!is.na(at)) df$group[at] else df$group[1],
          condition      = if (!is.na(at)) df$condition_lab[at] else NA_character_,
          stage          = if (!is.na(at)) df$stage[at] else NA_character_,
          block          = bb,
          block_starts   = bstart,
          block_duration = bdur,
          event          = "none",
          timestamp      = NA_real_,
          max            = NA_real_,
          duration       = NA_real_,
          stringsAsFactors = FALSE
        )
      })
      out <- dplyr::bind_rows(out, do.call(rbind, none_rows))
    }
    
    out
  })
  
  events <- do.call(rbind, ev_list)
  
  events %>%
    mutate(
      condition = factor(condition, levels = c("absent","present")),
      stage     = factor(stage, levels = c("before","after")),
      group     = factor(group, levels = c("E8","E16","E24","NE"))
    ) %>%
    arrange(id, block, timestamp)
}

events <- build_events(all_data)

####################################################BLOCKWISE DATA FRAME

safe_div <- function(num, den) ifelse(is.finite(den) & den > 0, num / den, NA_real_)

blockwise <- events %>%
  group_by(id, group, condition, stage, block) %>%
  summarise(
    block_starts   = dplyr::first(block_starts),
    block_duration = dplyr::first(block_duration),

    n_criterion    = sum(event == "criterion",    na.rm = TRUE),
    n_subthreshold = sum(event == "subthreshold", na.rm = TRUE),

    avg_d_criterion = {
      v <- duration[event == "criterion"]; if (length(v)) mean(v, na.rm = TRUE) else NA_real_
    },
    avg_d_subthreshold = {
      v <- duration[event == "subthreshold"]; if (length(v)) mean(v, na.rm = TRUE) else NA_real_
    },

    tot_idle = {
      v <- duration[event == "idle"]; if (length(v)) sum(v, na.rm = TRUE) else 0
    },

    max_d_criterion = {
      v <- duration[event == "criterion"]; if (length(v)) max(v, na.rm = TRUE) else NA_real_
    },
    max_d_subthreshold = {
      v <- duration[event == "subthreshold"]; if (length(v)) max(v, na.rm = TRUE) else NA_real_
    },

    avg_peak_criterion = {
      v <- max[event == "criterion"]; if (length(v)) mean(v, na.rm = TRUE) else NA_real_
    },
    avg_peak_subthreshold = {
      v <- max[event == "subthreshold"]; if (length(v)) mean(v, na.rm = TRUE) else NA_real_
    },
    avg_dip_idle = {
      v <- max[event == "idle"]; if (length(v)) mean(v, na.rm = TRUE) else NA_real_
    },

    block_peak_criterion = {
      v <- max[event == "criterion"]; if (length(v)) max(v, na.rm = TRUE) else NA_real_
    },
    block_peak_subthreshold = {
      v <- max[event == "subthreshold"]; if (length(v)) max(v, na.rm = TRUE) else NA_real_
    },
    block_trough_idle = {
      v <- max[event == "idle"]; if (length(v)) min(v, na.rm = TRUE) else NA_real_
    },

    force_variability = {
      v <- max[event %in% c("criterion","subthreshold")]
      if (length(v) > 1) stats::sd(v, na.rm = TRUE) else NA_real_
    },

    duration_variability = {
      v <- duration[event %in% c("criterion","subthreshold")]
      if (length(v) > 1) stats::sd(v, na.rm = TRUE) else NA_real_
    },
    .groups = "drop"
  ) %>%
  group_by(id) %>%
  arrange(block, .by_group = TRUE) %>%
  mutate(
    criterion_rate    = ifelse(n_criterion    == 0L, 0, safe_div(n_criterion,    block_duration)),
    subthreshold_rate = ifelse(n_subthreshold == 0L, 0, safe_div(n_subthreshold, block_duration)),
    idle_quotient     = ifelse(tot_idle       == 0,  0, safe_div(tot_idle,       block_duration)),
    
    block_diff        = block_duration - dplyr::lag(block_duration)
  )%>%
  ungroup() %>%
  arrange(id, block)
```
