## Supplementary material for "Evidence for the extinction burst in grip force: A preregistered study": Electronic Supplementary Materials: 4.1_ebf_bootstrap_inference.R

```
# =============================================================================
# WARNING
# =============================================================================
#
# Run the ENTIRE script at once by selecting all and then hitting run. Do NOT execute it line-by-line or chunk-
# by-chunk with Ctrl+Enter, or it will hang.


#Load required packages
if (!require("dplyr"))    install.packages("dplyr");    library(dplyr)
if (!require("lme4"))     install.packages("lme4");     library(lme4)
if (!require("lmerTest")) install.packages("lmerTest"); library(lmerTest)
if (!require("ggplot2"))  install.packages("ggplot2");  library(ggplot2)
if (!require("parallel")) install.packages("parallel"); library(parallel)


#number of bootstrap resamples per outcome x analysis
nsim_boot <- 10000

#Seed for reproducibility
master_seed <- 1401

#Outcomes variables
outcomes_of_interest <- c(
  "block_peak_criterion",
  "criterion_rate",
  "block_trough_idle",
  "avg_d_criterion",
  "force_variability"
)

#This sets the number of parallel cores to execute this computationally
#...intensive analysis, leaving one whole core free for other tasks.
n_cores <- max(1L, parallel::detectCores() - 1L)

#Number of chunks to split each bootstrap into for progress bar ticks
n_chunks <- 50L

#Standardize outcome variables to report betas on a common scale across behavioral indices
standardize_safe <- function(x) {
  x <- as.numeric(x)
  m <- mean(x, na.rm = TRUE)
  s <- stats::sd(x, na.rm = TRUE)

  #return NAs rather than producing Inf or NaN
  if (!is.finite(m) || !is.finite(s) || s == 0) {
    return(rep(NA_real_, length(x)))
  }

  (x - m) / s
}

#Fit the mixed model and pull the interaction term
fit_interaction_model <- function(d) {

  d <- d[
    is.finite(d$y) &
      !is.na(d$treatment) &
      !is.na(d$stage) &
      is.finite(d$block),
    , drop = FALSE
  ]

  if (nrow(d) < 10L ||
      length(unique(as.character(d$treatment))) < 2L ||
      length(unique(as.character(d$stage))) < 2L) {
    return(NULL)
  }

  #By fixing levels to c("absent","present") and c("before","after"), the
  #...interaction is always called "treatmentpresent:stageafter"
  d$participant <- factor(d$participant)
  d$treatment   <- factor(as.character(d$treatment), levels = c("absent", "present"))
  d$stage       <- factor(as.character(d$stage),     levels = c("before", "after"))
  d$block       <- as.numeric(d$block)

  fit <- tryCatch(
    lmer(
      y ~ treatment * stage + block + (1 | participant),
      data = d,
      REML = FALSE,
      control = lmerControl(
        optimizer = "bobyqa",
        optCtrl = list(maxfun = 2e5)
      )
    ),
    error = function(e) NULL
  )

  if (is.null(fit)) return(NULL)

  #Extract the interaction row from the fixed-effects coefficient table
  co <- summary(fit)$coefficients
  rn <- rownames(co)

  inter <- rn[grepl("treatment.*:.*stage|stage.*:.*treatment", rn)]
  if (!length(inter)) inter <- "treatmentpresent:stageafter"
  if (!inter %in% rn) return(NULL)

  list(
    fit     = fit,
    data    = d,
    inter   = inter,
    beta    = unname(co[inter, "Estimate"]),
    se      = unname(co[inter, "Std. Error"]),
    t       = unname(co[inter, "t value"]),
    df      = if ("df" %in% colnames(co)) unname(co[inter, "df"]) else NA_real_,
    p_model = if ("Pr(>|t|)" %in% colnames(co)) unname(co[inter, "Pr(>|t|)"]) else NA_real_,
    n       = nrow(d)
  )
}

#When a participant is sampled twice, the two copies of their data must
#...be treated as two separate participants, not one participant
#...with twice the data. Otherwise, the very heterogeneity the bootstrap
#...is trying to capture collapses.
boot_chunk <- function(chunk_size, obs_data, ids, chunk_seed, chunk_offset) {

  set.seed(chunk_seed)

  n_ids <- length(ids)
  out_beta <- rep(NA_real_, chunk_size)
  out_se   <- rep(NA_real_, chunk_size)

  for (i in seq_len(chunk_size)) {

    #sample n_ids participants with replacement.
    sampled_ids <- sample(ids, size = n_ids, replace = TRUE)

    #pull each sampled cluster's full set of rows and relabel.
    boot_list <- vector("list", length(sampled_ids))
    for (j in seq_along(sampled_ids)) {
      tmp <- obs_data[
        as.character(obs_data$participant) == sampled_ids[j],
        c("participant", "treatment", "stage", "block", "y"),
        drop = FALSE
      ]

      tmp$participant <- paste0("boot_", chunk_offset + i, "_", j)
      boot_list[[j]]  <- tmp
    }

    db <- do.call(rbind, boot_list)

    fit_b <- fit_interaction_model(db)

    if (!is.null(fit_b)) {
      out_beta[i] <- fit_b$beta
      out_se[i]   <- fit_b$se
    }
  }

  cbind(beta = out_beta, se = out_se)
}

#run the bootstrap in parallel chunks by splitting nsim total iterations into n_chunks chunks of equal size
#After all chunks return, the matrix of (beta, se) draws is assembled and
#...the studentized confidence interval and p-value are computed.
cluster_boot_interaction <- function(d,
                                     nsim        = 5000,
                                     seed        = NULL,
                                     n_cores     = 1L,
                                     n_chunks    = 50L,
                                     show_progress = TRUE) {

  if (!is.null(seed)) set.seed(seed)

  obs <- fit_interaction_model(d)
  if (is.null(obs)) return(NULL)

  ids <- unique(as.character(obs$data$participant))

  cut_points <- floor(seq(0, nsim, length.out = n_chunks + 1L))
  chunk_sizes <- diff(cut_points)

  #Each chunk gets a different seed so its random draws
  #...are independent of other chunks
  chunk_seeds <- if (is.null(seed)) {
    sample.int(.Machine$integer.max, n_chunks)
  } else {
    seed + seq_len(n_chunks)
  }

  chunk_offsets <- cut_points[-length(cut_points)]

  cl <- parallel::makeCluster(n_cores)
  on.exit(parallel::stopCluster(cl), add = TRUE)

  parallel::clusterEvalQ(cl, {
    library(lme4)
    library(lmerTest)
  })
  parallel::clusterExport(
    cl,
    varlist = c("fit_interaction_model", "boot_chunk"),
    envir   = environment()
  )

  #Progress bar that ticks once per completed chunk
  if (show_progress) {
    pb <- txtProgressBar(min = 0, max = n_chunks, style = 3)
  }

  results <- vector("list", n_chunks)
  next_chunk <- 1L

  while (next_chunk <= n_chunks) {

    batch_end <- min(next_chunk + n_cores - 1L, n_chunks)
    batch_idx <- next_chunk:batch_end

    batch_result <- parallel::clusterApply(
      cl,
      x   = batch_idx,
      fun = function(k, sizes, seeds, offsets, obs_data, ids) {
        boot_chunk(
          chunk_size   = sizes[k],
          obs_data     = obs_data,
          ids          = ids,
          chunk_seed   = seeds[k],
          chunk_offset = offsets[k]
        )
      },
      sizes    = chunk_sizes,
      seeds    = chunk_seeds,
      offsets  = chunk_offsets,
      obs_data = obs$data,
      ids      = ids
    )

    for (k in seq_along(batch_idx)) {
      results[[batch_idx[k]]] <- batch_result[[k]]
    }

    if (show_progress) setTxtProgressBar(pb, batch_end)
    next_chunk <- batch_end + 1L
  }

  if (show_progress) close(pb)

  big <- do.call(rbind, results)
  boot_beta <- big[, "beta"]
  boot_se   <- big[, "se"]

  #build the studentized t* distribution
  keep_t <- is.finite(boot_beta) & is.finite(boot_se) & boot_se > 0
  t_star <- (boot_beta[keep_t] - obs$beta) / boot_se[keep_t]
  t_star <- t_star[is.finite(t_star)]

  keep_beta <- is.finite(boot_beta)
  boot_beta_ok <- boot_beta[keep_beta]

  n_boot_beta <- length(boot_beta_ok)
  n_boot_t    <- length(t_star)

  if (n_boot_beta < 200L) {
    warning("Fewer than 200 successful bootstrap beta draws.")
  }

  ci_low  <- NA_real_
  ci_high <- NA_real_
  p_boot  <- NA_real_
  ci_method <- NA_character_
  p_method  <- NA_character_

  if (n_boot_t >= 200L && is.finite(obs$se) && obs$se > 0 && is.finite(obs$t)) {

    #2.5 and 97.5 percentiles of the t* distribution
    q025 <- unname(stats::quantile(t_star, 0.025, na.rm = TRUE, type = 6))
    q975 <- unname(stats::quantile(t_star, 0.975, na.rm = TRUE, type = 6))

    ci_low  <- obs$beta - q975 * obs$se
    ci_high <- obs$beta - q025 * obs$se

    #Two-sided bootstrap p-value with the +1 correction.
    #Because plain (sum >= obs)/B can return 0 when no resample matches or exceeds
    #...the observed |t|, which is a misleading "p = 0", +1 in numerator
    #...and denominator is included to yield a strictly positive p-value.
    p_boot <- (sum(abs(t_star) >= abs(obs$t)) + 1) / (n_boot_t + 1)

    ci_method <- "cluster bootstrap-t"
    p_method  <- "cluster bootstrap-t"

  } else if (n_boot_beta >= 200L) {

    ci_low  <- unname(stats::quantile(boot_beta_ok, 0.025, na.rm = TRUE, type = 6))
    ci_high <- unname(stats::quantile(boot_beta_ok, 0.975, na.rm = TRUE, type = 6))

    centered_dev <- boot_beta_ok - obs$beta
    p_boot <- (sum(abs(centered_dev) >= abs(obs$beta)) + 1) / (n_boot_beta + 1)

    ci_method <- "cluster bootstrap percentile"
    p_method  <- "cluster bootstrap centered"
  }

  #significance nomenclature for the forest plot
  sig <- ifelse(is.na(p_boot), NA_character_,
                ifelse(p_boot < 0.001, "***",
                       ifelse(p_boot < 0.01,  "**",
                              ifelse(p_boot < 0.05,  "*", "ns"))))

  data.frame(
    n           = obs$n,
    beta        = obs$beta,
    se          = obs$se,
    df          = obs$df,
    t           = obs$t,
    p           = p_boot,
    p_model     = obs$p_model,
    ci_low      = ci_low,
    ci_high     = ci_high,
    sig         = sig,
    n_boot_beta = n_boot_beta,
    n_boot_t    = n_boot_t,
    ci_method   = ci_method,
    p_method    = p_method
  )
}


#run all outcome x analysis combinations
make_bw_forest_results <- function(blockwise,
                                   outcomes,
                                   nsim_boot = 5000,
                                   seed      = 1401,
                                   n_cores   = 1L,
                                   n_chunks  = 50L) {

  #Drop warm-up blocks (block 0 and 1)
  bw0 <- blockwise %>%
    dplyr::filter(!(as.numeric(block) %in% c(0, 1))) %>%
    dplyr::mutate(
      participant   = factor(id),
      block         = as.numeric(block),
      condition_chr = as.character(condition),
      stage_chr     = as.character(stage)
    )

  #standardization
  for (oc in intersect(outcomes, names(bw0))) {
    bw0[[oc]] <- standardize_safe(bw0[[oc]])
  }

  #early extinction analyses
  bw_early <- bw0 %>%
    dplyr::filter(!is.na(condition_chr) & !is.na(stage_chr)) %>%
    dplyr::mutate(
      treatment = factor(condition_chr, levels = c("absent", "present")),
      stage     = factor(stage_chr,     levels = c("before", "after")),
      analysis  = "early"
    )

  #late extinction analyses
  bw_late <- bw0 %>%
    dplyr::filter(!(!is.na(condition_chr) & !is.na(stage_chr) &
                      condition_chr == "present" & stage_chr == "after")) %>%
    dplyr::mutate(
      condition_chr = ifelse(is.na(condition_chr), "present", condition_chr),
      stage_chr     = ifelse(is.na(stage_chr),     "after",   stage_chr),
      treatment     = factor(condition_chr, levels = c("absent", "present")),
      stage         = factor(stage_chr,     levels = c("before", "after")),
      analysis      = "late"
    )

  #run the bootstrap for one (outcome, extinction stage) combination.
  run_one <- function(df, outcome, nsim_boot, seed) {

    y <- df[[outcome]]
    if (!is.numeric(y)) y <- suppressWarnings(as.numeric(y))

    d <- data.frame(
      participant = df$participant,
      treatment   = df$treatment,
      stage       = df$stage,
      block       = df$block,
      y           = y     # already standardized upstream
    )

    d <- d[
      is.finite(d$y) &
        !is.na(d$treatment) &
        !is.na(d$stage) &
        is.finite(d$block),
      , drop = FALSE
    ]

    na_row <- data.frame(
      outcome      = outcome,
      n            = nrow(d),
      beta         = NA_real_,
      se           = NA_real_,
      df           = NA_real_,
      t            = NA_real_,
      p            = NA_real_,
      p_model      = NA_real_,
      ci_low       = NA_real_,
      ci_high      = NA_real_,
      sig          = NA_character_,
      n_boot_beta  = NA_real_,
      n_boot_t     = NA_real_,
      ci_method    = NA_character_,
      p_method     = NA_character_
    )

    if (nrow(d) < 10L ||
        length(unique(as.character(d$treatment))) < 2L ||
        length(unique(as.character(d$stage))) < 2L) {
      return(na_row)
    }

    res <- cluster_boot_interaction(
      d,
      nsim          = nsim_boot,
      seed          = seed,
      n_cores       = n_cores,
      n_chunks      = n_chunks,
      show_progress = TRUE
    )

    if (is.null(res)) return(na_row)

    res$outcome <- outcome
    res
  }

  outcomes <- intersect(outcomes, names(bw0))

  cat("Bootstrapping early extinction models...\n")
  res_early <- do.call(rbind, lapply(seq_along(outcomes), function(i) {
    o <- outcomes[i]
    cat("\nEARLY:", o, "\n")
    run_one(bw_early, o, nsim_boot = nsim_boot, seed = seed + i)
  }))

  cat("\nBootstrapping late extinction models...\n")
  res_late <- do.call(rbind, lapply(seq_along(outcomes), function(i) {
    o <- outcomes[i]
    cat("\nLATE:", o, "\n")
    run_one(bw_late, o, nsim_boot = nsim_boot, seed = seed + 100 + i)
  }))

  res_early$analysis <- "early"
  res_late$analysis  <- "late"

  dplyr::bind_rows(res_early, res_late)
}

#Run
bw_forest_results <- make_bw_forest_results(
  blockwise  = blockwise,
  outcomes   = outcomes_of_interest,
  nsim_boot  = nsim_boot,
  seed       = master_seed,
  n_cores    = n_cores,
  n_chunks   = n_chunks
)

#Forest plot
bw_forest_plot_df <- bw_forest_results
bw_forest_plot_df$Extinction <- factor(bw_forest_plot_df$analysis, levels = c("late", "early"))

#Display names for the y-axis
name_map <- c(
  "block_peak_criterion" = "peak force",
  "criterion_rate"       = "response rate",
  "block_trough_idle"    = "minimum idle force",
  "avg_d_criterion"      = "response duration",
  "force_variability"    = "force variability"
)

bw_forest_plot_df$outcome <- as.character(bw_forest_plot_df$outcome)
bw_forest_plot_df$outcome[bw_forest_plot_df$outcome %in% names(name_map)] <-
  unname(name_map[bw_forest_plot_df$outcome[bw_forest_plot_df$outcome %in% names(name_map)]])

bw_forest_plot_df <- bw_forest_plot_df[
  !is.na(bw_forest_plot_df$outcome) &
    bw_forest_plot_df$outcome %in% unname(name_map),
  , drop = FALSE
]

bw_forest_plot_df$outcome <- factor(bw_forest_plot_df$outcome, levels = rev(unname(name_map)))

p_forest <- ggplot(bw_forest_plot_df, aes(x = beta, y = outcome, color = Extinction)) +
  geom_vline(xintercept = 0, linetype = 2) +
  geom_pointrange(aes(xmin = ci_low, xmax = ci_high),
                  position = position_dodge2(width = 0.60, preserve = "single"),
                  linewidth = 0.9) +
  geom_text(
    aes(label = sig, vjust = ifelse(Extinction == "early", -0.6, 1.9)),
    position = position_dodge2(width = 0.60, preserve = "single"),
    size = 3.6, show.legend = FALSE
  ) +
  scale_color_manual(
    values = c("early" = "#FF4D00", "late" = "gray55"),
    breaks = c("early", "late"),
    drop = FALSE,
    name = "Extinction"
  ) +
  labs(x = "Standardized treatment x stage interaction beta with 95% bootstrap CI") +
  theme_classic(base_size = 13) +
  theme(
    legend.position = "top",
    axis.title.y = element_blank(),
    legend.title = element_text(size = 10)
  )

p_forest

bw_forest_results
#750x450

#####################################################################################
```
