## Supplementary material for "Evidence for the extinction burst in grip force: A preregistered study": Electronic Supplementary Materials: 4_ebf_effect_size_visualization.R

```
#Visualization of standardized interaction betas

#Load packages
if (!require("dplyr"))      install.packages("dplyr");      library(dplyr)
if (!require("lme4"))       install.packages("lme4");       library(lme4)
if (!require("lmerTest"))   install.packages("lmerTest");   library(lmerTest)
if (!require("ggplot2"))    install.packages("ggplot2");    library(ggplot2)
if (!require("effectsize")) install.packages("effectsize"); library(effectsize)

outcomes_of_interest <- c(
  "block_peak_criterion",
  "criterion_rate",
  "block_trough_idle",
  "avg_d_criterion",
  "force_variability"
)

make_bw_forest_results <- function(blockwise, outcomes) {
  
  bw0 <- blockwise %>%
    dplyr::filter(!(as.numeric(block) %in% c(0, 1))) %>%
    dplyr::mutate(
      participant   = factor(id),
      block         = as.numeric(block),
      condition_chr = as.character(condition),
      stage_chr     = as.character(stage)
    )
  
  #Early extinction contrasts
  bw_early <- bw0 %>%
    dplyr::filter(!is.na(condition_chr) & !is.na(stage_chr)) %>%
    dplyr::mutate(
      treatment = factor(condition_chr, levels = c("absent","present")),
      stage     = factor(stage_chr,     levels = c("before","after")),
      analysis  = "early"
    )
  
  #Late extinction
  bw_late <- bw0 %>%
    dplyr::filter(!( !is.na(condition_chr) & !is.na(stage_chr) &
                       condition_chr == "present" & stage_chr == "after")) %>%
    dplyr::mutate(
      condition_chr = ifelse(is.na(condition_chr), "present", condition_chr),
      stage_chr     = ifelse(is.na(stage_chr),     "after",   stage_chr),
      treatment     = factor(condition_chr, levels = c("absent","present")),
      stage         = factor(stage_chr,     levels = c("before","after")),
      analysis      = "late"
    )
  
  run_one <- function(df, outcome) {
    y <- df[[outcome]]
    if (!is.numeric(y)) y <- suppressWarnings(as.numeric(y))
    
    d <- data.frame(
      participant = df$participant,
      treatment   = df$treatment,
      stage       = df$stage,
      block       = df$block,
      y           = y
    )
    
    d <- d[is.finite(d$y) &
             !is.na(d$treatment) & !is.na(d$stage) &
             is.finite(d$block), , drop = FALSE]
    
    if (nrow(d) < 10L ||
        length(unique(d$treatment)) < 2L ||
        length(unique(d$stage)) < 2L) {
      return(data.frame(outcome=outcome, n=nrow(d),
                        beta=NA_real_, se=NA_real_, df=NA_real_, t=NA_real_, p=NA_real_,
                        ci_low=NA_real_, ci_high=NA_real_, sig=NA_character_))
    }
    
    d$z_y <- as.numeric(effectsize::standardize(d$y))
    d <- d[is.finite(d$z_y), , drop = FALSE]
    
    if (nrow(d) < 10L) {
      return(data.frame(outcome=outcome, n=nrow(d),
                        beta=NA_real_, se=NA_real_, df=NA_real_, t=NA_real_, p=NA_real_,
                        ci_low=NA_real_, ci_high=NA_real_, sig=NA_character_))
    }
    
    fit <- lmer(z_y ~ treatment * stage + block + (1 | participant),
                data = d, REML = FALSE)
    
    co <- summary(fit)$coefficients
    rn <- rownames(co)
    
    inter <- rn[grepl("treatment.*:.*stage|stage.*:.*treatment", rn)]
    inter <- if (length(inter)) inter[1] else "treatmentpresent:stageafter"
    
    if (!inter %in% rn) {
      return(data.frame(outcome=outcome, n=nrow(d),
                        beta=NA_real_, se=NA_real_, df=NA_real_, t=NA_real_, p=NA_real_,
                        ci_low=NA_real_, ci_high=NA_real_, sig=NA_character_))
    }
    
    beta <- unname(co[inter, "Estimate"])   
    se   <- unname(co[inter, "Std. Error"])
    df_  <- unname(co[inter, "df"])
    t_   <- unname(co[inter, "t value"])
    p_   <- unname(co[inter, "Pr(>|t|)"])
    
    crit <- if (is.finite(df_)) stats::qt(0.975, df_) else 1.96
    ci_low  <- beta - crit * se
    ci_high <- beta + crit * se
    
    sig <- ifelse(is.na(p_), NA_character_,
                  ifelse(p_ < 0.001, "***",
                         ifelse(p_ < 0.01,  "**",
                                ifelse(p_ < 0.05,  "*", "ns"))))
    
    data.frame(outcome=outcome, n=nrow(d),
               beta=beta, se=se, df=df_, t=t_, p=p_,
               ci_low=ci_low, ci_high=ci_high, sig=sig)
  }
  
  outcomes <- intersect(outcomes, names(bw0))
  
  res_early <- do.call(rbind, lapply(outcomes, function(o) run_one(bw_early, o)))
  res_late  <- do.call(rbind, lapply(outcomes, function(o) run_one(bw_late,  o)))
  
  res_early$analysis <- "early"
  res_late$analysis  <- "late"
  
  dplyr::bind_rows(res_early, res_late)
}

bw_forest_results <- make_bw_forest_results(blockwise, outcomes_of_interest)
bw_forest_plot_df <- bw_forest_results

bw_forest_plot_df$Extinction <- if ("Extinction" %in% names(bw_forest_plot_df)) {
  bw_forest_plot_df$Extinction
} else {
  bw_forest_plot_df$analysis
}

bw_forest_plot_df$Extinction <- as.character(bw_forest_plot_df$Extinction)
bw_forest_plot_df$Extinction[bw_forest_plot_df$Extinction == "confirmatory"] <- "early"
bw_forest_plot_df$Extinction[bw_forest_plot_df$Extinction == "exploratory"]  <- "late"
bw_forest_plot_df$Extinction <- factor(bw_forest_plot_df$Extinction, levels = c("late","early"))

#Outcome display names
name_map <- c(
  "block_peak_criterion" = "peak force",
  "criterion_rate"       = "response rate",
  "block_trough_idle"    = "minimum idle force",
  "avg_d_criterion"      = "response duration",
  "force_variability"    = "force variability"
)

bw_forest_plot_df$outcome <- as.character(bw_forest_plot_df$outcome)
bw_forest_plot_df$outcome[bw_forest_plot_df$outcome %in% names(name_map)] <-
  unname(name_map[bw_forest_plot_df$outcome[bw_forest_plot_df$outcome %in% names(name_map)]])

bw_forest_plot_df <- bw_forest_plot_df[
  !is.na(bw_forest_plot_df$outcome) &
    bw_forest_plot_df$outcome %in% unname(name_map),
  , drop = FALSE
]

bw_forest_plot_df$outcome <- factor(bw_forest_plot_df$outcome, levels = rev(unname(name_map)))

if (!("sig" %in% names(bw_forest_plot_df))) {
  bw_forest_plot_df$sig <- ifelse(is.na(bw_forest_plot_df$p), NA_character_,
                                  ifelse(bw_forest_plot_df$p < 0.001, "***",
                                         ifelse(bw_forest_plot_df$p < 0.01,  "**",
                                                ifelse(bw_forest_plot_df$p < 0.05,  "*", "ns"))))
}

#Plot
pos <- tryCatch(position_dodge2(width = 0.60, preserve = "single"),
                error = function(e) position_dodge(width = 0.60))

geom_pr <- function(...) {
  tryCatch(geom_pointrange(..., linewidth = 0.9),
           error = function(e) geom_pointrange(..., size = 0.9))
}

p_forest <- ggplot(bw_forest_plot_df, aes(x = beta, y = outcome, color = Extinction)) +
  geom_vline(xintercept = 0, linetype = 2) +
  geom_pr(aes(xmin = ci_low, xmax = ci_high), position = pos) +
  geom_text(
    aes(label = sig, vjust = ifelse(Extinction == "early", -0.6, 1.9)),
    position = pos, size = 3.6, show.legend = FALSE
  ) +
  scale_color_manual(
    values = c("early" = "#FF4D00", "late" = "gray55"),
    breaks = c("early","late"),
    drop = FALSE,
    name = "Extinction"
  ) +
  labs(x = "Standardized treatment × stage interaction β ± 95% CI") +
  theme_classic(base_size = 13) +
  theme(
    legend.position = "top",
    axis.title.y = element_blank(),
    legend.title = element_text(size = 10)
  )

p_forest
```
