## Supplementary material for "Evidence for the extinction burst in grip force: A preregistered study": Electronic Supplementary Materials: 5_ebf_episode-matched_analysis.R

```
#Load required packages
if (!require("ggplot2")) install.packages("ggplot2"); library(ggplot2)

#SETTINGS

B <- 4000 #Number of iterations for bootstrapping
ci_level <- 0.68              #For visualization only

col_absent  <- "#228B22"       
col_present <- "#8B008B"      
cols <- c(absent = col_absent, present = col_present)

#Tier structure 
tier_map <- data.frame(
  tier = c("t1","t2","t3"),
  pre_block  = c(2, 4, 6),
  post_block = c(3, 5, 7),
  stringsAsFactors = FALSE
)

#AUGMENT EVENTS DATA FRAME
events_aug <- events
events_aug$stage_raw <- events_aug$stage
events_aug$condition_raw <- events_aug$condition

events_aug$stage <- as.character(events_aug$stage)
events_aug$condition <- as.character(events_aug$condition)

#Late extinction NAs treatment
events_aug$stage[is.na(events_aug$stage)] <- "after"
events_aug$condition[is.na(events_aug$condition)] <- "present"

events_aug$condition <- factor(events_aug$condition, levels = c("absent","present"))
events_aug$event <- as.character(events_aug$event)

#Build data frame containing metrics per criterion response

crit_aug <- events_aug[events_aug$event == "criterion", , drop = FALSE]
crit_aug <- crit_aug[order(crit_aug$id, crit_aug$timestamp), , drop = FALSE]

#previous criterion timestamp
crit_aug$prev_crit_ts_global <- ave(
  crit_aug$timestamp, crit_aug$id,
  FUN = function(x) c(NA_real_, x[-length(x)])
)

#Idle computations
idle_tbl <- events_aug[events_aug$event == "idle", c("id","timestamp","max"), drop = FALSE]
idle_tbl <- idle_tbl[order(idle_tbl$id, idle_tbl$timestamp), , drop = FALSE]

min_idle_between_times <- function(pid, t_prev, t_curr) {
  if (is.na(t_prev) || is.na(t_curr)) return(NA_real_)
  idle_p <- idle_tbl[idle_tbl$id == pid, , drop = FALSE]
  if (!nrow(idle_p)) return(NA_real_)
  w <- idle_p$timestamp > t_prev & idle_p$timestamp < t_curr
  if (!any(w)) return(NA_real_)
  min(idle_p$max[w], na.rm = TRUE)
}

#Initialize tier columns 
for (tt in tier_map$tier) {
  crit_aug[[paste0("case_id_", tt)]]   <- NA_character_
  crit_aug[[paste0("case_cond_", tt)]] <- NA_character_
  crit_aug[[paste0("rel_ep_", tt)]]    <- NA_integer_
  crit_aug[[paste0("ict_", tt)]] <- NA_real_
  crit_aug[[paste0("irr_", tt)]] <- NA_real_
  crit_aug[[paste0("mib_", tt)]] <- NA_real_
}

#Populate tier instances
for (pid in unique(crit_aug$id)) {
  
  d_all  <- events_aug[events_aug$id == pid, , drop = FALSE]
  d_crit <- crit_aug[crit_aug$id == pid, , drop = FALSE]
  
  glob <- which(crit_aug$id == pid)
  
  #Detect extinction onset timestamp t0 from ALL events
  ext_mask_all <- d_all$block >= 2 &
    as.character(d_all$condition) == "present" &
    (as.character(d_all$stage) == "after" | is.na(d_all$stage))
  
  ext_start_ts <- if (any(ext_mask_all)) min(d_all$timestamp[ext_mask_all], na.rm = TRUE) else Inf
  
  for (ii in seq_len(nrow(tier_map))) {
    
    tt <- tier_map$tier[ii]
    b_pre  <- tier_map$pre_block[ii]
    b_post <- tier_map$post_block[ii]
    
    #tier eligibility by group
    pid_group <- unique(as.character(d_all$group))
    pid_group <- pid_group[!is.na(pid_group)][1]
    
    max_tier_allowed <- switch(
      pid_group,
      "E8"  = 1L,
      "E16" = 2L,
      "E24" = 3L,
      "NE"  = 3L,
      3L  
    )
    
    tier_i <- as.integer(sub("^t", "", tt))
    if (!is.finite(tier_i) || tier_i > max_tier_allowed) next
    
    pre_idx <- which(d_crit$block == b_pre)
    if (!length(pre_idx)) next
    pre_idx <- pre_idx[order(d_crit$timestamp[pre_idx])]
    n_pre <- length(pre_idx)
    
    #Tier boundary condition label (block-level) based on the 2 blocks defining the tier
    cond_vals <- unique(as.character(d_all$condition[
      d_all$block %in% c(b_pre, b_post) & !is.na(d_all$condition)
    ]))
    if (!length(cond_vals)) next
    if (length(cond_vals) > 1) {
      warning(sprintf(
        "id=%s tier=%s has multiple condition labels across blocks %d/%d: %s",
        pid, tt, b_pre, b_post, paste(cond_vals, collapse = ", ")
      ))
    }
    case_cond <- cond_vals[1]
    
    #Post borrowing rule:
    #-if boundary is present: borrow all later PRESENT-condition criterion episodes
    #-if boundary is absent: borrow any later criterion episodes that occur BEFORE extinction onset (t0),
    #...even if their block-level condition is "present" (because they can still be "before").
    if (case_cond == "present") {
      post_idx <- which(d_crit$block > b_pre & as.character(d_crit$condition) == "present")
    } else {
      post_idx <- which(d_crit$block > b_pre & d_crit$timestamp < ext_start_ts)
    }
    post_idx <- post_idx[order(d_crit$timestamp[post_idx])]
    
    #Case membership
    case_id <- paste(pid, tt, sep = "__")
    idx_all <- c(pre_idx, post_idx)
    
    crit_aug[[paste0("case_id_", tt)]][glob[idx_all]]   <- case_id
    crit_aug[[paste0("case_cond_", tt)]][glob[idx_all]] <- case_cond
    
    crit_aug[[paste0("rel_ep_", tt)]][glob[pre_idx]] <- seq(-n_pre, -1)
    if (length(post_idx)) crit_aug[[paste0("rel_ep_", tt)]][glob[post_idx]] <- seq_len(length(post_idx))
    
    #compute interval metrics (minimum idle and response rate)
    #Ensure case rows are in chronological order
    idx_all <- idx_all[order(d_crit$timestamp[idx_all])]
    
    if (length(idx_all) >= 2) {
      for (pos in seq_len(length(idx_all) - 1)) {
        
        jj      <- idx_all[pos]
        jj_next <- idx_all[pos + 1]
        
        g_row  <- glob[jj]
        
        t_curr <- d_crit$timestamp[jj]
        t_next <- d_crit$timestamp[jj_next]
        
        if (is.na(t_curr) || is.na(t_next)) next
        
        ict <- t_next - t_curr
        if (!is.finite(ict) || ict <= 0) next
        
        crit_aug[[paste0("ict_", tt)]][g_row] <- ict
        crit_aug[[paste0("irr_", tt)]][g_row] <- 1 / ict
        crit_aug[[paste0("mib_", tt)]][g_row] <- min_idle_between_times(pid, t_curr, t_next)
      }
    }
    
  }
}

#INSPECTION HELPER
inspect_case <- function(id, tier = "t3") {
  cols <- c("id","group","block","stage","condition","timestamp","max","duration",
            "prev_crit_ts_global",
            paste0("case_id_", tier), paste0("case_cond_", tier), paste0("rel_ep_", tier),
            paste0("ict_", tier), paste0("irr_", tier), paste0("mib_", tier))
  crit_aug[crit_aug$id == id & !is.na(crit_aug[[paste0("rel_ep_", tier)]]), cols, drop = FALSE]
}
#example
inspect_case("01", "t1")

#BUILD CASE-EPISODE DATA FRAME IN LONG FORMAT
to_long_metric <- function(tier, metric = c("peak","duration","irr","mib")) {
  metric <- match.arg(metric)
  rel_col  <- paste0("rel_ep_", tier)
  case_col <- paste0("case_id_", tier)
  cond_col <- paste0("case_cond_", tier)
  
  keep <- !is.na(crit_aug[[rel_col]]) & !is.na(crit_aug[[case_col]])
  d <- crit_aug[keep, , drop = FALSE]
  
  val <- switch(metric,
                peak = d$max,
                duration = d$duration,
                irr = d[[paste0("irr_", tier)]],
                mib = d[[paste0("mib_", tier)]])
  
  data.frame(
    tier = tier,
    case_id = d[[case_col]],
    id = d$id,
    group = d$group,
    condition = d[[cond_col]],
    rel_ep = d[[rel_col]],
    value = val,
    stringsAsFactors = FALSE
  )
}

case_long <- rbind(
  transform(to_long_metric("t1","peak"),     metric = "Peak force"),
  transform(to_long_metric("t1","duration"), metric = "Response duration"),
  transform(to_long_metric("t1","mib"),      metric = "Minimum idle force"),
  transform(to_long_metric("t1","irr"),      metric = "Instantaneous response rate"),
  
  transform(to_long_metric("t2","peak"),     metric = "Peak force"),
  transform(to_long_metric("t2","duration"), metric = "Response duration"),
  transform(to_long_metric("t2","mib"),      metric = "Minimum idle force"),
  transform(to_long_metric("t2","irr"),      metric = "Instantaneous response rate"),
  
  transform(to_long_metric("t3","peak"),     metric = "Peak force"),
  transform(to_long_metric("t3","duration"), metric = "Response duration"),
  transform(to_long_metric("t3","mib"),      metric = "Minimum idle force"),
  transform(to_long_metric("t3","irr"),      metric = "Instantaneous response rate")
)

case_long$condition <- factor(as.character(case_long$condition), levels = c("absent","present"))
case_long$metric <- factor(case_long$metric, levels = c(
  "Peak force","Response duration","Minimum idle force","Instantaneous response rate"
))

#Deviations from anchor (rel_ep == -1), per CASE and metric
key <- paste(case_long$case_id, case_long$metric, sep="__")

anchor_vals <- tapply(seq_len(nrow(case_long)), key, function(ii) {
  vv <- case_long$value[ii]
  rr <- case_long$rel_ep[ii]
  w <- which(rr == -1 & !is.na(vv))
  if (!length(w)) return(NA_real_)
  vv[w[1]]
})

case_long$dev <- case_long$value - anchor_vals[key]

#SET COMMON POST LENGTH L (based on Peak force, present cases)
L_common <- max(case_long$rel_ep[case_long$metric == "Peak force" &
                                   case_long$condition == "present" &
                                   case_long$rel_ep > 0], na.rm = TRUE)

case_long <- case_long[case_long$rel_ep < 0 | (case_long$rel_ep > 0 & case_long$rel_ep <= L_common), , drop = FALSE]

#BOOTSTRAP CONFIDENCE OVER CASES
bootstrap_ribbon <- function(dat, B = 4000, ci_level = 0.68) {
  dat <- dat[!is.na(dat$dev) & !is.na(dat$rel_ep), , drop = FALSE]
  if (!nrow(dat)) return(data.frame())
  
  cases_u <- unique(dat$case_id)
  rel_u <- sort(unique(dat$rel_ep))
  
  mat <- matrix(NA_real_, nrow = length(cases_u), ncol = length(rel_u),
                dimnames = list(cases_u, rel_u))
  
  for (ci in seq_along(cases_u)) {
    dd <- dat[dat$case_id == cases_u[ci], , drop = FALSE]
    v <- tapply(dd$dev, dd$rel_ep, mean, na.rm = TRUE)
    cols <- match(names(v), rel_u)
    mat[ci, cols] <- as.numeric(v)
  }
  
  m <- colMeans(mat, na.rm = TRUE)
  n_cases <- colSums(!is.na(mat))
  
  alpha <- (1 - ci_level) / 2
  lo_q <- alpha
  hi_q <- 1 - alpha
  
  boot <- matrix(NA_real_, nrow = B, ncol = ncol(mat))
  pb <- txtProgressBar(min = 0, max = B, style = 3)
  for (b in 1:B) {
    idx <- sample.int(nrow(mat), size = nrow(mat), replace = TRUE)
    boot[b, ] <- colMeans(mat[idx, , drop = FALSE], na.rm = TRUE)
    if (b %% 50 == 0) setTxtProgressBar(pb, b)
  }
  close(pb)
  
  lo <- apply(boot, 2, quantile, probs = lo_q, na.rm = TRUE)
  hi <- apply(boot, 2, quantile, probs = hi_q, na.rm = TRUE)
  
  data.frame(
    rel_ep = as.integer(rel_u),
    mean = as.numeric(m),
    lo = as.numeric(lo),
    hi = as.numeric(hi),
    n_cases = as.integer(n_cases),
    stringsAsFactors = FALSE
  )
}

sum_list <- list()
k <- 0L
for (met in levels(case_long$metric)) {
  for (cond in levels(case_long$condition)) {
    dd <- case_long[case_long$metric == met & case_long$condition == cond, , drop = FALSE]
    s <- bootstrap_ribbon(dd, B = B, ci_level = ci_level)
    if (!nrow(s)) next
    s$metric <- met
    s$condition <- cond
    k <- k + 1L
    sum_list[[k]] <- s
  }
}
sum_df <- do.call(rbind, sum_list)
sum_df$metric <- factor(sum_df$metric, levels = levels(case_long$metric))
sum_df$condition <- factor(sum_df$condition, levels = levels(case_long$condition))


#CAP BY n_present 
counts_pf <- case_long[case_long$metric == "Peak force" & !is.na(case_long$dev), , drop = FALSE]

cc <- aggregate(case_id ~ condition + rel_ep, data = counts_pf,
                FUN = function(x) length(unique(x)))
names(cc)[3] <- "n_cases"

#determine L_cap 
min_present <- 10
present_cc <- cc[cc$condition == "present" & cc$rel_ep > 0, , drop = FALSE]
L_cap <- max(present_cc$rel_ep[present_cc$n_cases >= min_present], na.rm = TRUE)

if (!is.finite(L_cap)) L_cap <- 1

#cap the plotting summaries
sum_df_plot <- sum_df[sum_df$rel_ep < 0 | sum_df$rel_ep <= L_cap, , drop = FALSE]

# rebuild counts restricted to the plotted range (for clean top axis)
cc_plot <- cc[cc$rel_ep < 0 | cc$rel_ep <= L_cap, , drop = FALSE]

#−1 AND +1 ARE ADJACENT
map_x <- function(rel_ep) ifelse(rel_ep >= 1, rel_ep - 1, rel_ep)

sum_df_plot$x_plot <- map_x(sum_df_plot$rel_ep)

#breaks/labels
rel_breaks <- sort(unique(sum_df_plot$rel_ep))
x_breaks <- map_x(rel_breaks)
x_labels <- as.character(rel_breaks)

#top-axis labels "absent/present" per episode index (using Peak force counts)
n_abs <- setNames(rep(0, length(rel_breaks)), rel_breaks)
n_pre <- setNames(rep(0, length(rel_breaks)), rel_breaks)

for (r in rel_breaks) {
  a <- cc_plot$n_cases[cc_plot$rel_ep == r & cc_plot$condition == "absent"]
  p <- cc_plot$n_cases[cc_plot$rel_ep == r & cc_plot$condition == "present"]
  n_abs[as.character(r)] <- if (length(a)) a[1] else 0
  n_pre[as.character(r)] <- if (length(p)) p[1] else 0
}

top_labels <- paste0(n_abs, "/", n_pre)
names(top_labels) <- as.character(rel_breaks)

if ("0" %in% names(top_labels)) top_labels["0"] <- ""

#PLOT
p <- ggplot(sum_df_plot, aes(x = x_plot, y = mean, color = condition, fill = condition)) +
  geom_hline(yintercept = 0, linewidth = 0.6) +
  geom_vline(xintercept = -0.5, linetype = "dashed", linewidth = 0.6) +
  geom_ribbon(aes(ymin = lo, ymax = hi), alpha = 0.20, color = NA) +
  geom_line(linewidth = 0.9) +
  geom_point(size = 2.2) +
  facet_wrap(~ metric, ncol = 1, scales = "free_y") +
  scale_color_manual(values = cols) +
  scale_fill_manual(values = cols) +
  scale_x_continuous(
    breaks = x_breaks,
    labels = x_labels,
    sec.axis = dup_axis(
      name = "Aligned-episode counts [absent/present]",
      breaks = x_breaks,
      labels = top_labels[as.character(rel_breaks)]
    )
  ) +
  labs(
    x = "Episode number (relative to extinction onset)",
    y = "Deviation from reference (episode −1)",
    color = "Treatment",
    fill = "Treatment"
  ) +
  theme_classic(base_size = 13) +
  theme(
    legend.position = "top",
    strip.background = element_blank(),
    strip.text = element_text(face = "bold"),
    axis.text.x.top = element_text(size = 9)
  )

print(p)
#500x750
```
