## Supplementary material for "Evidence for the extinction burst in grip force: A preregistered study": Electronic Supplementary Materials: 6_ebf_individual_display_helper.R

```
#Plot specification for individual participant displays

#Load packages
if (!require("ggplot2")) install.packages("ggplot2"); library(ggplot2)

nonidle_spans <- function(time, state) {
  ok <- is.finite(time) & !is.na(state)
  time <- time[ok]
  st   <- tolower(trimws(as.character(state[ok])))
  
  if (!length(time)) return(data.frame(xmin=numeric(0), xmax=numeric(0), fill=character(0)))
  
  o <- order(time); time <- time[o]; st <- st[o]
  nonidle <- st != "idle"
  
  r <- rle(nonidle)
  ends <- cumsum(r$lengths)
  starts <- c(1L, head(ends, -1L) + 1L)
  
  out <- list(); k <- 0L
  for (i in seq_along(r$values)) {
    if (!isTRUE(r$values[i])) next
    idx <- starts[i]:ends[i]
    has_lag <- any(st[idx] == "lag", na.rm = TRUE)
    k <- k + 1L
    out[[k]] <- data.frame(
      xmin = min(time[idx], na.rm = TRUE),
      xmax = max(time[idx], na.rm = TRUE),
      fill = if (has_lag) "yellow" else "aquamarine3",
      stringsAsFactors = FALSE
    )
  }
  if (!length(out)) return(data.frame(xmin=numeric(0), xmax=numeric(0), fill=character(0)))
  do.call(rbind, out)
}

micro_vline_segments <- function(boundary_x, y_min, y_max) {
  if (!length(boundary_x) || !is.finite(y_min) || !is.finite(y_max) || y_max <= y_min) {
    return(data.frame(x=numeric(0), y=numeric(0), yend=numeric(0)))
  }
  
  yr <- y_max - y_min
  seg_len <- max(0.012, min(0.040, 0.030 * yr))
  gap_len <- max(0.010, min(0.035, 0.022 * yr))
  
  y_starts <- seq(y_min, y_max, by = seg_len + gap_len)
  y_ends   <- pmin(y_starts + seg_len, y_max)
  
  do.call(rbind, lapply(boundary_x, function(x) {
    data.frame(x = x, y = y_starts, yend = y_ends)
  }))
}

plot_participant_force_std <- function(all_data, file_pattern, use_block = c("block_final", "block_alt")) {
  
  use_block <- match.arg(use_block)
  
  hit <- grepl(file_pattern, all_data$file_name, ignore.case = TRUE)
  if (!any(hit)) stop("No rows matched file_pattern in all_data$file_name")
  
  df <- all_data[hit, , drop = FALSE]
  df <- df[order(df$t_abs_s), , drop = FALSE]
  
  blk0 <- suppressWarnings(as.integer(df[[use_block]]))
  
  bo0 <- tolower(trimws(as.character(df$block_original)))
  is_conf0 <- !is.na(bo0) & bo0 == "extinction_conf"
  is_expl0 <- !is.na(bo0) & bo0 == "extinction_expl"
  
  conf_blk <- suppressWarnings(as.integer(blk0[is_conf0][which(!is.na(blk0[is_conf0]))[1]]))
  blk_plot <- blk0
  if (is.finite(conf_blk) && any(is_expl0, na.rm = TRUE)) {
    blk_plot[is_expl0] <- conf_blk + 1L
  }
  df$block_plot <- blk_plot
  
  keep <- is.finite(df$block_plot) & df$block_plot >= 2L & is.finite(df$t_abs_s)
  df <- df[keep, , drop = FALSE]
  if (!nrow(df)) stop("No rows remain after dropping blocks 0/1 and NAs.")
  
  t0_shift <- suppressWarnings(min(df$t_abs_s[df$block_plot == 2L], na.rm = TRUE))
  if (!is.finite(t0_shift)) t0_shift <- min(df$t_abs_s, na.rm = TRUE)
  df$t0 <- df$t_abs_s - t0_shift
  df <- df[order(df$t0), , drop = FALSE]
  
  stg <- tolower(trimws(as.character(df$stage)))
  df$stage_plot <- ifelse(!is.na(stg) & stg %in% c("before","after"), stg, NA_character_)
  
  bo <- tolower(trimws(as.character(df$block_original)))
  blk_chr <- as.character(df$block_plot)
  
  split_bo <- split(bo, blk_chr)
  block_type <- vapply(split_bo, function(v) {
    v <- v[!is.na(v)]
    if (any(v == "extinction_conf")) return("ext_conf")
    if (any(v == "extinction_expl")) return("ext_expl")
    "other"
  }, character(1))
  
  df$trace_cat <- unname(block_type[blk_chr])
  df$trace_run <- cumsum(c(TRUE, df$trace_cat[-1] != df$trace_cat[-nrow(df)]))
  df$block_run <- cumsum(c(TRUE, df$block_plot[-1] != df$block_plot[-nrow(df)]))
  
  y_max <- suppressWarnings(max(df$force_std, na.rm = TRUE))
  y_min <- suppressWarnings(min(df$force_std, na.rm = TRUE))
  if (!is.finite(y_min)) y_min <- 0
  if (!is.finite(y_max)) y_max <- 1
  y_lim_top <- y_max + 0.1
  
  y_range_total <- y_lim_top - y_min
  band_frac <- 0.07
  top_margin_frac <- 0.01
  band_thickness <- band_frac * y_range_total
  band_top_margin <- top_margin_frac * y_range_total
  band_ymax <- y_lim_top - band_top_margin
  band_ymin <- band_ymax - band_thickness
  
  band_base <- data.frame(
    xmin = min(df$t0, na.rm = TRUE),
    xmax = max(df$t0, na.rm = TRUE),
    ymin = band_ymin,
    ymax = band_ymax,
    fill = "black",
    stringsAsFactors = FALSE
  )
  band_nonidle <- nonidle_spans(df$t0, df$state)
  if (nrow(band_nonidle)) {
    band_nonidle$ymin <- band_ymin
    band_nonidle$ymax <- band_ymax
  }
  
  run_ids <- sort(unique(df$block_run))
  run_df <- do.call(rbind, lapply(run_ids, function(rid) {
    ii <- df$block_run == rid
    data.frame(
      block_run  = rid,
      xmin       = min(df$t0[ii], na.rm = TRUE),
      xmax       = max(df$t0[ii], na.rm = TRUE),
      stage_plot = df$stage_plot[which(ii)[1]],
      run_max    = suppressWarnings(max(df$force_std[ii], na.rm = TRUE)),
      stringsAsFactors = FALSE
    )
  }))
  
  run_df$stage_fill <- NA_character_
  run_df$stage_fill[run_df$stage_plot == "after"]  <- "antiquewhite"
  run_df$stage_fill[run_df$stage_plot == "before"] <- "#B7C9E2"
  
  stage_rects <- run_df[!is.na(run_df$stage_fill), , drop = FALSE]
  stage_rects <- data.frame(
    xmin = stage_rects$xmin, xmax = stage_rects$xmax,
    ymin = -Inf, ymax = Inf,
    fill = stage_rects$stage_fill,
    stringsAsFactors = FALSE
  )
  
  orange_rects <- run_df[is.finite(run_df$run_max), , drop = FALSE]
  orange_rects$ymin <- orange_rects$run_max
  orange_rects$ymax <- y_max
  orange_rects$fill <- "#FE5000"
  orange_rects <- orange_rects[is.finite(orange_rects$ymin) & orange_rects$ymax > orange_rects$ymin, , drop = FALSE]
  
  boundary_x <- run_df$xmin[order(run_df$xmin)]
  if (length(boundary_x) > 1) boundary_x <- boundary_x[-1] else boundary_x <- numeric(0)
  vseg_df <- micro_vline_segments(boundary_x, y_min, y_lim_top)
  
  hline_df <- data.frame(y = c(0.3, 0.7))
  
  fn <- unique(df$file_name); fn <- fn[which(!is.na(fn))[1]]
  lead_digits <- regmatches(fn, regexpr("^\\d+", fn))
  pid <- if (length(lead_digits) && nchar(lead_digits)) substr(lead_digits, 1, 2) else substr(fn, 1, 2)
  grp <- unique(df$group); grp <- grp[which(!is.na(grp))[1]]
  plot_title <- paste0("Participant ", pid, ", Group ", grp)
  
  x_min <- min(df$t0, na.rm = TRUE)
  x_max <- max(df$t0, na.rm = TRUE)
  title_x <- x_min + 0.01 * (x_max - x_min)
  title_y <- band_ymax - 0.15 * (band_ymax - band_ymin)
  
  p <- ggplot() +
    { if (nrow(stage_rects))
      geom_rect(
        data = stage_rects,
        aes(xmin = xmin, xmax = xmax, ymin = ymin, ymax = ymax, fill = fill),
        alpha = 0.8, inherit.aes = FALSE
      ) else NULL } +
    { if (nrow(orange_rects))
      geom_rect(
        data = orange_rects,
        aes(xmin = xmin, xmax = xmax, ymin = ymin, ymax = ymax, fill = fill),
        alpha = 0.5, inherit.aes = FALSE
      ) else NULL } +
    geom_rect(
      data = band_base,
      aes(xmin = xmin, xmax = xmax, ymin = ymin, ymax = ymax, fill = fill),
      alpha = 0.95, inherit.aes = FALSE
    ) +
    { if (nrow(band_nonidle))
      geom_rect(
        data = band_nonidle,
        aes(xmin = xmin, xmax = xmax, ymin = ymin, ymax = ymax, fill = fill),
        alpha = 0.95, inherit.aes = FALSE
      ) else NULL } +
    { if (nrow(vseg_df))
      geom_segment(
        data = vseg_df,
        aes(x = x, xend = x, y = y, yend = yend),
        linewidth = 0.22, alpha = 0.60, inherit.aes = FALSE
      ) else NULL } +
    geom_hline(
      data = hline_df,
      aes(yintercept = y),
      linetype = "dashed", linewidth = 0.35
    ) +
    geom_line(
      data = df,
      aes(x = t0, y = force_std),
      linewidth = 0.3, alpha = 0.3, color = "black"
    ) +
    geom_line(
      data = df,
      aes(x = t0, y = force_std, color = trace_cat, group = trace_run),
      linewidth = 0.4, alpha = 0.9
    ) +
    annotate(
      "text",
      x = title_x, y = title_y,
      label = plot_title,
      hjust = 0, vjust = 0.5,
      color = "gray80",
      size = 2.6
    ) +
    scale_color_manual(values = c(
      ext_conf = "#8B008B",
      ext_expl = "gray55",
      other    = "#228B22"
    )) +
    scale_fill_identity() +
    coord_cartesian(ylim = c(y_min, y_lim_top), expand = FALSE) +
    labs(x = NULL, y = NULL, title = NULL) +
    theme_classic(base_size = 14) +
    theme(
      legend.position = "none",
      plot.title = element_blank(),
      axis.title.x = element_blank(),
      axis.title.y = element_blank(),
      axis.text.x = element_text(size = 2.5),
      axis.text.y = element_blank(),
      axis.ticks  = element_line(linewidth = 0.25),
      axis.ticks.y = element_blank(),
      axis.ticks.length = grid::unit(1.5, "pt"),
      plot.margin = margin(1, 1, 1, 1, "pt")
    )
  
  attr(p, "x_span") <- diff(range(df$t0, na.rm = TRUE))
  p
}


compute_widths_in <- function(plots, sec_per_in,
                              left_right_overhead_in = 0.55,
                              min_total_width_in = 0.90) {
  
  spans <- vapply(plots, function(p) {
    s <- attr(p, "x_span")
    if (is.null(s) || !is.finite(s) || s < 0) {
      xr <- get_x_range(p)
      s <- diff(xr)
      if (!is.finite(s) || s < 0) s <- 0
    }
    s
  }, numeric(1))
  
  w <- left_right_overhead_in + spans / sec_per_in
  pmax(w, min_total_width_in)
}


pack_rows_bfd <- function(widths, max_w) {
  n <- length(widths)
  if (!n) return(list(rows = list(), rem = numeric(0)))
  
  idx <- order(widths, decreasing = TRUE)
  rows <- list()
  rem  <- numeric(0)
  
  for (i in idx) {
    w <- widths[i]
    if (!is.finite(w)) next
    if (w > max_w) return(NULL)
    
    can <- which(rem >= w)
    if (length(can)) {
      j <- can[which.min(rem[can] - w)]
      rows[[j]] <- c(rows[[j]], i)
      rem[j] <- rem[j] - w
    } else {
      rows[[length(rows) + 1L]] <- i
      rem <- c(rem, max_w - w)
    }
  }
  
  rows <- lapply(rows, function(ii) ii[order(widths[ii], decreasing = TRUE)])
  
  list(rows = rows, rem = rem)
}

layout_for_scale_autoH <- function(plots, sec_per_in,
                                   page_w_in, page_h_in,
                                   margins_in = c(left=0.35, right=0.15, top=0.25, bottom=0.25),
                                   row_gap_in = 0.04,
                                   left_right_overhead_in = 1.25,
                                   min_total_width_in = 1.6) {
  
  inner_w <- page_w_in - margins_in["left"] - margins_in["right"]
  inner_h <- page_h_in - margins_in["top"]  - margins_in["bottom"]
  
  w_in <- compute_widths_in(
    plots, sec_per_in,
    left_right_overhead_in = left_right_overhead_in,
    min_total_width_in = min_total_width_in
  )
  
  pack <- pack_rows_bfd(w_in, inner_w)
  if (is.null(pack)) {
    return(list(fits = FALSE, sec_per_in = sec_per_in, widths_in = w_in,
                rows = list(), n_rows = Inf, row_h_in = NA_real_,
                inner_w = inner_w, inner_h = inner_h,
                margins_in = margins_in, row_gap_in = row_gap_in,
                page_w_in = page_w_in, page_h_in = page_h_in))
  }
  
  n_rows <- length(pack$rows)
  row_h <- (inner_h - (n_rows - 1) * row_gap_in) / n_rows
  
  list(
    fits       = is.finite(row_h) && row_h > 0,
    sec_per_in = sec_per_in,
    widths_in  = w_in,
    rows       = pack$rows,
    rem_in     = pack$rem,
    n_rows     = n_rows,
    row_h_in   = row_h,
    inner_w    = inner_w,
    inner_h    = inner_h,
    margins_in = margins_in,
    row_gap_in = row_gap_in,
    page_w_in  = page_w_in,
    page_h_in  = page_h_in
  )
}

optimize_pack_one_page <- function(plots,
                                   page_w_in = 11, page_h_in = 8.5,
                                   margins_in = c(left=0.35, right=0.15, top=0.25, bottom=0.25),
                                   row_gap_in = 0.04,
                                   left_right_overhead_in = 1.25,
                                   min_total_width_in = 1.6,
                                   min_row_h_in = 0.45,
                                   sec_per_in_lo = 0.2, sec_per_in_hi = 2000,
                                   iters = 40) {
  
  hi <- layout_for_scale_autoH(plots, sec_per_in_hi, page_w_in, page_h_in,
                               margins_in, row_gap_in,
                               left_right_overhead_in, min_total_width_in)
  if (!hi$fits || hi$row_h_in < min_row_h_in) {
    stop("Cannot fit with the current min_row_h_in. Lower min_row_h_in, row_gap_in, margins, or increase page size.")
  }
  
  lo <- layout_for_scale_autoH(plots, sec_per_in_lo, page_w_in, page_h_in,
                               margins_in, row_gap_in,
                               left_right_overhead_in, min_total_width_in)
  
  if (lo$fits && lo$row_h_in >= min_row_h_in) return(lo)
  
  a <- sec_per_in_lo
  b <- sec_per_in_hi
  best <- hi
  
  for (k in seq_len(iters)) {
    mid <- (a + b) / 2
    m <- layout_for_scale_autoH(plots, mid, page_w_in, page_h_in,
                                margins_in, row_gap_in,
                                left_right_overhead_in, min_total_width_in)
    ok <- m$fits && is.finite(m$row_h_in) && m$row_h_in >= min_row_h_in
    if (ok) {
      best <- m
      b <- mid
    } else {
      a <- mid
    }
  }
  
  best
}

layout_placement_df <- function(lay, plot_names = NULL) {
  if (is.null(plot_names)) plot_names <- as.character(seq_along(lay$widths_in))
  out <- list(); k <- 0L
  for (r in seq_along(lay$rows)) {
    ii <- lay$rows[[r]]
    for (c in seq_along(ii)) {
      i <- ii[c]
      k <- k + 1L
      out[[k]] <- data.frame(
        row = r, col = c,
        plot_index = i,
        plot_name  = plot_names[i],
        width_in   = lay$widths_in[i],
        stringsAsFactors = FALSE
      )
    }
  }
  do.call(rbind, out)
}

render_pack_pdf <- function(plots, lay, out_pdf = "participants_optimized.pdf") {
  pdf(out_pdf, width = lay$page_w_in, height = lay$page_h_in, onefile = TRUE)
  on.exit(dev.off(), add = TRUE)
  
  grid.newpage()
  
  left <- lay$margins_in["left"]
  top  <- lay$margins_in["top"]
  gap  <- lay$row_gap_in
  rh   <- lay$row_h_in
  page_h <- lay$page_h_in
  
  for (r in seq_along(lay$rows)) {
    ii <- lay$rows[[r]]
    w_in <- lay$widths_in[ii]
    
    x_lefts <- left + c(0, cumsum(w_in))[seq_along(w_in)]
    y_top   <- page_h - top - (r - 1) * (rh + gap)
    y_c     <- y_top - rh / 2
    
    for (k in seq_along(ii)) {
      p <- plots[[ ii[k] ]]
      g <- ggplotGrob(p)
      
      vp <- viewport(
        x = unit(x_lefts[k] + w_in[k] / 2, "in"),
        y = unit(y_c, "in"),
        width  = unit(w_in[k], "in"),
        height = unit(rh, "in"),
        just = c("center","center")
      )
      pushViewport(vp); grid.draw(g); popViewport()
    }
  }
  
  invisible(out_pdf)
}

#build plots
files <- sort(unique(all_data$file_name))
files <- files[!grepl("^40_timeseries_all\\.csv$", files, ignore.case = TRUE)]

plots <- setNames(lapply(files, function(fn) {
 pat <- sub("\\.csv$", "", fn, ignore.case = TRUE)
 plot_participant_force_std(all_data, pat, use_block = "block_final")
}), files)

files <- sort(unique(all_data$file_name))
files <- files[!grepl("^40_timeseries_all\\.csv$", files, ignore.case = TRUE)]

plots <- setNames(lapply(files, function(fn) {
 pat <- sub("\\.csv$", "", fn, ignore.case = TRUE)
 plot_participant_force_std(all_data, pat, use_block = "block_final")
}), files)
```
