## Supplementary material for "Evidence for the extinction burst in grip force: A preregistered study": Electronic Supplementary Materials: 7_ebf_collective_viz_optimization_n_rendering.R

```
#One-page collective visualization of individual performance

#Load packages
if (!require("ggplot2")) install.packages("ggplot2"); library(ggplot2)
if (!require("grid"))   library(grid)

#Get plot time span 
plot_span_sec <- function(p) {
  s <- attr(p, "x_span", exact = TRUE)
  if (is.finite(s) && length(s) == 1) return(as.numeric(s))
  
  b <- ggplot_build(p)
  xd <- unlist(lapply(b$data, function(d) if ("x" %in% names(d)) d$x else NULL), use.names = FALSE)
  xd <- xd[is.finite(xd)]
  if (!length(xd)) return(0)
  diff(range(xd))
}

plot_fixed_width_in <- function(p) {
  g <- ggplotGrob(p)
  
  pan <- grepl("^panel", g$layout$name)
  if (!any(pan)) {
    return(convertWidth(sum(g$widths), "in", valueOnly = TRUE))
  }
  
  panel_cols <- unique(unlist(Map(seq, g$layout$l[pan], g$layout$r[pan])))
  total_in   <- convertWidth(sum(g$widths), "in", valueOnly = TRUE)
  panel_in   <- convertWidth(sum(g$widths[panel_cols]), "in", valueOnly = TRUE)
  
  fixed_in <- total_in - panel_in
  if (!is.finite(fixed_in) || fixed_in < 0) fixed_in <- 0
  fixed_in
}

compute_widths_in <- function(plots, sec_per_in, min_panel_in = 0.20) {
  spans <- vapply(plots, plot_span_sec, numeric(1))
  fixed <- vapply(plots, plot_fixed_width_in, numeric(1))
  
  spans[!is.finite(spans) | spans < 0] <- 0
  fixed[!is.finite(fixed) | fixed < 0] <- 0
  
  panel_w <- pmax(min_panel_in, spans / sec_per_in)
  fixed + panel_w
}

pack_rows_bfd <- function(widths, max_w) {
  n <- length(widths)
  if (!n) return(list(rows = list(), rem = numeric(0)))
  
  idx <- order(widths, decreasing = TRUE)
  rows <- list()
  rem  <- numeric(0)
  
  for (i in idx) {
    w <- widths[i]
    if (!is.finite(w)) next
    if (w > max_w) return(NULL)
    
    can <- which(rem >= w)
    if (length(can)) {
      j <- can[which.min(rem[can] - w)]
      rows[[j]] <- c(rows[[j]], i)
      rem[j] <- rem[j] - w
    } else {
      rows[[length(rows) + 1L]] <- i
      rem <- c(rem, max_w - w)
    }
  }
  
  rows <- lapply(rows, function(ii) ii[order(widths[ii], decreasing = TRUE)])
  list(rows = rows, rem = rem)
}

layout_for_scale_autoH <- function(plots, sec_per_in,
                                   page_w_in, page_h_in,
                                   margins_in = c(left=0.35, right=0.15, top=0.25, bottom=0.25),
                                   row_gap_in = 0.04,
                                   min_panel_in = 0.20) {
  
  inner_w <- page_w_in - margins_in["left"] - margins_in["right"]
  inner_h <- page_h_in - margins_in["top"]  - margins_in["bottom"]
  
  w_in <- compute_widths_in(plots, sec_per_in, min_panel_in = min_panel_in)
  
  pack <- pack_rows_bfd(w_in, inner_w)
  if (is.null(pack)) {
    return(list(fits = FALSE, sec_per_in = sec_per_in, widths_in = w_in,
                rows = list(), n_rows = Inf, row_h_in = NA_real_,
                inner_w = inner_w, inner_h = inner_h,
                margins_in = margins_in, row_gap_in = row_gap_in,
                page_w_in = page_w_in, page_h_in = page_h_in))
  }
  
  n_rows <- length(pack$rows)
  row_h <- (inner_h - (n_rows - 1) * row_gap_in) / n_rows
  
  list(
    fits       = is.finite(row_h) && row_h > 0,
    sec_per_in = sec_per_in,
    widths_in  = w_in,
    rows       = pack$rows,
    rem_in     = pack$rem,
    n_rows     = n_rows,
    row_h_in   = row_h,
    inner_w    = inner_w,
    inner_h    = inner_h,
    margins_in = margins_in,
    row_gap_in = row_gap_in,
    page_w_in  = page_w_in,
    page_h_in  = page_h_in
  )
}

optimize_pack_one_page <- function(plots,
                                   page_w_in = 11, page_h_in = 8.5,
                                   margins_in = c(left=0.35, right=0.15, top=0.25, bottom=0.25),
                                   row_gap_in = 0.04,
                                   min_panel_in = 0.20,
                                   min_row_h_in = 0.45,
                                   sec_per_in_lo = 0.2, sec_per_in_hi = 2000,
                                   iters = 40) {
  
  hi <- layout_for_scale_autoH(plots, sec_per_in_hi, page_w_in, page_h_in,
                               margins_in, row_gap_in, min_panel_in)
  if (!hi$fits || hi$row_h_in < min_row_h_in) {
    stop("Cannot fit with current min_row_h_in. Lower min_row_h_in/row_gap/margins or increase page size.")
  }
  
  lo <- layout_for_scale_autoH(plots, sec_per_in_lo, page_w_in, page_h_in,
                               margins_in, row_gap_in, min_panel_in)
  if (lo$fits && lo$row_h_in >= min_row_h_in) return(lo)
  
  a <- sec_per_in_lo
  b <- sec_per_in_hi
  best <- hi
  
  for (k in seq_len(iters)) {
    mid <- (a + b) / 2
    m <- layout_for_scale_autoH(plots, mid, page_w_in, page_h_in,
                                margins_in, row_gap_in, min_panel_in)
    ok <- m$fits && is.finite(m$row_h_in) && m$row_h_in >= min_row_h_in
    if (ok) { best <- m; b <- mid } else { a <- mid }
  }
  
  best
}

layout_placement_df <- function(lay, plot_names = NULL) {
  if (is.null(plot_names)) plot_names <- as.character(seq_along(lay$widths_in))
  out <- list(); k <- 0L
  for (r in seq_along(lay$rows)) {
    ii <- lay$rows[[r]]
    for (c in seq_along(ii)) {
      i <- ii[c]
      k <- k + 1L
      out[[k]] <- data.frame(
        row = r, col = c,
        plot_index = i,
        plot_name  = plot_names[i],
        width_in   = lay$widths_in[i],
        stringsAsFactors = FALSE
      )
    }
  }
  do.call(rbind, out)
}

render_pack_pdf <- function(plots, lay, out_pdf = "participants_optimized.pdf") {
  pdf(out_pdf, width = lay$page_w_in, height = lay$page_h_in, onefile = TRUE)
  on.exit(dev.off(), add = TRUE)
  
  grid.newpage()
  
  left   <- lay$margins_in["left"]
  top    <- lay$margins_in["top"]
  gap    <- lay$row_gap_in
  rh     <- lay$row_h_in
  page_h <- lay$page_h_in
  inner_w <- lay$inner_w
  
  for (r in seq_along(lay$rows)) {
    ii <- lay$rows[[r]]
    w_in <- lay$widths_in[ii]
    
    row_total <- sum(w_in)
    start_x <- left + (inner_w - row_total) / 2
    
    x_lefts <- start_x + c(0, cumsum(w_in))[seq_along(w_in)]
    y_top   <- page_h - top - (r - 1) * (rh + gap)
    y_c     <- y_top - rh / 2
    
    for (k in seq_along(ii)) {
      p <- plots[[ ii[k] ]]
      g <- ggplotGrob(p)
      
      vp <- viewport(
        x = unit(x_lefts[k] + w_in[k] / 2, "in"),
        y = unit(y_c, "in"),
        width  = unit(w_in[k], "in"),
        height = unit(rh, "in"),
        just = c("center","center")
      )
      pushViewport(vp); grid.draw(g); popViewport()
    }
  }
  
  invisible(out_pdf)
}

#Implementation

page_w_in <- 7
page_h_in <- 9

lay <- optimize_pack_one_page(
  plots,
  page_w_in = page_w_in,
  page_h_in = page_h_in,
  min_row_h_in = 0.45,
  min_panel_in = 0.20
)

cat("Chosen sec_per_in:", lay$sec_per_in, "\n")
cat("Rows:", lay$n_rows, "\n")
cat("Row height (in):", round(lay$row_h_in, 3), "\n")

placement <- layout_placement_df(lay, plot_names = names(plots))
print(placement[order(placement$row, placement$col), ], row.names = FALSE)

render_pack_pdf(plots, lay, out_pdf = "participants_optimized.pdf")
```
