## Supplementary material for "Evidence for the extinction burst in grip force: A preregistered study": Electronic Supplementary Materials: 8_ebf_continuous_force.R

```
# Continuous force trajectory analysis
#
# OVERVIEW:
#   The pre-extinction ("before") block is mapped to x = -1 to x = 0.
#   The post-transition block (early extinction or next reward block) extends
#   from x = 0 onward. This normalization makes every transition comparable
#   regardless of block duration in absolute time.
#
#   Sliding windows of width 0.25 (i.e., 1/4 of the pre-transition block) are computed
#   across the x axis. Each window mean is placed at its right edge. Force is
#   re-standardized within each transition so that the last pre-transition window
#   equals zero; this removes between-transition level differences and makes
#   the y-axis interpretable as deviation from the individual pre-transition
#   baseline.

#Load required packages
if (!require("ggplot2"))  install.packages("ggplot2");  library(ggplot2)
if (!require("dplyr"))    install.packages("dplyr");    library(dplyr)
if (!require("tidyr"))    install.packages("tidyr");    library(tidyr)
if (!require("ggridges")) install.packages("ggridges"); library(ggridges)

set.seed(1401)

N_BOOT    <- 1000   #bootstrap iterations
CI_LEVEL  <- 0.68   #CI width; 68% approximates ±1 SEM under normality
bin_width <- 0.01   #x-axis steps between consecutive window placements
#Note that, unlike typical bins, windows heavily overlap by design,
#...producing a smooth trajectory.
WINDOW_W  <- 0.25   #window width in normalized x-units.
#again, 0.25 means each window spans one quarter of the
#pre-transition block duration. Choice is reflects the aim to balance
#smoothness against temporal resolution.

#Important:
#Only tier-boundary "before" blocks are used as pre-transition anchors.
#Block 1 (warm-up) is excluded
ANCHOR_BLOCKS <- c(2L, 4L, 6L)
WARMUP_BLOCK  <- 1L

#Sampling rate detection
#Computes the median inter-sample interval (step_size_s) from consecutive
#timestamps within each participant file. 
#Documents the data's temporal resolution as requested by a reviewer.

resolution_raw <- all_data %>%
  arrange(file_name, t_abs_s) %>%
  group_by(file_name) %>%
  mutate(dt_ms = c(NA_real_, diff(t_abs_s)) * 1000) %>%
  filter(!is.na(dt_ms)) %>%
  ungroup() %>%
  mutate(pid = sub("^(\\d{2}).*", "P\\1", file_name))

resolution_check <- resolution_raw %>%
  group_by(file_name, pid) %>%
  summarise(median_dt_ms = median(dt_ms, na.rm = TRUE), .groups = "drop")

step_size_s <- median(resolution_check$median_dt_ms, na.rm = TRUE) / 1000
cat(sprintf("Raw logging resolution: %.4f s (%.2f ms)\n",
            step_size_s, step_size_s * 1000))
cat(sprintf("  Implied logging rate: ~%.1f Hz\n", 1 / step_size_s))

#Deduplication of oversampled force readings
#
#The dynamometer logs at ~120 Hz but the underlying force reading only
#updates at ~10 Hz, so each true reading is written to ~12 consecutive
#rows with byte-identical force_std values (confirmed empirically:
#100% exact match within runs, modal run length = 12, within-run
#inter-sample interval ~8.3 ms).
#
#We do NOT collapse every run of identical consecutive values to a
#single row. Some runs span 2-4 true reading periods (force genuinely
#unchanged across consecutive ~10 Hz cycles, not just oversampling of
#one reading) -- naive collapsing would wrongly flatten a real
#multi-cycle plateau into one point, and would silently down-weight
#stable periods when window means are later computed. Instead, each
#run is assigned an estimated number of true readings based on its
#duration relative to the empirical true period, and that many
#evenly-spaced representative rows are kept.

modal_run_len  <- 12  #confirmed empirically; see duplication characterization
true_period_s  <- step_size_s * modal_run_len
cat(sprintf("True update period: %.4f s (%.2f ms)\n",
            true_period_s, true_period_s * 1000))
cat(sprintf("  Implied true update rate: ~%.1f Hz\n", 1 / true_period_s))

dedup_dynamometer <- function(df, true_period_s) {
  df <- df %>% arrange(t_abs_s)
  if (nrow(df) < 2) return(df)

  run_id <- cumsum(c(TRUE, diff(df$force_std) != 0))
  df$.run_id <- run_id

  df %>%
    group_by(.run_id) %>%
    group_modify(function(run_df, key) {
      run_dur <- max(run_df$t_abs_s) - min(run_df$t_abs_s)
      n_true  <- max(1L, round(run_dur / true_period_s))
      n_true  <- min(n_true, nrow(run_df))  #never invent more rows than exist
      idx     <- unique(round(seq(1, nrow(run_df), length.out = n_true)))
      run_df[idx, ]
    }) %>%
    ungroup() %>%
    select(-.run_id)
}

cat("\n=== DEDUPLICATING RAW FORCE READINGS ===\n")
n_before <- nrow(all_data)
all_data <- all_data %>%
  group_by(file_name) %>%
  group_modify(~ dedup_dynamometer(.x, true_period_s = true_period_s)) %>%
  ungroup()
n_after <- nrow(all_data)
cat(sprintf("Rows before dedup: %d\n", n_before))
cat(sprintf("Rows after dedup:  %d (%.1f%% of raw)\n",
            n_after, 100 * n_after / n_before))

#Sliding window extractor

#Takes the raw samples for one before block and the
#immediately following block, maps them onto the normalized x-axis, and
#returns a data frame of window means across that axis.

#Windowing:
#Windows are placed at right-edge positions from (-1 + WINDOW_W) to
#x_limit, stepping by bin_width. Each window collects all samples
#whose x_norm falls within [right_edge - WINDOW_W, right_edge].

#y axis re-standarization:
#The anchor window is the last window whose right edge <= 0 AND contains only pre-transition samples.
#Its mean is subtracted from all window means, so the trace crosses
#y = 0 exactly at the last pre-onset window. 

extract_transition <- function(before_rows, after_rows,
                               transition_type, transition_id,
                               window_w = WINDOW_W,
                               x_limit  = 2) {
  
  before_rows <- before_rows %>% arrange(t_abs_s)
  after_rows  <- after_rows  %>% arrange(t_abs_s)
  
  #Require at least two before-block samples (needed to define a duration)
  #and at least one after-block sample.
  if (nrow(before_rows) < 2 || nrow(after_rows) == 0) return(NULL)
  
  #Time landmarks for x-normalization
  t_x_minus1     <- min(before_rows$t_abs_s, na.rm = TRUE)  #gets x = -1
  t_x_zero       <- min(after_rows$t_abs_s,  na.rm = TRUE)  #gets x =  0
  pre_duration_s <- t_x_zero - t_x_minus1                   #1 normalized unit
  
  if (!is.finite(pre_duration_s) || pre_duration_s <= 0) return(NULL)
  
  #Apply x-normalization to all samples from both blocks combined
  work_rows <- bind_rows(before_rows, after_rows) %>%
    arrange(t_abs_s) %>%
    mutate(x_norm_raw = (t_abs_s - t_x_zero) / pre_duration_s)
  
  #Define right-edge positions for all windows across the x-axis
  right_edges <- seq(-1 + window_w, x_limit, by = bin_width)
  
  #Compute mean force within each window
  window_means <- vapply(right_edges, function(re) {
    le     <- re - window_w
    in_win <- work_rows$force_std[work_rows$x_norm_raw >= le &
                                    work_rows$x_norm_raw <= re]
    if (!length(in_win)) return(NA_real_)
    mean(in_win, na.rm = TRUE)
  }, numeric(1))
  
  result <- data.frame(
    transition_id   = transition_id,
    transition_type = transition_type,
    x_norm          = right_edges,
    window_mean     = window_means,
    pre_duration_s  = pre_duration_s,  
    t_x_minus1      = t_x_minus1,
    t_x_zero        = t_x_zero,
    stringsAsFactors = FALSE
  )
  
  result <- result %>% filter(!is.na(window_mean))
  if (!nrow(result)) return(NULL)
  
  #Identify the y anchor: last window that falls entirely within the
  #before block
  anchor_wins  <- result %>% filter(x_norm <= 0, x_norm - window_w >= -1)
  if (!nrow(anchor_wins)) return(NULL)
  last_pre_val <- anchor_wins$window_mean[which.max(anchor_wins$x_norm)]
  
  #Subtract the anchor value from all windows (baseline correction)
  result %>%
    mutate(force_renorm = window_mean - last_pre_val) %>%
    filter(x_norm <= x_limit)
}

#Collect transitions
#For each tier-boundary before block,
#if the next block is extinction_conf (and participant is not NE),
#the transition is classified as extinction and the after block
#includes both extinction_conf (i.e., early extinction) and extinction_expl 
#(i.e., late extinction), so that we capture full extinction
#exposure and x_limit = 2;
#...and if the next block is a reward block, the transition
#is classified as "reward-reward" and the after block is the next
#reward tier only (x_limit = 1, matched to the extinction pre-period).

all_transitions <- list()
k <- 0L

participants <- sort(unique(all_data$file_name))
participants <- participants[!grepl("^40_timeseries_all", participants,
                                    ignore.case = TRUE)]

for (fn in participants) {
  df        <- all_data %>% filter(file_name == fn) %>% arrange(t_abs_s)
  grp       <- df$group[which(!is.na(df$group))[1]]
  grp_upper <- toupper(trimws(as.character(grp)))
  
  df <- df %>%
    mutate(
      stage_lc = tolower(trimws(as.character(stage))),
      bo_lc    = tolower(trimws(as.character(block_original))),
      bf_int   = suppressWarnings(as.integer(block_final))
    )
  
  #excludes warm-up and NA
  bf_ordered <- df %>%
    filter(!is.na(bf_int), bf_int != WARMUP_BLOCK) %>%
    arrange(t_abs_s) %>%
    pull(bf_int) %>%
    unique()
  
  for (i in seq_along(bf_ordered)) {
    bf <- bf_ordered[i]
    
    #Only tier-boundary before blocks are used as anchors
    if (!bf %in% ANCHOR_BLOCKS) next
    
    before_rows <- df %>% filter(bf_int == bf, stage_lc == "before")
    if (nrow(before_rows) < 2) next
    
    t_before_end <- max(before_rows$t_abs_s, na.rm = TRUE)
    
    if (i == length(bf_ordered)) next
    next_bf <- bf_ordered[i + 1]
    
    next_block_rows <- df %>%
      filter(bf_int == next_bf, t_abs_s > t_before_end)
    if (nrow(next_block_rows) == 0) next
    
    next_bo <- tolower(trimws(as.character(next_block_rows$bo_lc[1])))
    
    if (next_bo == "extinction_conf" && !grepl("NE", grp_upper)) {
      #identifies extinction transition
      #After block = both extinction phases (early + late), so x extends to 2
      after_rows <- df %>%
        filter(bo_lc %in% c("extinction_conf", "extinction_expl"),
               t_abs_s > t_before_end)
      k <- k + 1L
      result <- extract_transition(
        before_rows     = before_rows,
        after_rows      = after_rows,
        transition_type = "extinction",
        transition_id   = paste0(fn, "_ext_tier", bf),
        x_limit         = 2
      )
      if (!is.null(result)) {
        result$file_name <- fn
        result$group     <- grp
        all_transitions[[k]] <- result
      }
      
    } else if (next_bo != "extinction_conf" && next_bo != "extinction_expl") {
      #identifies reward-reward transition
      #After block = next reward tier only; x_limit = 1
      after_rows <- df %>%
        filter(bf_int == next_bf, t_abs_s > t_before_end)
      k <- k + 1L
      result <- extract_transition(
        before_rows     = before_rows,
        after_rows      = after_rows,
        transition_type = "reward-reward",
        transition_id   = paste0(fn, "_rr_tier", bf),
        x_limit         = 1
      )
      if (!is.null(result)) {
        result$file_name <- fn
        result$group     <- grp
        all_transitions[[k]] <- result
      }
    }
  }
}

trans_df <- bind_rows(all_transitions) %>%
  mutate(pid = sub("^(\\d{2}).*", "P\\1", file_name))

cat(sprintf("Total transitions: %d\n", length(unique(trans_df$transition_id))))
cat(sprintf("  Reward-Extinction:    %d\n",
            length(unique(trans_df$transition_id[trans_df$transition_type == "extinction"]))))
cat(sprintf("  Reward-Reward: %d\n",
            length(unique(trans_df$transition_id[trans_df$transition_type == "reward-reward"]))))
cat(sprintf("Participants with data: %d\n\n", length(unique(trans_df$file_name))))

cat("=== TRANSITION COUNT PER PARTICIPANT ===\n")
trans_summary <- trans_df %>%
  group_by(pid, group, transition_type) %>%
  summarise(n = n_distinct(transition_id), .groups = "drop") %>%
  pivot_wider(names_from = transition_type, values_from = n, values_fill = 0L)
print(trans_summary)

#Verify that x ranges match expectations:
#extinction transitions should span x = -0.75 to 2
#...whereas reward-reward transitions should span x = -0.75 to 1
cat("\n=== X RANGE CHECK ===\n")
trans_df %>%
  group_by(transition_type) %>%
  summarise(x_min = round(min(x_norm), 3),
            x_max = round(max(x_norm), 3),
            .groups = "drop") %>%
  print()
#All is good :)

#Bootstrapping
#Resampling is done at the transition level rather than the participant level.
#The transition event is the natural unit
#of observation here i.e., what varies across the y-axis at any given x_bin is
#how different transitions behaved at that point in normalized time, and that
#is what the CI should reflect. Each bootstrap iteration therefore draws
#transitions (48 R-E and 45 R-R) with replacement, computes bin means across the resampled set,
#and stores them. CIs are percentile-based.

cat("\n=== BOOTSTRAPPING ===\n")
#Reward-reward source: NE group only
rr_source <- trans_df %>%
  filter(transition_type == "reward-reward",
         toupper(trimws(group)) == "NE")

cat(sprintf("Reward-reward transitions used (NE only): %d\n",
            length(unique(rr_source$transition_id))))

#Rounds x_norm to bin_width grid
trans_df_binned  <- trans_df  %>%
  mutate(x_bin = round(x_norm / bin_width) * bin_width)
rr_source_binned <- rr_source %>%
  mutate(x_bin = round(x_norm / bin_width) * bin_width)

boot_ci <- function(df_type, n_boot = N_BOOT, ci = CI_LEVEL, label = "") {
  trans_ids <- unique(df_type$transition_id)
  x_bins    <- sort(unique(df_type$x_bin))
  alpha     <- (1 - ci) / 2
  
  boot_mat  <- matrix(NA_real_, nrow = n_boot, ncol = length(x_bins))
  
  cat(sprintf("  %s (%d transitions)...\n", label, length(trans_ids)))
  pb <- txtProgressBar(min = 0, max = n_boot, style = 3)
  
  for (b in seq_len(n_boot)) {
    #Draw transitions with replacement
    samp_ids  <- sample(trans_ids, length(trans_ids), replace = TRUE)
    samp_df   <- df_type %>% filter(transition_id %in% samp_ids)
    bin_means <- samp_df %>%
      group_by(x_bin) %>%
      summarise(m = mean(force_renorm, na.rm = TRUE), .groups = "drop")
    idx <- match(bin_means$x_bin, x_bins)
    boot_mat[b, idx] <- bin_means$m
    setTxtProgressBar(pb, b)
  }
  close(pb)
  
  data.frame(
    x_bin      = x_bins,
    mean_force = colMeans(boot_mat, na.rm = TRUE),
    ci_lo      = apply(boot_mat, 2, quantile, probs = alpha,     na.rm = TRUE),
    ci_hi      = apply(boot_mat, 2, quantile, probs = 1 - alpha, na.rm = TRUE),
    n_trans    = colSums(!is.na(boot_mat))
  )
}

ext_binned <- trans_df_binned %>% filter(transition_type == "extinction")

ext_boot <- boot_ci(ext_binned,       label = "Extinction")    %>%
  mutate(transition_type = "extinction")
rr_boot  <- boot_ci(rr_source_binned, label = "Reward-reward") %>%
  mutate(transition_type = "reward-reward")

boot_df <- bind_rows(ext_boot, rr_boot)


#Plot continuous force dynamics
#
#Inset: histogram + rug of the normalized x-position of peak rolling-average
#force for each extinction transition. Illustrates that
#individual peaks are spread across the post-onset window and do not cluster
#tightly at x = 0, cautioning against interpreting the group average as
#implying synchrony of individual peaks. When multiple x-bins share the
#maximum force_renorm within a transition, the first occurrence is used.

type_colors <- c(extinction      = "#8B008B",
                 "reward-reward" = "#228B22")
type_labels <- c(extinction      = "Reward-Extinction",
                 "reward-reward" = "Reward-Reward")

#Peak x-position per extinction transition (post-onset only: x >= 0)
peak_locs <- trans_df %>%
  filter(transition_type == "extinction", x_norm >= 0) %>%
  group_by(transition_id) %>%
  slice(which.max(force_renorm)) %>%   #first occurrence if tied
  ungroup() %>%
  pull(x_norm)
print(sort(peak_locs))

#Inset: histogram with rug, spanning the full post-onset extinction window
set.seed(1401)
p_inset <- ggplot(data.frame(x = peak_locs), aes(x = x)) +
  geom_histogram(binwidth = 0.1, fill = "#4B0050", color = "gray",
                 alpha = 1, boundary = 0) +
  geom_rug(data = data.frame(x = jitter(peak_locs, amount = 0.0005)),
           aes(x = x), color = "#4B0050", alpha = 0.5, linewidth = 0.6, length = unit(0.035, "npc")) +
  geom_vline(xintercept = 1, linetype = "dotted",
             color = "gray30", linewidth = 0.4) +
  annotate("text", x =  0.65, y = 9, label = "Early ext.",
           vjust = 1.5, size = 2, color = "gray40") +
  annotate("text", x =  1.35, y = 9, label = "Late ext.",
           vjust = 1.5, size = 2, color = "gray40") +
  scale_x_continuous(limits = c(0, 2.05),
                     breaks = c(0, .25, .5, .75, 1, 1.25, 1.5, 1.75, 2),
                     labels = scales::number_format(drop0trailing = TRUE)) +
  scale_y_continuous(breaks = scales::breaks_pretty(n = 4),
                     labels = scales::number_format(accuracy = 1)) +
  labs(x = "Peak location", y = "Count") +
  theme_classic(base_size = 7) +
  theme(axis.title      = element_text(size = 6),
        axis.text       = element_text(size = 5),
        panel.background = element_rect(fill = "white"),
        plot.background = element_rect(fill = NA, color = "NA",
                                       linewidth = 0.3))

#Main figure
p_main <- ggplot(boot_df,
                 aes(x = x_bin, y = mean_force,
                     color = transition_type, fill = transition_type)) +
  geom_ribbon(aes(ymin = ci_lo, ymax = ci_hi), alpha = 0.25, color = NA) +
  geom_line(linewidth = 0.9) +
  geom_vline(xintercept = 0, linetype = "dashed",
             color = "gray30", linewidth = 0.4) +
  geom_vline(xintercept = 1, linetype = "dashed",
             color = "gray30", linewidth = 0.4) +
  geom_hline(yintercept = 0, linetype = "dotted",
             color = "black",  linewidth = 0.3) +
  annotate("text", x = -0.425, y = -0.22, label = "Pre-transition",
           vjust = 1.5, size = 3, color = "gray40") +
  annotate("text", x =  0.5, y = -0.22, label = "Post-transition",
           vjust = 1.5, size = 3, color = "gray40") +
  annotate("text", x =  1.5, y = -0.22, label = "Late extinction",
           vjust = 1.5, size = 3, color = "gray40") +
  annotation_custom(
    grob = ggplotGrob(p_inset),
    xmin = 0.50, xmax = 2.15,
    ymin = 0.00, ymax = 0.185
  ) +
  scale_color_manual(values = type_colors, labels = type_labels, name = "Transition type") +
  scale_fill_manual( values = type_colors, labels = type_labels, name = "Transition type") +
  labs(x = "Normalized time (pre-transition block = 1 unit)",
       y = "Continuous force") +
  theme_classic(base_size = 12) +
  theme(legend.position  = c(0.15, 0.88),
        legend.background = element_rect(fill = NA, color = NA),
        legend.key.size = unit(0.35, "cm"),
        legend.title = element_text(size = 8),
        legend.text  = element_text(size = 7))

print(p_main)
#save at 500x400

#Plots all individual transition traces, faceted by participant.
#Useful for identifying individual patterns.
p_per_part <- ggplot(
  trans_df,
  aes(x = x_norm, y = force_renorm,
      group = transition_id, color = transition_type)
) +
  geom_line(linewidth = 0.35, alpha = 0.75) +
  geom_vline(xintercept = 0, linetype = "dashed",
             color = "gray30", linewidth = 0.3) +
  geom_hline(yintercept = 0, linetype = "dotted",
             color = "black",  linewidth = 0.25) +
  facet_wrap(~ pid + group, scales = "free_y", labeller = label_both) +
  scale_color_manual(values = type_colors, labels = type_labels, name = NULL) +
  labs(x = "Normalized time",
       y = "Re-standardized force") +
  theme_classic(base_size = 7) +
  theme(
    legend.position = "top",
    strip.text      = element_text(size = 5),
    axis.text       = element_text(size = 4)
  )

print(p_per_part)
#This figure is large, so open in a full window or export at high resolution
```
