## Supplementary material for "Evidence for the extinction burst in grip force: A preregistered study": Electronic Supplementary Materials: README.txt

This folder contains R scripts for the complete data-analysis pipeline, including power analysis, data processing, statistical analyses, and generation of the figures reported in the manuscript.

To run the pipeline:

*Open R (or RStudio).

*Set ebf_raw_data as your working directory.

*Run the scripts in the order indicated by their filenames.
