## Supplementary figures and images for "Evidence for the extinction burst in grip force: A preregistered study"

### 5pesos.png

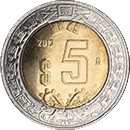

### consentimiento.jpg

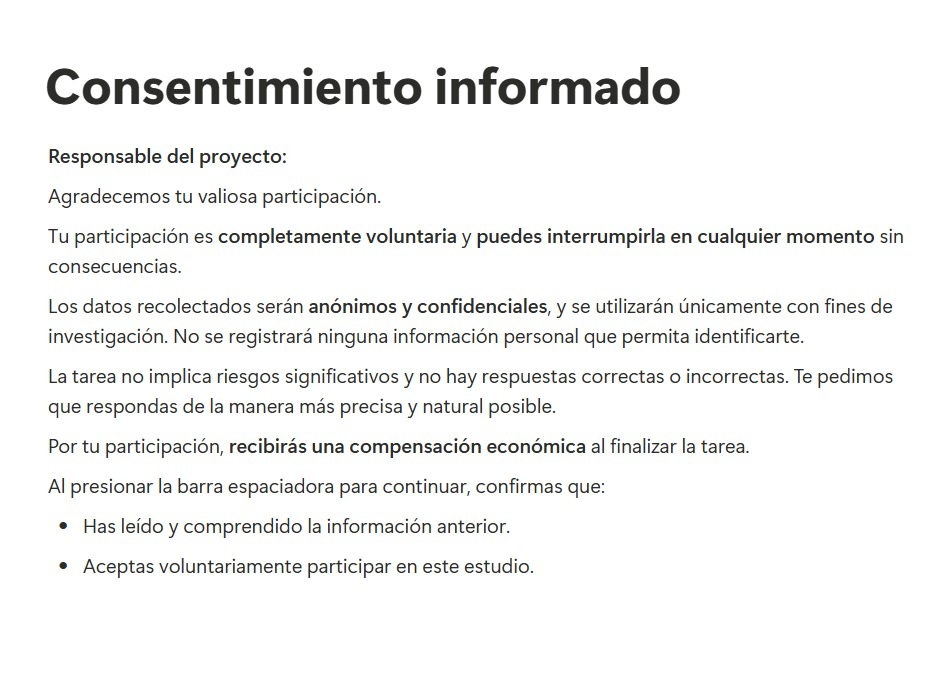

### IMG-20250925-WA0005.jpg

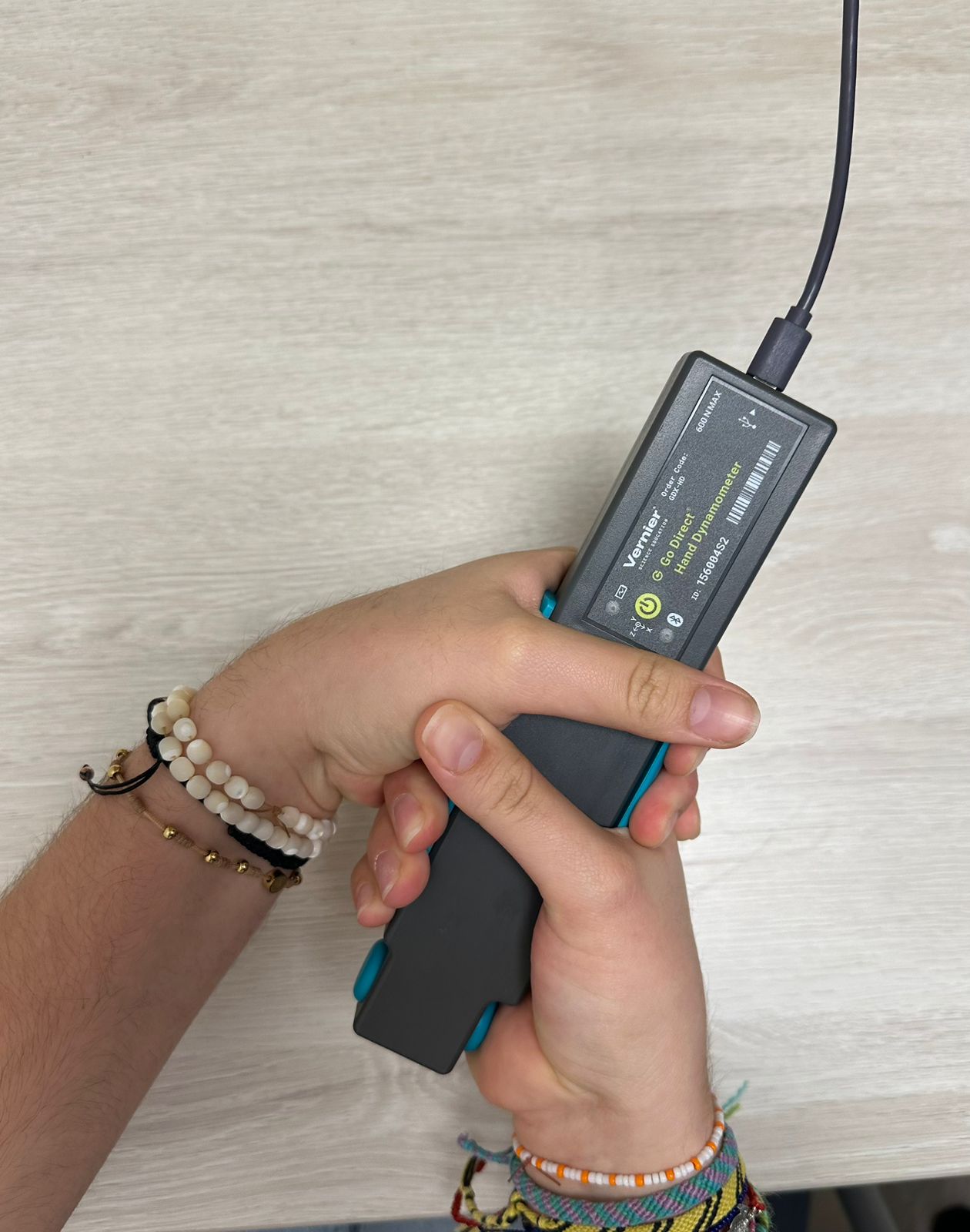

### IMG-20250925-WA0006.jpg

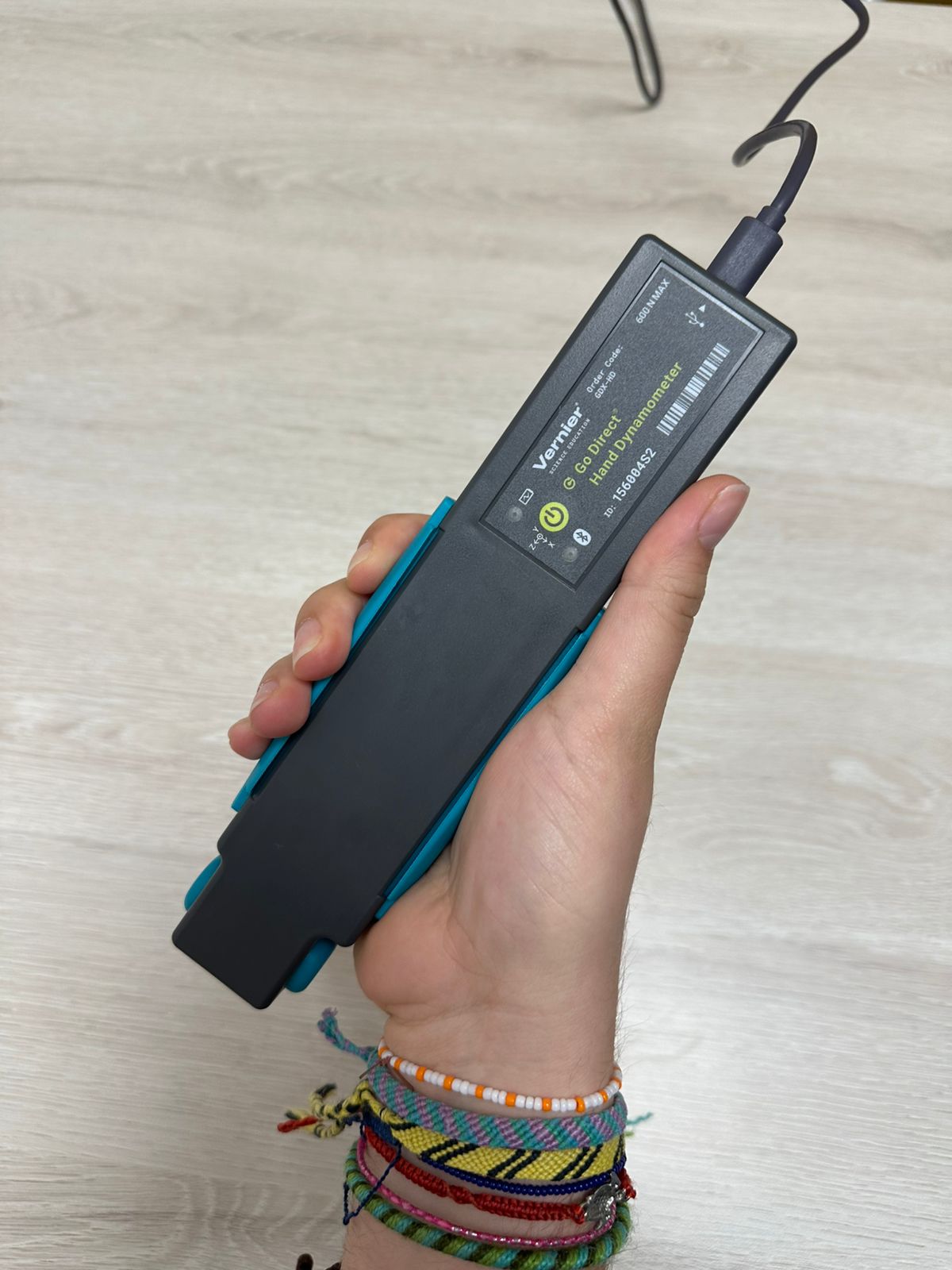

### IMG-20250925-WA0008.jpg

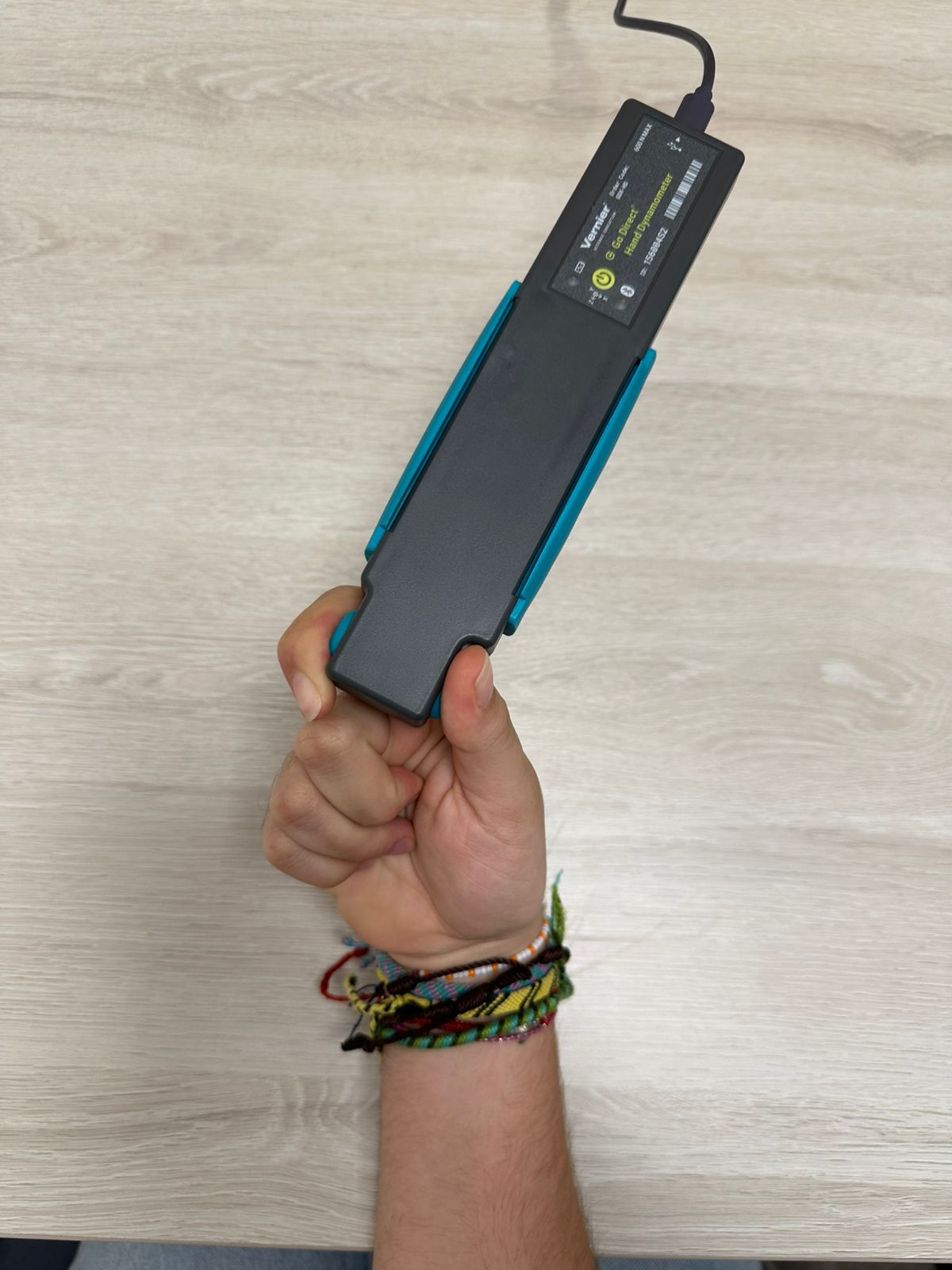

### IMG-20250925-WA0009.jpg

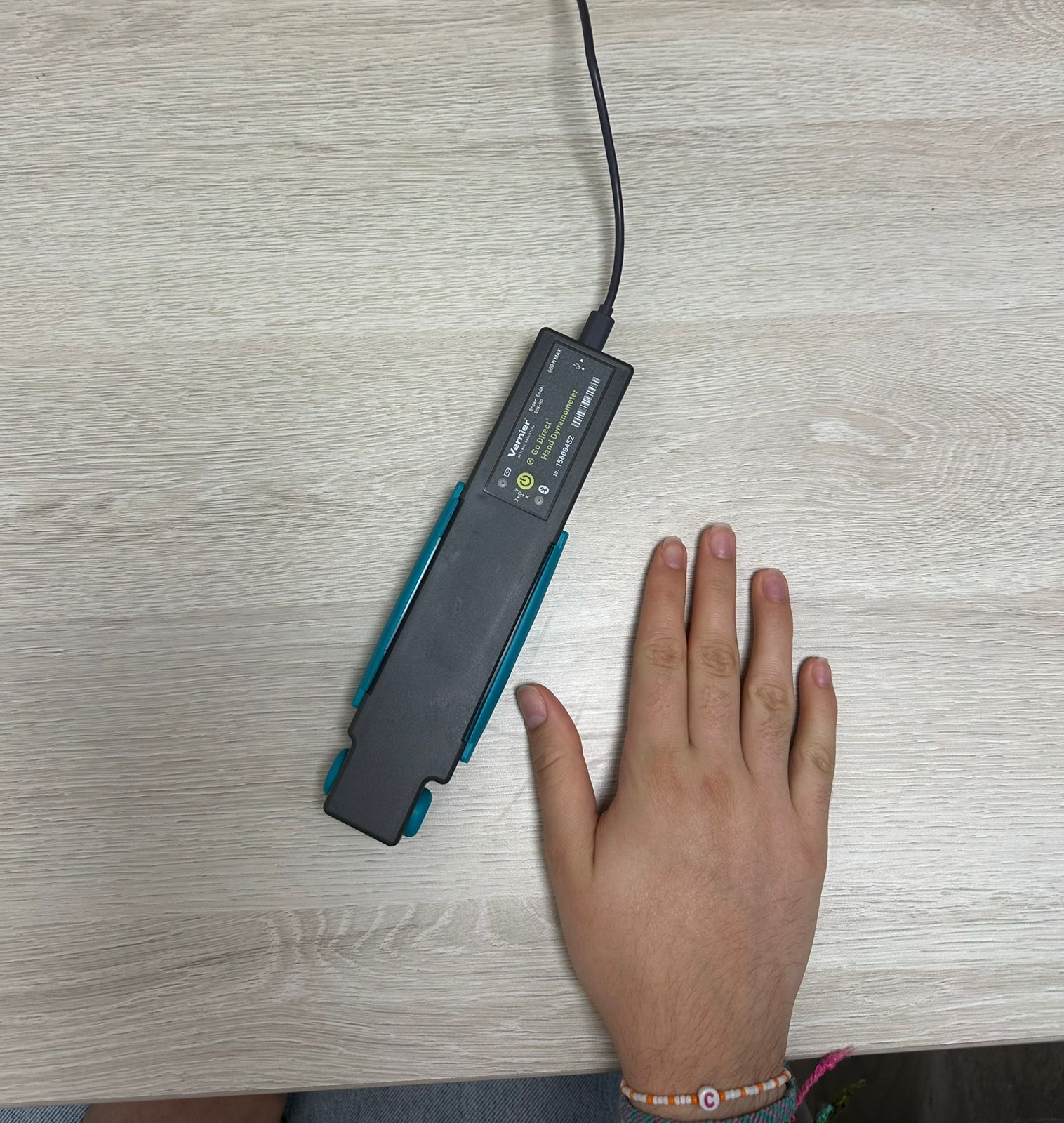

### palomita agarrar.png

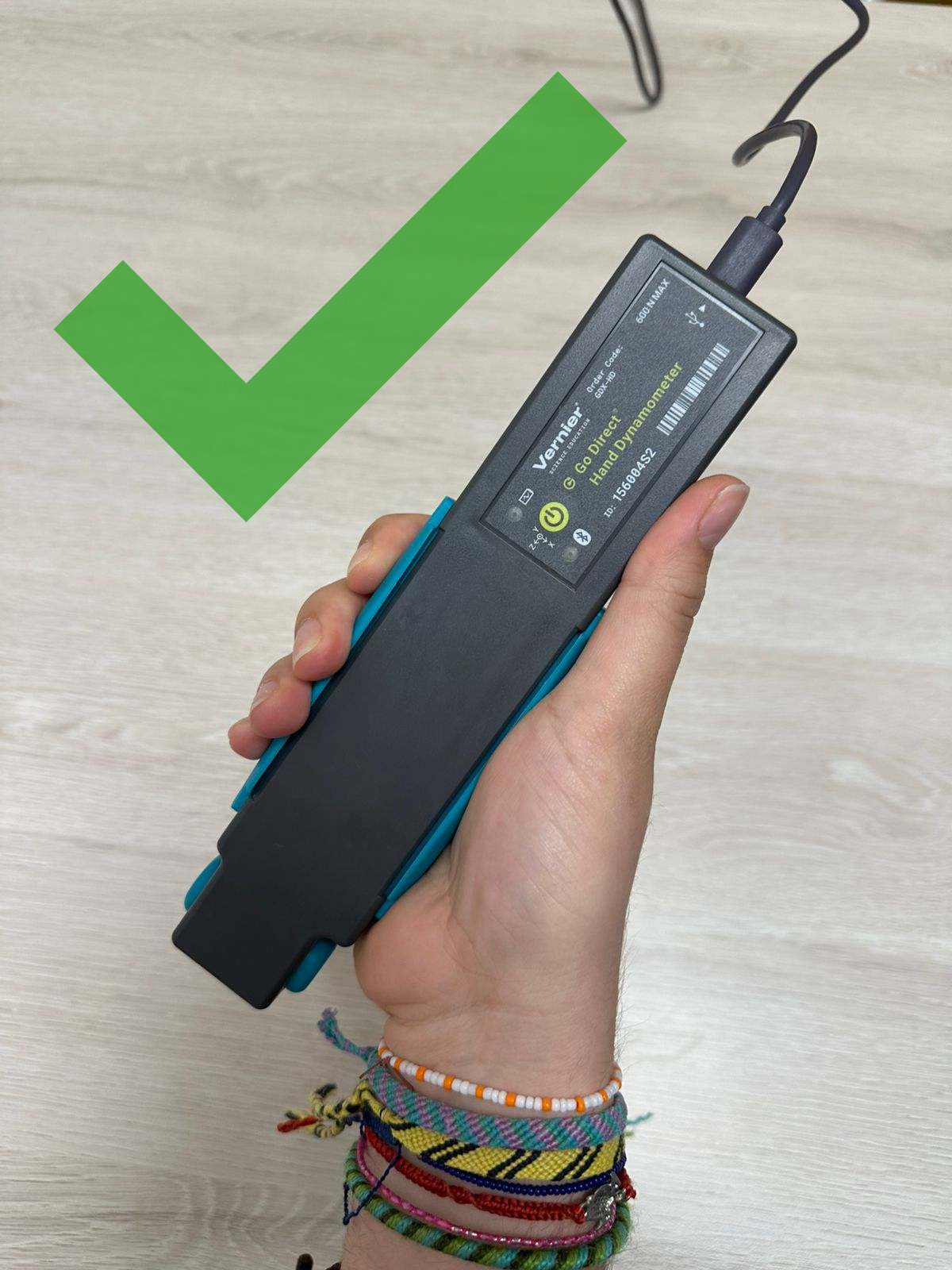

### tache circulos.png

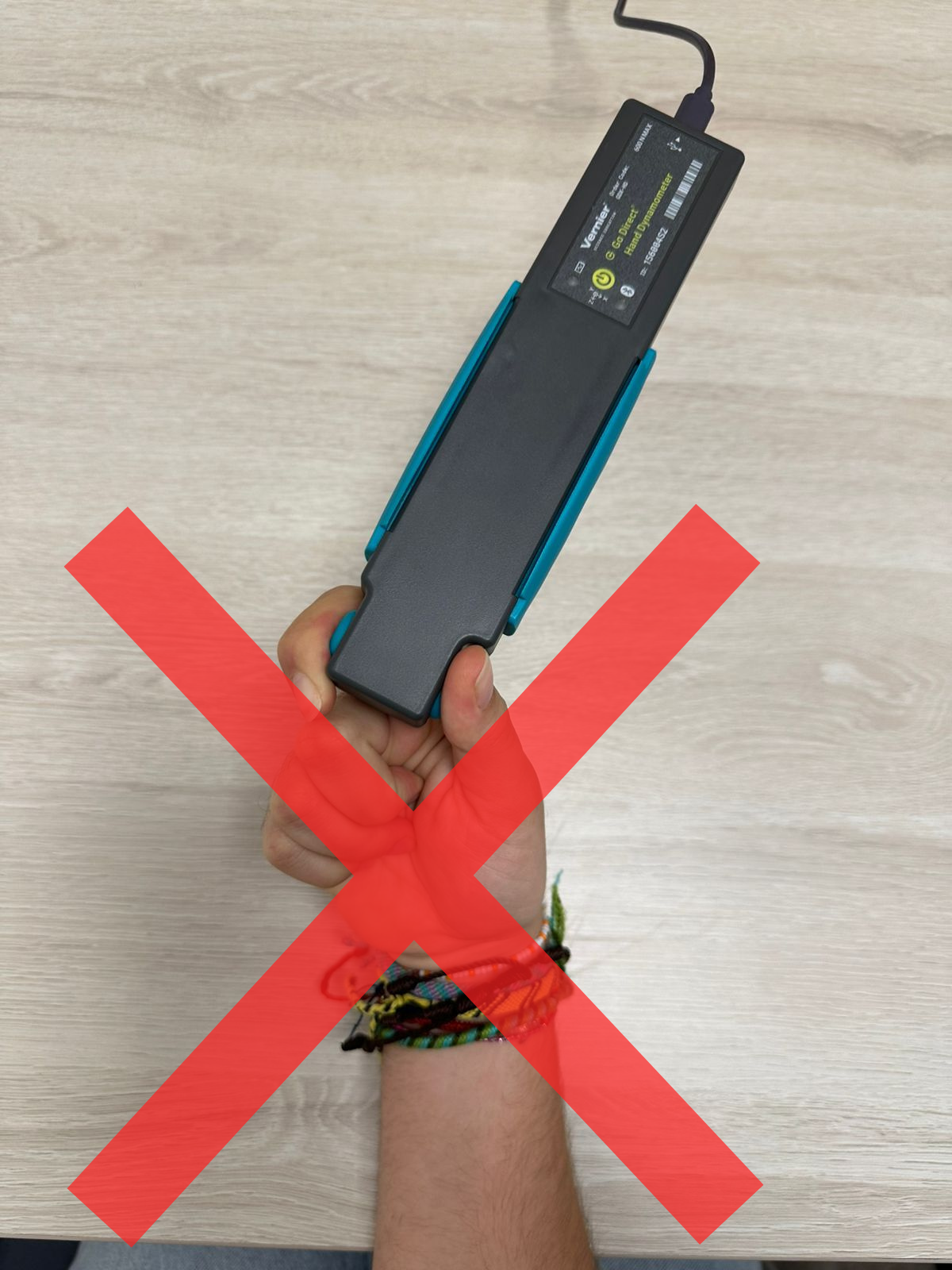

### tache dos manos.png

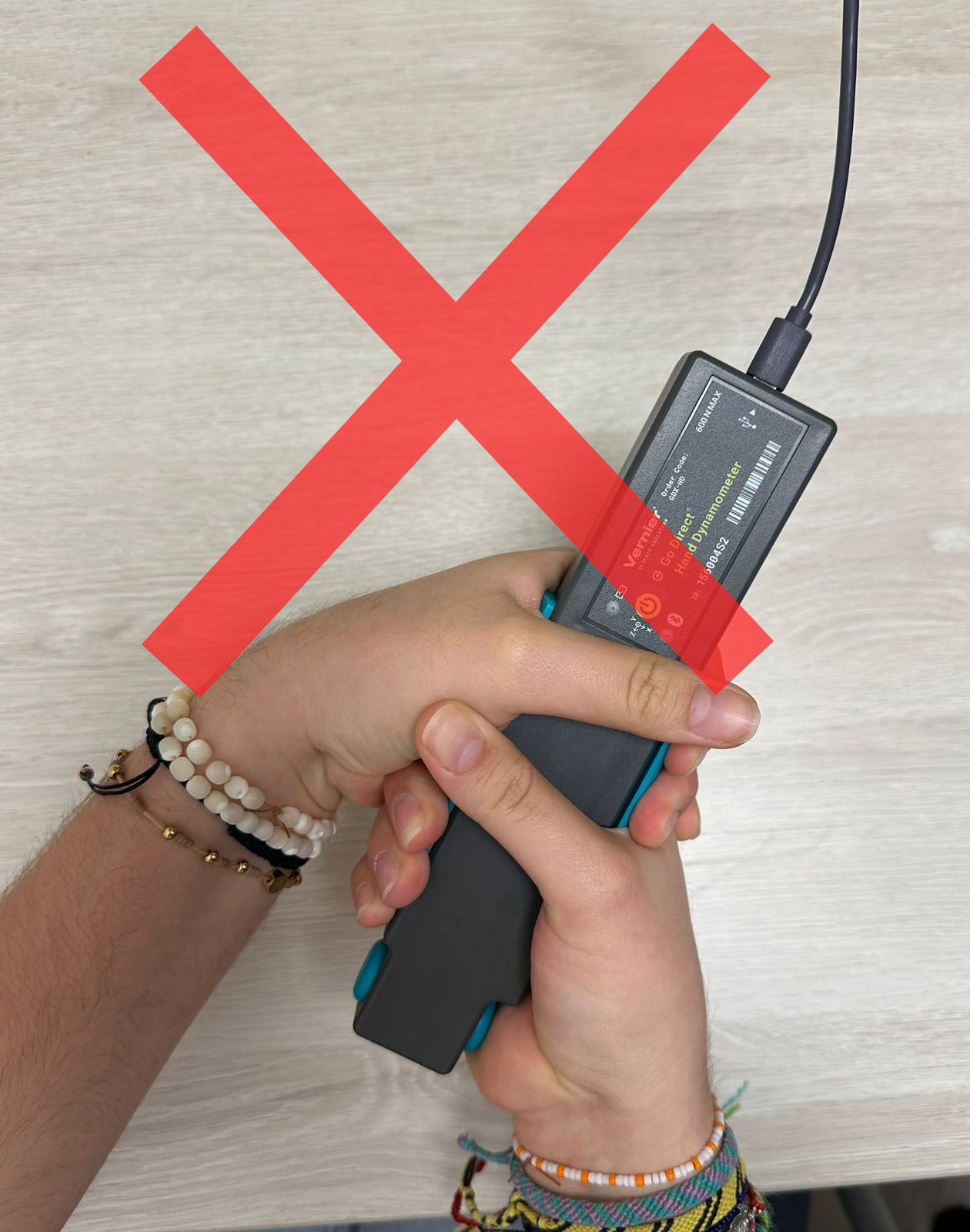

### tache soltar.png

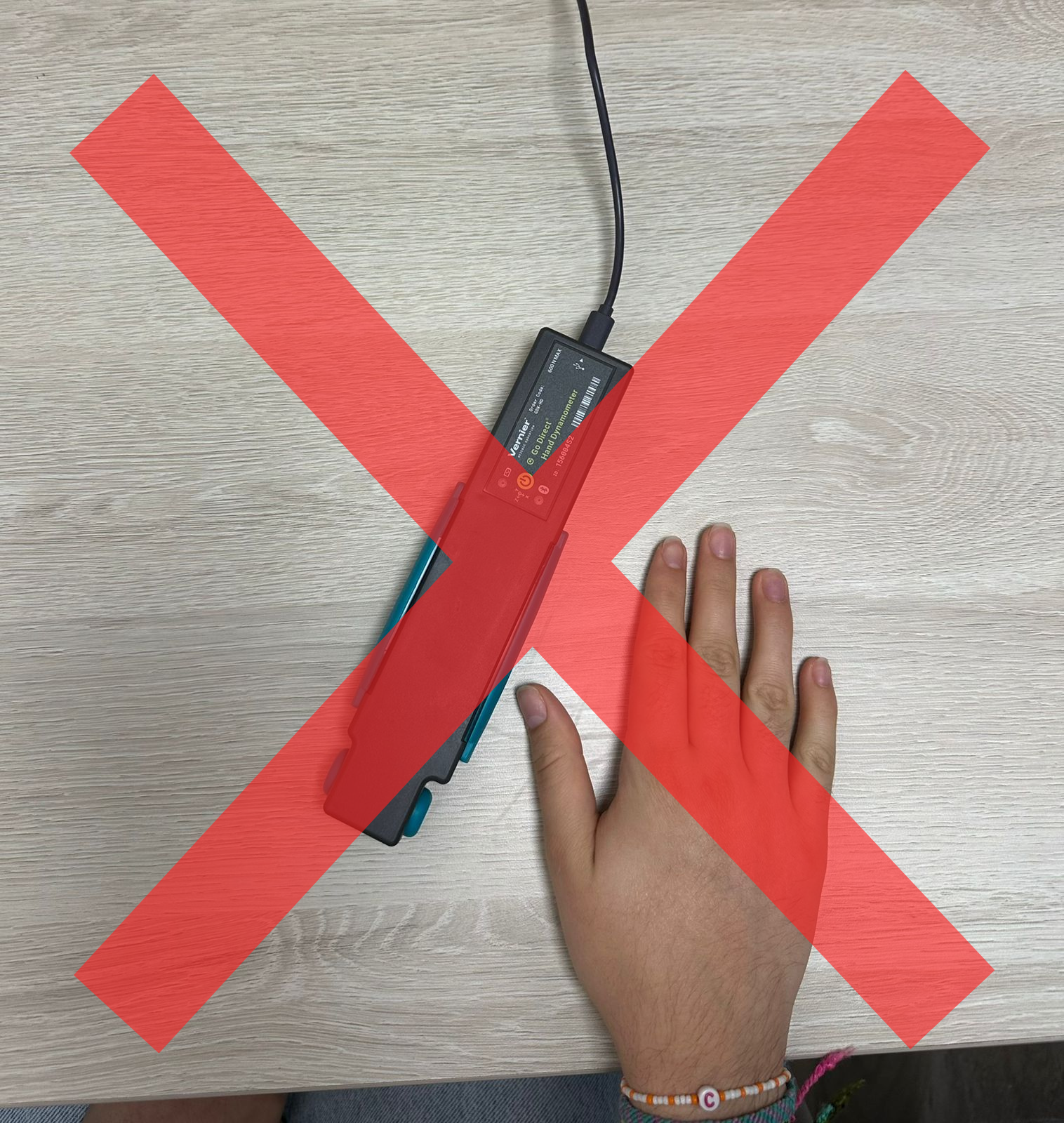
